# Drug screens of NGLY1 Deficiency worm and fly models reveal catecholamine, NRF2 and anti-inflammatory pathway activation as potential clinical approaches

**DOI:** 10.1101/563478

**Authors:** Sangeetha Iyer, Joshua D. Mast, Hillary Tsang, Tamy P. Rodriguez, Nina DiPrimio, Madeleine Prangley, Feba S. Sam, Zachary Parton, Ethan O. Perlstein

**Affiliations:** Perlara PBC, 6000 Shoreline Court, South San Francisco, California USA

**Keywords:** *N*-glycanase 1 Deficiency, *NGLY1*, *SKN-1*, *Pngl*, congenital disorder of deglycosylation, disease model

## Abstract

*N*-glycanase 1/*NGLY1* Deficiency is an ultra-rare and complex monogenic glycosylation disorder that affects fewer than 40 patients globally. *NGLY1* Deficiency has been studied in model organisms such as yeast, worms, flies and mice. Proteasomal and mitochondrial homeostasis gene networks are controlled by the evolutionarily conserved transcriptional regulator Nrf1, whose activity requires deglycosylation by NGLY1. Hypersensitivity to the proteasome inhibitor bortezomib is a common phenotype observed in whole animal and cellular models of *NGLY1* Deficiency. Here we describe unbiased phenotypic drug screens to identify FDA approved drugs, generally recognized as safe natural products and novel chemical entities that rescue growth and development of *NGLY1*-deficient worm and fly larvae treated with a toxic dose of bortezomib. We used image-based larval size and number assays for use in screens of a 2,560-member drug repurposing library and a 20,240-member lead discovery library. A total of 91 validated hit compounds from primary invertebrate screens were tested in a human cell line in a NRF2 activity assay. NRF2 is a transcriptional regulator that regulates cellular redox homeostasis and it can compensate for loss of Nrf1. Plant-based polyphenols comprise the largest class of hit compounds and NRF2 inducers. Catecholamines and catecholamine receptor activators comprise the second largest class of hits. Steroidal and non-steroidal anti-inflammatory drugs comprise the third largest class. Only one compound was active in all assays and species: the atypical antipsychotic and dopamine receptor agonist aripiprazole. Worm and fly models of *NGLY1* Deficiency validate therapeutic rationales for activation of NRF2 and anti-inflammatory pathways based on results in mice and human cell models and suggest a novel therapeutic rationale for boosting catecholamine levels and/or signaling in the brain.

## Introduction

NGLY1 Deficiency is the first congenital disorder of deglycosylation (CDDG) described in the biomedical literature (Lam et al., 2017; Enns et al., 2014). NGLY1 Deficiency has multi-organ presentation and clinical features in patients such as global developmental delay, a complex hyperkinetic movement disorder, small body size, seizures, and alacrimia. *NGLY1* is an ancient gene encoding a cytosolic enzyme called *N*-glycanase 1 – also referred to as PNGase – which catalyzes the hydrolysis and release of *N*-glycans from *N*-glycosylated proteins (Suzuki et al., 2016). NGLY1 is thought to function primarily in an evolutionarily conserved protein surveillance and disposal pathway called ERAD, or endoplasmic reticulum-associated degradation (Suzuki et al., 2015). NGLY1 also regulates the activity of specific glycoproteins by using deglycosylation as a post-translational on/off switch. Cellular models of NGLY1 Deficiency have shown that the transcriptional regulator Nrf1 is a specific deglycosylation target of NGLY1 and that knocking out *NGLY1* phenocopies knocking out *Nrf1* (Tomlin el al., 2017). Only deglycosylated Nrf1 can be proteolytically processed into the mature nuclear-active form. Once in the nucleus, Nrf1 controls the expression of proteasomal subunit genes in response to protein folding stress in mammalian cells (Radhakrishnan et al., 2014), worms (Lehrbach & Ruvkun, 2016) and flies (Grimberg et al., 2011). In flies NGLY1 regulates the glycosylation status of the ortholog of a bone morphogenetic protein (BMP) signaling ligand (Galeone et al., 2017). Demonstrating the complexity of how loss-of-function mutations in the gene lead to pathophysiology in humans, NGLY1 regulates mitochondrial physiology in human and mouse fibroblasts and in worms through mechanisms that are still under investigation (Kong et al. 2018). Interestingly, mitophagy defects caused by loss of Nrf1 function can be rescued by activation of the related transcriptional regulator NRF2, which controls the expression of genes involved in antioxidant and redox stress responses (Yang et al., 2018).

In the five years since the publication of the first *NGLY1* Deficiency diagnostic cohort of eight patients (Enns et al., 2014) multiple research groups have contributed to our understanding of disease-causing and loss-of-function mutations in the *NGLY1* gene and its orthologs by generating and characterizing small and large animal models as well as patient-derived cell models. From this marketplace of disease models, a common phenotype emerged: hypersensitivity to proteasome inhibition by bortezomib (Fenteany et al., 1995). In worms, hypersensitivity to bortezomib toxicity was observed in an otherwise normally developing *PNG-1/NGLY1* null mutant, which has a half-maximal growth inhibitory concentration (IC_50_) several hundred times lower than wildtype worms (Lehrbach & Ruvkun, 2016). Using a chemically related proteasome inhibitor lacking the reactive boronic acid group, it was shown that mouse embryonic fibroblasts derived from *NGLY1*-knockout mice and *NGLY1*-knockdown human cell lines are several-fold more sensitive to carfilzomib toxicity compared to controls (Tomlin et al., 2017).

In an effort to phenotype and screen a homozygous loss-of-function *Pngl/NGLY1* fly modeling the patient-derived C-terminal premature stop codon allele R401X, we showed that *Pngl*^−/−^ homozygous mutant larvae are 25-fold more sensitive than heterozygotes to the toxic effects of bortezomib (Rodriguez et al., 2018). Another group reported that a *Pngl* RNAi-knockdown fly model of *NGLY1* Deficiency has constitutively reduced expression of Nrf1-dependent proteasomal subunit genes, consistent with findings of hypersensitivity to bortezomib toxicity in the other models (Owings et al., 2018). Loss of NGLY1 causes intolerance to bortezomib that is as evolutionarily conserved as the underlying Nrf1-dependent proteasome bounce-back response because they go hand in hand. The prediction that has been confirmed so far in mammalian cells and in nematodes (Lehrbach et al., 2019) is that NGLY1 and its orthologs deglycosylate Nrf1 and its orthologs. We reason that small molecule suppressors of bortezomib will safely activate bypass pathways that rescue or compensate for loss of NGLY1 in a whole animal and will have a higher probability of exhibiting a favorable therapeutic index in mammals.

We used our *Pngl*^−/−^ fly model in a drug repurposing screen to identify compounds that rescue larval developmental delay (Rodriguez et al., 2018). Homozygote flies fail to thrive in the absence of any exogenous stressor or enhancer such as bortezomib. There are two limitations of our previous screen. First, flies homozygous for a patient-derived nonsense allele are extremely sick and only one fly-specific validated hit was identified – the insect molting hormone 20-hydroxyecdysone. Second, flies homozygous for a patient-derived nonsense allele are further sickened by the organic solvent dimethyl sulfoxide (DMSO) in which all test compounds are solubilized, and all stock solutions prepared. Therefore, drug screens were conducted at the limit of detection and a high false negative rate was due to under-dosing of test compounds.

Here we addressed the shortcomings of the initial fly-only drug repurposing campaign. First, we discovered that the *Pngl*^+/-^ heterozygote is two-fold more sensitive to bortezomib toxicity than wildtype animals, it tolerates DMSO, and it has no detectable developmental delay. We hypothesize that it will be easier for a small molecule to suppress larval developmental delay in bortezomib-treated healthy heterozygotes versus constitutively sick homozygotes because otherwise healthy heterozygotes tolerate higher levels DMSO and thus allow for all test compounds to be screened at a five-fold higher concentration. Second, we extended the bortezomib intolerance results of Lehrbach & Ruvkun to a liquid-based, 384-well-plate quantitative worm larval growth and development assay for drug screening. We hypothesize that a two-species bortezomib-suppressor screen conducted in parallel would reveal hit compounds that target evolutionarily conserved disease modifiers and disease-modifying pathways. Third, we not only cross-tested all hit compounds from the primary worm screen in the fly assay, and vice versa, but we also cross-tested all hit compounds in two different worm assay paradigms (bortezomib treatment versus carfilzomib treatment), in two different fly assay paradigms (bortezomib treatment of heterozygotes versus homozygotes), and in a human cell NRF2 transcriptional activity reporter assay. This cross-validation scheme generated a short list of generic drugs and nutriceuticals in which we had the highest degree of confidence of reproducibility and clinical trial actionability. Finally, we screened not only the same repurposing library as described in Rodriguez *et al* but also a ten-fold larger lead discovery library. We hypothesize that in some instances the same known mechanisms of action of repurposable drugs will be targeted by novel chemical entities, which serve as starting points for lead optimization toward a best-in-class clinical candidate.

## Materials and Methods

### Strains and compound libraries

The *png-1* deletion mutant ok1654 was previously described (Lehrback & Ruvkun, 2016) and is available from the CGC Stock Center (strain ID: RB1452). The nonsense allele *Pngl* fly was previously described (Rodriguez et al., 2018) and is available from the Bloomington Stock Center. The 2,560-compound Microsource Spectrum Collection was available for purchase before the distributor went out of business. The 20,240-compound lead discovery library was purchased from Chembridge Corporation. All compounds were dissolved in DMSO and stored at −80°C in Labcyte-compatible LDV plates until use.

#### High-throughput larval growth assays in worms

Positive controls for all screens are *png-1* mutants + DMSO (vehicle control) and the negative controls are *png-1* mutants + DMSO + 205nM bortezomib. Both drug libraries were screened in triplicate with controls on each 384-well drug screening plate. Using the Echo550 (Labcyte Inc.), Bortezomib was acoustically dispensed into the destination plates the day before the worm larvae sort. 5μL of HB101 bacteria were dispensed into 384-well plates, containing S Medium in each well. Using the BioSorter (Union Biometrica), 15 L1 *png-1* mutant larvae were sorted into each well, and plates were incubated at 20°C while shaking. After five days of incubation, 15μL of 8mM sodium azide was added to each well to immobilize worms prior to imaging in a custom worm imager. Finally, automated image processing was run on each plate.

#### High-throughput fly larval growth assays

Drug screens of *Pngl* heterozygous flies were conducted in the presence of 9μM bortezomib. The negative controls in the screen are *Pngl* heterozygous flies + DMSO + bortezomib and the positive controls are *Pngl* heterozygous flies + DMSO. Both drug libraries were screened in triplicate. Using the Echo550 (Labcyte Inc.), test compound was acoustically dispensed and standard fly food media (molasses, agar, yeast, propionic acid) lacking cornmeal, but carrying 0.025% bromophenol blue was dispensed using a Multi-Flo (Bio-Tek Instruments) into each well of a 96-well plate for drug screens or a 12-well plate for hit retests. Then, the BioSorter was used to dispense *Pngl* heterozygous larvae, three per well. At three days post incubation with the compounds, the plates were scored for larval size rescue in a custom fly imager.

#### High-throughput drug screening and analysis

Hit compounds rescued worm development, i.e., increased the number of worms (total area taken up by worms) in the well. Therefore, the output of the image processing was worm area per well. Each plate contained 32 wells of positive controls (worms raised in the absence of bortezomib) and 32 wells of negative controls (worms raised in the presence of bortezomib). The average area occupied by worms in positive and negative control wells as well standard deviation of control wells were calculated. Outlier elimination was performed by identifying those wells that were greater than one and half times the interquartile range (1.5 X IQR). Z scores for test wells were calculated by normalizing by mean area standard deviation of negative control wells. For each replicate a test well that had a Z score ≥ 2 was counted as a hit. Quality control was performed to ensure that the control well grew as expected, the *png-1* mutants are sensitive to bortezomib as expected, there are no obvious plate effects, and dispense errors were eliminated. Because image artifacts can often confound area measurements, we also manually verified each well by visual analysis after the unbiased quantitative analysis to make sure no false positives were counted as true hits. We count wells as hits that have a Z-score greater than 2 across all three replicates of the screen and were free of image artifacts.

Fly screening conditions were previously described (Rodriguez et al., 2018). Briefly, we imaged the plates using a custom fly imager and analysis tool to determine the area the larvae occupy per well. Then, we counted the number of larvae per well to normalize the total area per well data and identified suppressors of the phenotype. We also manually scored the wells for contaminants that may give false positives. Similar to the statistical analysis for the nematode assay, test wells were assigned Z scores relative to mean negative controls. Any well that had Z score > 2.5 in at least two out of three replicates was counted as a hit.

#### Keap1-NRF2 activation luciferase reporter assay in U2OS cells

We tested compounds using the PathHunter^®^ eXpress Keap1-NRF2 Nuclear Translocation Assay (DiscoverX, San Diego). Briefly, PathHunter cells are plated and incubated for 24 hours at 37°C. 10μL of test compound was added and cells were incubated with compound for 6 hours at room temperature. Working detection reagent solution was added and plates were incubated for 60 minutes at room temperature. Chemiluminescence signal was read by a SpectraMax M3.

### Results

There are currently no FDA approved treatments for NGLY1 Deficiency despite the unmet medical need. Drug repurposing involves finding new uses for old drugs and is the shortest path to a therapy for ultra-rare disease communities with limited financial resources and few dedicated researchers (Pushpakom et al., 2019). We used a model-organism-based disease modeling and phenotypic drug screening approach, which is enabling precision medicine to bridge bench to bedside (Li et al., 2019).

#### Developing high-throughput bortezomib-modifier assays for nematode and fly larvae

The nematode ortholog of *NGLY1* is *PNG-1*. The strain used here is ok1654, which contains an 800-base-pair deletion in the *PNG-1* open reading frame resulting in a null mutant. We confirmed that ok1654 does not have an intrinsic growth defect but is markedly hypersensitive to bortezomib toxicity (**Figure 1A**). Bortezomib exacerbates the proteasomal stress that *NGLY1*-deficient worms are already experiencing because of the concomitant loss of Nrf1 activity and results in an exaggerated disease phenotype, in this case toxicity in the form of early larval growth arrest (Lehrbach & Ruvkun, 2016). Because *png-1* homozygous mutant worms do not have a constitutive growth or developmental defect, we decided to use bortezomib to induce larval arrest and screen for compounds that restore normal growth and development as measured by the size and number of worms in each well of a 384-well plate. From the bortezomib dose-response data we established that a *png-1* null mutant is sensitive to bortezomib toxicity down to the low nanomolar range. After further assay optimization studies, we selected 205nM bortezomib as the concentration for the primary screens and for secondary retest and cross-test experiments to validate hit compounds.

**Figure 1.**
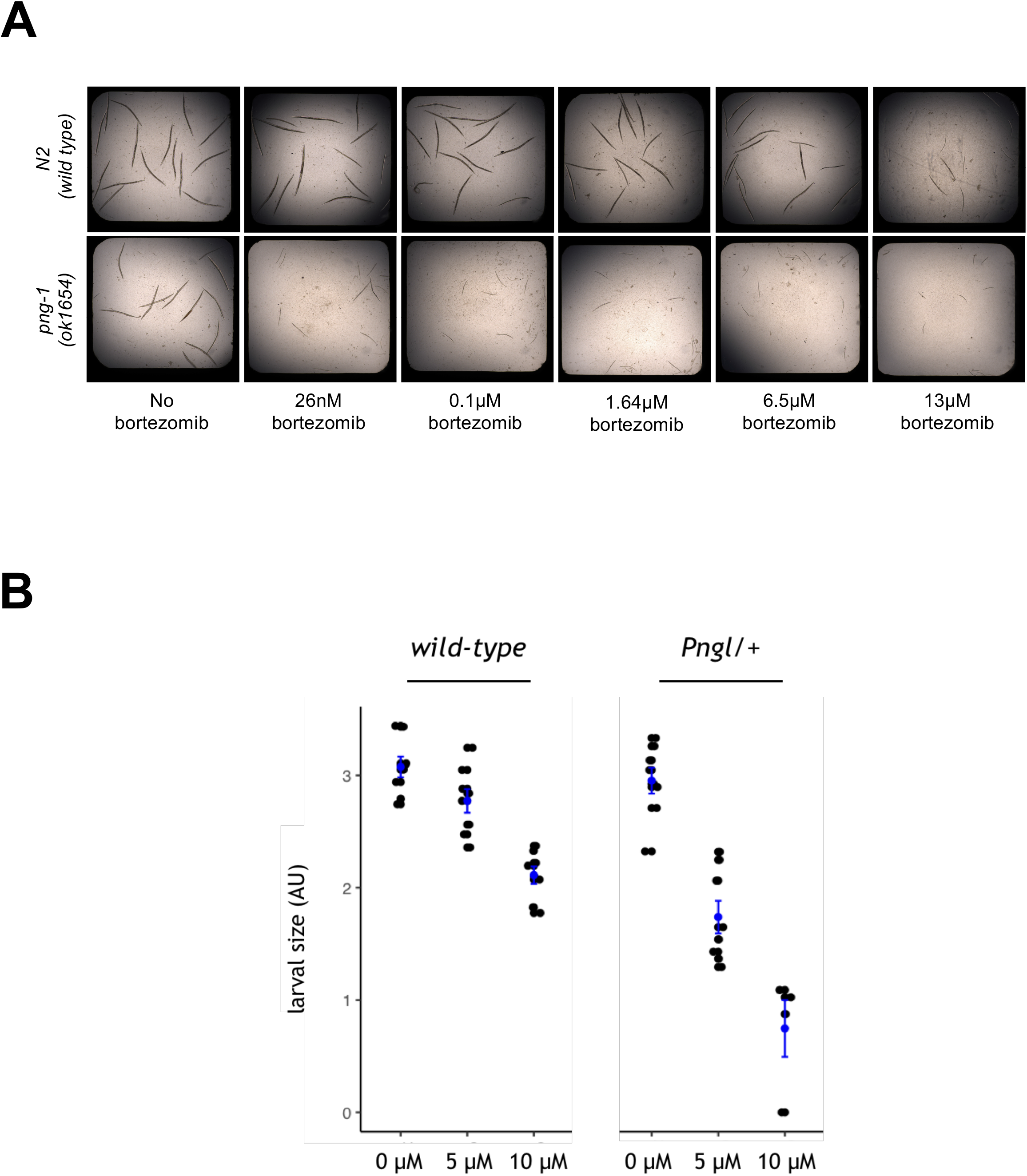
Determining a half-maximal effective concentration (EC_50_) for bortezomib in NGLY1 deficient worms and flies. **(A)** Wildtype N2 worms (top row) and *png-1* homozygous mutant worms (bottom row) were grown in liquid media in the presence of increasing concentrations of bortezomib (right to left). **(B)** Wildtype and *Pngl*^+/-^ heterozygous mutant fly larvae were grown on solid media in the presence of either 5μM or 10μM bortezomib. Fly larvae were imaged and their sizes were plotted in arbitrary units (AU).

The ortholog of *NGLY1* in flies is *Pngl*. We previously developed a *Pngl*^−/−^ fly model based on a recurrent C-terminal nonsense allele found in NGLY1 patients, optimized a high-throughput larval size assay, and screened a 2,560-member drug repurposing library on *Pngl*^−/−^ larvae (Rodriguez *et al*, 2018). That effort culminated in a single validated hit compound, 20-hydroxyecdysone (20E). 20E is an insect-specific sterol-derived molting hormone. In order to identify clinically actionable hits, we increased the overall hit rate and robustness of fly screens. We generated bortezomib toxicity dose-response data for *Pngl*^+/-^ heterozygote larvae and determined that these animals are two-fold more sensitive than wildtype animals (**Figure 1B**). After further assay optimization studies with doses between 5μM and 10μM, we selected 9μM bortezomib as the concentration for the primary screens and for secondary retests and cross-tests to validate hit compounds.

#### Worm repurposing screen and hit validation

In the presence of 205nM bortezomib, *png-1* null mutant worms arrest as L1 larvae while wildtype animals treated with the same dose of bortezomib develop normally. A worm repurposing hit is defined as a compound that rescued bortezomib-treated *png-1* null worms such that their size and number were indistinguishable from control wells containing vehicle-treated *png-1* null worms. All test compounds were screened at a final concentration of 25μM. Images of a representative positive control well and a representative negative control well, and two examples of wells containing suppressors and enhancers/toxic compounds are shown in **Figure 2A**. The Venn diagram in **Figure 2B** summarizes the overlap of screening positives between three independent replicates. A total of 63 suppressors have Z-scores greater than two in all three replicates. The hit rate for worm suppressors is 63/2560 or 2.5%. A total of 51 enhancers have Z-scores less than two in all three replicates. The hit rate for enhancers/toxic compounds is 51/2560 or 2%. Enhancers/toxic compounds were not further investigated in this study. 60/63 worm suppressors were available for reorder as fresh powder stocks, retested in the primary bortezomib assay paradigm, and then scored in a secondary non-bortezomib assay paradigm.

**Figure 2.**
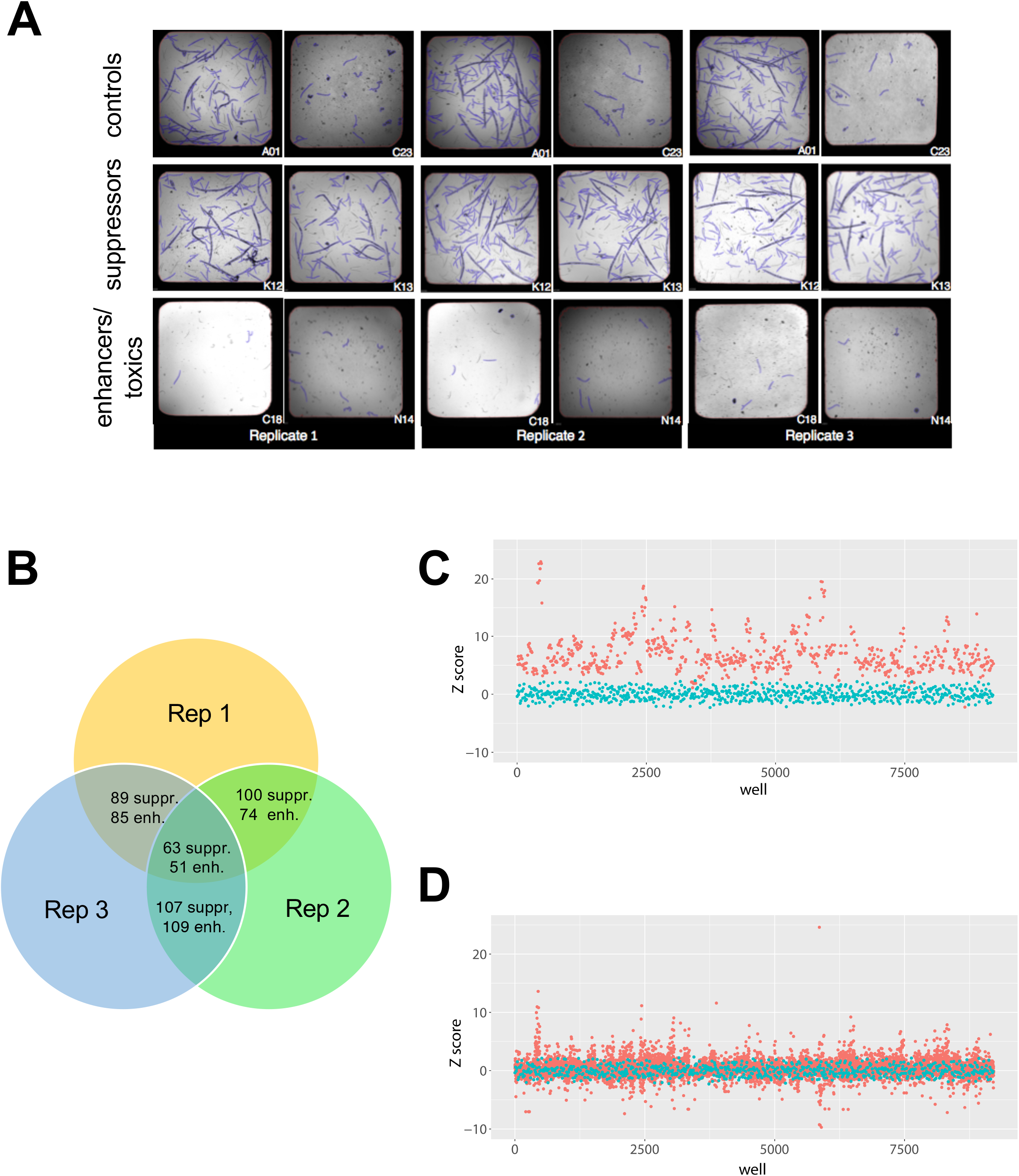
2,560-compound drug repurposing screens of *png-1* homozygous mutant worms and *Pngl* heterozygous mutant flies. **(A)** Worm screen images of a representative positive control well (A01), a representative negative control well (C23), two presumptive suppressors (K12, K13), and two presumptive enhancers/toxic compounds (C18, N14). Worms were pseudocolored blue by during image processing and analysis. **(B)** Venn diagram of overlapping hits from three replicate screens. **(C)** Z-score plot of three replicates of fly repurposing screen positive control (red circles) versus negative control (cyan circles) wells. **(D)** Z-score plot of three replicates of fly repurposing screen test compounds (red circles) versus negative (cyan circles) control wells.

Some compounds can directly inactivate bortezomib which leads to a false positive result. For example, polyphenols and in particular polyphenols containing multiple catechols are known to form covalent adducts with bortezomib by boronate-catechol complexation (Glynn et al., 2015; Golden et al., 2009). In order to filter out those compounds, we tested the 60 worm suppressors on the *png-1* null mutant treated with the chemically related proteasome inhibitor carfilzomib, which lacks the reactive boronic acid that renders bortezomib vulnerable to covalent attack. We established a carfilzomib dose-response curve for *png-1* null mutant worms (**Supplemental Figure 1**). We directly compared these data to worms treated with 205nM bortezomib in order to determine at which concentration we observe comparable larval growth arrest. We retested all 60 worm suppressors in the presence of 27.3μM carfilzomib, meaning carfilzomib is significantly less potent than bortezomib.

A total of 48/60 (80%) worm suppressors retested in the bortezomib assay paradigm. 15/60 (25%) worm suppressors scored positively in the carfilzomib assay paradigm. Overall, 15/60 (25%) worm suppressors rescued in both worm assay paradigms, i.e., in the presence of either bortezomib or carfilzomib. Those 15 compounds are the worm repurposing hits (**Table 1**): aripiprazole, benserazide, ellagic acid, epicatechin monogallate, epigallocatechin-3-monogallate, ethylnorepinephrine, gossypetin, koparin, phenylbutazone, pomiferin, purpurogallin-4-carboxylic acid, quercetin, theaflavin monogallate, triamcinolone, and 3,4-didesmethyl-5-deshydroxy-3’-ethoxyscleroin.

**Table 1.** Summary description of worm and fly repurposing hits

| Compound name | Pharmacological class | FDA approved? |
| --- | --- | --- |
| 3,4-didesmethyl-5-deshydroxy-3'-ethoxyscleroin | polyphenol | no |
| aminacrine | topical antiseptic | no |
| β-Amyrin | steroidal anti-inflammatory | no |
| aripiprazole | catecholamine receptor modulator | yes |
| benserazide | aromatic L-amino acid decarboxylase inhibitor | yes |
| butopyronoxyl | insect repellent | no |
| clindamycin | bacterial protein synthesis inhibitor (antibiotic) | yes |
| dyphylline | bronchodilator | yes |
| ellagic acid | polyphenol | no |
| epicatechin monogallate | polyphenol | no |
| epigallocatechin-3-monogallate | polyphenol | no |
| ethylnorepinephrine | adrenergic receptor agonist | yes |
| gossypetin | polyphenol | no |
| L-glutamine | amino acid supplement | no |
| ketorolac | nonsteroidal anti-inflammatory | yes |
| koparin | polyphenol | no |
| lumefantrine | antimalarial | yes |
| phenylbutazone | nonsteroidal anti-inflammatory | yes |
| pomiferin | polyphenol | no |
| promazine | phenothiazine antipsychotic | yes |
| purpurogallin-4-carboxylic acid | polyphenol | no |
| pyrimethamine | dihydrofolate reductase inhibitor (antimalarial) | yes |
| quercetin | polyphenol | no |
| quinizarin | dye | no |
| theaflavin monogallate | polyphenol | no |
| thioguanosine | antimetabolite antineoplastic | yes |
| triamcinolone | synthetic corticosteroid anti-inflammatory | yes |

10/15 (66%) worm repurposing hits are plant-based polyphenols. Epicatechin monogallate and epigallocatechin-3-monogallate are structural analogs, the latter containing an additional hydroxyl group. They are both abundant in tea leaves. The two plant flavonols quercetin and gossypetin are structural analogs, the latter containing an additional hydroxyl group. Quercetin is found in many foods and gossypetin is found is *Hibiscus* flowers. Aripiprazole, benserazide and ethylnorepinephrine act on catecholamine levels and/or signaling in the brain. Phenylbutazone is a decades-old first-generation non-steroidal anti-inflammatory drug, or NSAID, but is not currently approved and marketed in the United States. Triamcinolone is a synthetic corticosteroid and generic topical anti-inflammatory drug. These worm data suggest the existence of at least three mechanistic classes of suppressors: 1) polyphenolic antioxidants; 2) catecholamine pathway modulators; 3) anti-inflammatories.

#### Fly repurposing screen and hit validation

In the presence of 9μM bortezomib, *Pngl*^+/-^ flies are developmentally delayed as first-instar larvae. A fly repurposing hit is defined as any compound that in triplicate rescued bortezomib-treated *Pngl*^+/-^ larval growth and size such that they were indistinguishable from control wells containing DMSO-treated heterozygotes. All test compounds were screened at a final concentration of 32μM. Separation between positive and negative control wells is shown in **Figure 2C**. As expected, most test compound wells do not affect bortezomib-treated heterozygotes as determined by quantifying larval area per well and remain close to the negative control Z-score, (**Figure 2D**). We identified 31 fly suppressors that rescued larval size with Z-score greater than 2.5 in at least two out of three replicates, yielding a hit rate of 1.2%, or half the hit rate of the worm repurposing screen (**Supplemental Figure 2**) but thirty times the hit rate of the previously published *Pngl*^−/−^ drug repurposing screen (Rodriguez et al., 2018).

In order to eliminate suppressors that directly inactivate bortezomib or that fail to rescue in the absence of bortezomib-induced proteasomal stress, we retested suppressors in a 12-well petri dish assay at 0, 10μM, 50μM, 100μM of compound on either *Pngl*^+/-^ larvae in the presence of bortezomib or *Pngl^−/−^* larvae without bortezomib. We could not retest suppressors in a carfilzomib assay paradigm because *Pngl*^+/-^ larvae were resistant to carfilzomib toxicity at the highest concentration tested. Note in the DMSO-hypersensitive homozygous animals that we might not observe rescue at 100μM because of the increased amount of DMSO (0.2%) needed to test that concentration of compound. Although we did not confirm by chemical analysis, any compound that strongly rescued only in the bortezomib assay paradigm is likely a bortezomib inactivator. The top candidate bortezomib inactivators are tannic acid and gossypetin (**Supplemental Material**). Gossypetin contains two catechols per molecule and tannic acid contains five catechols per molecule. These two compounds are the only overlapping primary screening hits between the worm and fly repurposing screens.

30/31 fly suppressors were reordered as fresh powder stocks (one compound failed to dissolve at 10mM as stock solution). 21/30 (70%) fly suppressors retested in the primary heterozygote assay paradigm, which is comparable to the retest rate of worm suppressors. 20/30 (66%) fly suppressors scored positively in the secondary homozygote assay paradigm. 13/30 (43%) suppressors validated in both homozygote and heterozygote assay paradigms, which is higher than the worm cross-validation rate. Those 13 compounds are the fly repurposing hits (**Table 1**): aminacrine, aripiprazole, β-Amyrin, butopyronoxyl, clindamycin, dyphylline, L-glutamine, ketorolac, lumefantrine, promazine, pyrimethamine, quinizarin and thioguanosine.

The fly repurposing hits span a range of mechanisms, including an overlap with the worm repurposing hits that fall into the three aforementioned pharmacological classes. Only one of the fly repurposing hits appears to be fly-specific, e.g., the insect repellent butopyronoxyl. Ketorolac is a first-generation NSAID anti-inflammatory drug but is structurally distinct from the worm repurposing hit and NSAID phenylbutazone. β-Amyrin is triterpene natural product with antiinflammatory effects. As stated above, aripiprazole is a catecholamine receptor modulator. The remaining fly repurposing hits appear to be mechanistic singletons and do not have obvious functional or structural analogs with worm repurposing hits. There are two antimalarial compounds, pyrimethamine and lumefantrine, and two DNA-damaging cancer drugs, aminacrine and thioguanosine. Dyphylline is a xanthine derivative with bronchodilator and vasodilator effects. Clindamycin is a lincosamide antibiotic used for a range of bacterial infections.

#### Cross-validation of worm and fly repurposing hits

In order to quantify the overlap between worm and fly repurposing hits, we selected 11 worm repurposing hits for cross-testing in both fly assay paradigms, and 19 fly repurposing hits for cross-testing in both worm assay paradigms. The fraction of worm repurposing hits that rescue in flies is higher than the fraction of fly repurposing hits that rescue in worms. 11/11 (100%) of worm repurposing hits scored positively in the fly heterozygote assay paradigm. 6/11 (55%) of these compounds cross-validated in both fly assay paradigms. There may be several reasons why a compound scored positively in a secondary retest in one organism but was not a hit in the primary screen in the same organism. Especially in flies because primary screens in flies have more variance than primary screens in worms due to fewer animals per well, greater complexity of food source, and increased developmental stochasticity.

However, 3/19 (16%) fly repurposing hits scored positively in the worm bortezomib paradigm. 3/19 (16%) fly repurposing hits scored positively in the worm carfilzomib paradigm. Only 1/19 (5%) fly repurposing hit cross-validated in both worm assay paradigms: aripiprazole. Of the 30 repurposing hits that were cross-tested, seven compounds (23%) are active both worm assay paradigms and in both fly assay paradigms: aripiprazole, benserazide, phenylbutazone, pomiferin, quercetin, theaflavin monogallate and 3,4-didesmethyl-5-deshydroxy-3’-ethoxyscleroin. Their chemical structures are shown in **Figure 3**.

**Figure 3.**
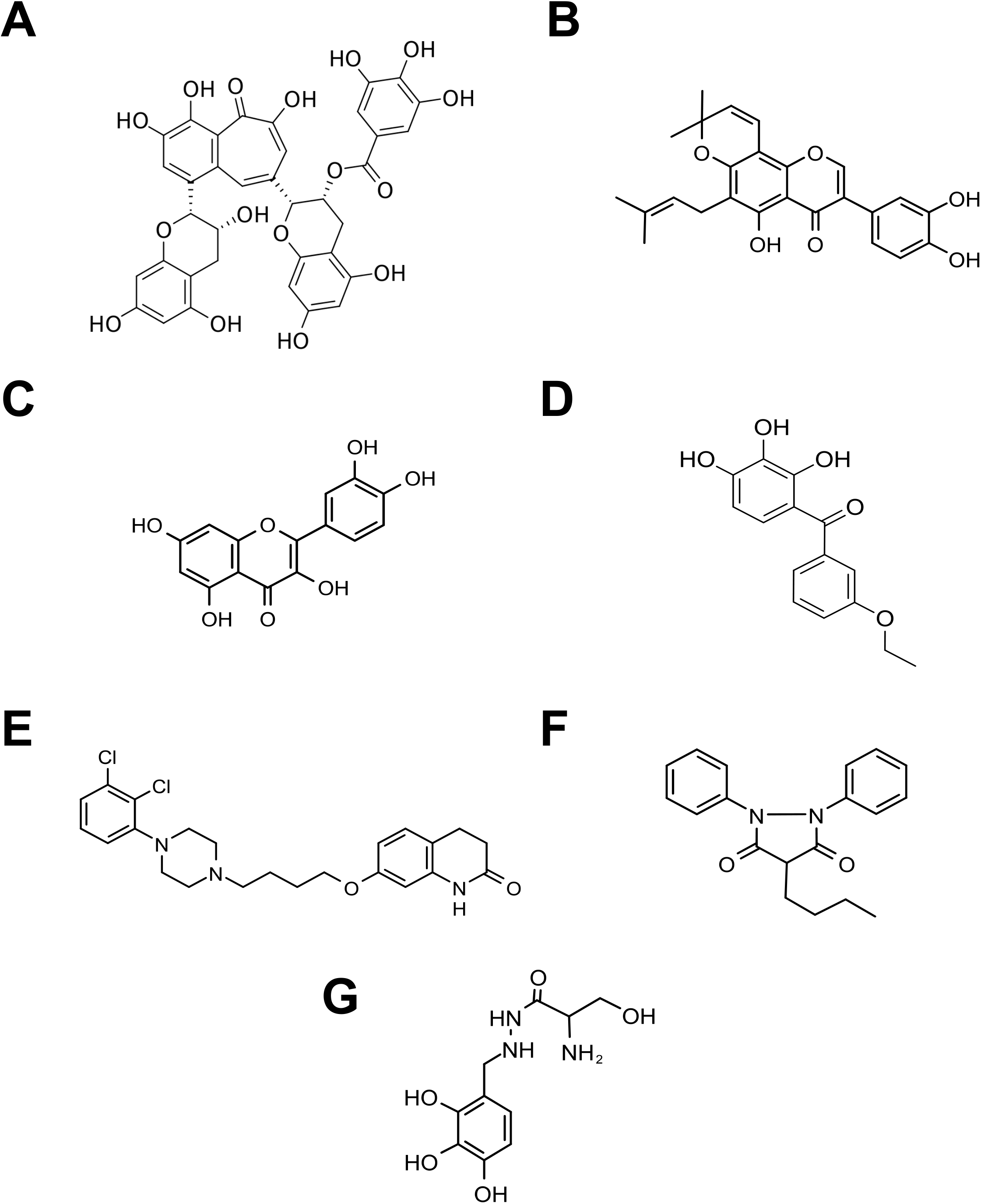
Chemical structures of cross-validated drug repurposing hits. **(A)** theaflavin monogallate. **(B)** pomiferin. **(C)** quercetin. **(D)** 3,4-didesmethyl-5-deshydroxy-3’-ethoxyscleroin. **(E)** aripiprazole. **(F)** phenylbutazone. **(G)** benserazide.

Aripiprazole is an atypical antipsychotic drug and dopamine receptor agonist approved for use alone or in combination in adults and children for the treatment of symptoms of many CNS diseases, including schizophrenia, autism spectrum disorder and bipolar disorder (Shapiro et al., 2003; Kikuchi et al., 1995). Aripiprazole has been available in generic form since 2015. Benserazide is an aromatic L-amino acid decarboxylase inhibitor which has been coadministered with levodopa (L-Dopa) for decades to boost dopamine levels in the brain in the treatment of Parkinson disease. As mentioned above, phenylbutazone is decades-old NSAID and quercetin is plant flavonol with complex pharmacology, including anti-inflammatory effects. Theaflavin monogallate is a polyphenol found in black tea leaves. Pomiferin is prenylated isoflavone found in osage orange trees.

#### Drug discovery screens, secondary retests and cross-validation of novel hit compounds

Using the same screening conditions and hit-calling analysis as the worm repurposing screen, we scaled up efforts with *png-1* null mutant larvae and a 20,240-member lead discovery library. We identified 28 novel worm suppressors and 6 novel worm enhancers/toxic compounds with a combined hit rate of 0.143% (**Figure 4A; Supplemental Material**). Enhancers/toxic compounds were not further investigated in this study. 12/28 (43%) novel worm suppressors retested in the bortezomib assay paradigm. 10/28 (36%) novel worm suppressors scored positively in the carfilzomib assay paradigm. A total of five novel worm suppressors (18%) validated in both paradigms, which is comparable to the cross-validation rate observed for worm repurposing hits. Consistent with the results of the repurposing screens, the fraction of novel worm suppressors that are active in flies is higher than the fraction of novel fly suppressors that are active in worms. 4/5 (80%) of novel worm suppressors cross-validated in both fly assay paradigms.

**Figure 4.**
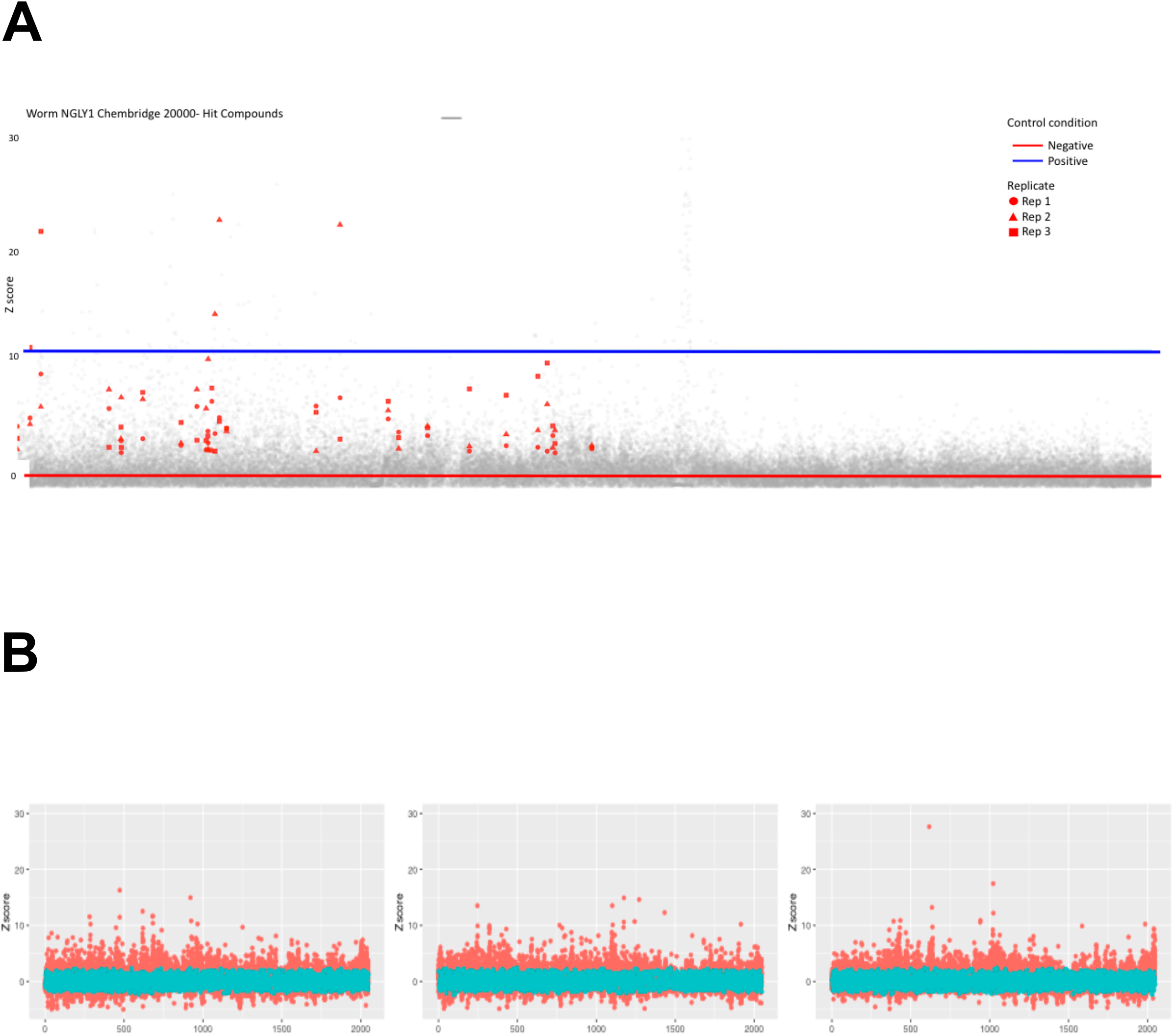
20,240-compound novel lead discovery screens of *png-1* homozygous mutant worms and *Pngl* heterozygous mutant flies. **(A)** Worm screen Z-score plot of 20,240 test compounds in triplicate. Replicate 1 is shown as red circles. Replicate 2 is shown as red triangles. Replicate 3 is shown as red squares. The mean negative control Z-score is indicated by the red line. The mean positive control Z-score is indicated by the blue line **(B)** Fly screen Z-score plot of 20,240 test compounds in triplicate. Replicate 1 is the left panel. Replicate 2 is the center panel. Replicate 3 is the right panel.

**Figure 5.**
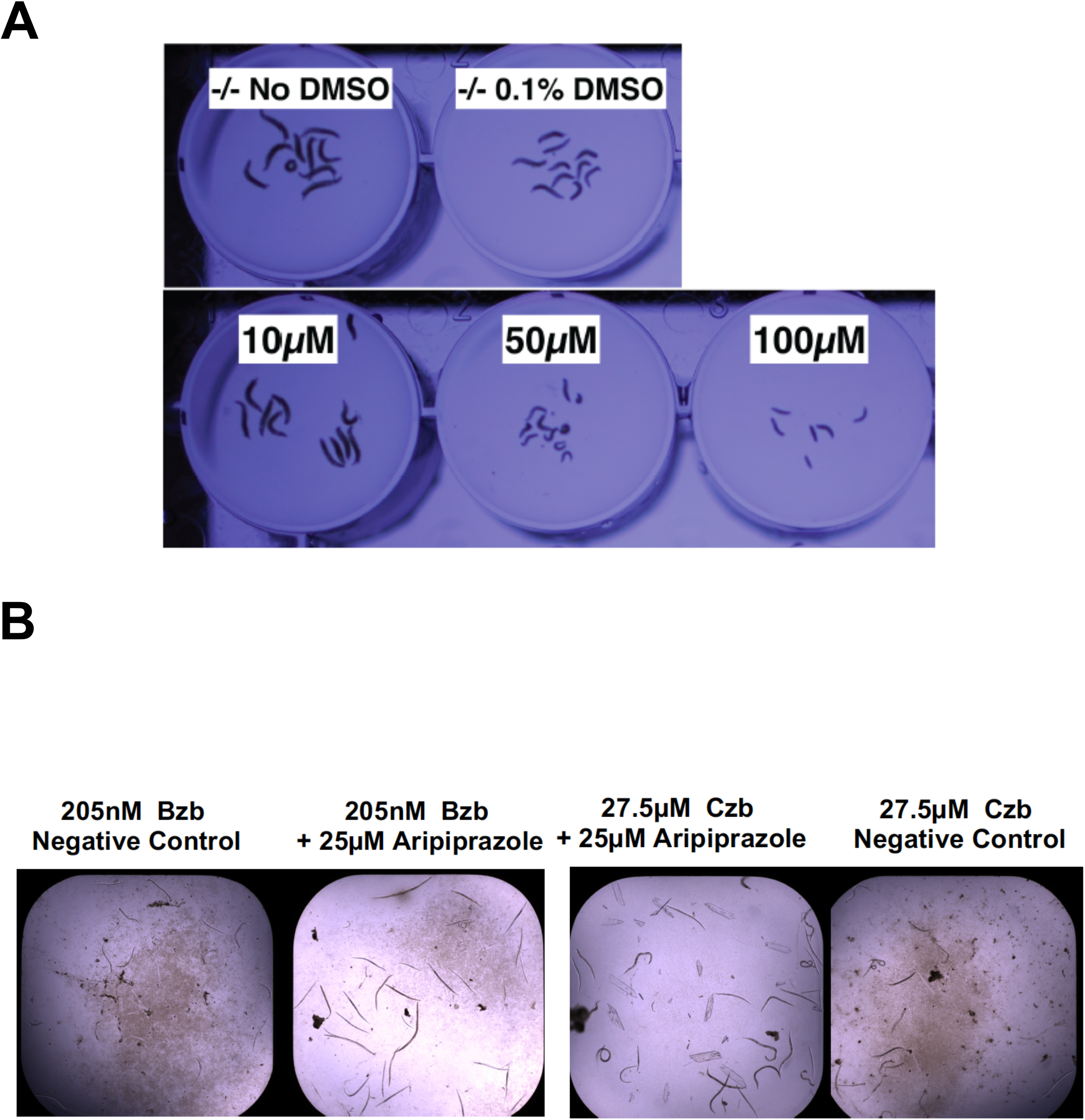
Aripiprazole worm and fly cross-validation data. **(A)** *Pngl*^−/−^ homozygote larvae treated with 10μM, 50μM and 100μM aripiprazole compared to untreated controls. **(B)** *png-1* homozygote larvae treated with 25μM aripiprazole in the presence of 205nM Bortezomib (Bzb) or 27.5μM carfilzomib (Czb).

Using the same screening conditions and hit calling analysis for the fly repurposing screen, we scaled up efforts with *Pngl*^+/-^ larvae and a 20,240-member lead discovery library. We identified 16 novel fly suppressors, resulting in hit rate of 0.07% (**Figure 4B; Supplemental Material**). 15/16 novel fly suppressors were reordered as fresh powder stocks. 10/15 (67%) novel fly suppressors retested in the primary heterozygote assay paradigm. 13/15 (87%) novel fly suppressors scored positively in the secondary homozygote assay paradigm. A total of 10 novel fly suppressors (67%) validated in both paradigms. Of the 13 novel fly suppressors, 12 were retested in both worm assay paradigms. None of the novel fly suppressors scored positively in the bortezomib assay paradigm, but 8/12 (66%) compounds scored positively in the worm carfilzomib assay paradigm.

The known NRF2 activator sulforaphane did not rescue in either worm assays but it did score positively in the *Pngl*^−/−^ homozygote assay paradigm, rescuing this mutant to 50% of the control (**Supplementary Material**). In the most striking example of overlap between the repurposing library and discovery library, we identified two novel fly suppressors that are structural analogs of pyrimethamine, sharing a diaminopteridine group present in folic acid analog inhibitors of DNA synthesis enzymes, such as methotrexate. Both pyrimethamine and the two novel fly suppressors rescue homozygotes to near 100% of the heterozygote control. A second novel fly suppressor also rescues homozygotes to near 100% of the heterozygote control.

#### Keap1-NRF2 activation assay in human cells

A total of 91 repurposing hits and novel suppressors from both worm and fly screens were cross-tested in a Keap1-NRF2 activation assay in the U2OS osteosarcoma cell line. 7/91 (8%) compounds are active with a half-maximal effective concentration (EC_50_) between 2-30μM (**Table 2**), or 10-100 fold less potent than the positive control NRF2 activators, e.g., sulforophane. The remaining 84 compounds have EC_50_ values above 50μM and were not considered further for lack of potency. Listed here in order of percentage of maximal response normalized to the positive control CDDO methyl ester: aripiprazole (78%) = pyrogallin (78%) > fisetin (59%) > purpurogallin-4-carboxylic acid (53%) = 3-methoxycatechol (50%) > rhamnetin (43%) > pyrimethamine (27%). Of these seven only fisetin has been previously shown to activate NRF2 target gene expression (Smirnova et al., 2011). To increase confidence in the assay, we also tested three additional known NRF2 inducers: sulforaphane, omaveloxolone and dimethyl fumarate. None of the novel lead worm or fly suppressors are NRF2 inducers.

**Table 2.**
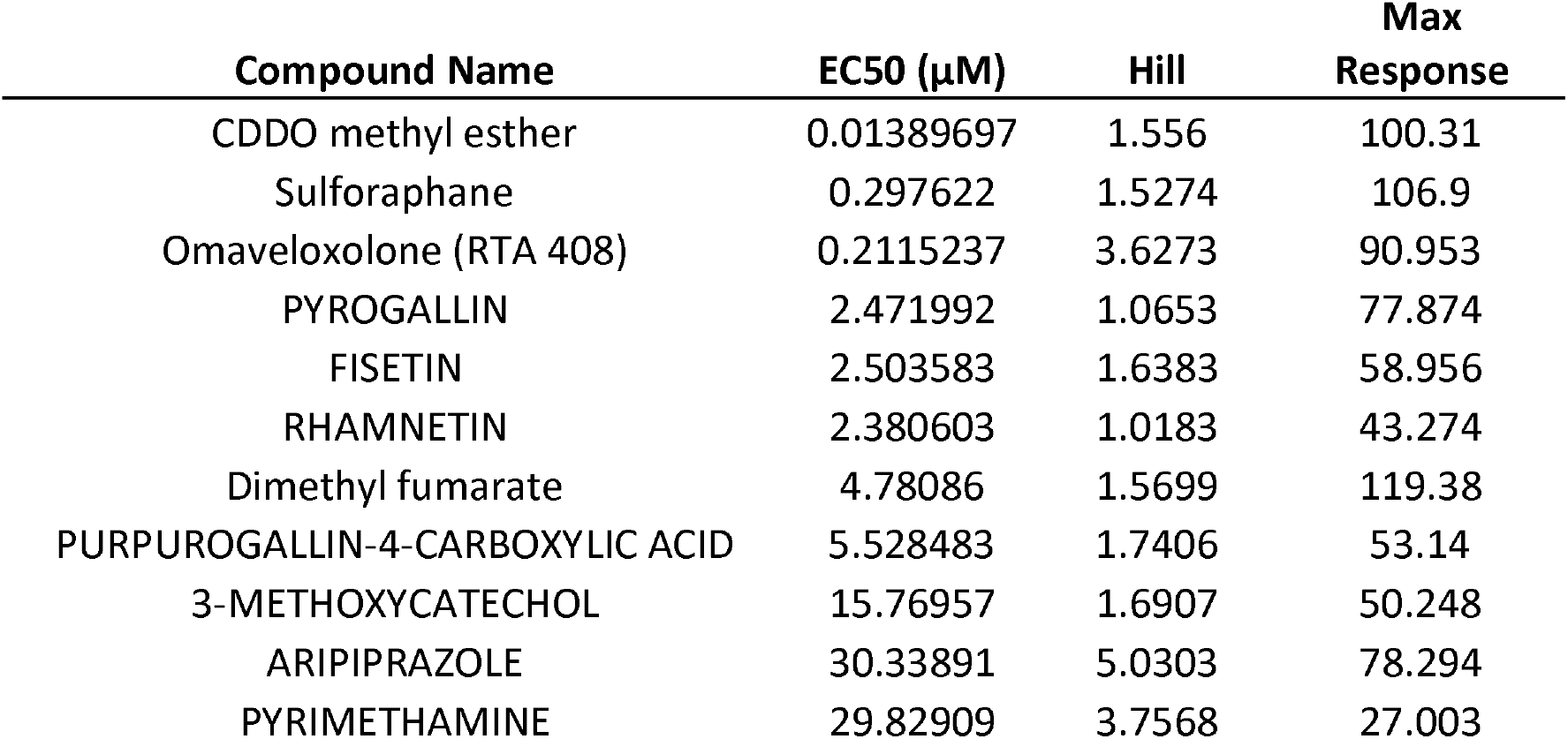
Keap1-NRF2 transcriptional activity reporter data

### Discussion

In summary, we demonstrated the therapeutic relevance of unbiased phenotypic drug screens of worm and fly models of NGLY1 Deficiency. We addressed the major shortcoming of our previous drug screening effort by increasing hit rate and reproducibility with the more permissive *Pngl^+/-^* heterozygote assay as the primary screen followed by a secondary retest in the more restrictive *Pngl*^−/−^ homozygote assay paradigm, which selected for the subset of hit compounds that specifically rescues loss of NGLY1. We addressed the other significant shortcoming of our previous drug screening effort by including worms as a second species in the primary screening stage, which increased the total number and mechanistic diversity of hits. And by including human cells as a third species in the hit validation stage we selected for hit compounds that act on conserved targets and pathways. We identified three clinic-ready therapeutic approaches by focusing on the subset of hit compounds that rescue larval growth and development in both worms and flies and in the presence and absence of bortezomib: (1) NRF2 activators/inducers; (2) catecholamine boosting drugs; (3) anti-inflammatory drugs. Only one compound was found to be active in all three species (worm, fly and human cells) and in all assay paradigms (plus bortezomib, plus carfilzomib, and NRF2 reporter): the atypical antipsychotic aripiprazole.

Before discussing why we observed those three pharmacological classes and the evidence that supports their testing in clinical trials, we consider reasons why any given hit compound might be active in one or a combination of the species and assay paradigms tested herein. Some reasons have to do with compound stability, compound solubility, bioavailability, drug metabolism and pharmacokinetics, not to mention the differences between worm screens versus fly screens. The following is by no means an exhaustive list of salient factors. Worm screens were all performed in simple liquid media that is a buffered salt solution plus essential nutrients and contains bacteria as a food source. By contrast, the fly screens were performed in a nutrient-rich solid media (“fly food”) with a molasses base. The worm screen is a six-day assay that spans all stages of the worm life cycle while the fly screen is a three-day assay that only spans the larval stages of fly development. The worm screen was performed with 205nM bortezomib while the fly screen was performed with 9μM bortezomib. Primary drug screens with flies involve as few as three animals per test well, while primary drug screens with worms involve a minimum of 15 animals per test well.

Other reasons why a hit compound is active in one species but not the other may have to do with pharmacodynamics, that is the drug targets are not evolutionarily conserved or the drug binding sites of shared drug targets are too divergent between worm and fly orthologs. Resolving which pharmacokinetic and pharmacodynamic variables and their relative contributions explain the failure of a hit compound in any given assay or species paradigm is a very important research question that is beyond the scope of the present study. However, what we can conclude from this study is that NRF2, anti-inflammatory and catecholamine pathways appear to be evolutionarily conserved between worms, flies and human cells.

Indeed, NRF2 pathway activators and anti-inflammatory drugs have been proposed as clinical approaches to address the underlying defects in mitochondrial physiology observed in **NGLY1*-* deficient cells, specifically a defect in mitophagy and excessive mitochondrial fragmentation (Yang et al., 2018). In addition to its more well studied role as a transcriptional regulator of the proteasome bounce-back response, Nrf1 also controls the expression of mitochondrial homeostasis and mitophagy gene not only in human cells but also in worms (Paek et al., 2012) and flies (Tsakiri et al., 2013). A unifying theory of NGLY1 Deficiency is that disease phenotypes are primary or secondary consequences of the loss of conserved Nrf1-dependent gene expression programs in cell types and tissues that are vulnerable to proteotoxic, oxidative or redox stress. A recent n-of-1 clinical study of a NGLY1 patient who died of complications due to adrenal insufficiency suggest that steroidogenic secretory tissues fit the criteria of a vulnerable cell type (van Keulen et al., 2019). The *Pngl*^−/−^ fly has defects in its physiologically analogous neuroendocrine tissues that fail to produce and secrete the cholesterol-derived hormone 20-hydroxyecdysone (Rodriguez et al., 2018). That said, there is only a single ortholog of both Nrf1 and NRF2 in worms and flies, so the functions of worm *SKN-1* and fly *cnc* may not recapitulate all functionality of mammalian Nrf1 and NRF2.

Why do anti-inflammatory drugs rescue worm and fly models of NGLY1 Deficiency? Defects in mitophagy lead to release of mitochondrial genomic DNA and mitochondrial-encoded RNA from fragmented and damaged mitochondria into the cytoplasm drives constitutive cGAS-STING and related innate immunity responses in *NGLY1^−/−^* mice and cell lines (Yang et al., 2018). These innate immunity pathways are conserved in flies (Martin et al., 2018) and worms (Wu et al., 2014). We speculate that mitochondrial fragmentation and mitophagy defects are present in cells and tissues of *png-1* null mutant worms and *Pngl*^−/−^ mutant flies. Mitophagy defects and oxidative stress were in observed in *SKN-1* mutant worms (Palikaras et al., 2015). We therefore hypothesize that the steroidal and nonsteroidal anti-inflammatory drugs that rescue worm and fly models of *NGLY1* Deficiency, e.g., phenylbutazone, are suppressing basally elevated innate immune responses. There is evidence for this inflammation hypothesis in the recent NGLY1 literature. *SKN-1* is required for pathogen resistance in worms (Papp et al., 2012). Transcriptional profiling of *Pngl* knockdown flies revealed that genes involved in the innate immune response are up-regulated (Owings et al., 2018). The expression of interferon genes is constitutively up-regulated in *Ngly1^−/−^* murine embryonic fibroblasts (Yang et al. 2018). It’s tempting to speculate further that hyperactive innate immunity responses contribute to larval growth arrest and developmental delay in *NGLY1*-deficient animals and that these antiinflammatory responses are dampened by inhibitors of DNA synthesis, specifically purine biosynthetic enzymes targeted by the fly repurposing hits thioguanosine and the dihydrofolate reductase-like inhibitor pyrimethamine (Dziekan et al., 2019), and possibly the diaminopteridine-containing novel lead compounds that resemble pyrimethamine. Even more so because the same mechanism of action appears to have been revealed by parallel repurposing and discovery screens.

At this time, catecholamine boosters are supported by measurements of reduced catecholamine precursors in the cerebral spinal fluid of NGLY1 patients (Lam et al., 2017), as well as the fact that an adult and wheelchair-bound NGLY1 patient has received the dopamine precursor levodopa. Interestingly, there is evidence for dopamine insufficiency from a Pngl RNAi knockdown fly model of NGLY1 Deficiency, where the expression of genes involved in dopamine biosynthesis (e.g., tyrosine hydroxylase and the dopamine transporter DAT) are constitutively down-regulated (Owings et al., 2018). Future natural history studies of *NGLY1* Deficiency should focus on catecholamine insufficiency as a potential driver or axis of pathophysiology. The effects of aripiprazole in worms depends on catecholamine pathway genes (Osuna-Lugue et al., 2018). In flies, aripiprazole was shown to reduce the levels of an aggregated polyglutamine-expanded mutant protein in a model of Machado-Joseph disease, or spinocerebellar ataxia type 3 (Costa et al., 2016). The mechanism of rescue appears to rely on activation of proteotoxic and antioxidant stress responses. It is tempting to conjecture that aripiprazole achieved its rescue effects in the Machado-Joseph fly model and in our *NGLY1* Deficiency fly model by activating both NRF2 and catecholamine pathways; and further that the unique polypharmacology of aripiprazole combines catecholamine pathway activation and NRF2 pathway activation.

## Supporting information

All supplementary material, figures and legends

## Acknowledgements

We acknowledge Grace Science Foundation as a funding source, and we thank Dr. Matthew Might for feedback on the manuscript. We also thank the reviewers for their comments.

## Supplemental Material

Summary of cross-validation and retest experiments for worm and fly hit compounds from the Microsource Spectrum library and a Chembridge diversity library.

**Supplemental Figure 1.**
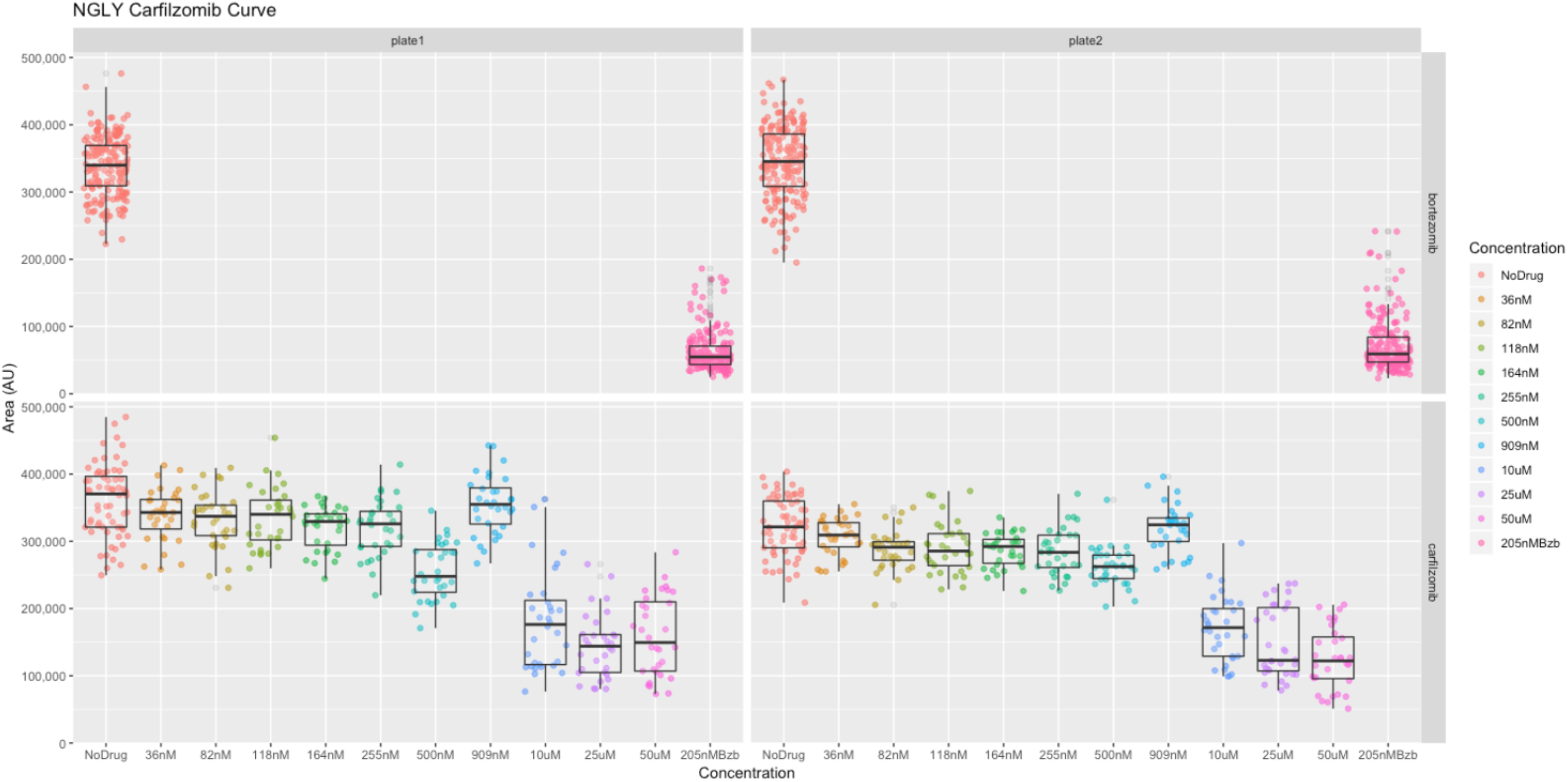
Carfilzomib dose-response experiment in *png-1* homozygous worms. Results from two technical replicates are shown. The y-axis is well area occupied by worms. As a comparison, the top panels show *png-1* homozygous worms treated with 205nM bortezomib. The bottom panels show *png-1* homozygous worms treated with ascending doses of carfilzomib.

**Supplemental Figure 2.**
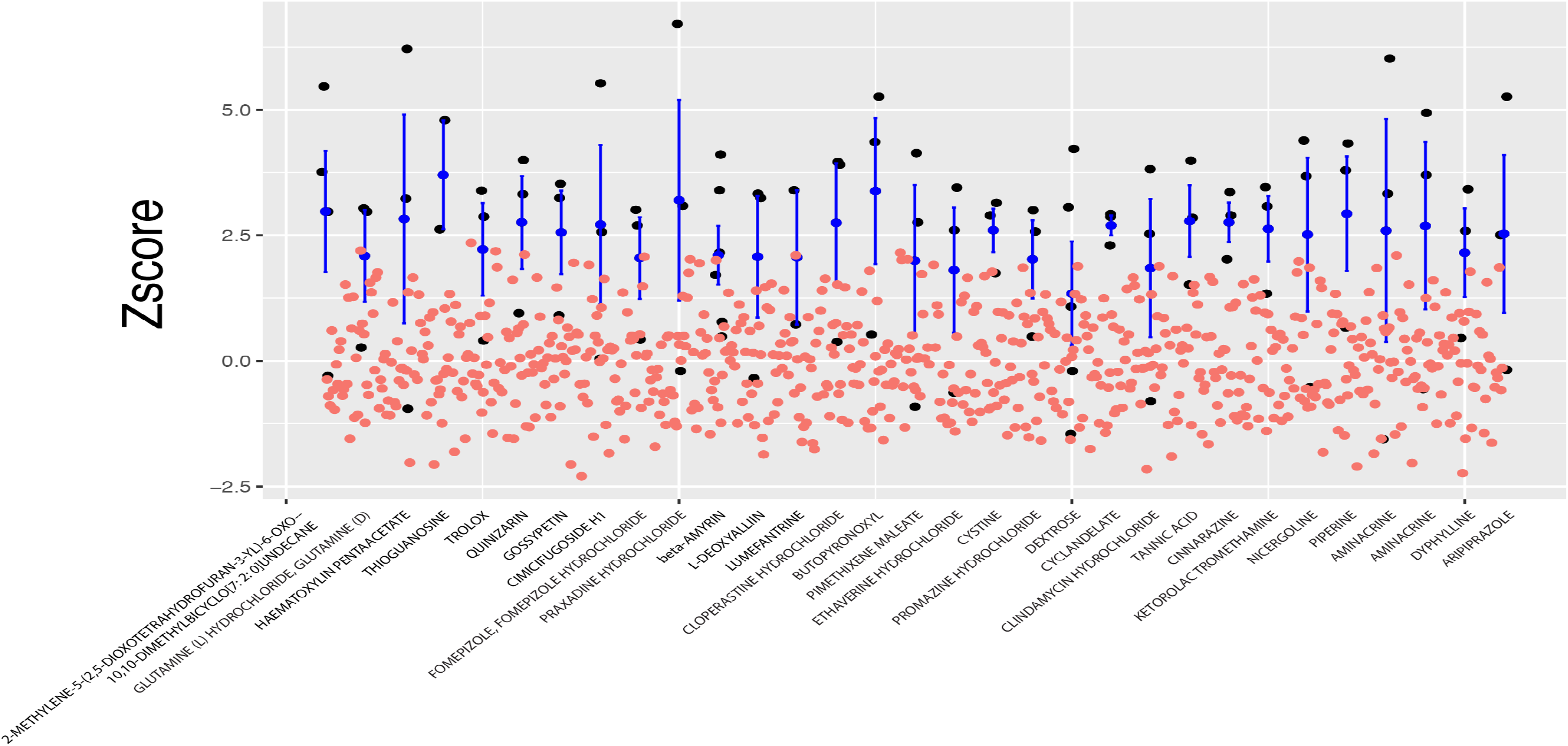
Z-score plot of 31 Microsource Spectrum repurposing hits from the *Pngl*^+/-^ fly primary screen. Red circles denote negative controls, black circles denote hit compounds wells, and blue circles correspond to the average of replicates of hit compounds.

**Supplemental Figure 3.**
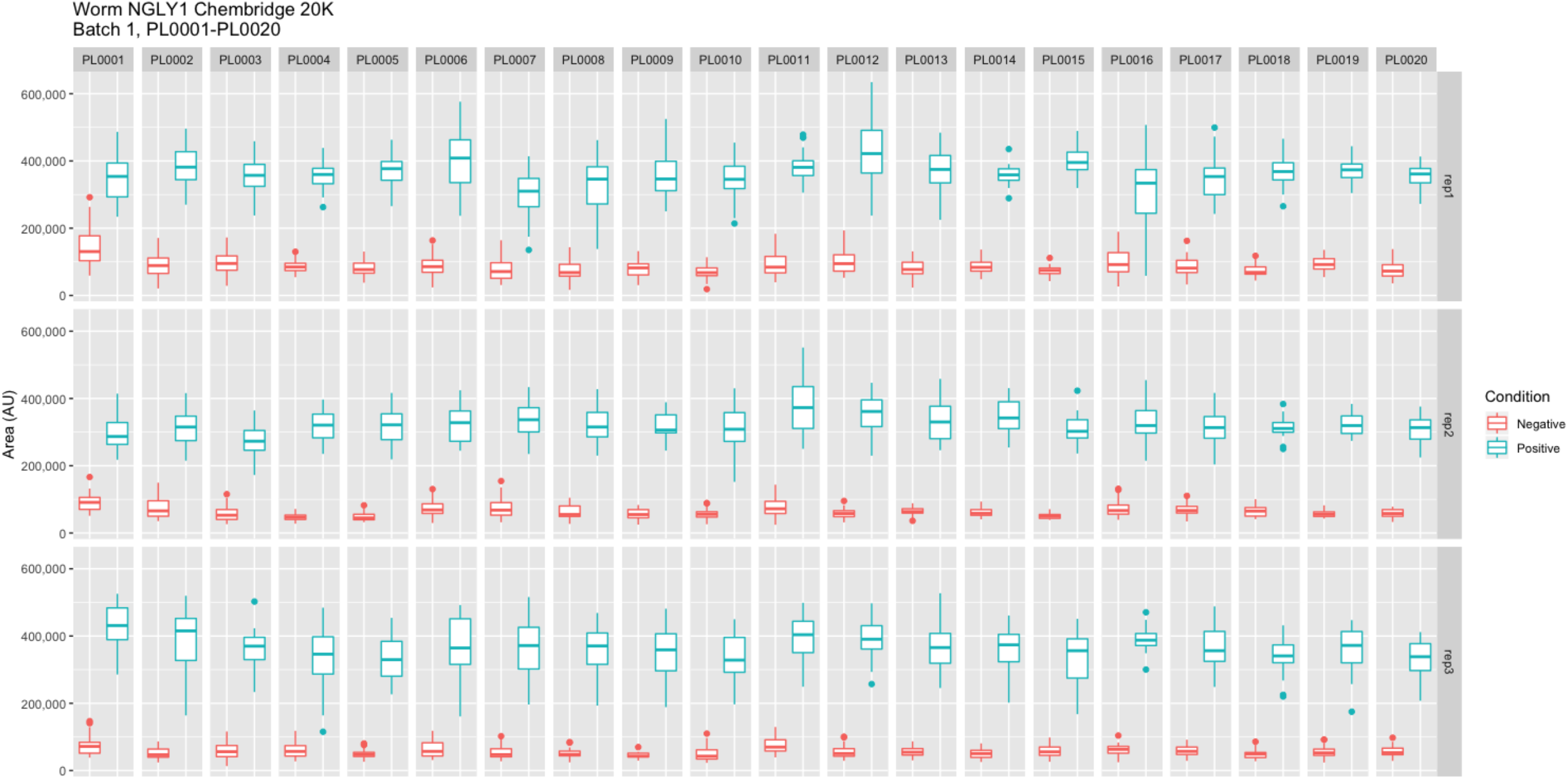
Box plots showing size separation of positive control versus negative control wells in the 20,240-compound lead discovery screen of *png-1* homozygous worms. Area (Arbitrary Units) refers to the area of each well that is occupied by worms.

**Supplemental Figure 4.**
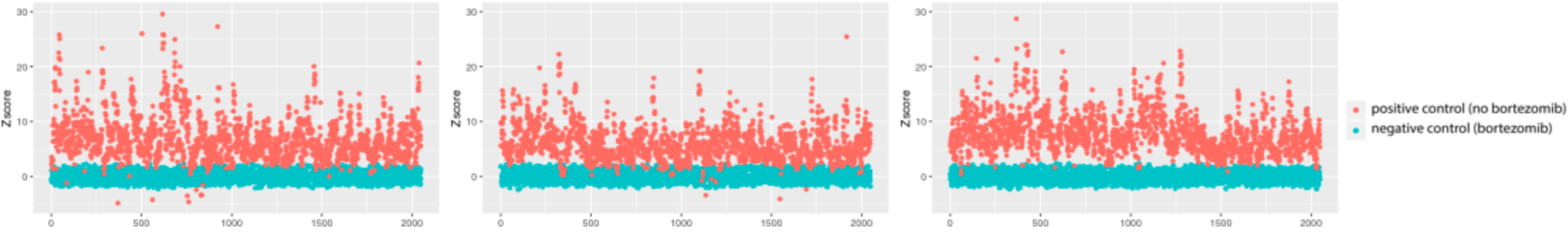
Box plots showing size separation of positive control versus negative control wells in the 20,240-compound lead discovery screen of *Pngl*^+/-^ fly larvae. Z scores were calculated as described in Methods and Materials.

