## Supplementary material for "Drug screens of NGLY1 Deficiency worm and fly models reveal catecholamine, NRF2 and anti-inflammatory pathway activation as potential clinical approaches": All supplementary material, figures and legends

**Supplemental Material**. Summary of cross-validation and retest experiments for worm and fly hit compounds from the Microsource Spectrum library and a Chembridge diversity library.

**Supplemental Figure 1**. Carfilzomib dose-response experiment in *png-1* homozygous worms. Results from two technical replicates are shown. The y-axis is well area occupied by worms. As a comparison, the top panels show *png-1* homozygous worms treated with 205nM bortezomib. The bottom panels show *png-1* homozygous worms treated with ascending doses of carfilzomib.

**Supplemental Figure 2**. Z-score plot of 31 Microsource Spectrum repurposing hits from the *Pngl* $^{+/-}$ fly primary screen. Red circles denote negative controls, black circles denote hit compounds wells, and blue circles correspond to the average of replicates of hit compounds.

**Supplemental Figure 3**. Box plots showing size separation of positive control versus negative control wells in the 20,240-compound lead discovery screen of *png-1* homozygous worms. Area (Arbitrary Units) refers to the area of each well that is occupied by worms.

**Supplemental Figure 4**. Box plots showing size separation of positive control versus negative control wells in the 20,240-compound lead discovery screen of *Pngl* $^{+/-}$ fly larvae. Z scores were calculated as described in Methods and Materials.

### Supplementary Figure 1

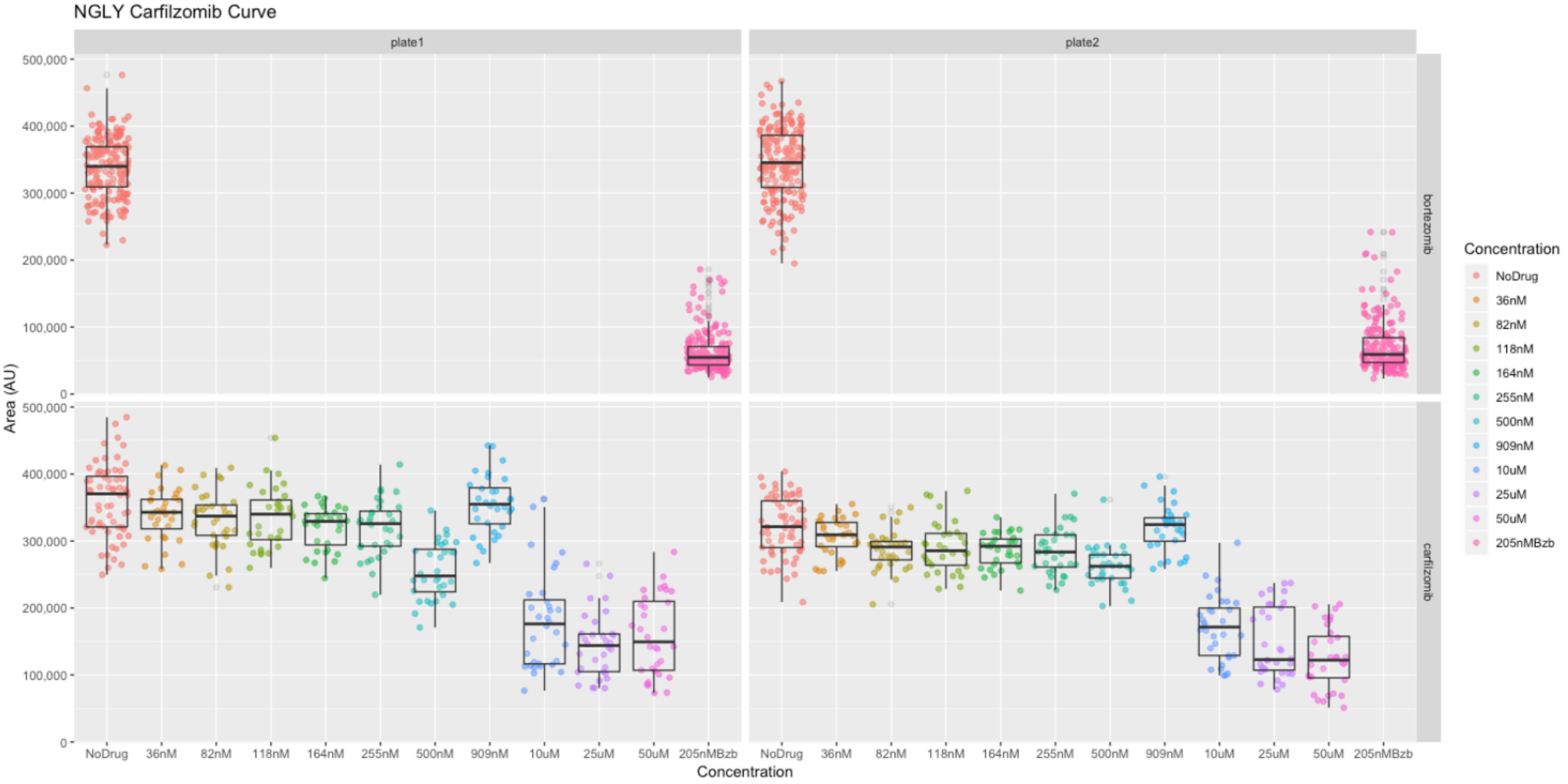

#### Supplementary Figure 2

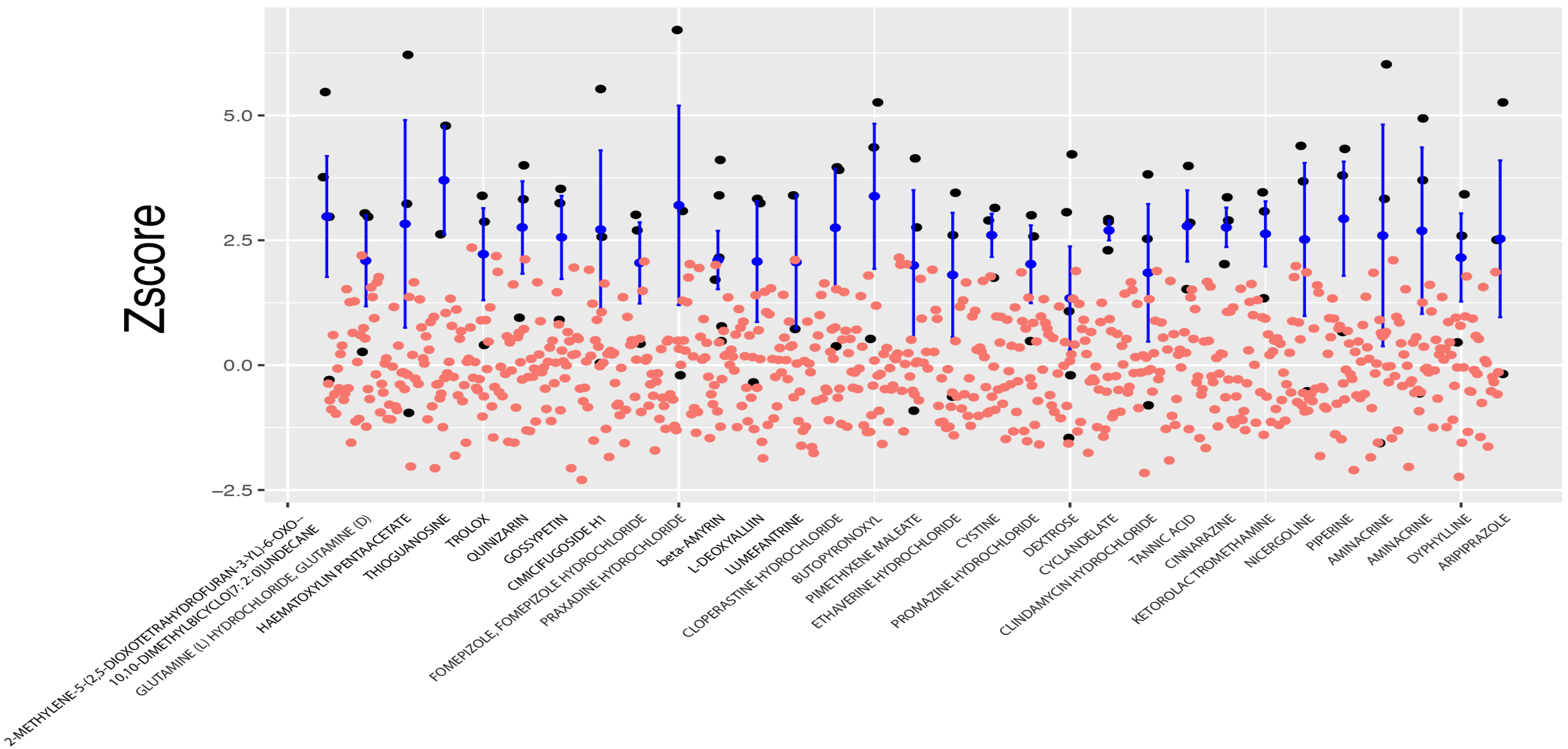

### Supplementary Figure 3

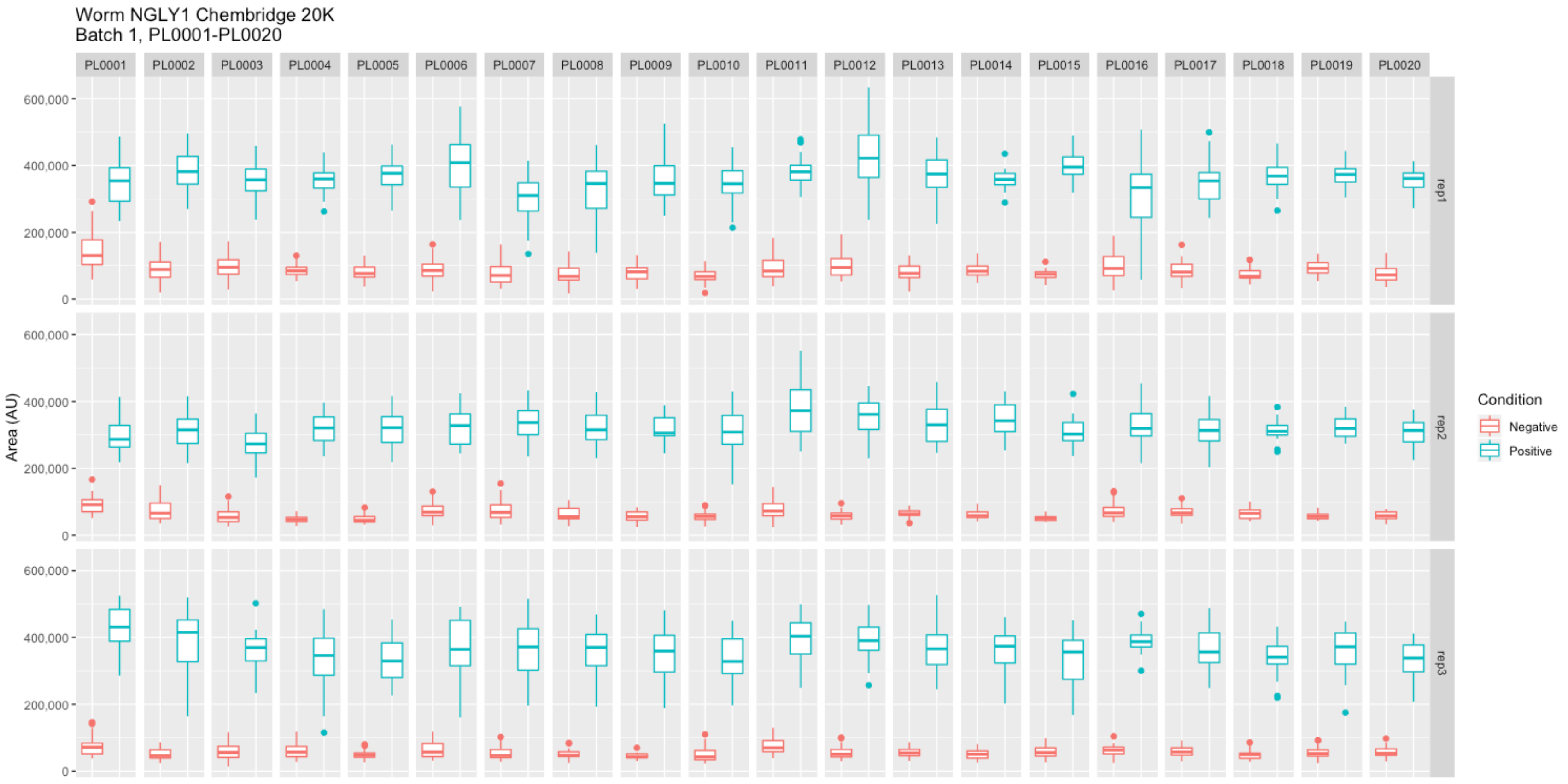

#### Supplementary Figure 4

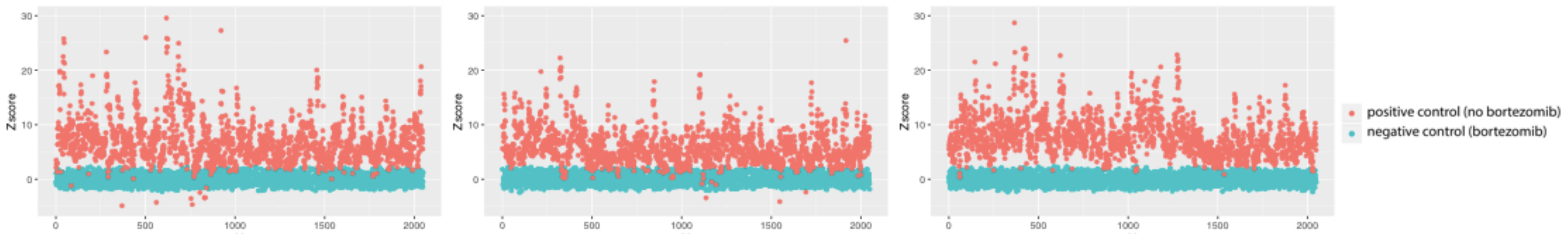

### Fly retest of Chembridge Worm hits

### Fly Chembridge retest – Supplier ID: 4141722

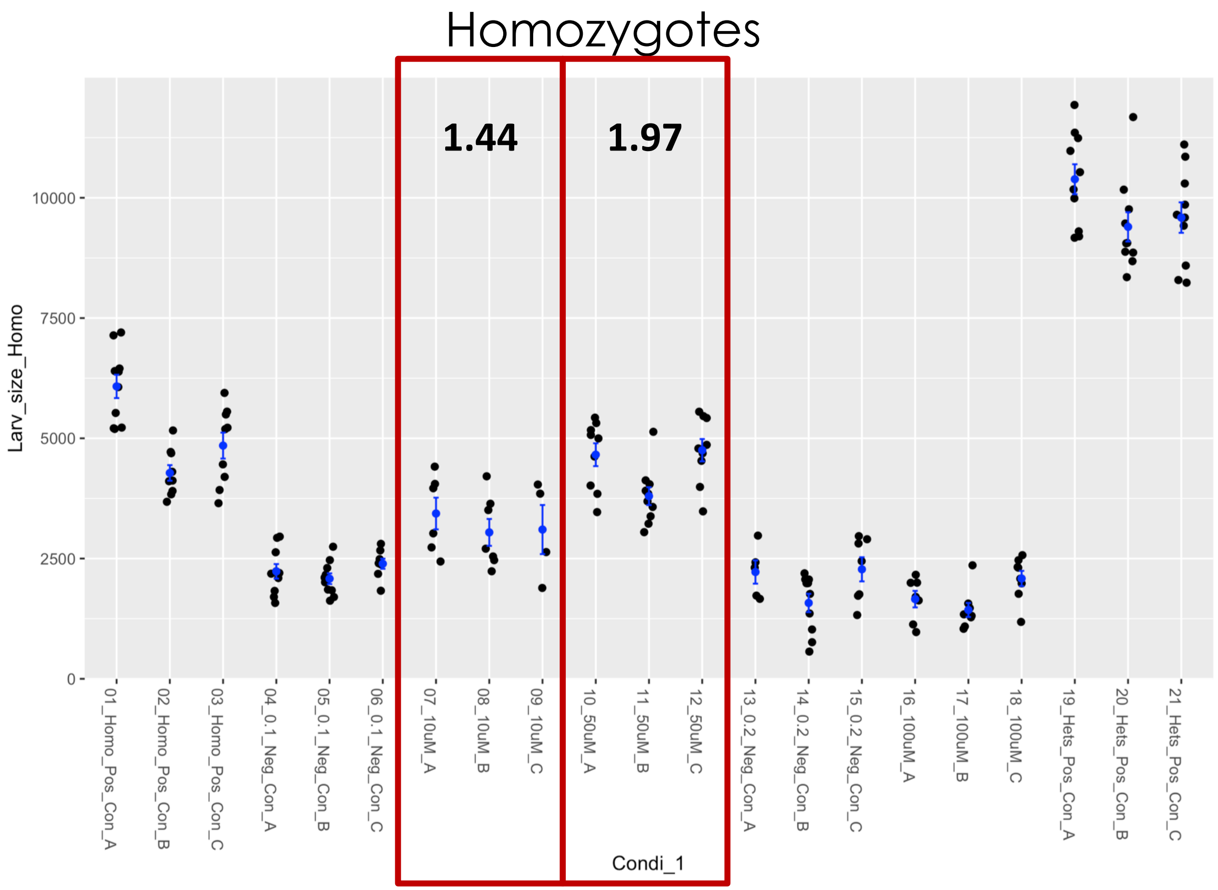


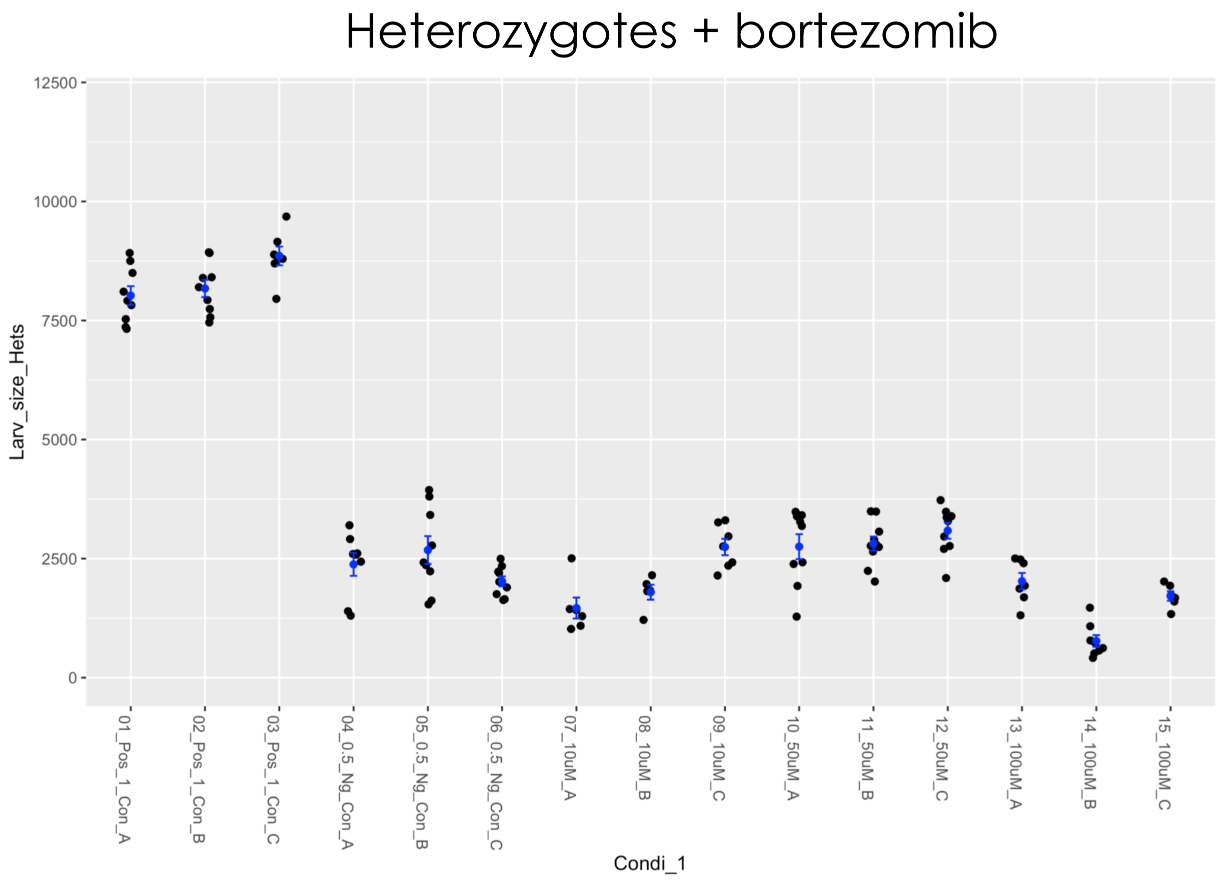

### Fly Chembridge retest – Supplier ID: 9223573

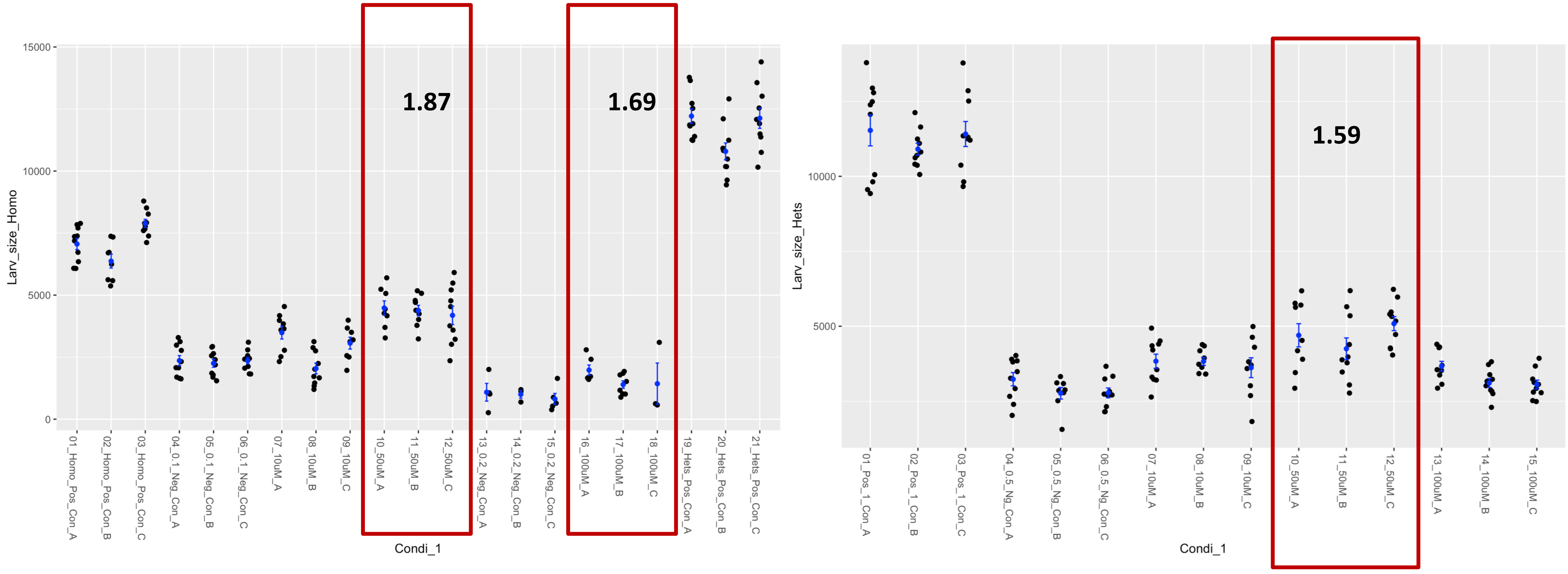

### Fly Chembridge retest – Supplier ID: 9268787

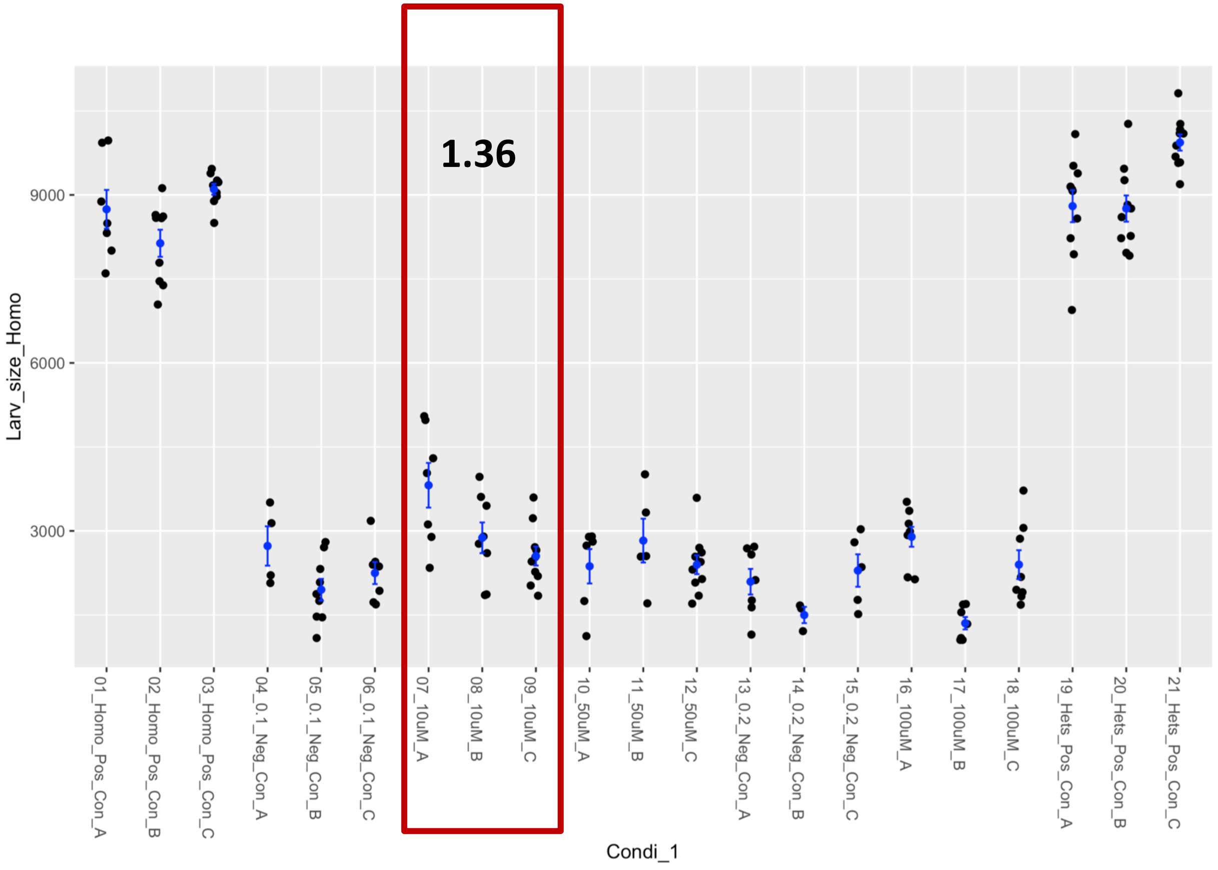


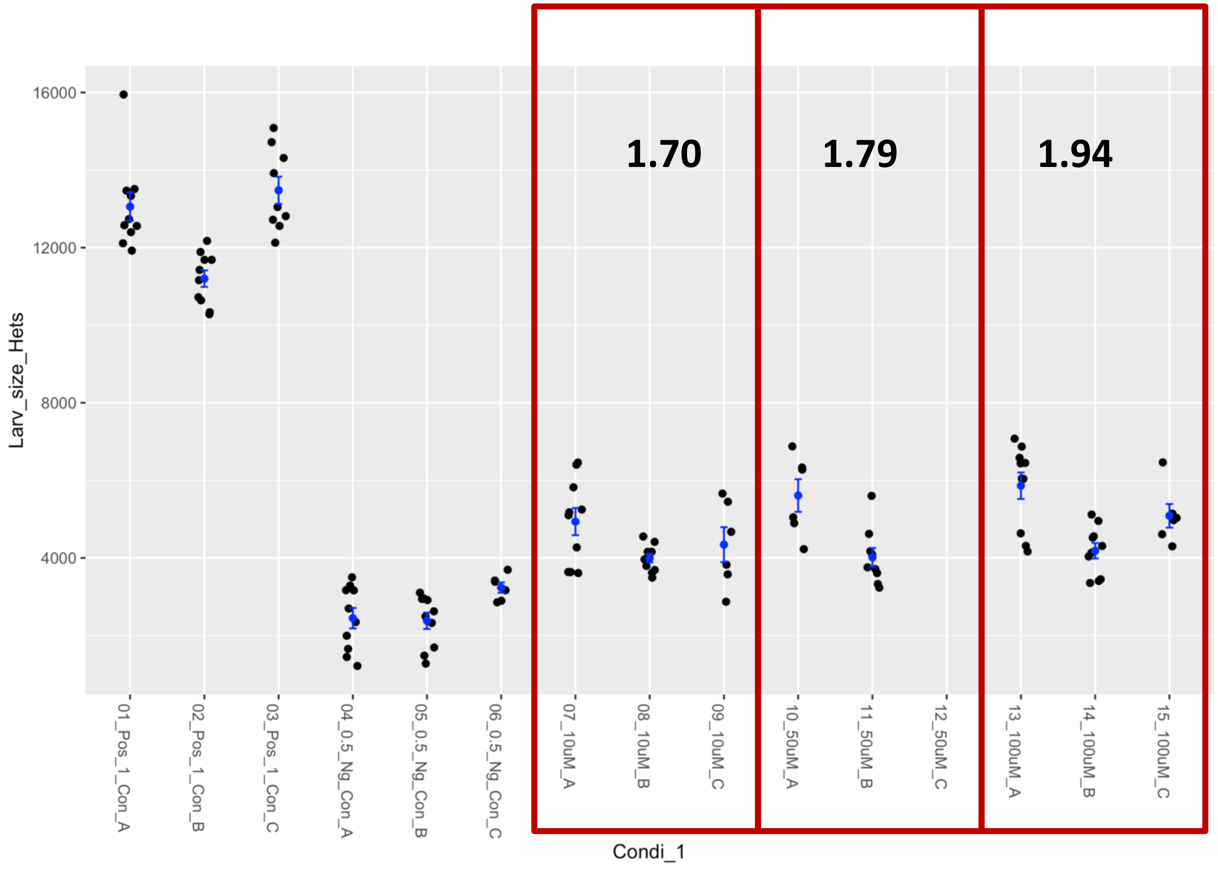

### Fly Chembridge retest – Supplier ID: 9283120

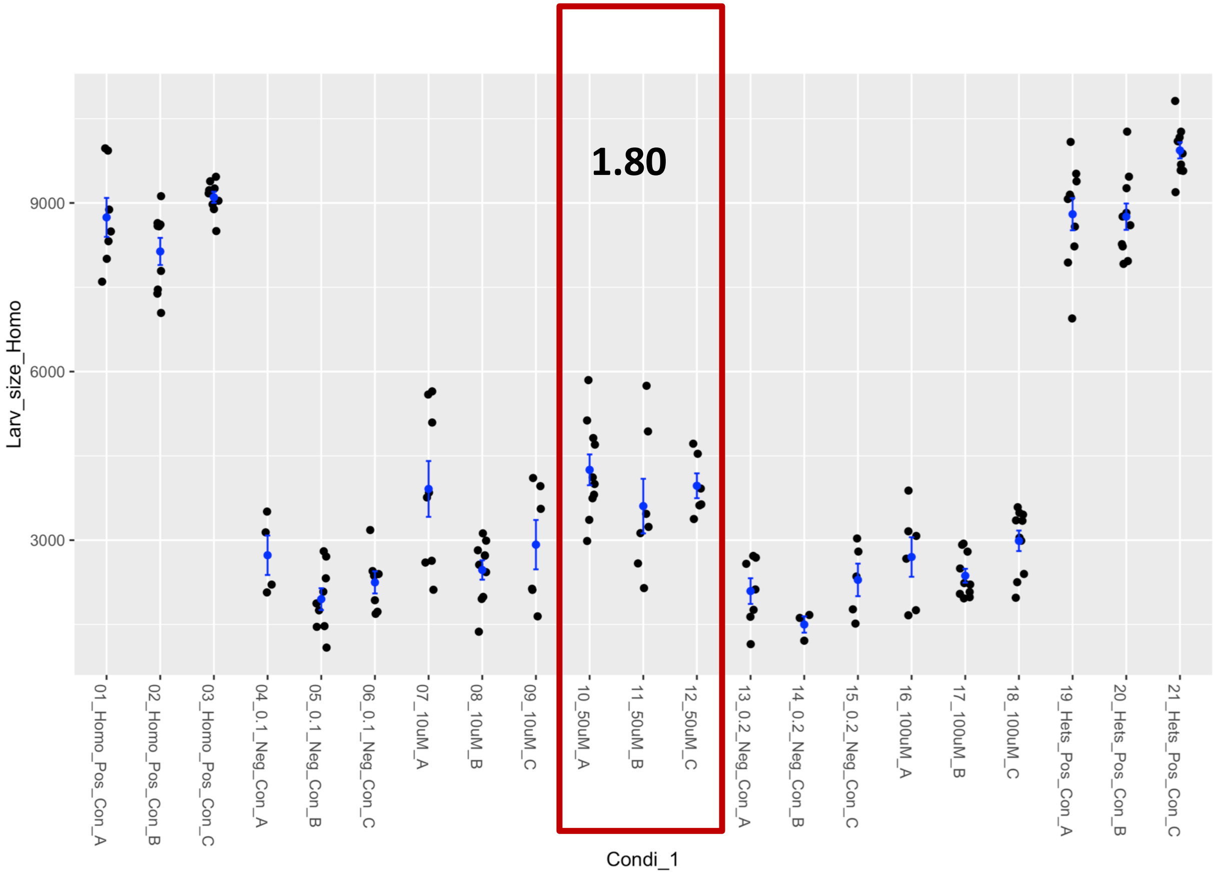


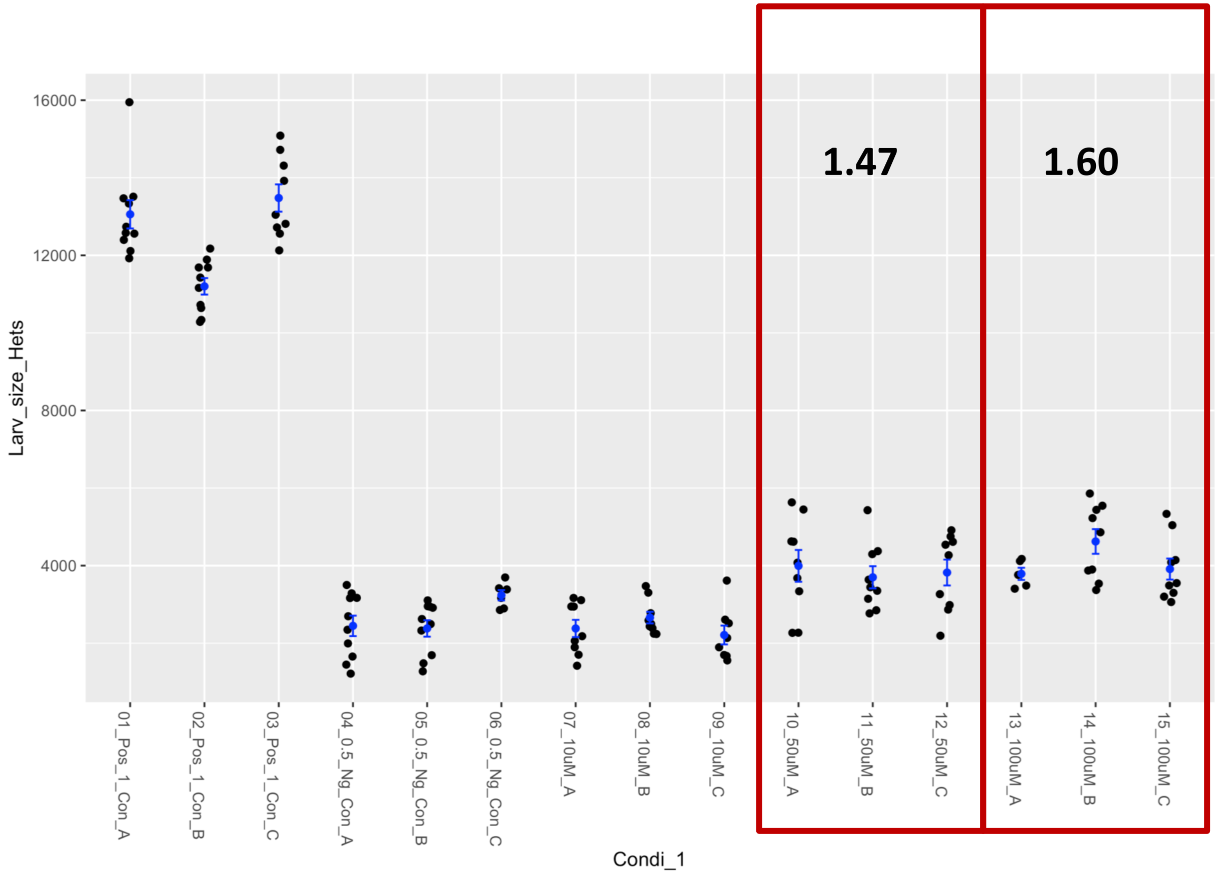

### Fly Chembridge retest – Supplier ID: 9313992

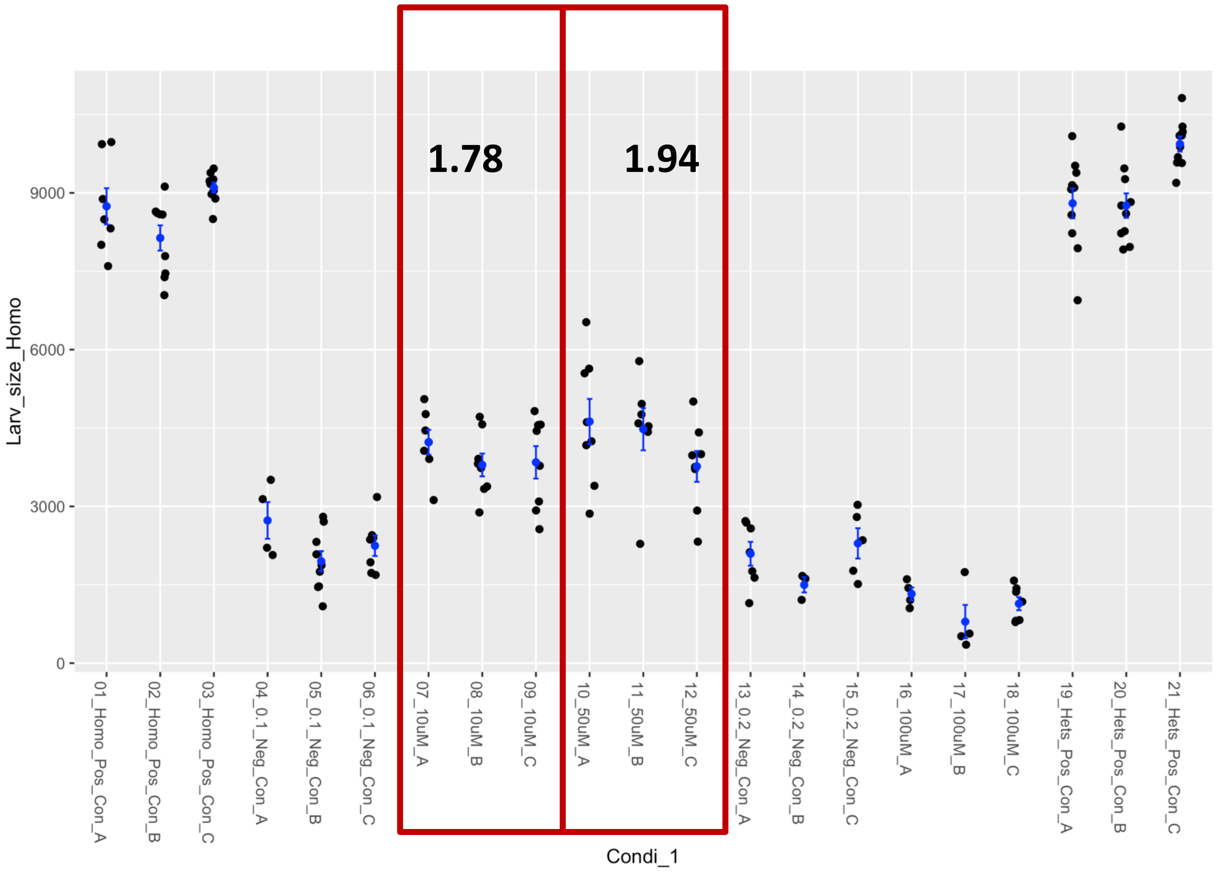


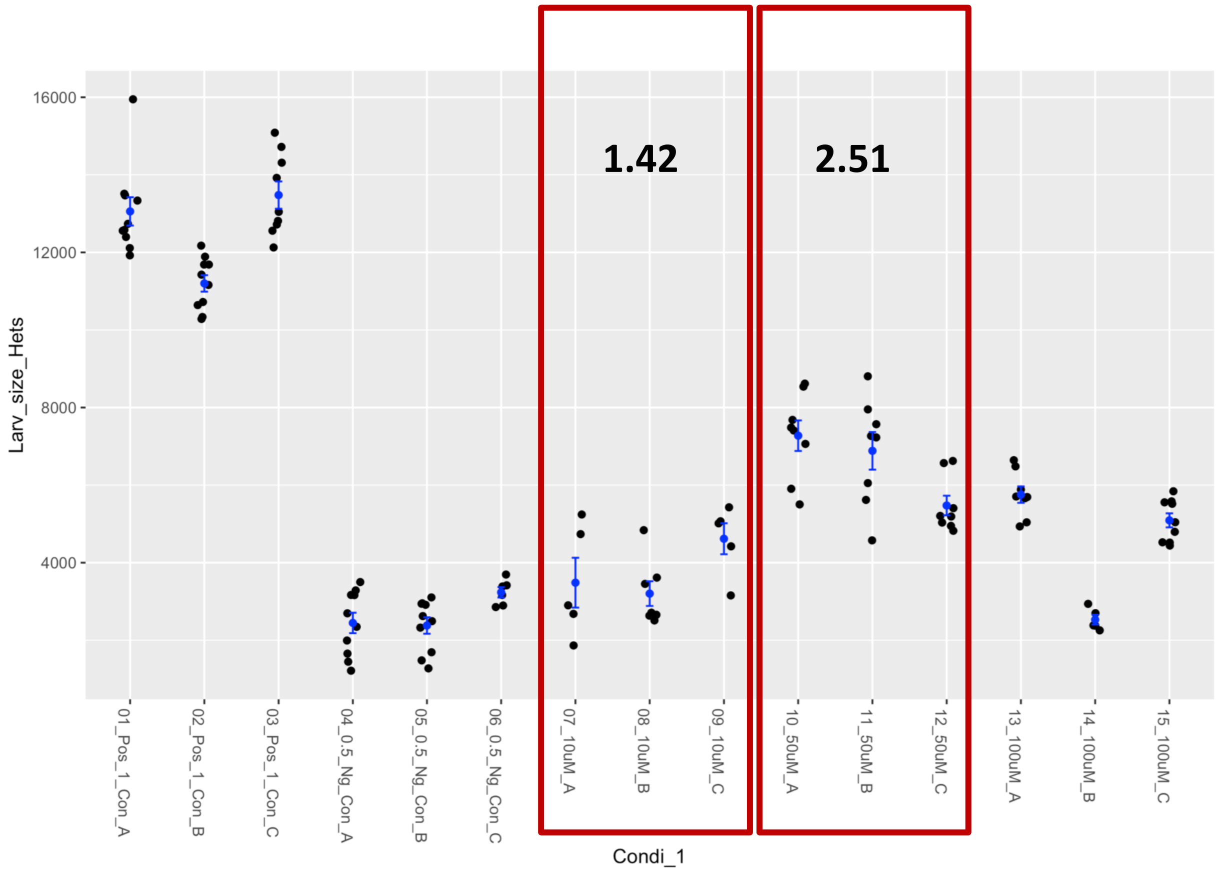

### Fly Chembridge retest – Supplier ID: 10343708

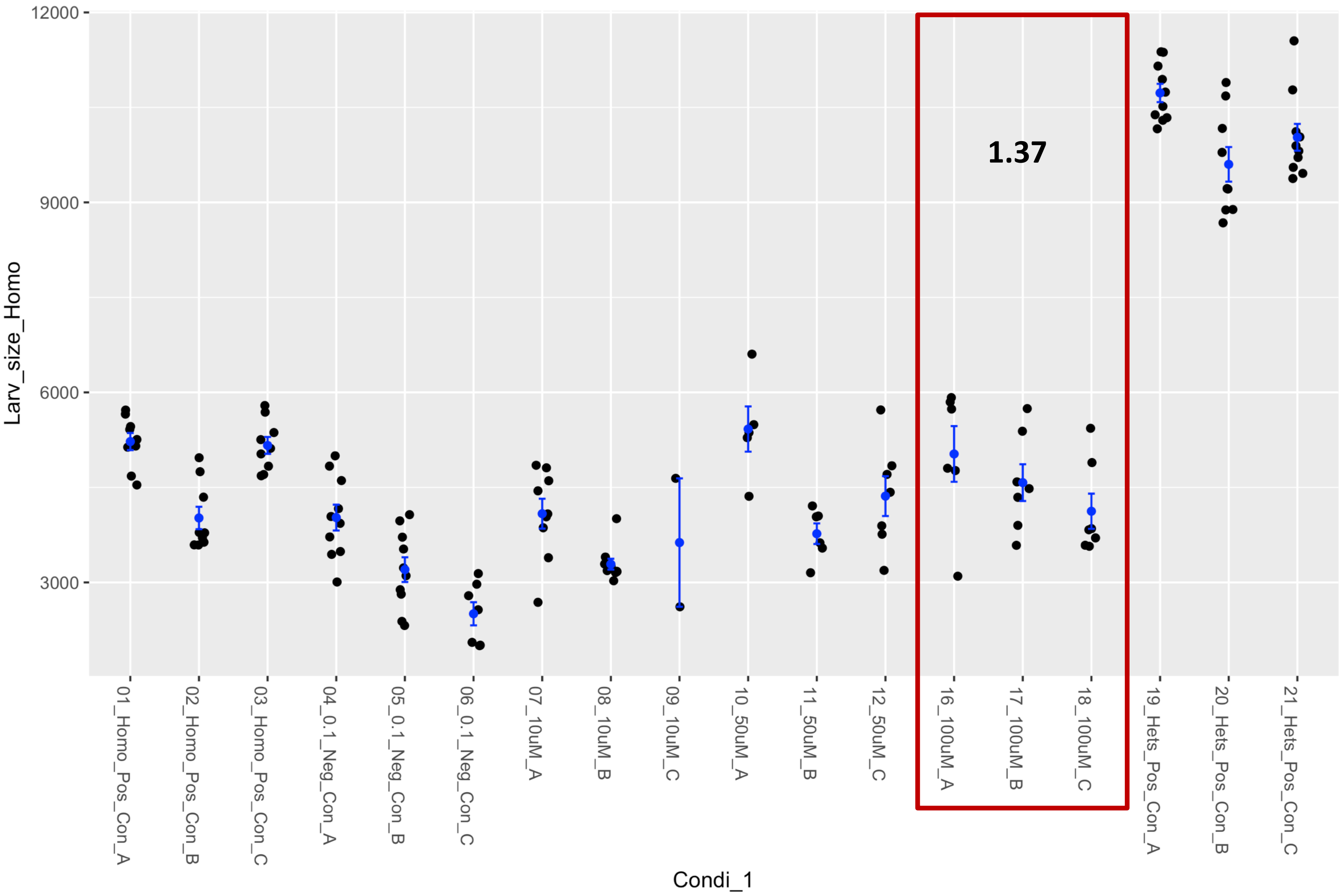


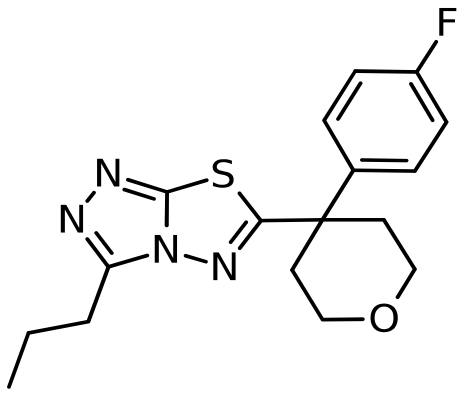

### Fly Chembridge retest – Supplier ID: 11646952

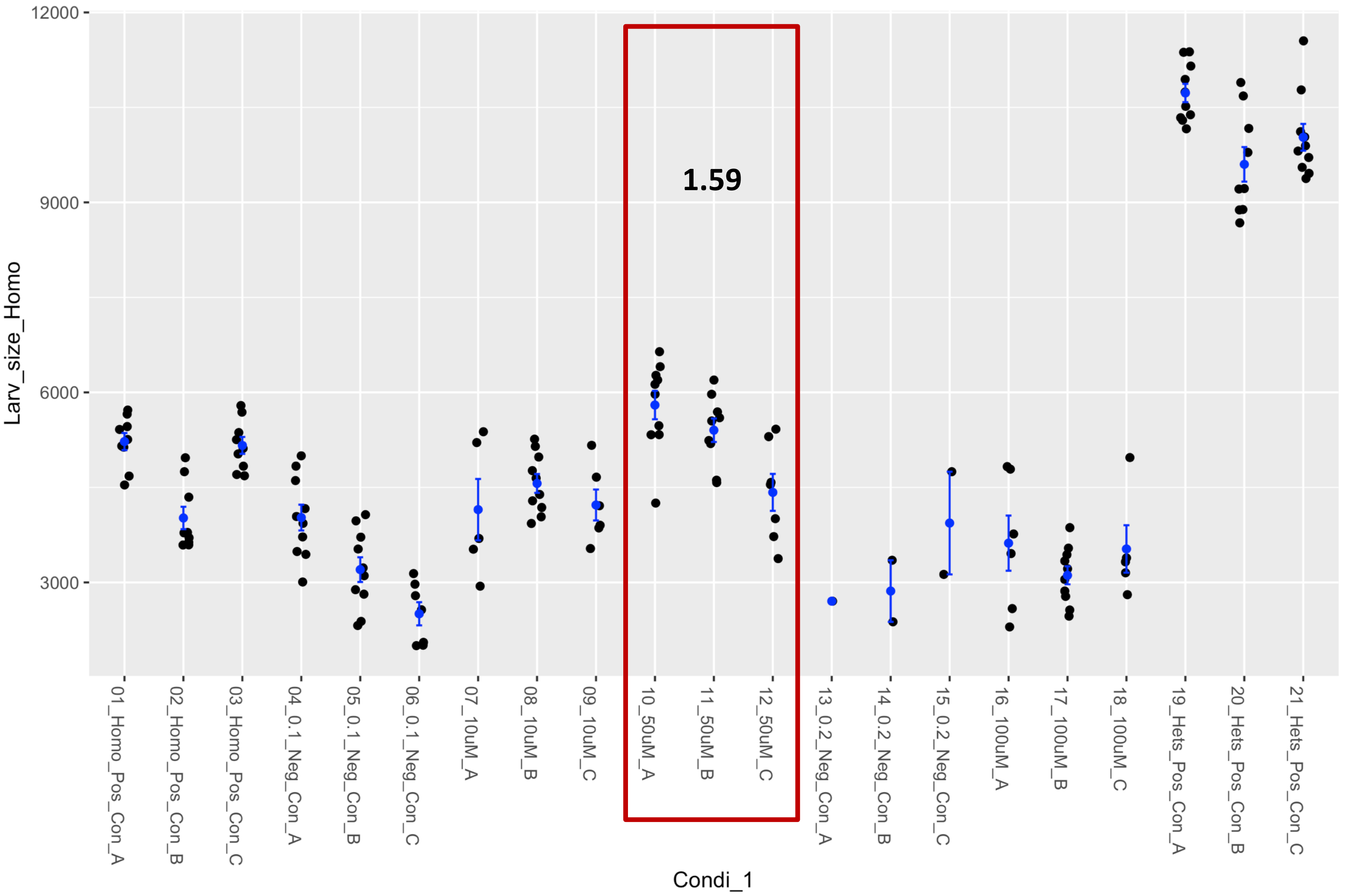


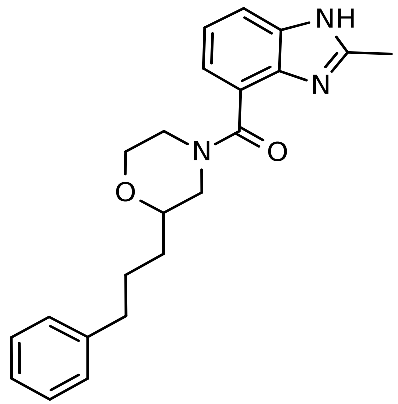

### Fly Chembridge retest – Supplier ID: 22934709

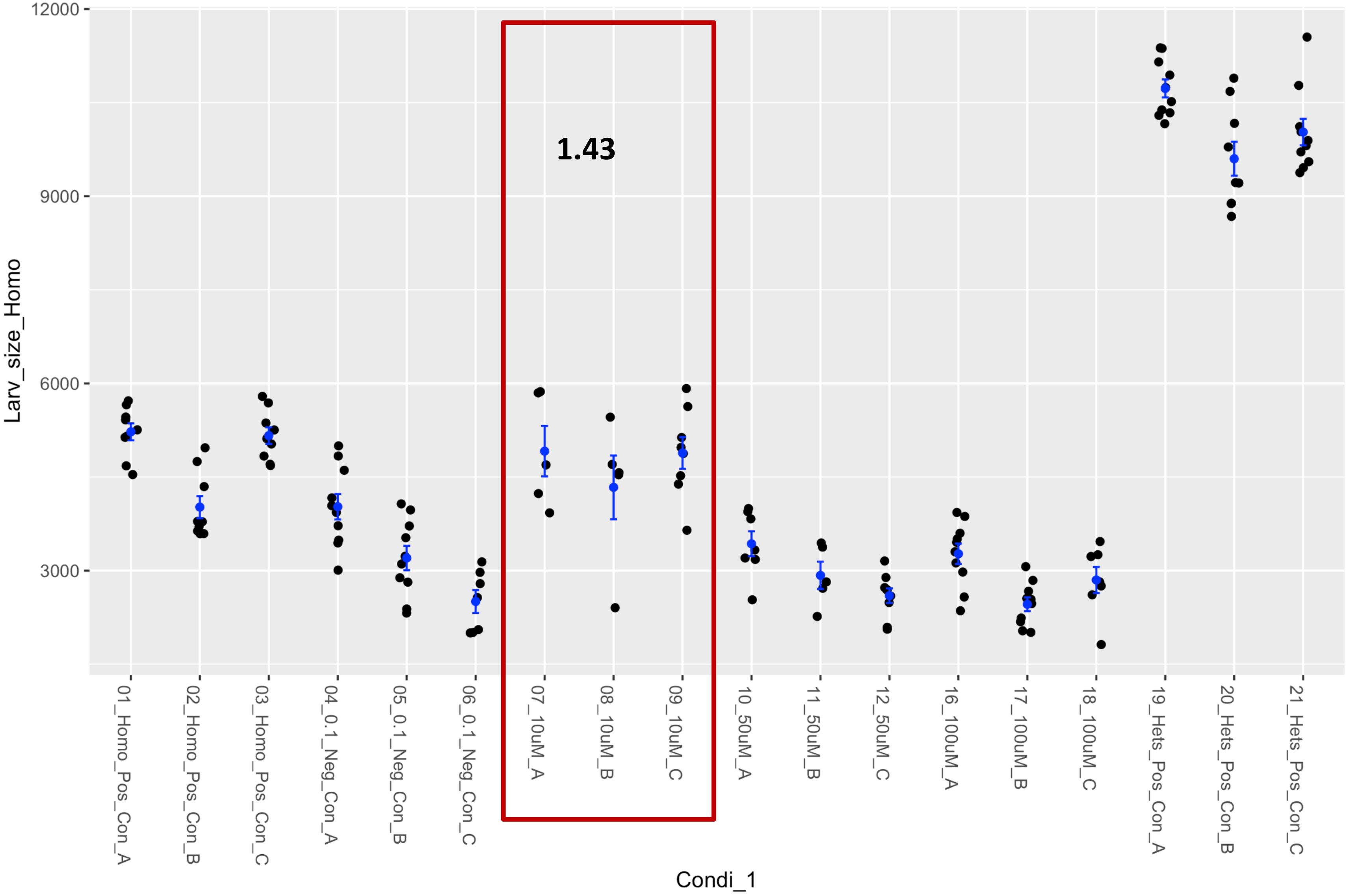


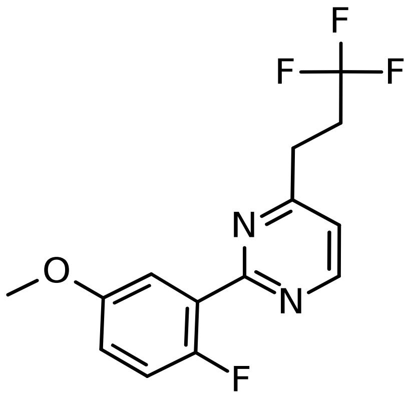

### Fly Chembridge retest – Supplier ID: 26889292

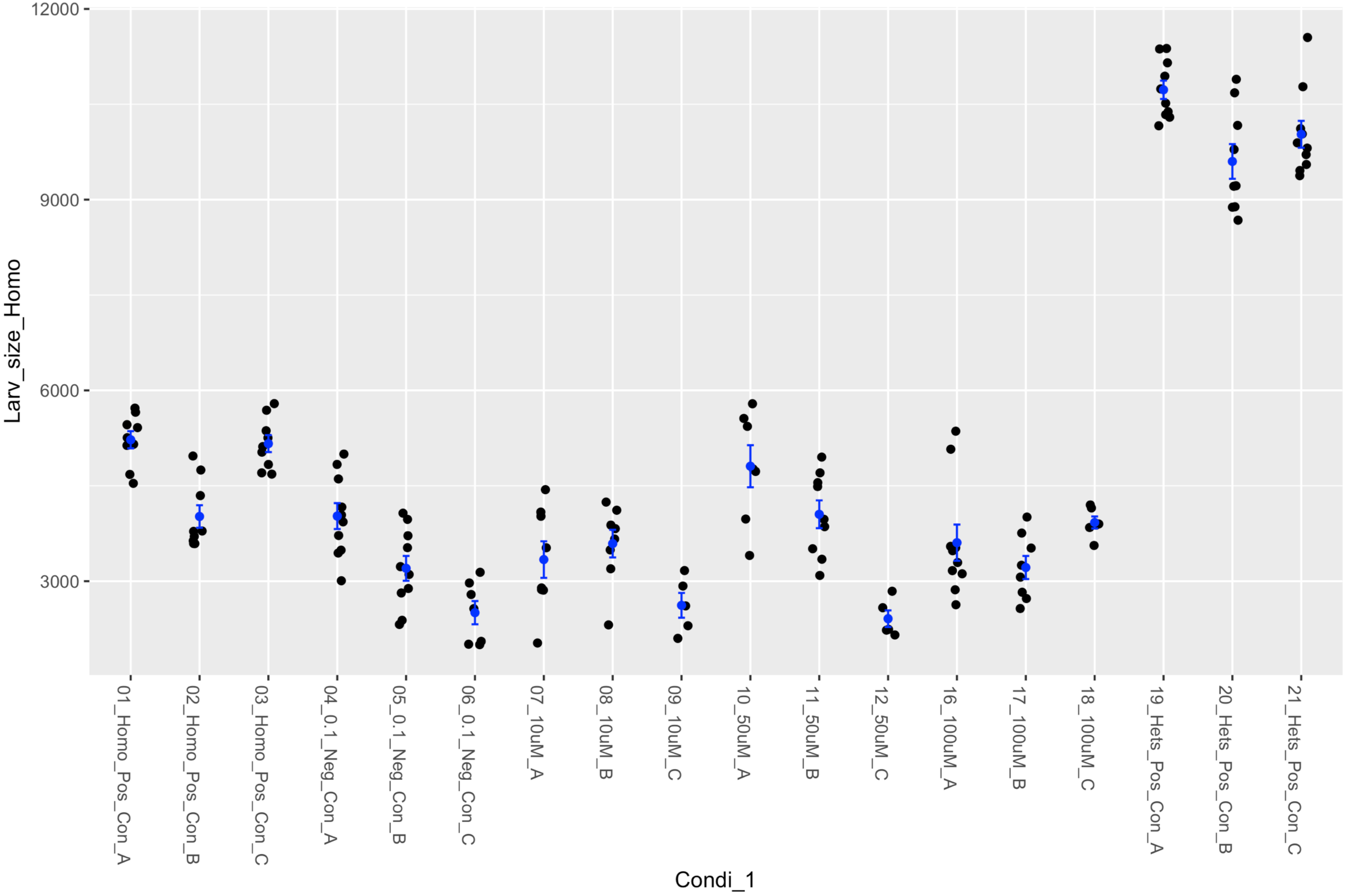


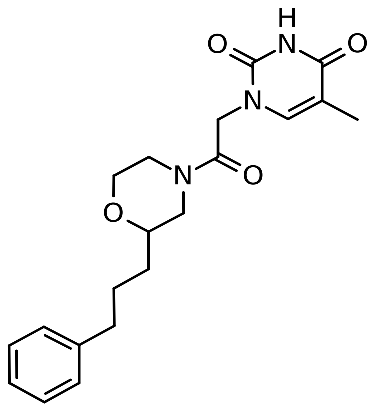

### Fly Chembridge retest – Supplier ID: 35527962

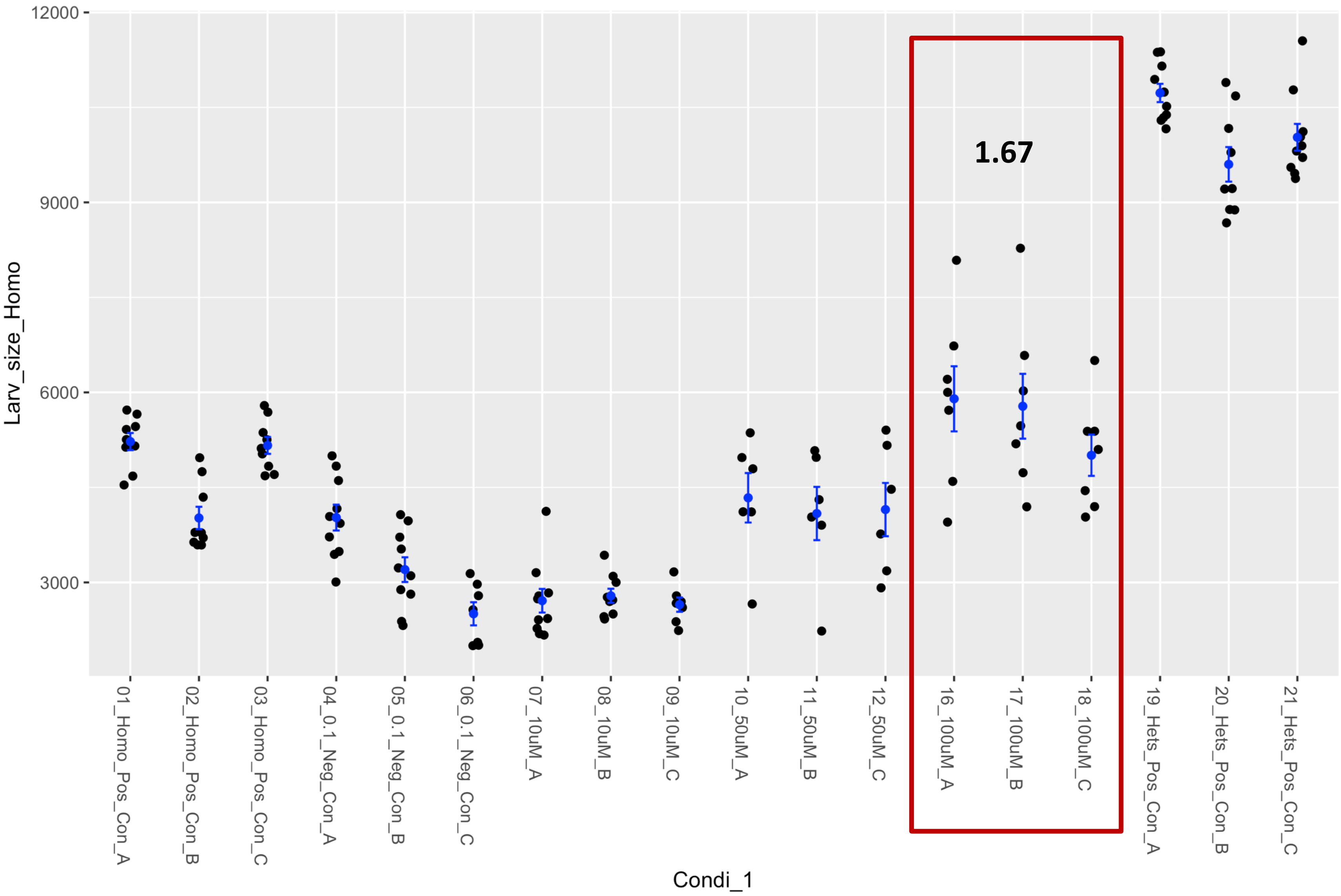


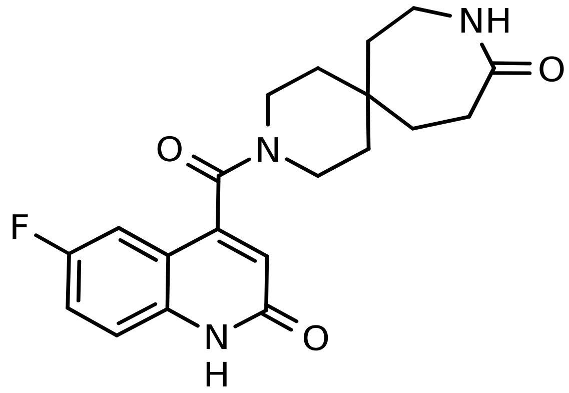

### Fly Chembridge retest – Supplier ID: 40966617

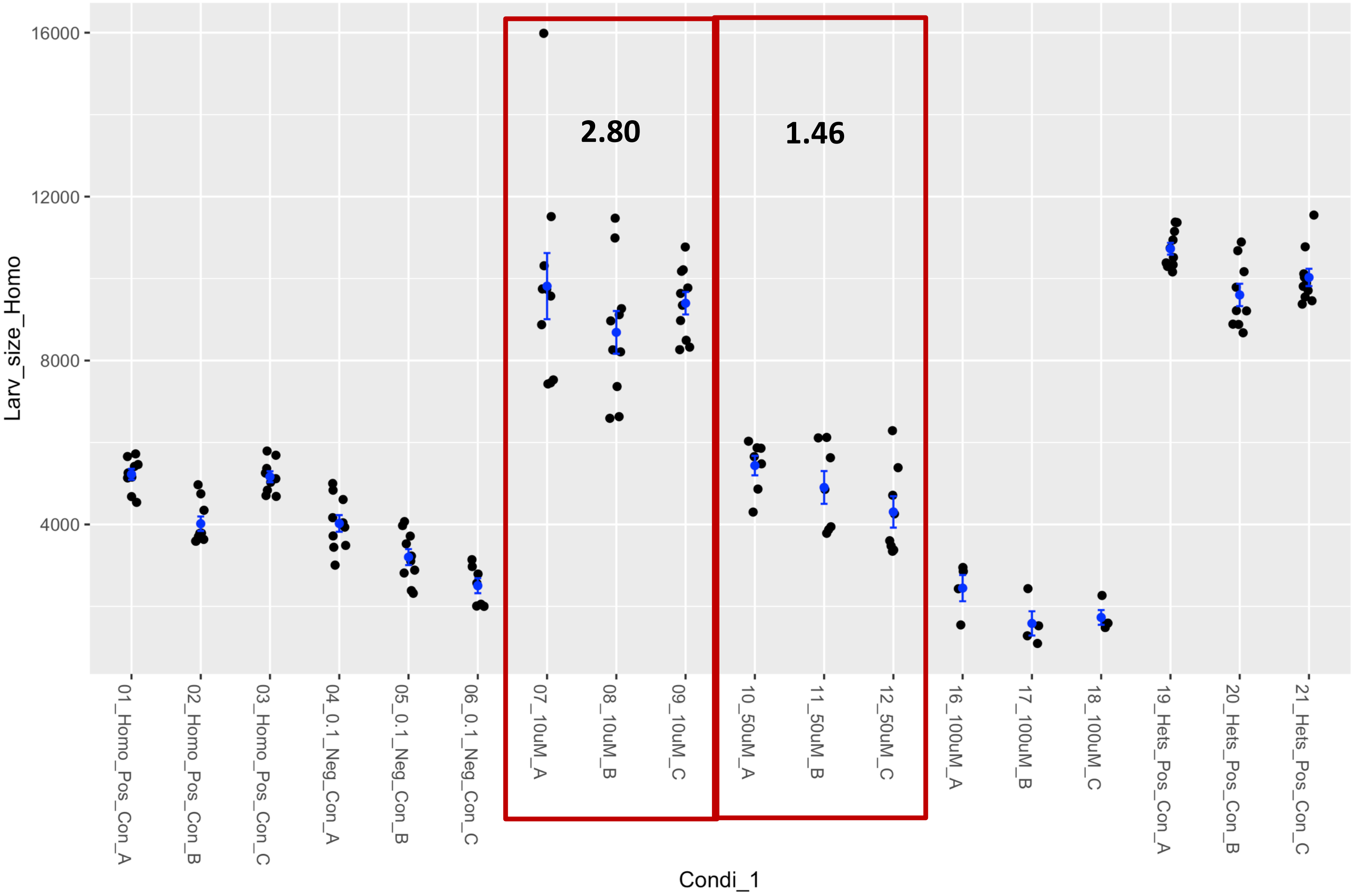


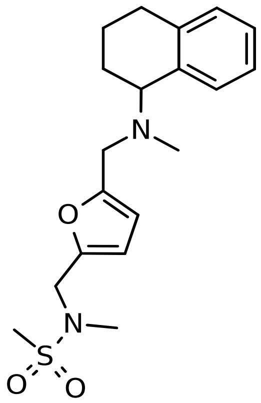

### Fly Chembridge retest – Supplier ID: 58095314

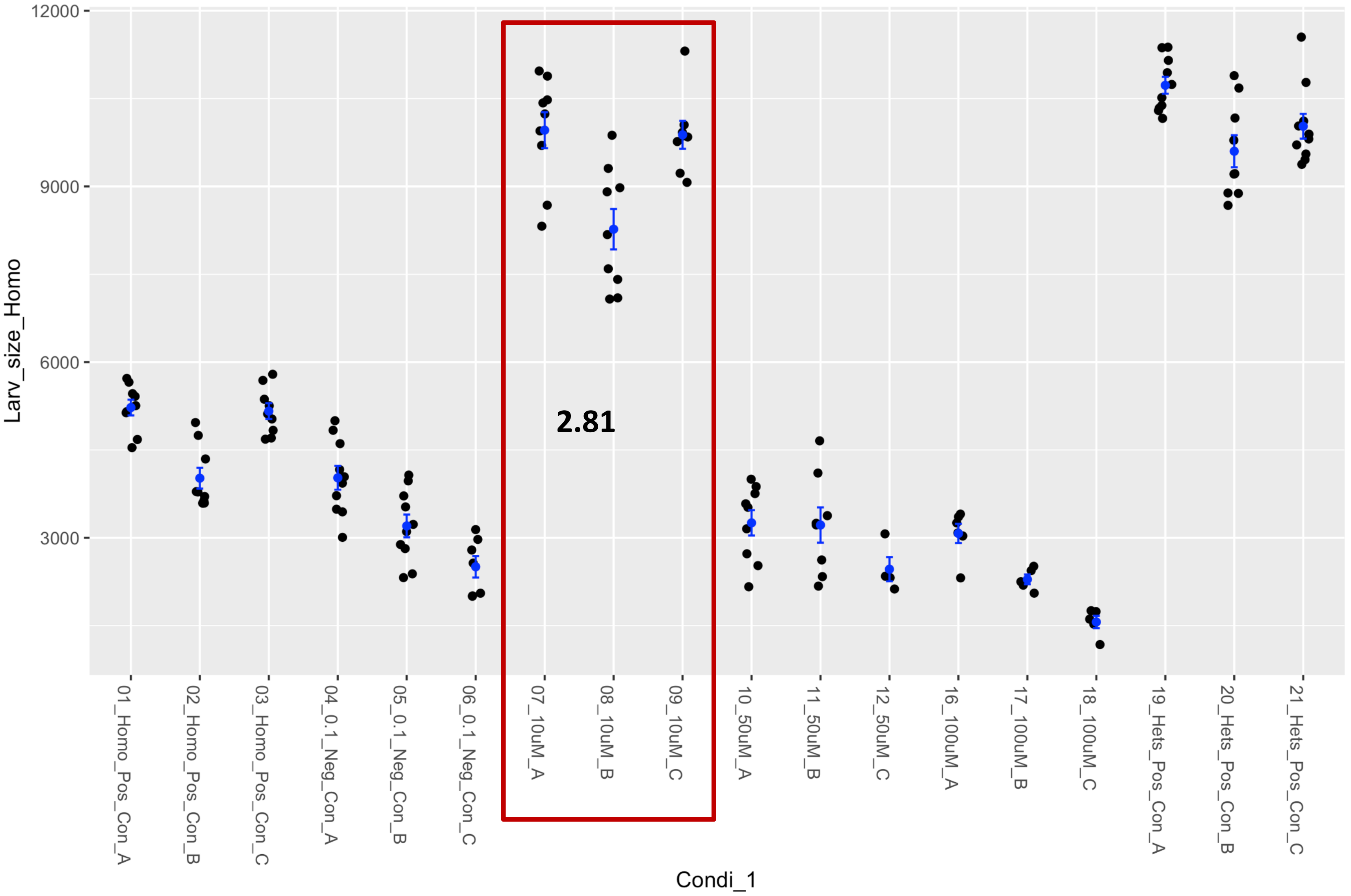


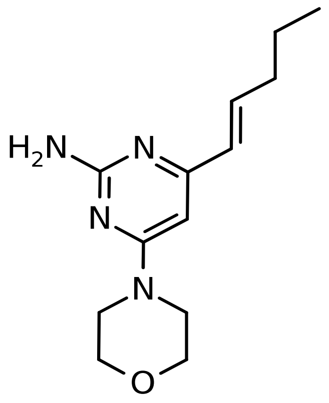

### Fly Chembridge retest – Supplier ID: 89890646

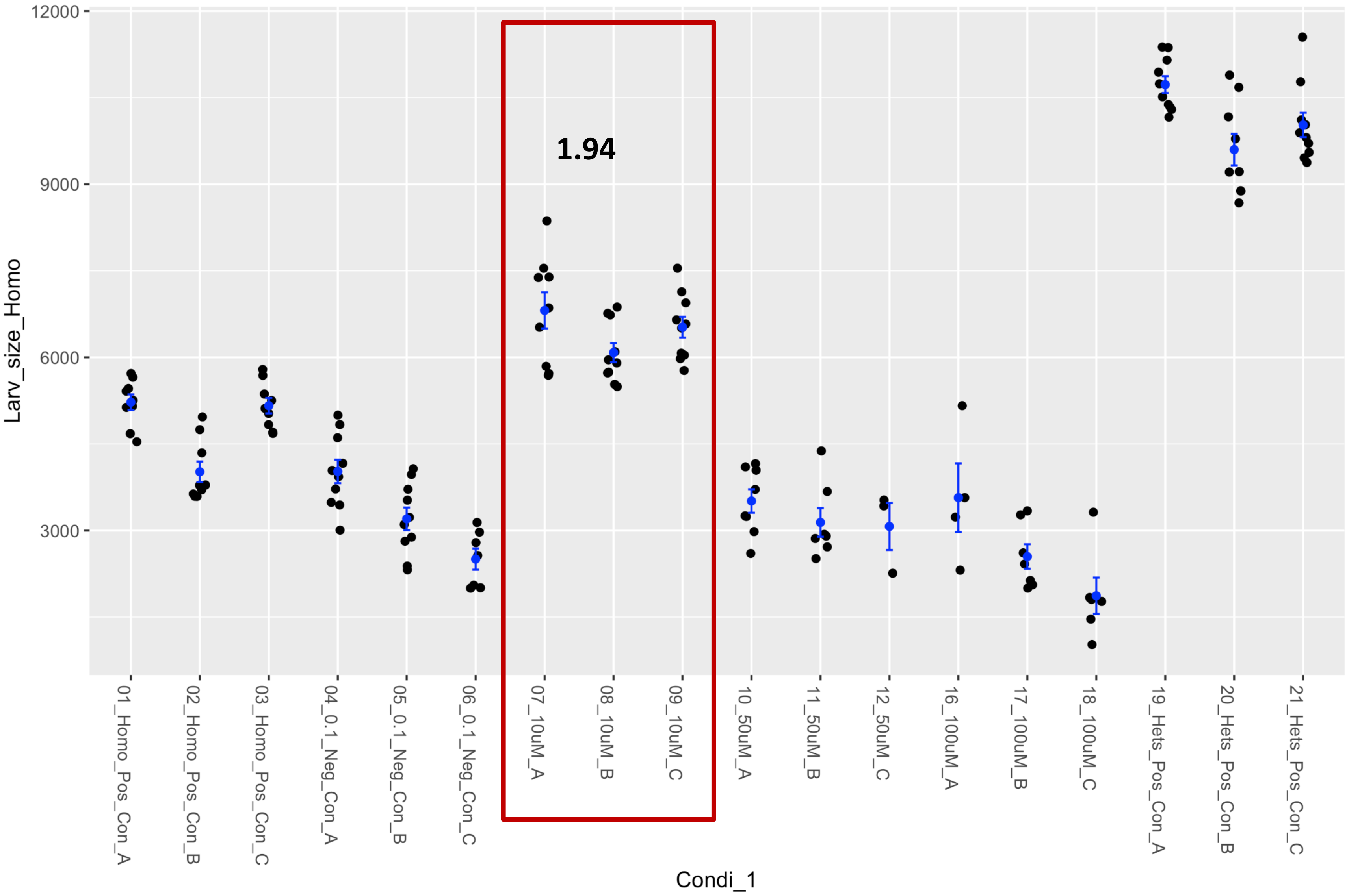


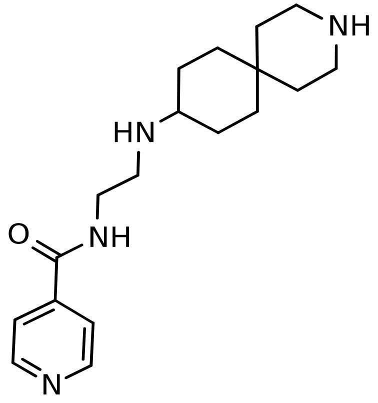

### Fly Chembridge retest – Supplier ID: 9206326

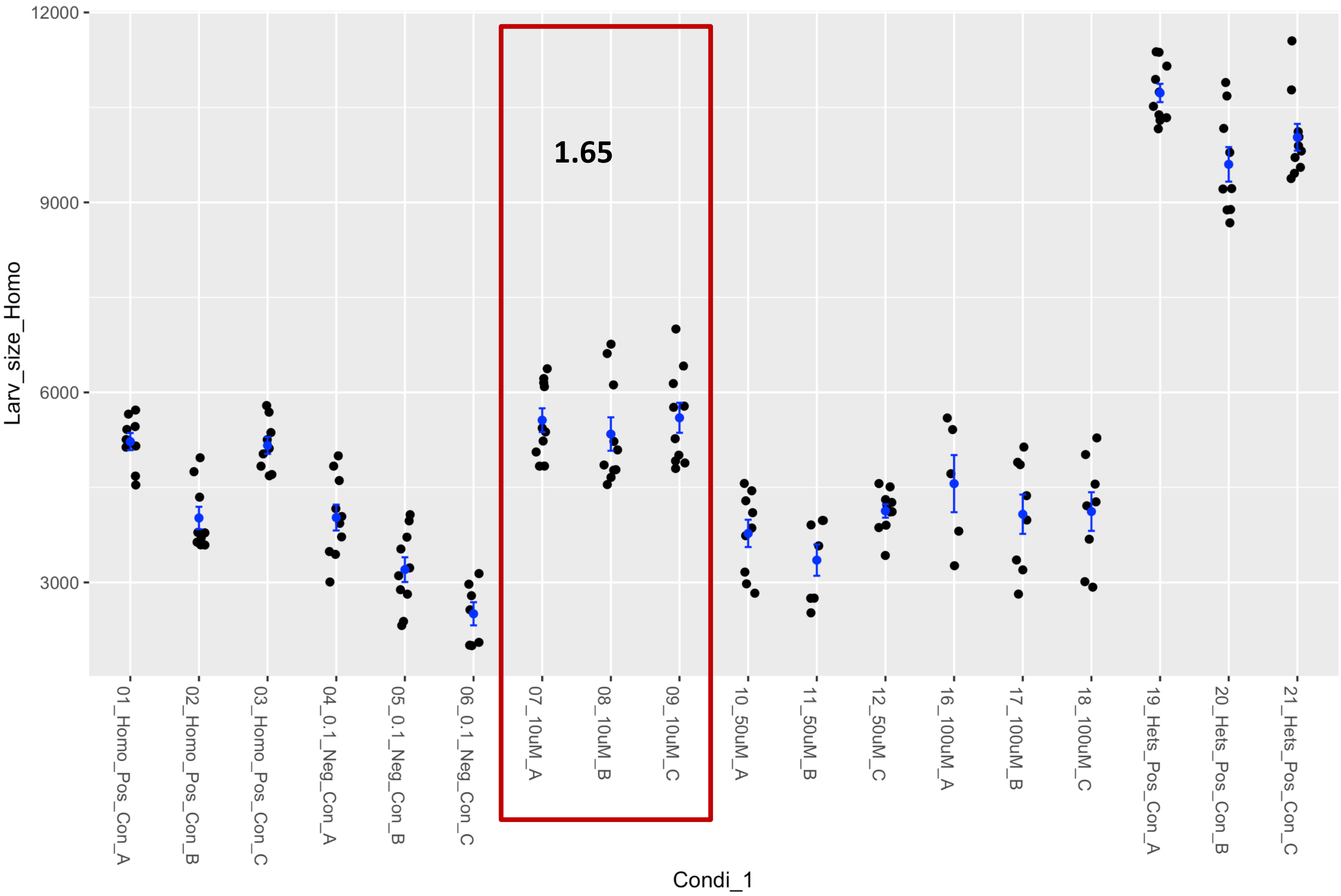


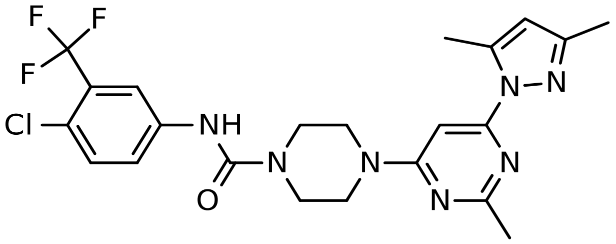

### Fly Chembridge retest – Supplier ID: 9233974

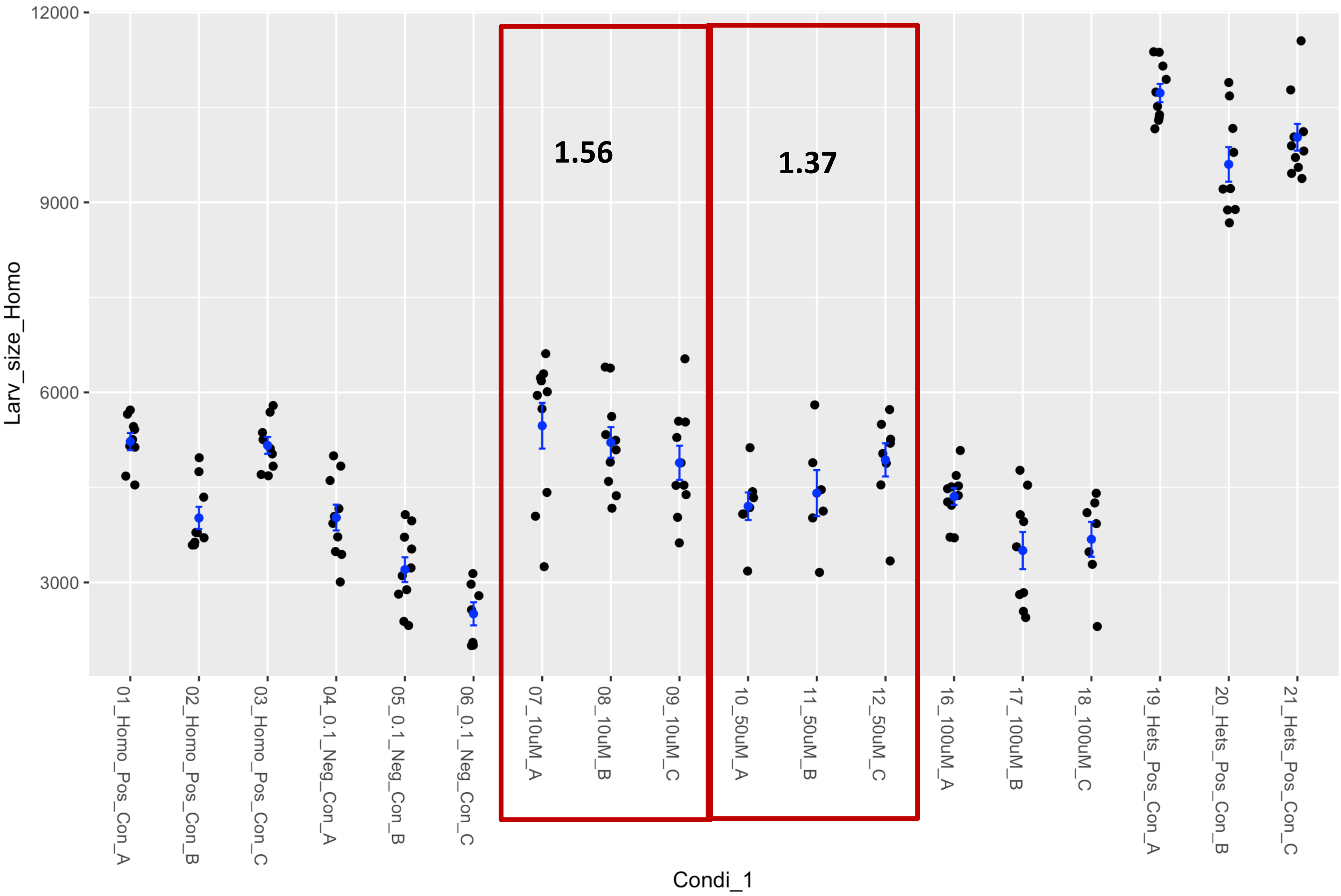


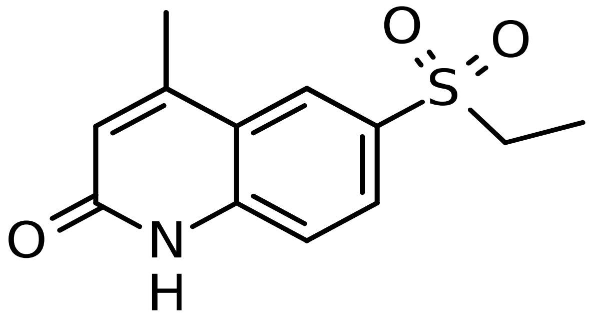

### Fly Chembridge retest – Supplier ID: 9251119

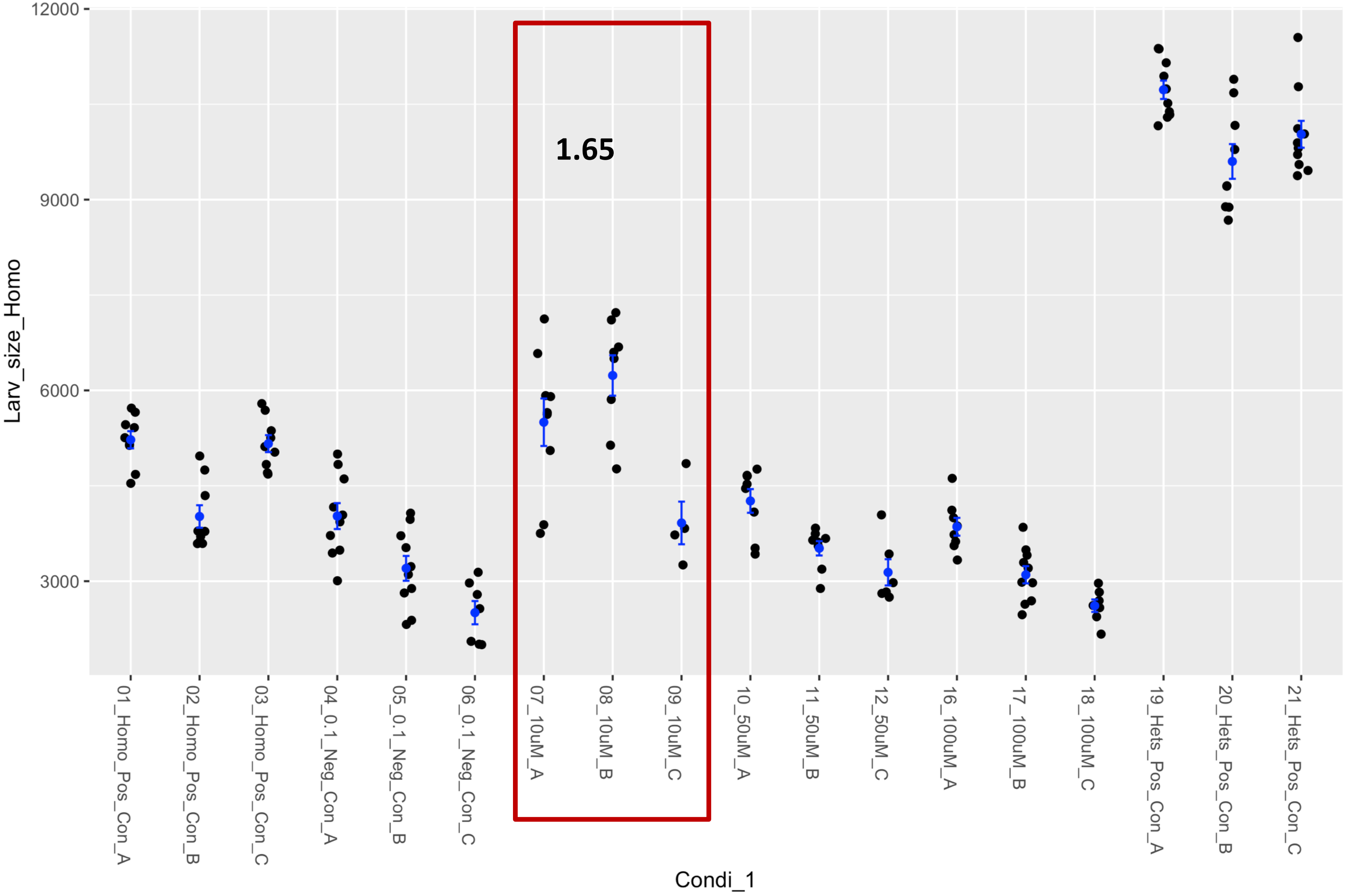


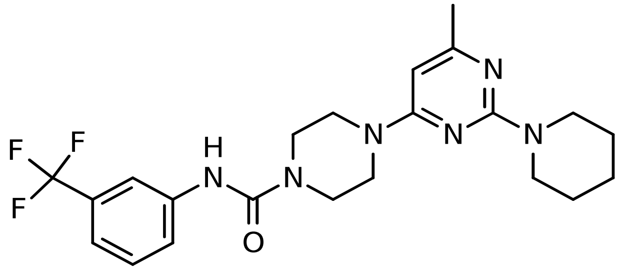

### Fly Chembridge retest – Supplier ID: 9251633

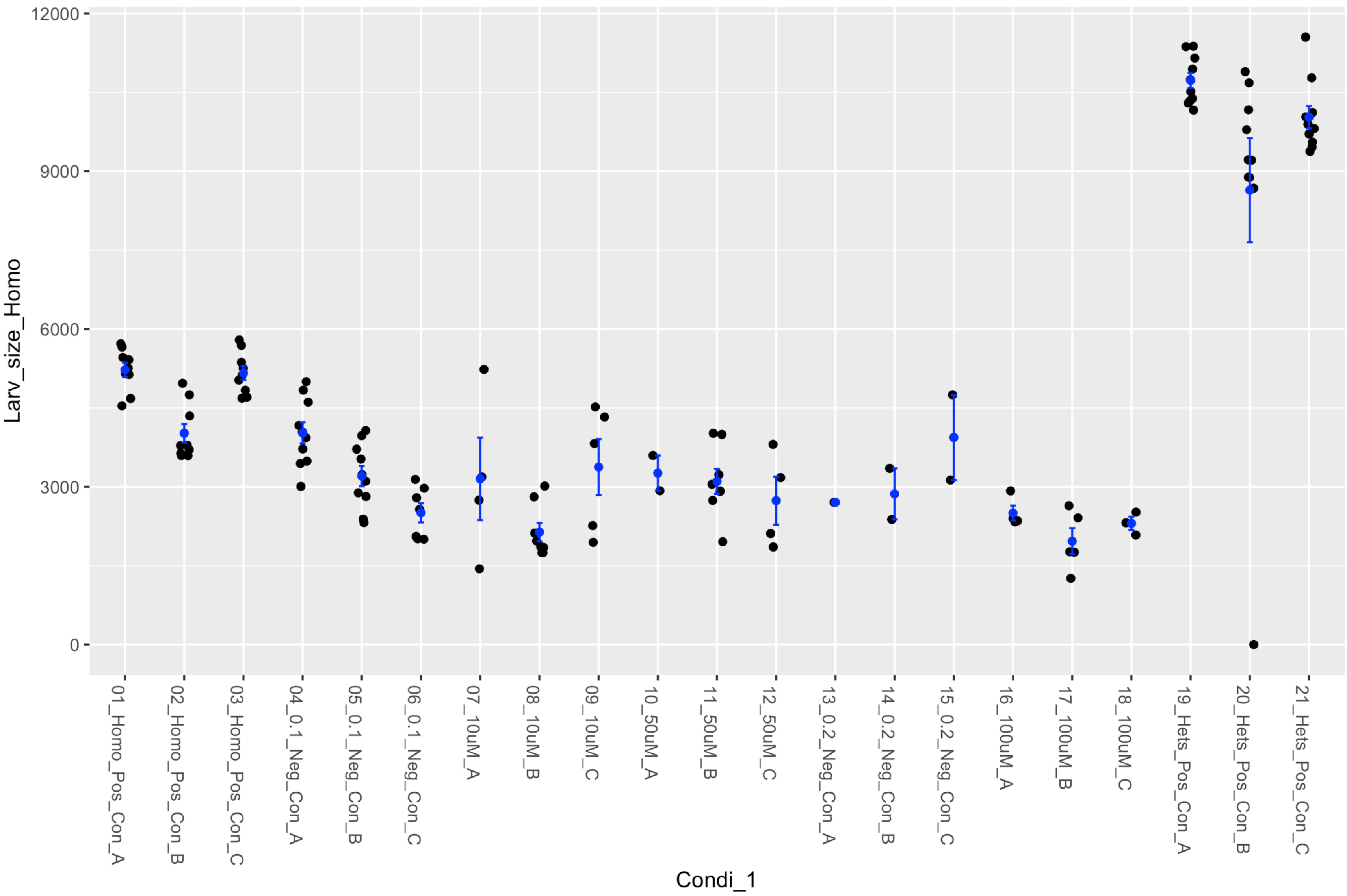


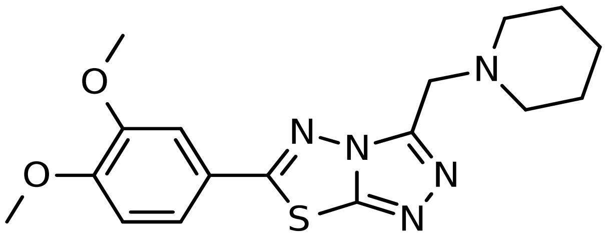

### Fly Chembridge retest – Supplier ID: 9262220

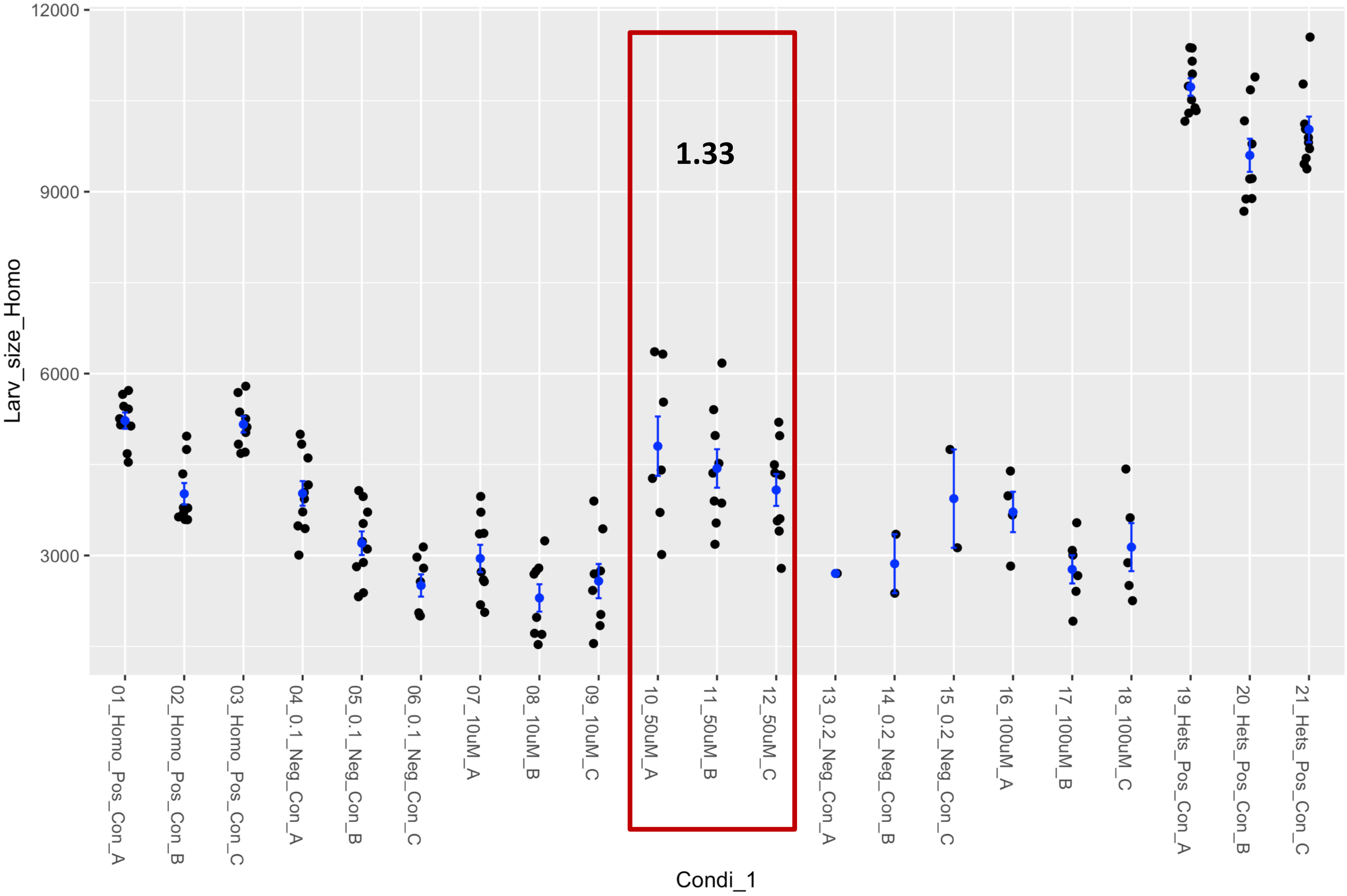


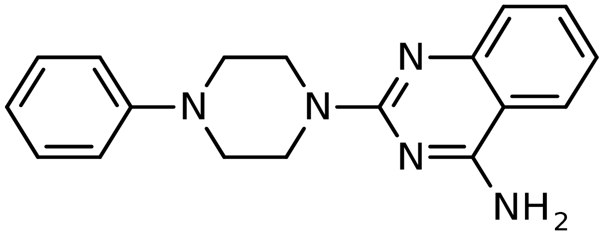

### Fly Chembridge retest – Supplier ID: 9262955

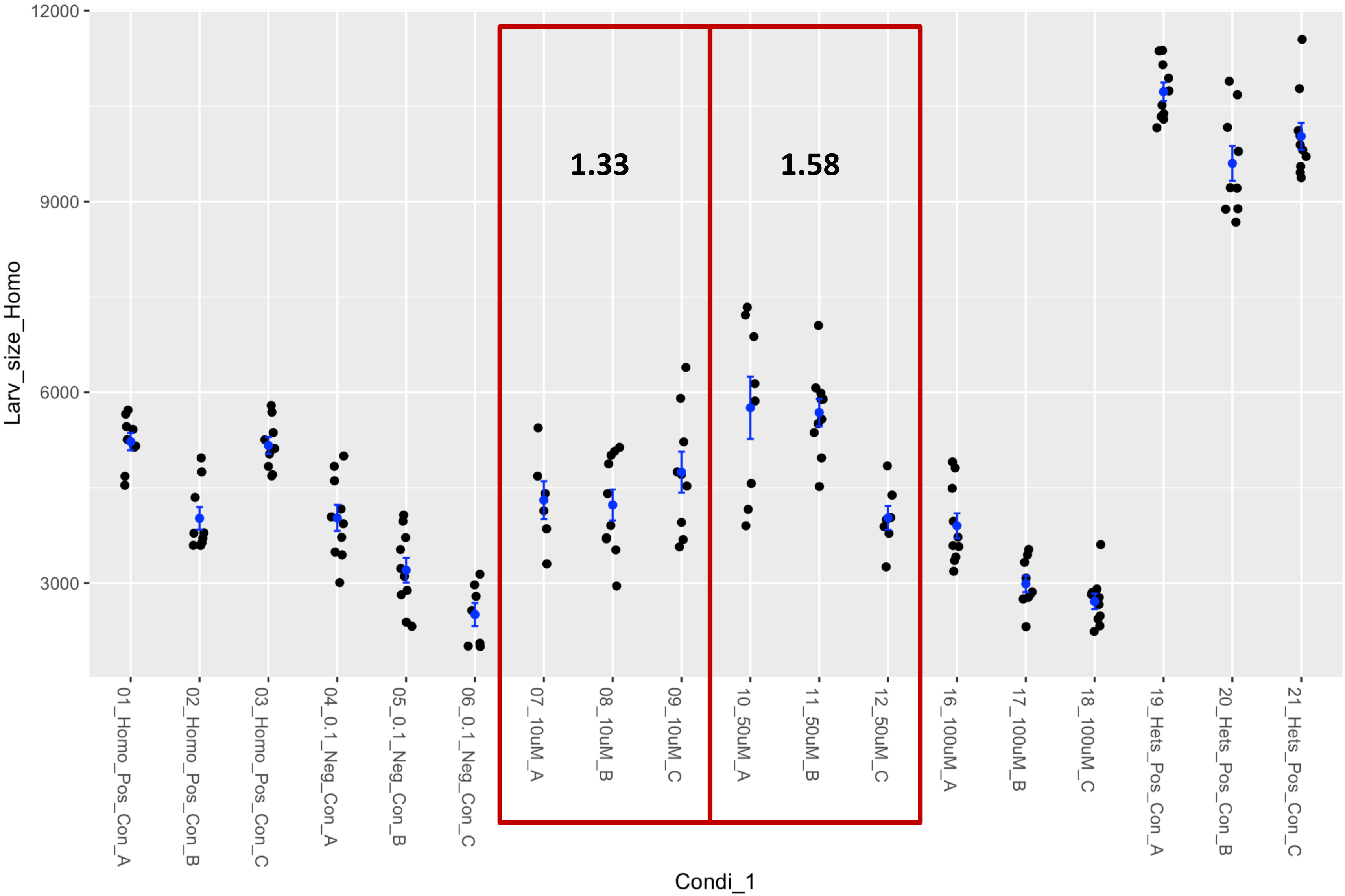


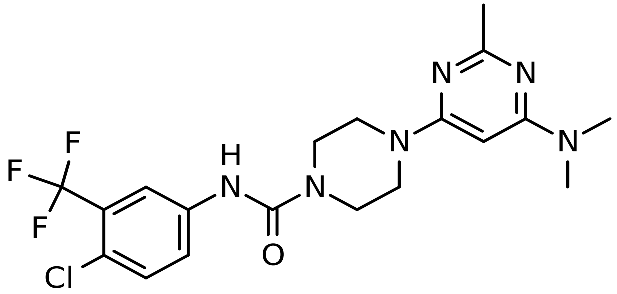

### Fly Chembridge retest – Supplier ID: 9290684

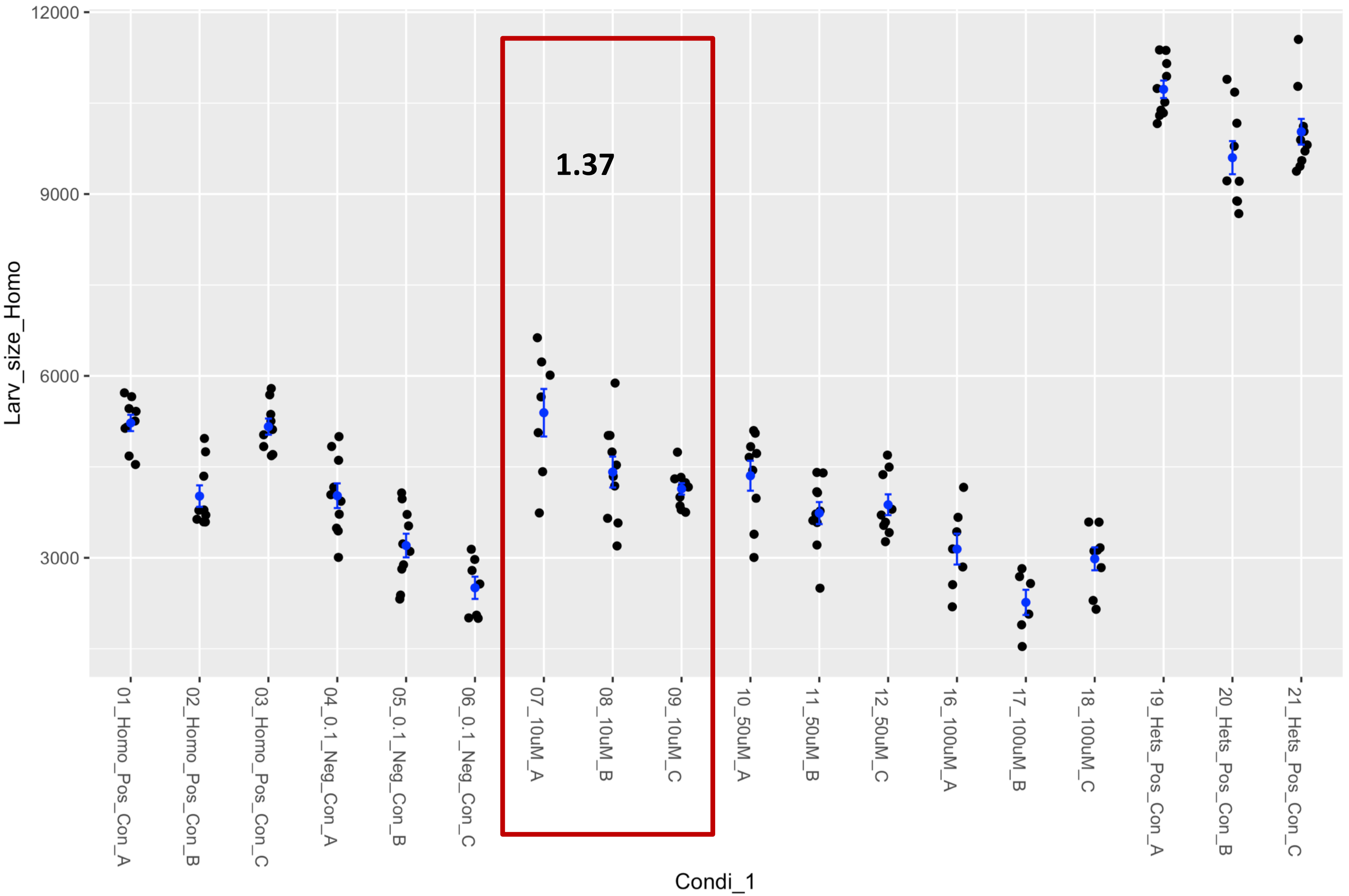


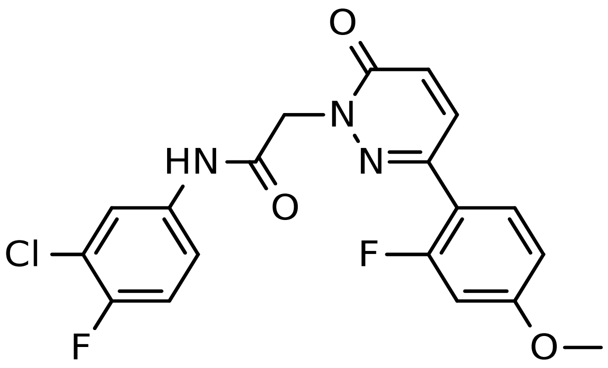

### Fly Chembridge retest – Supplier ID: 10343708

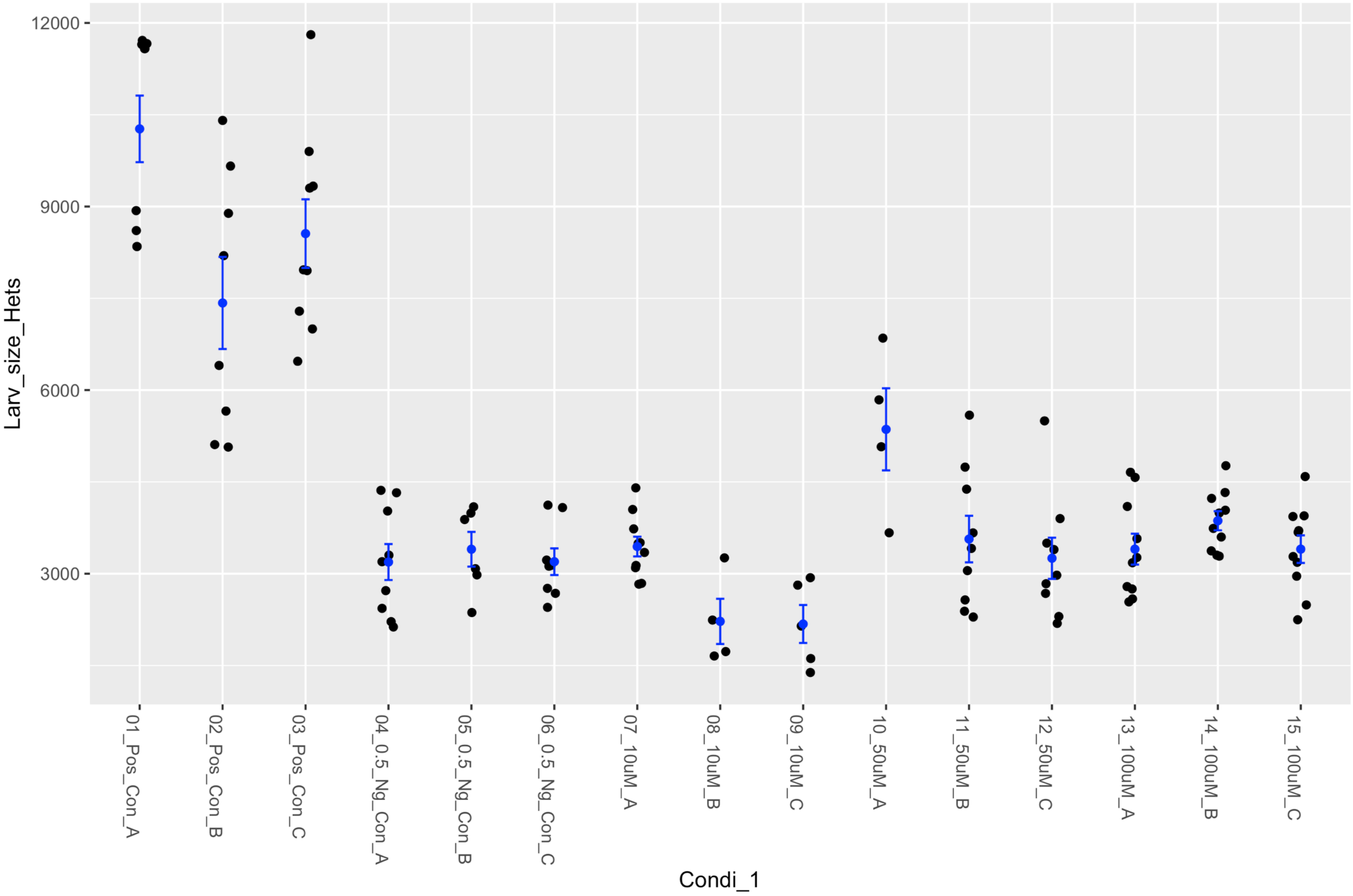


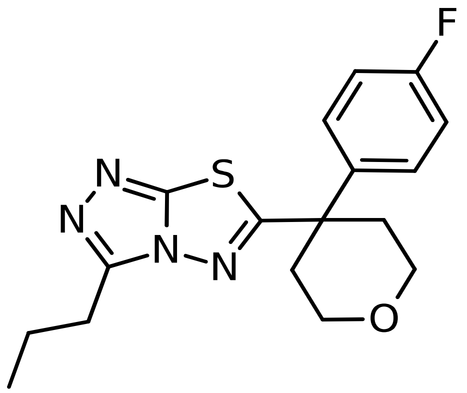

### Fly Chembridge retest – Supplier ID: 11646952

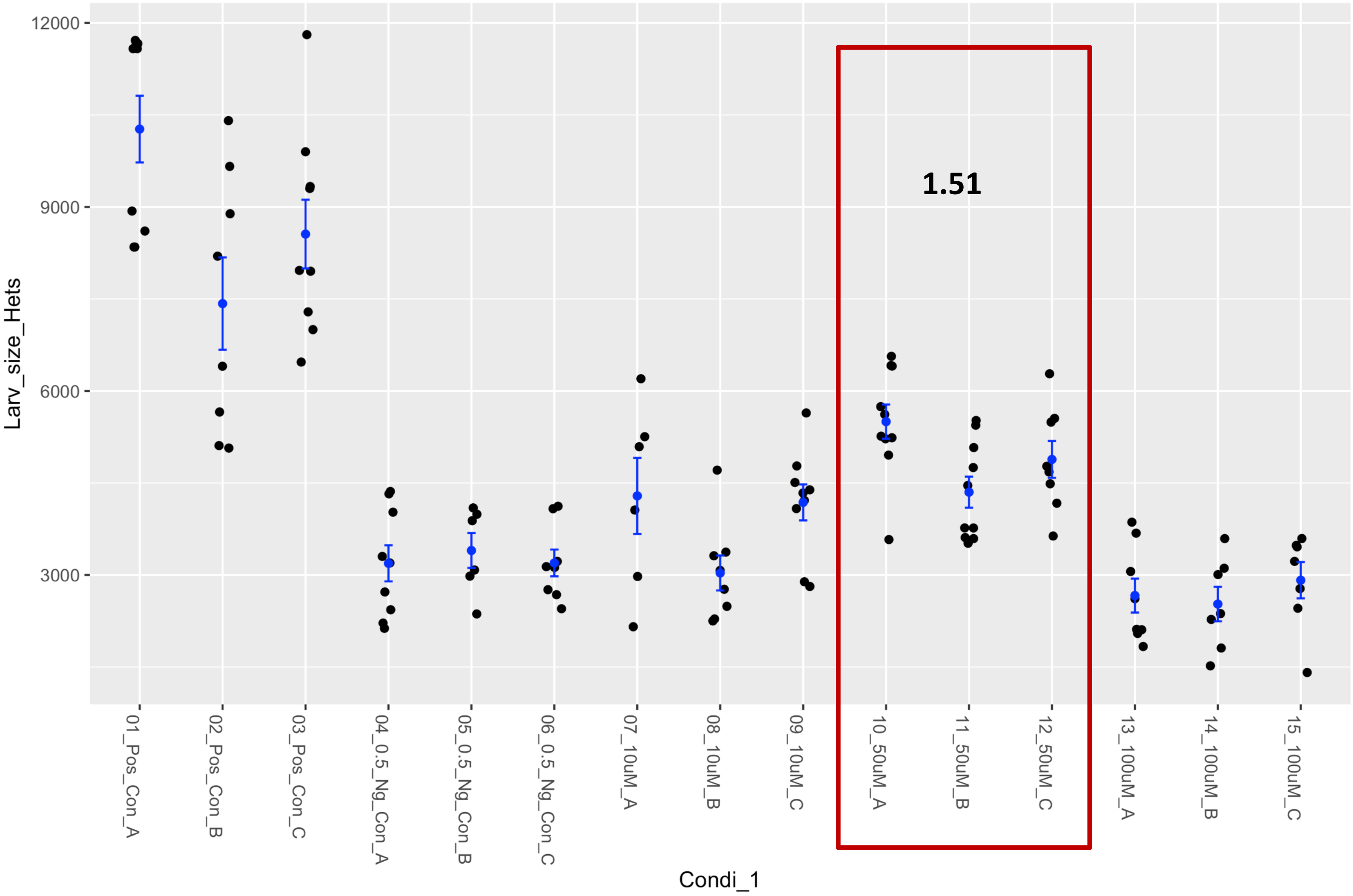


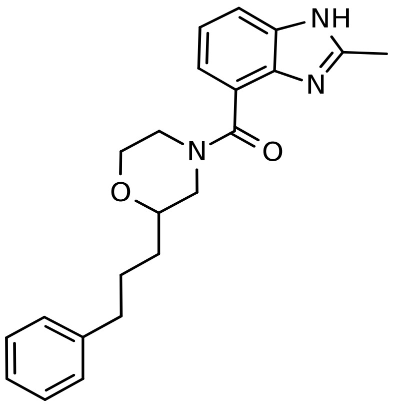

### Fly Chembridge retest – Supplier ID: 22934709

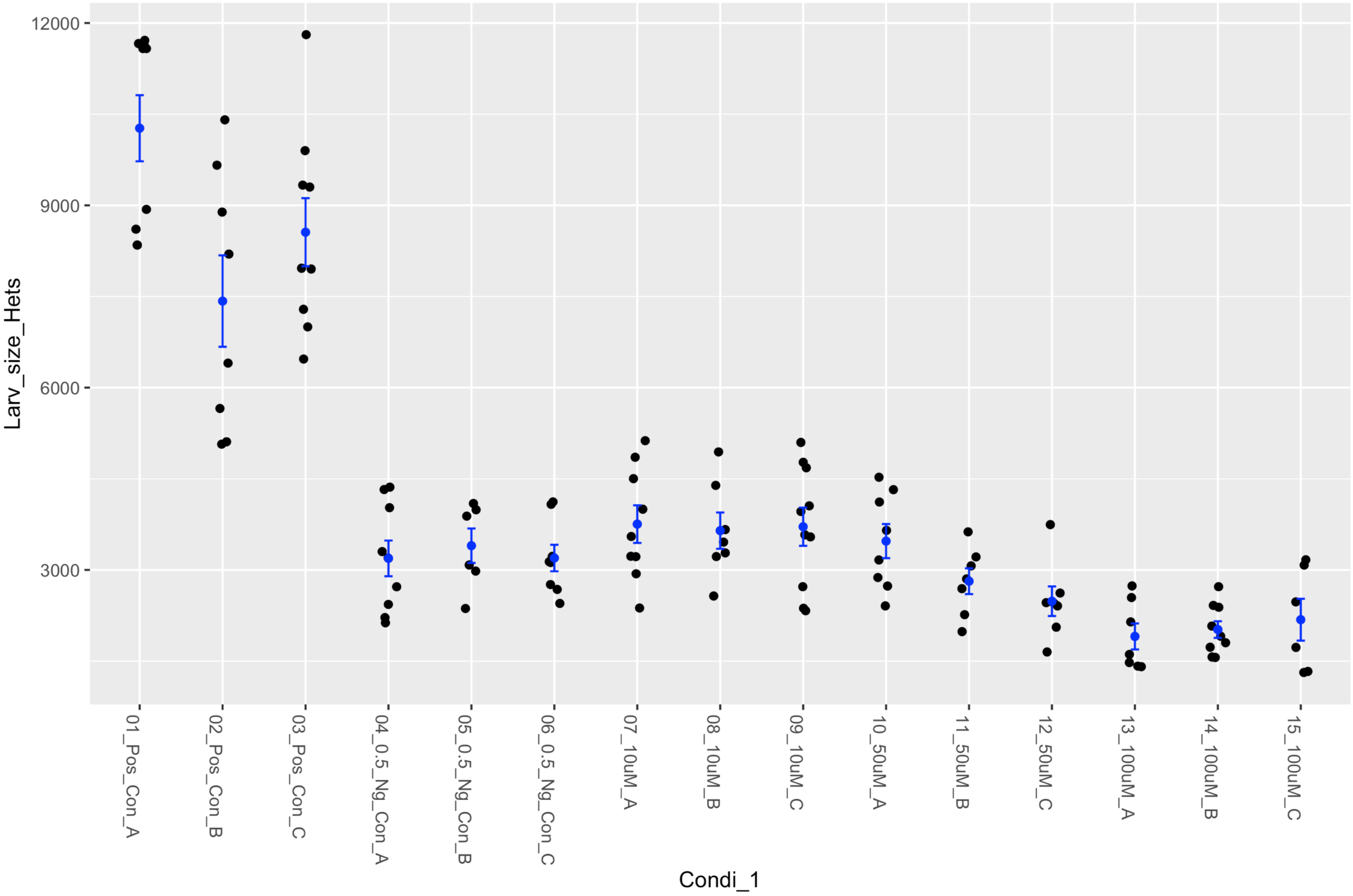


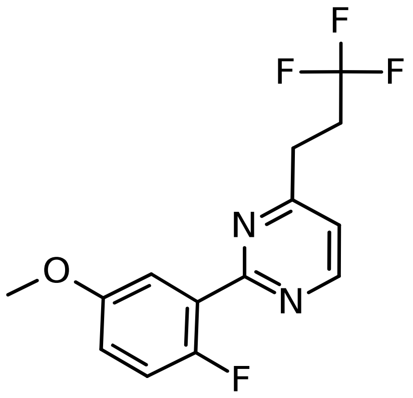

### Fly Chembridge retest – Supplier ID: 26889292

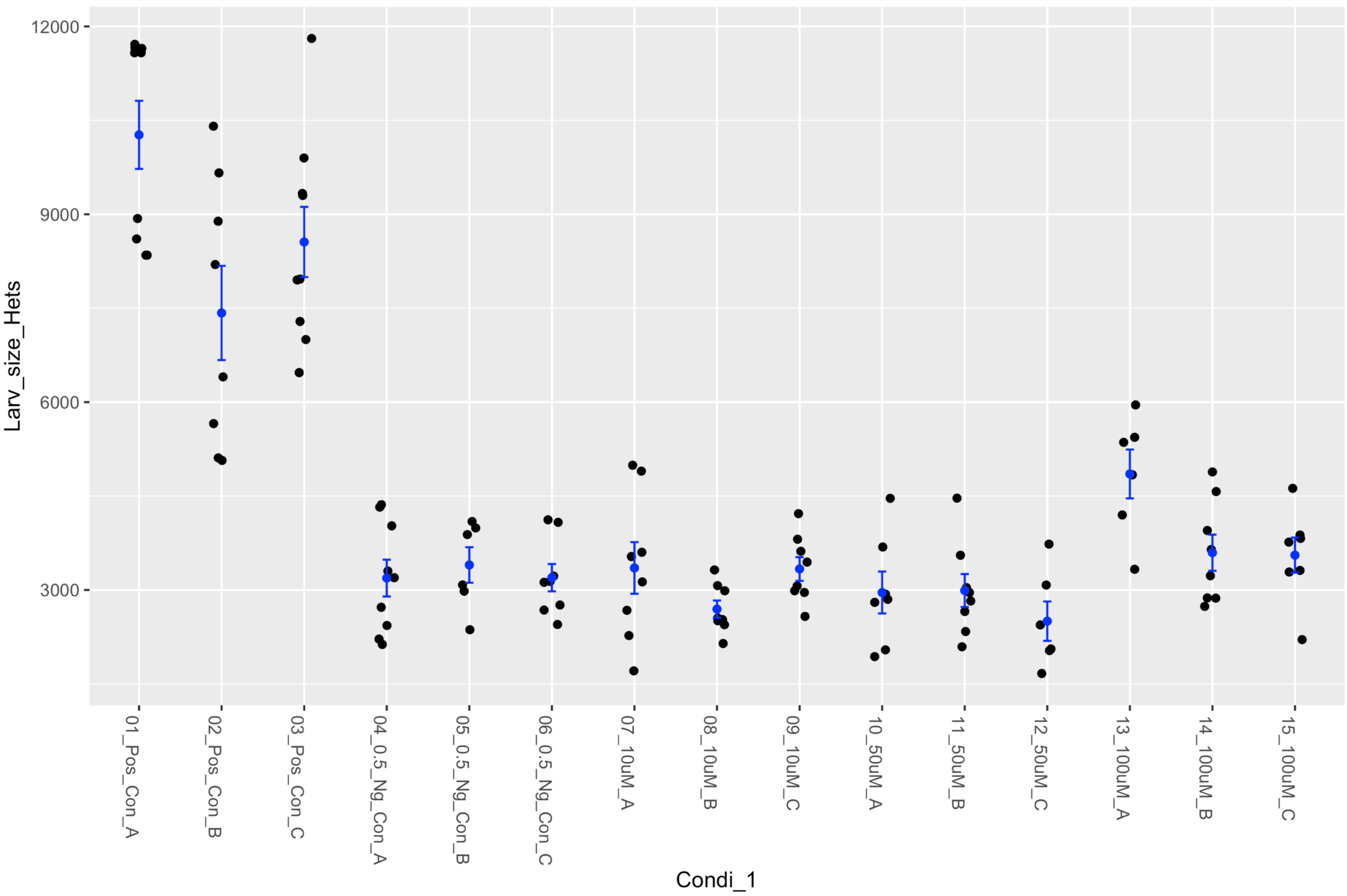


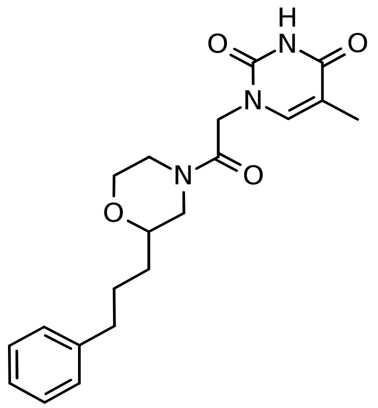

### Fly Chembridge retest – Supplier ID: 35527962

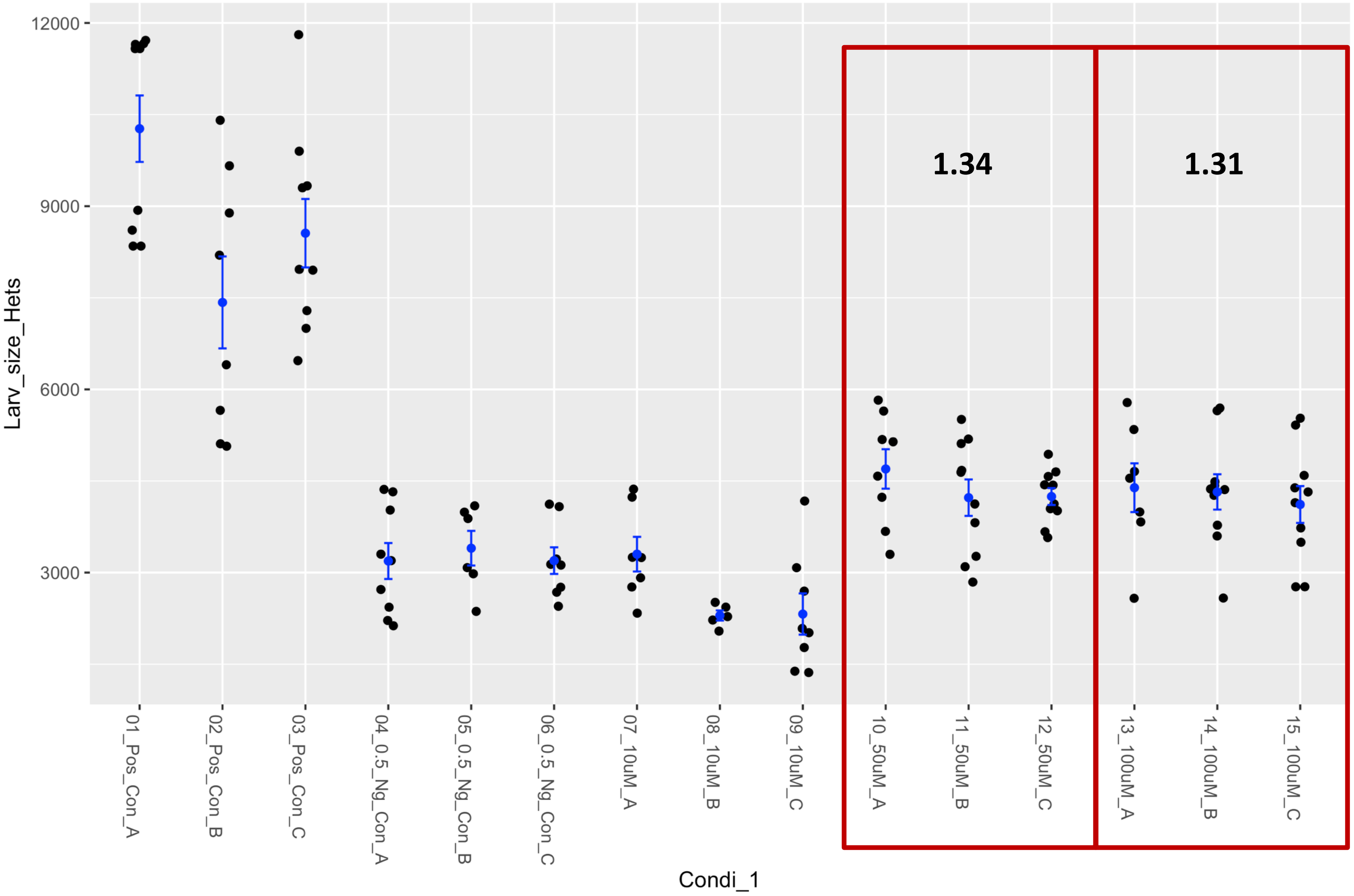


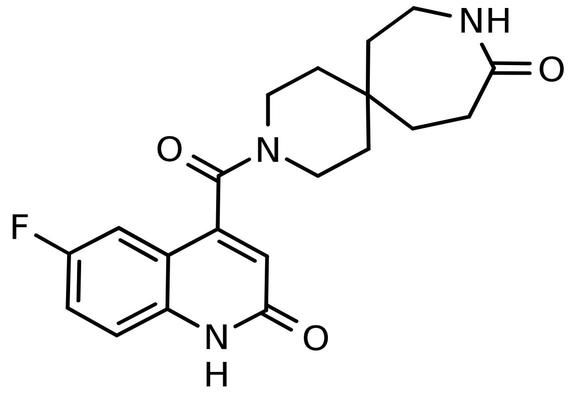

### Fly Chembridge retest – Supplier ID: 40966617

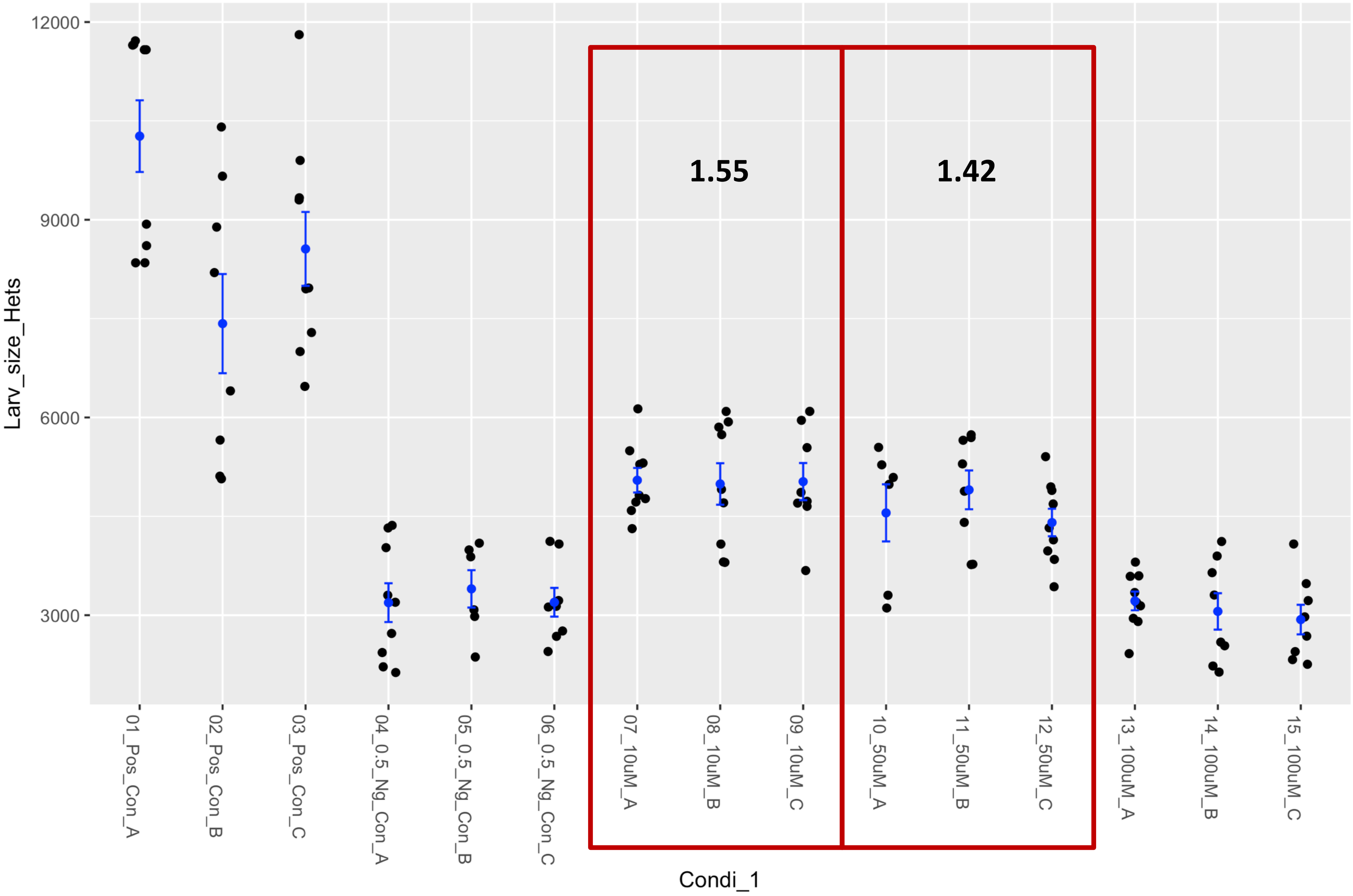


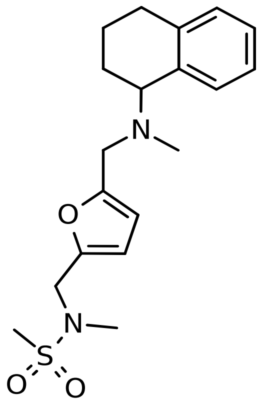

### Fly Chembridge retest – Supplier ID: 58095314

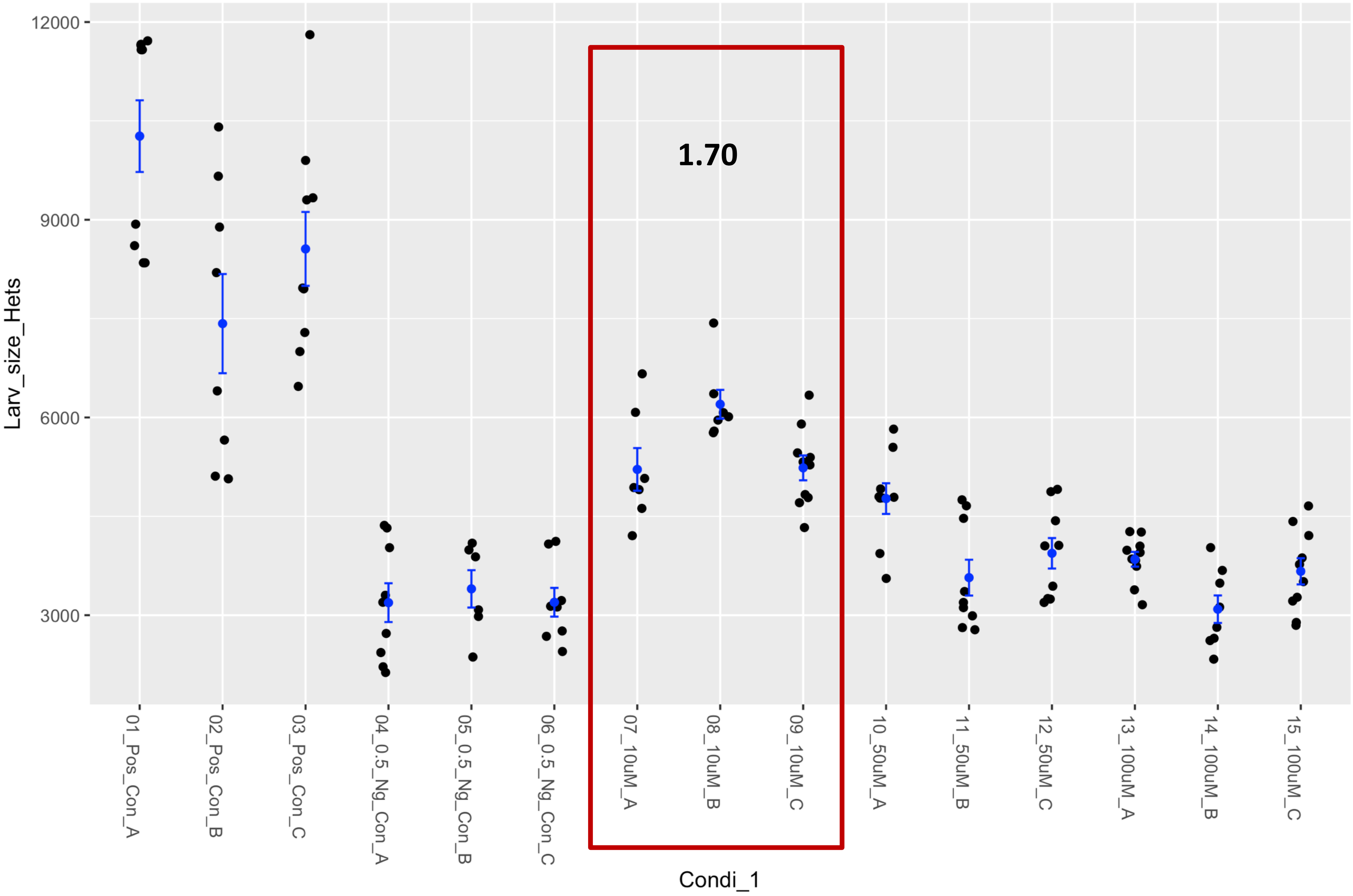


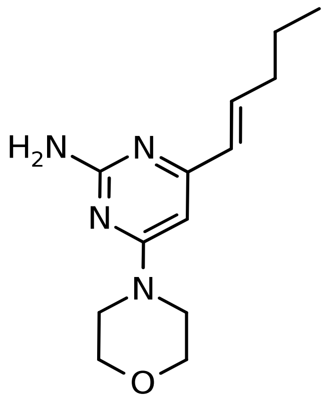

### Fly Chembridge retest – Supplier ID: 89890646

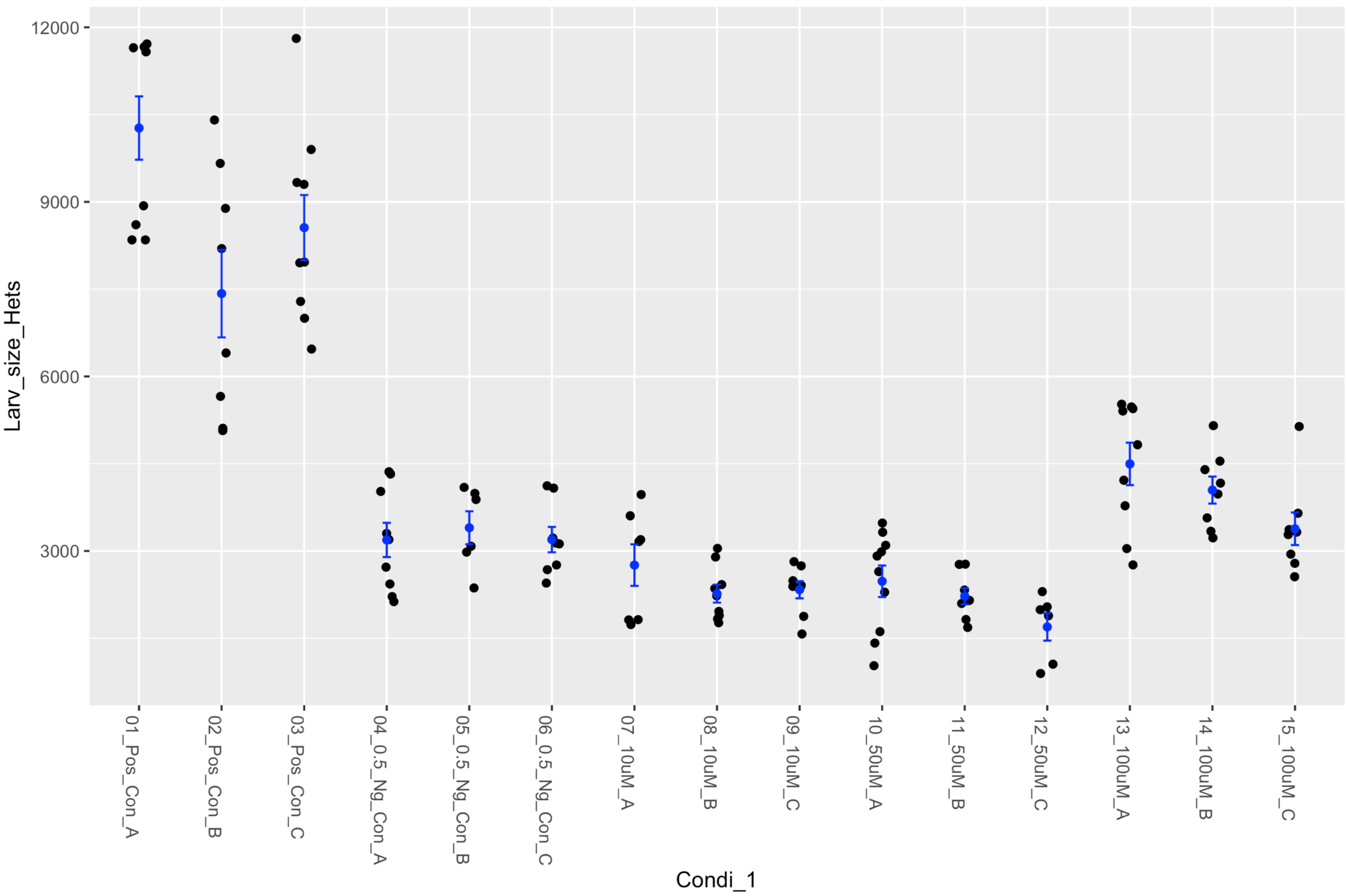


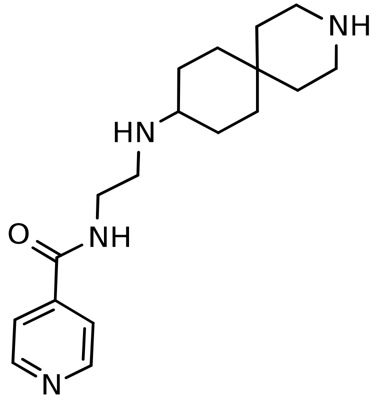

### Fly Chembridge retest – Supplier ID: 9206326

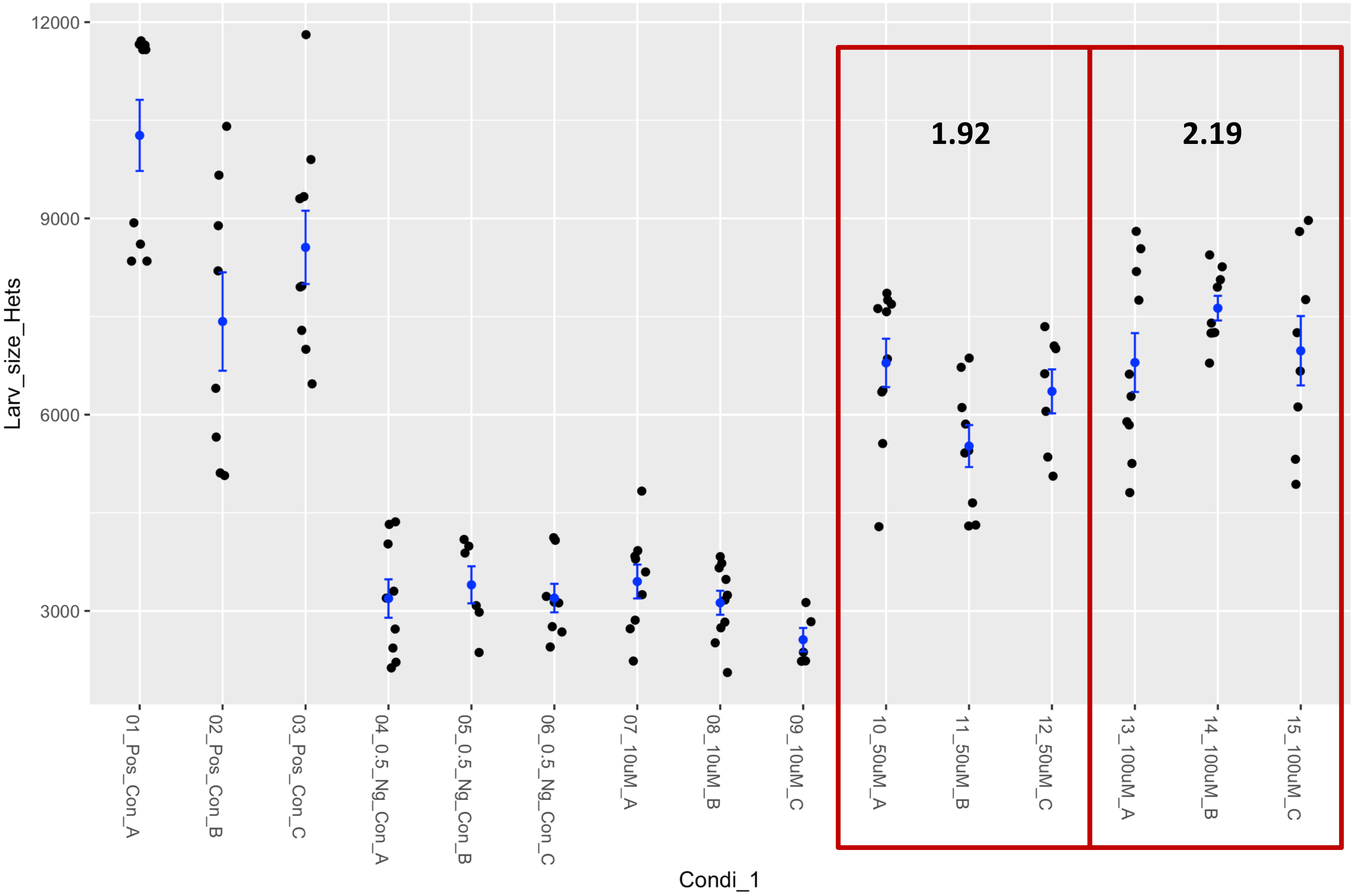


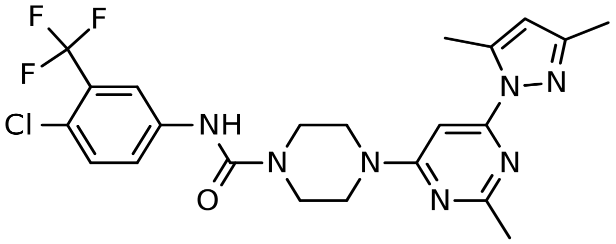

### Fly Chembridge retest – Supplier ID: 9233974

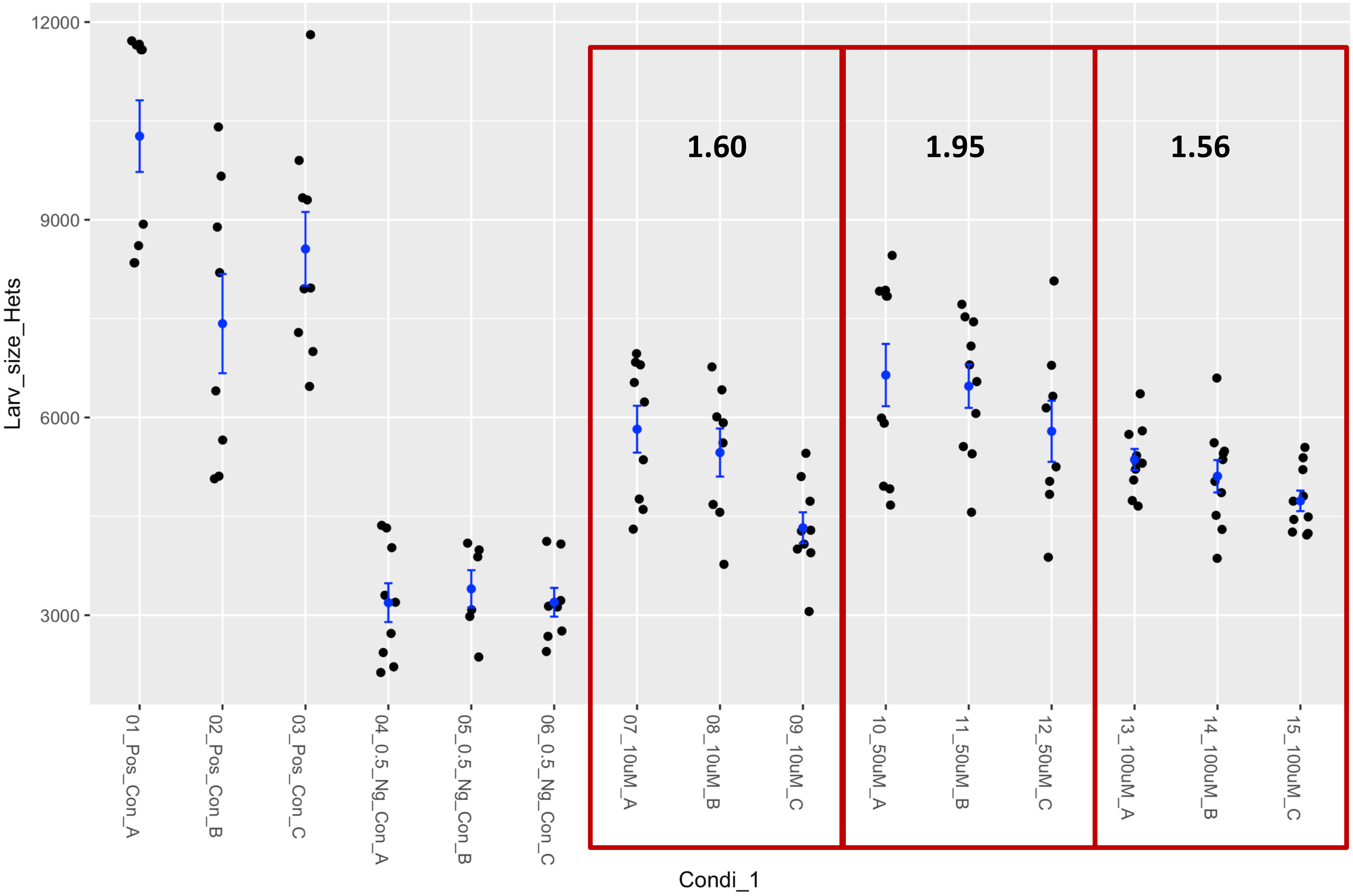


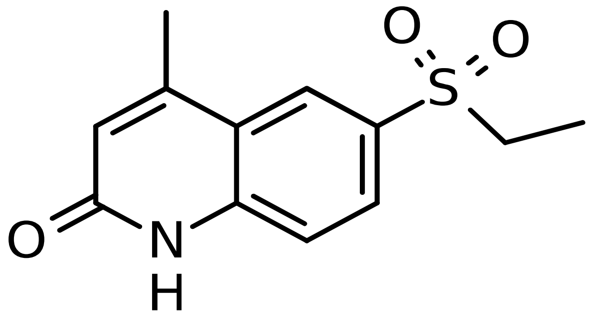

### Fly Chembridge retest – Supplier ID: 9251119

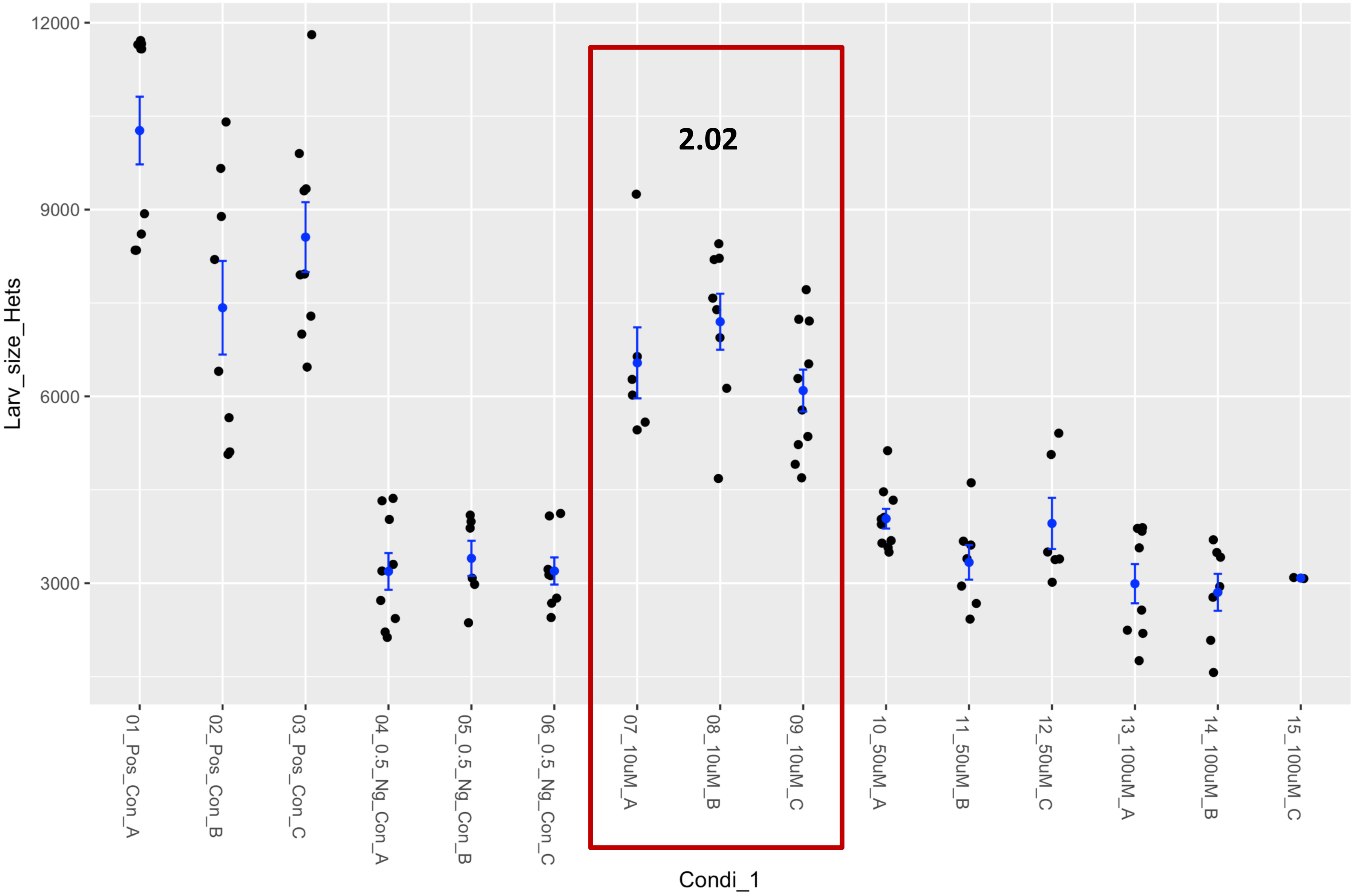


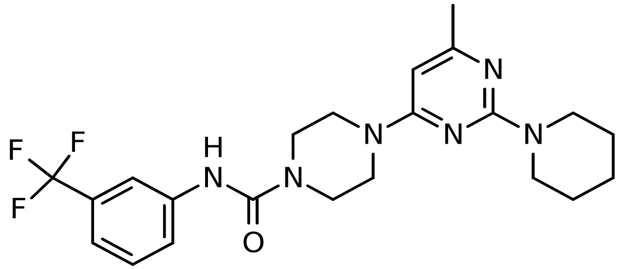

### Fly Chembridge retest – Supplier ID: 9251633

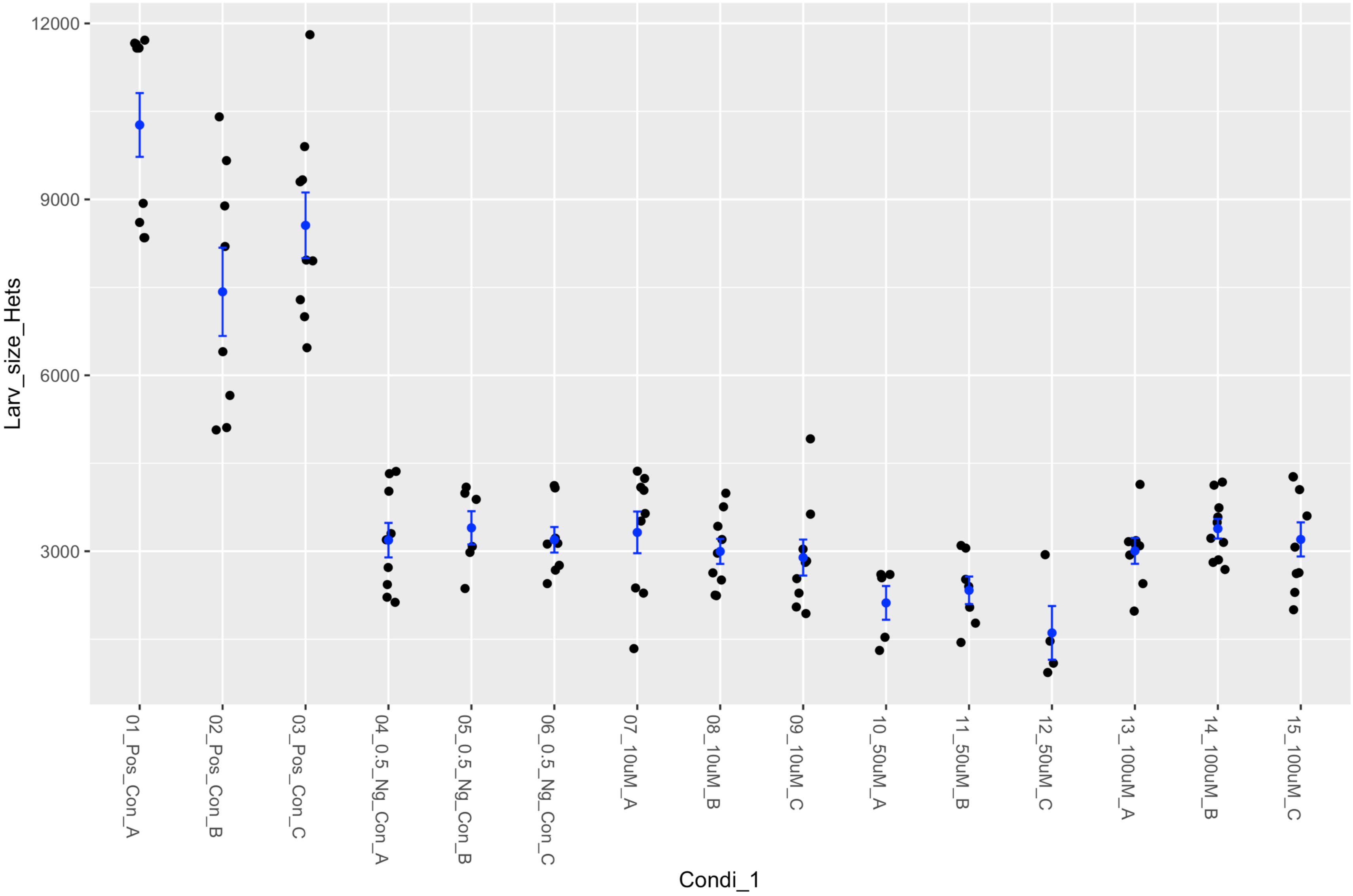


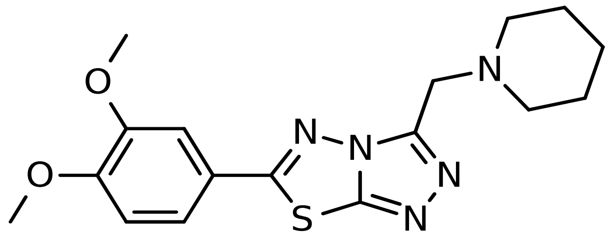

### Fly Chembridge retest – Supplier ID: 9262220

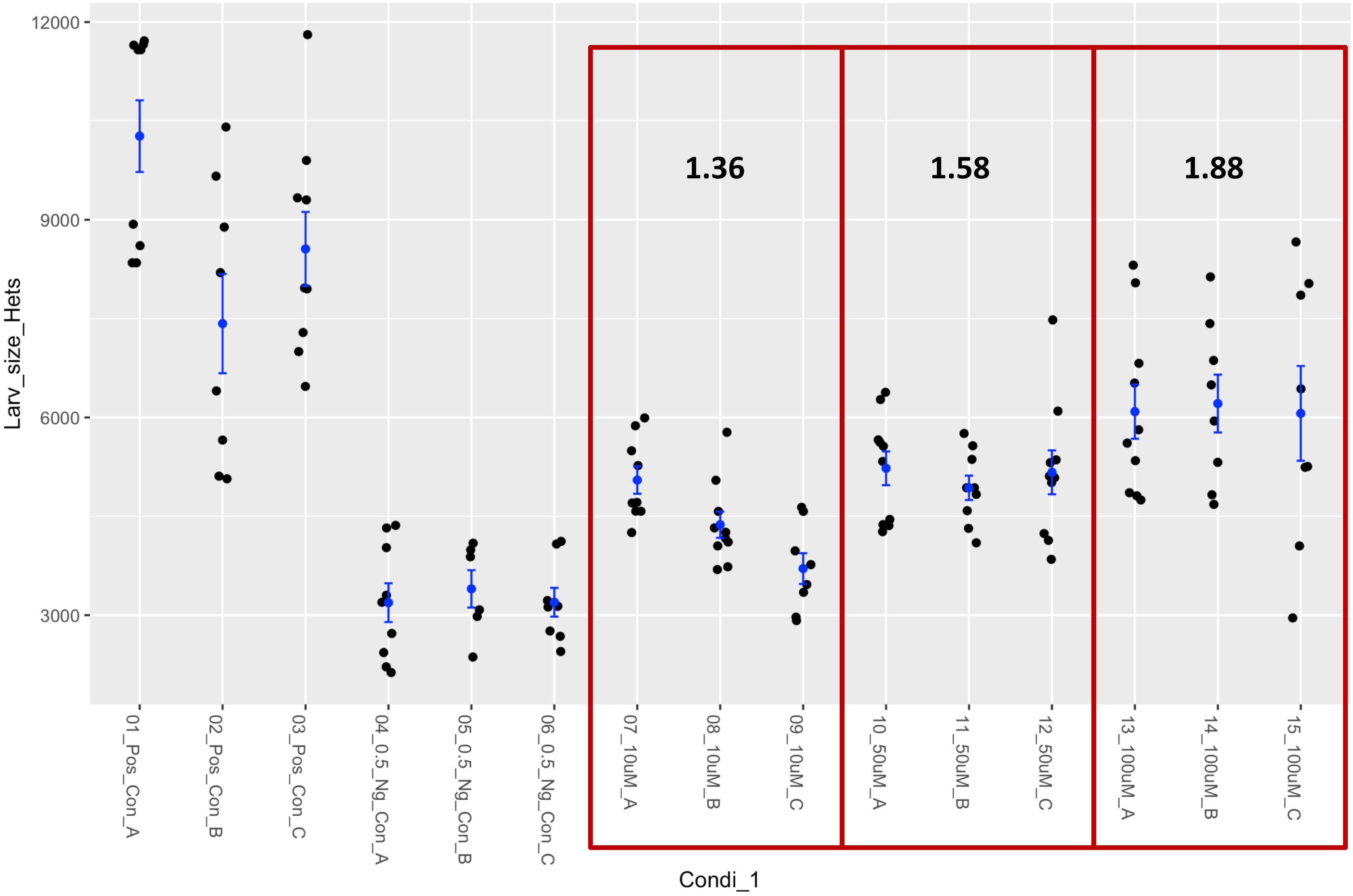


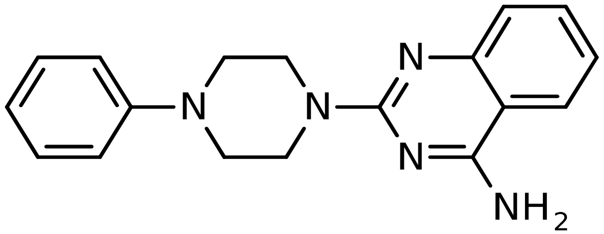

### Fly Chembridge retest – Supplier ID: 9262955

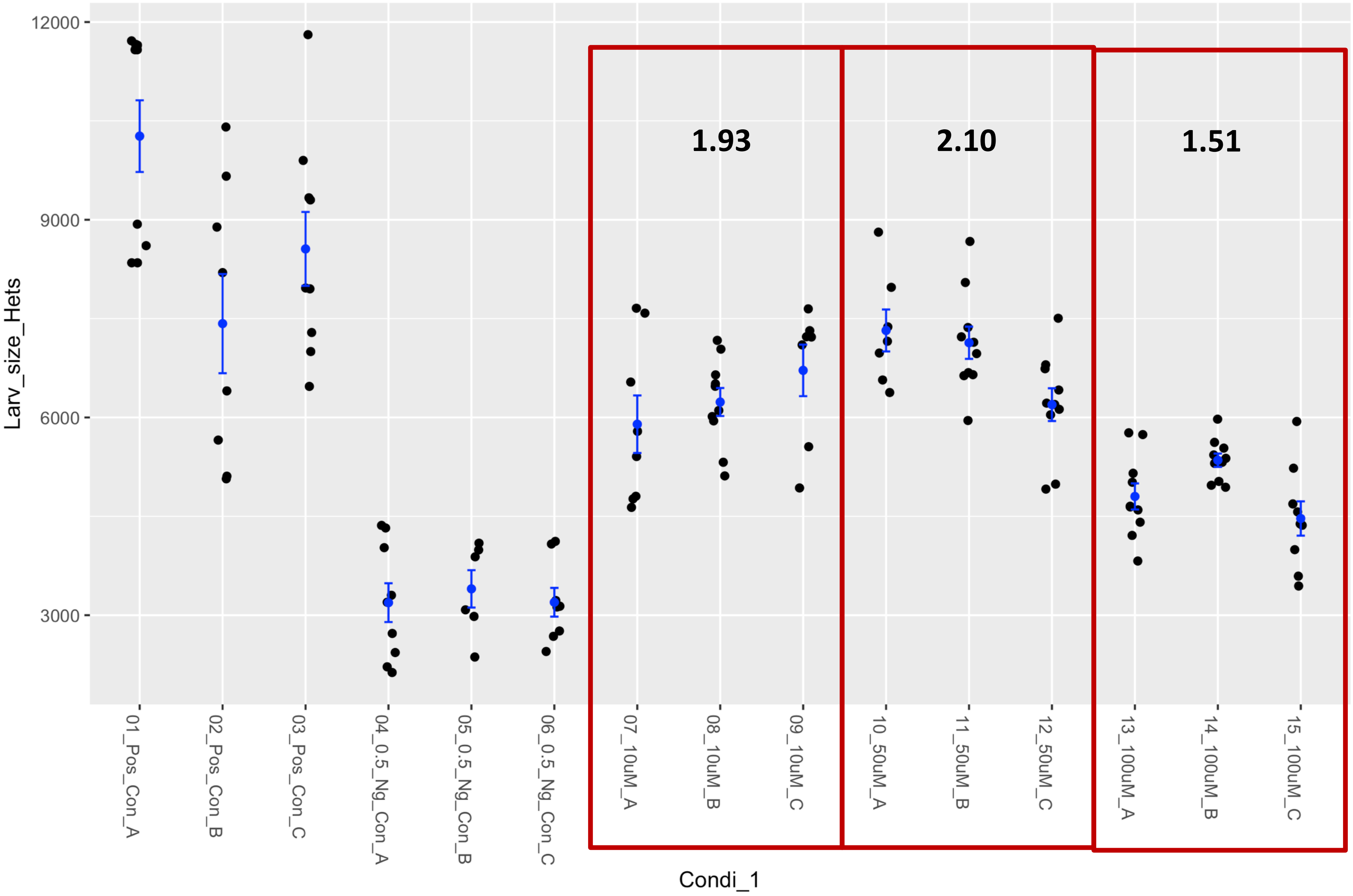


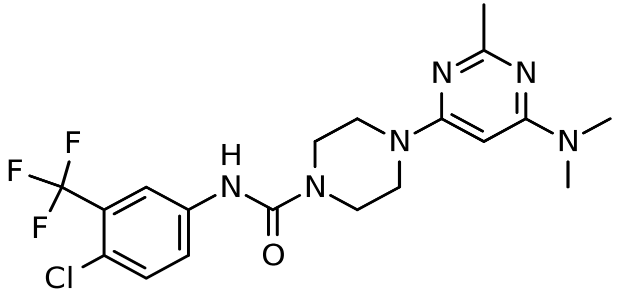

### Fly Chembridge retest – Supplier ID: 9290684

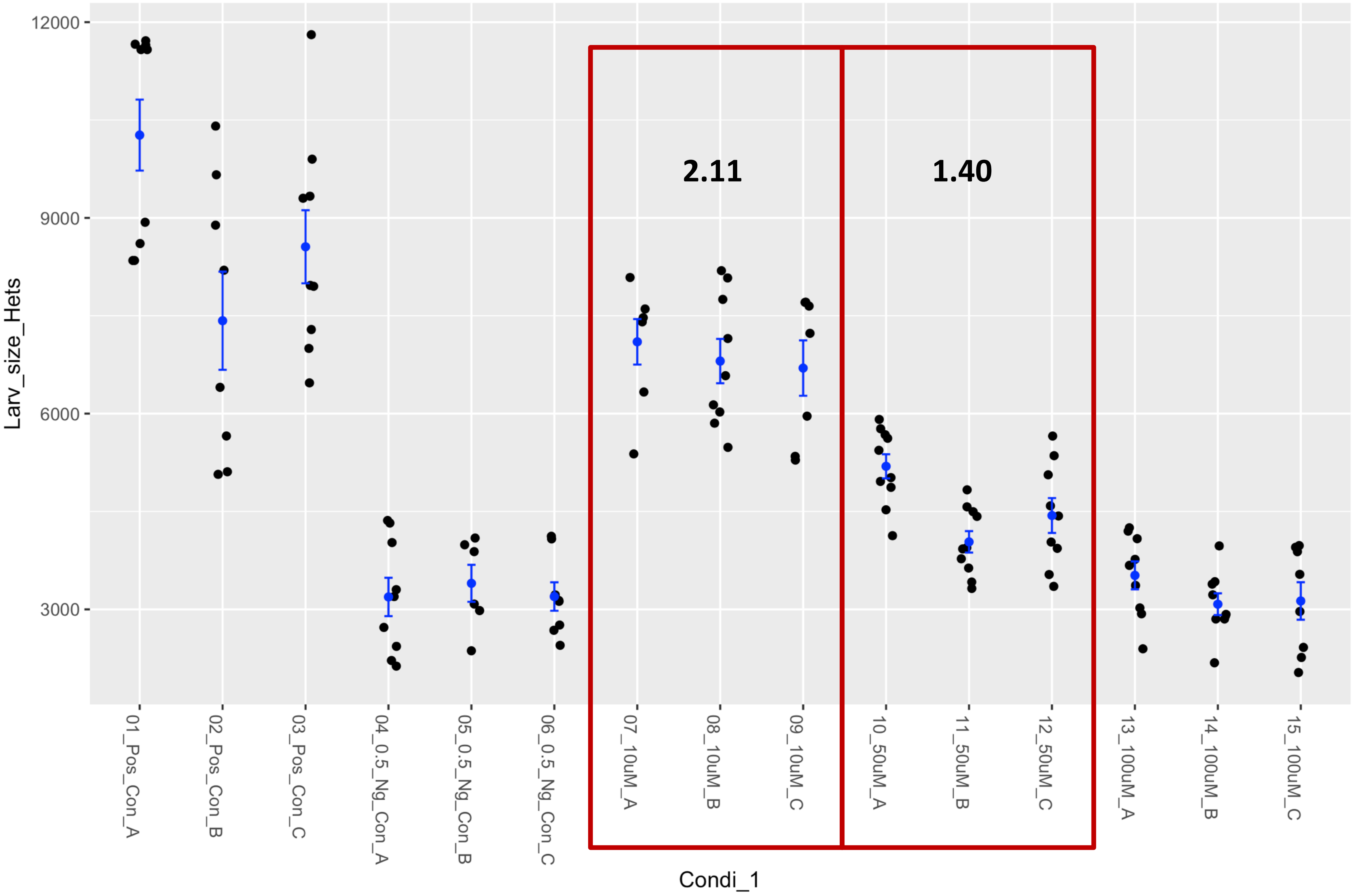


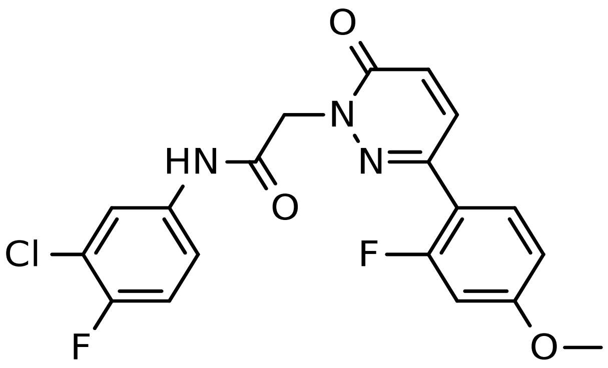

### Fly Worm MS cross validation

### Fly WormMS cross validation –
#### 3,4-DIDESMETHYL-5-DESHYDROXY-3'-ETHOXYSCLEROIN

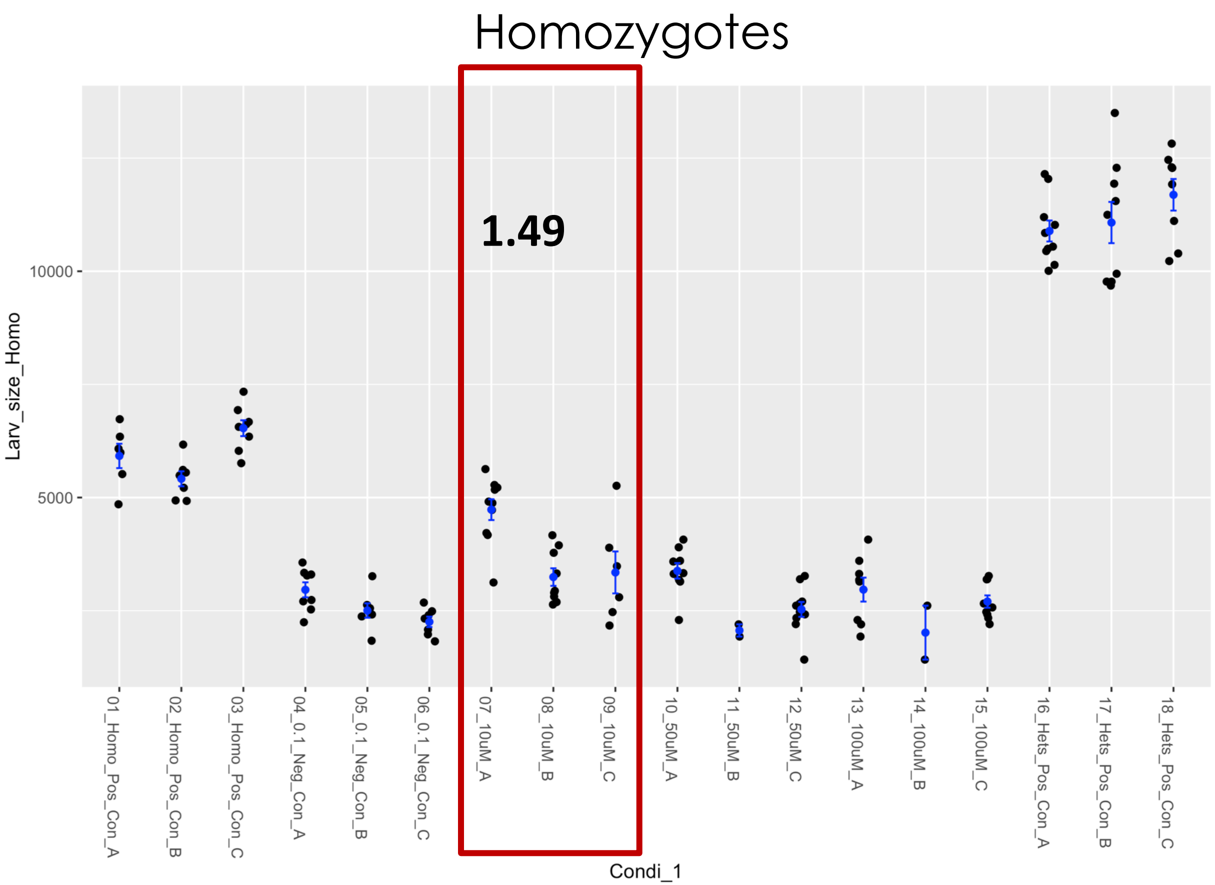


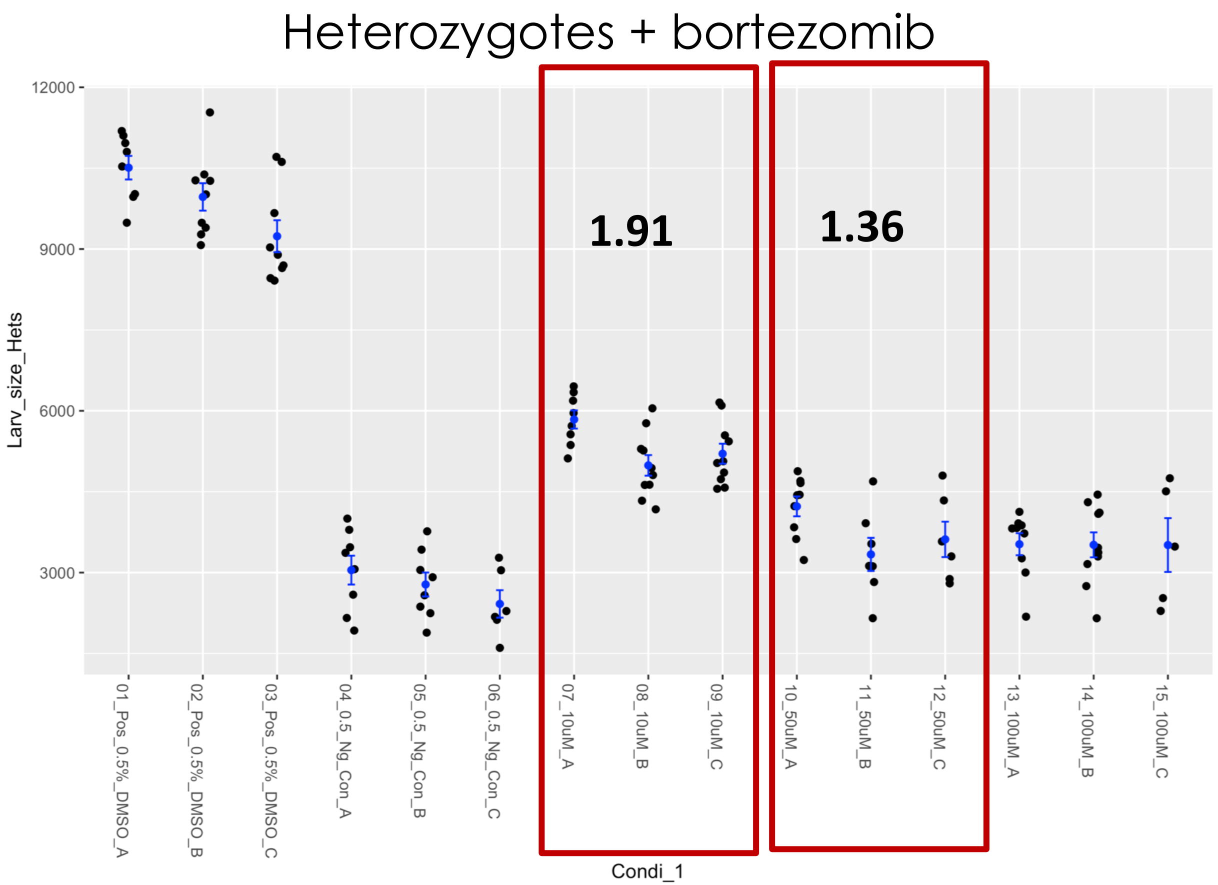

### Fly WormMS cross validation – BENSERAZIDE

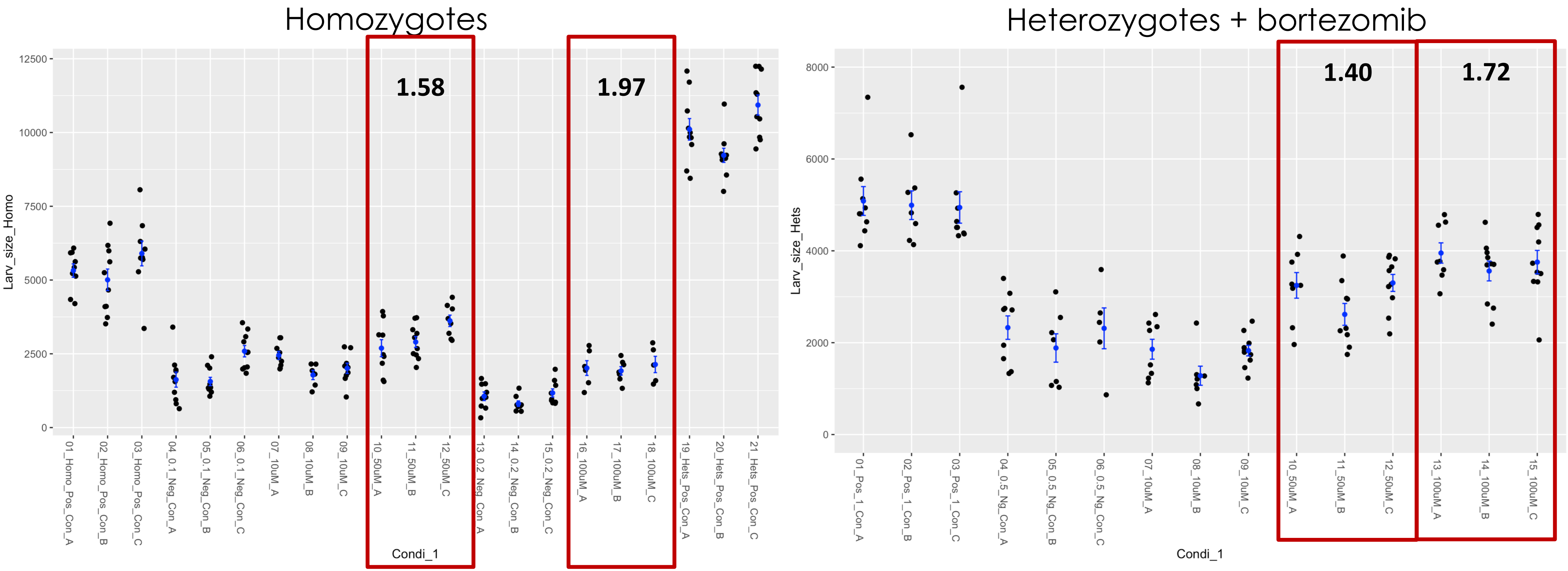

### Fly WormMS cross validation – ELLAGIC ACID

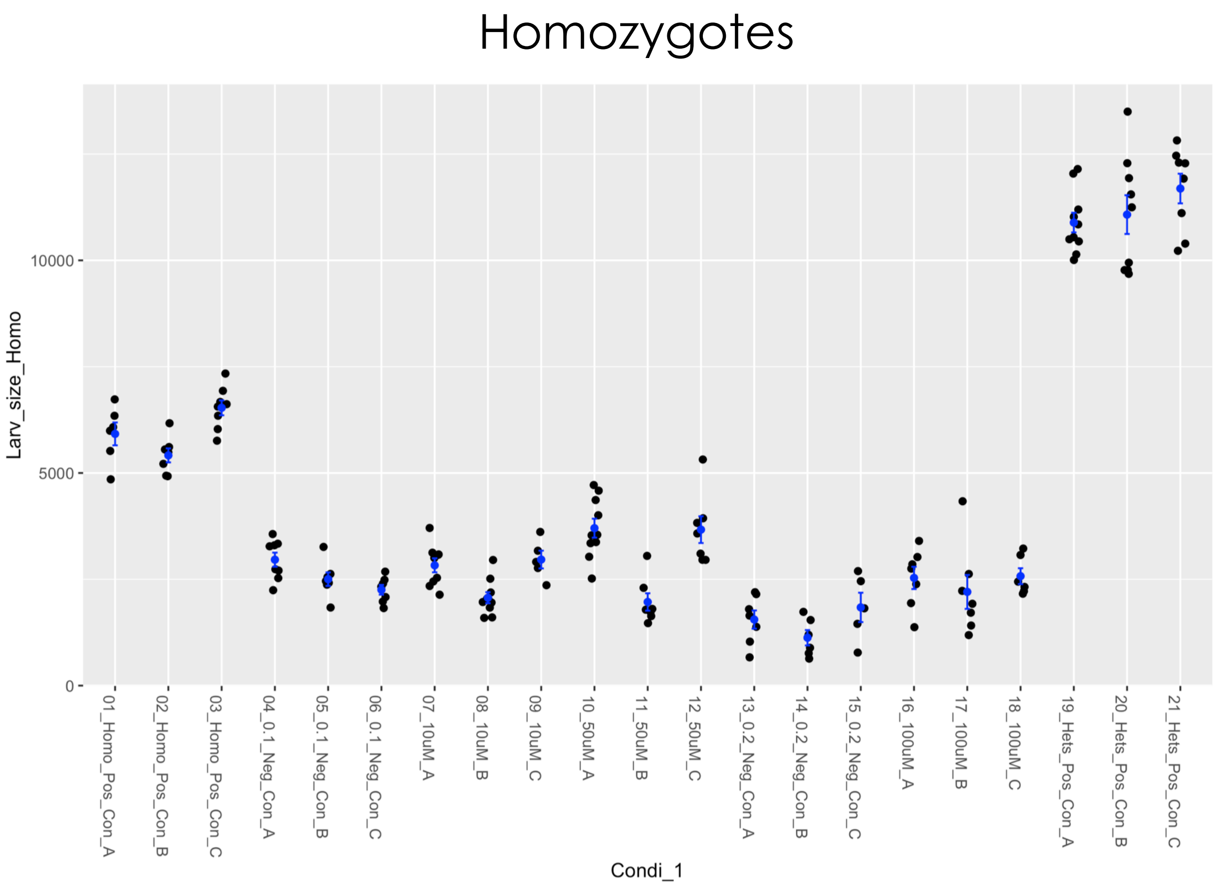


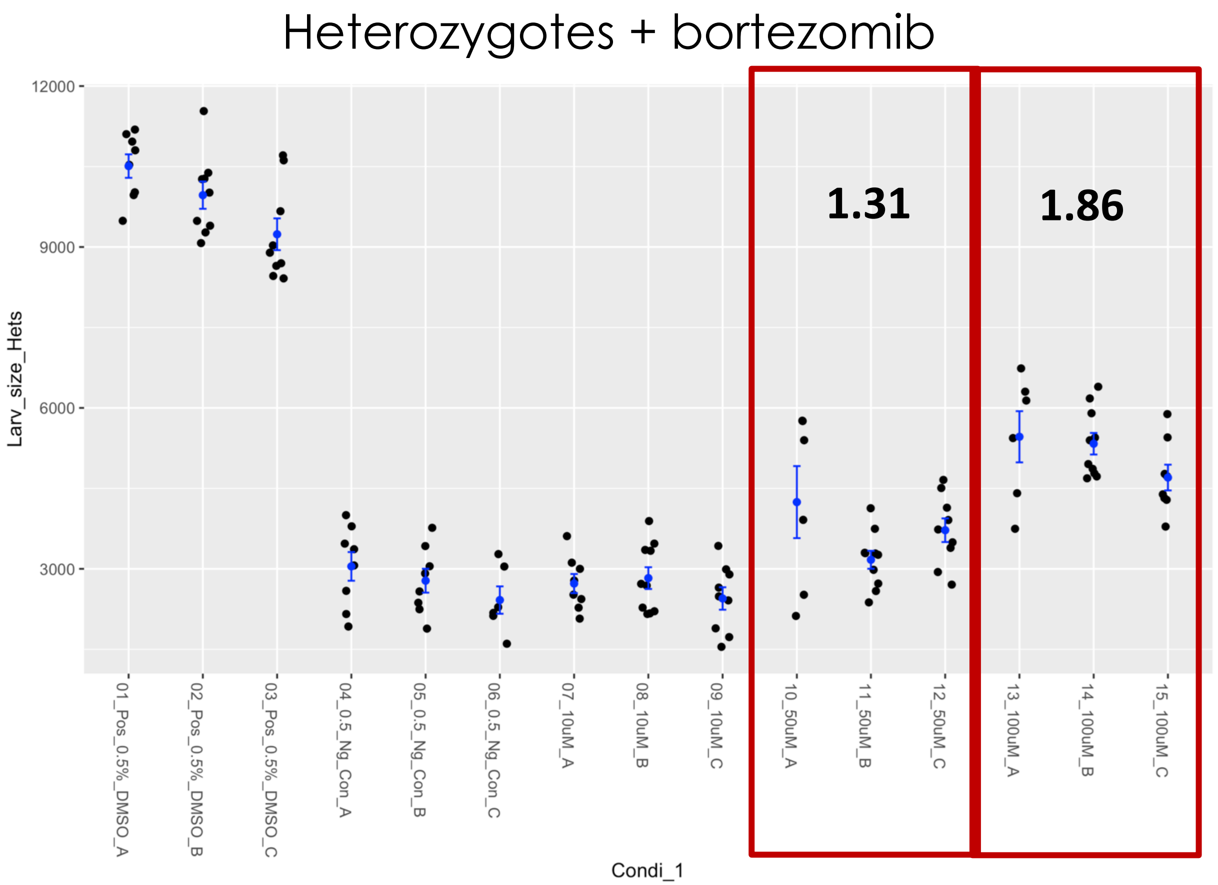

### Fly WormMS cross validation – EPICATECHIN MONOGALLATE

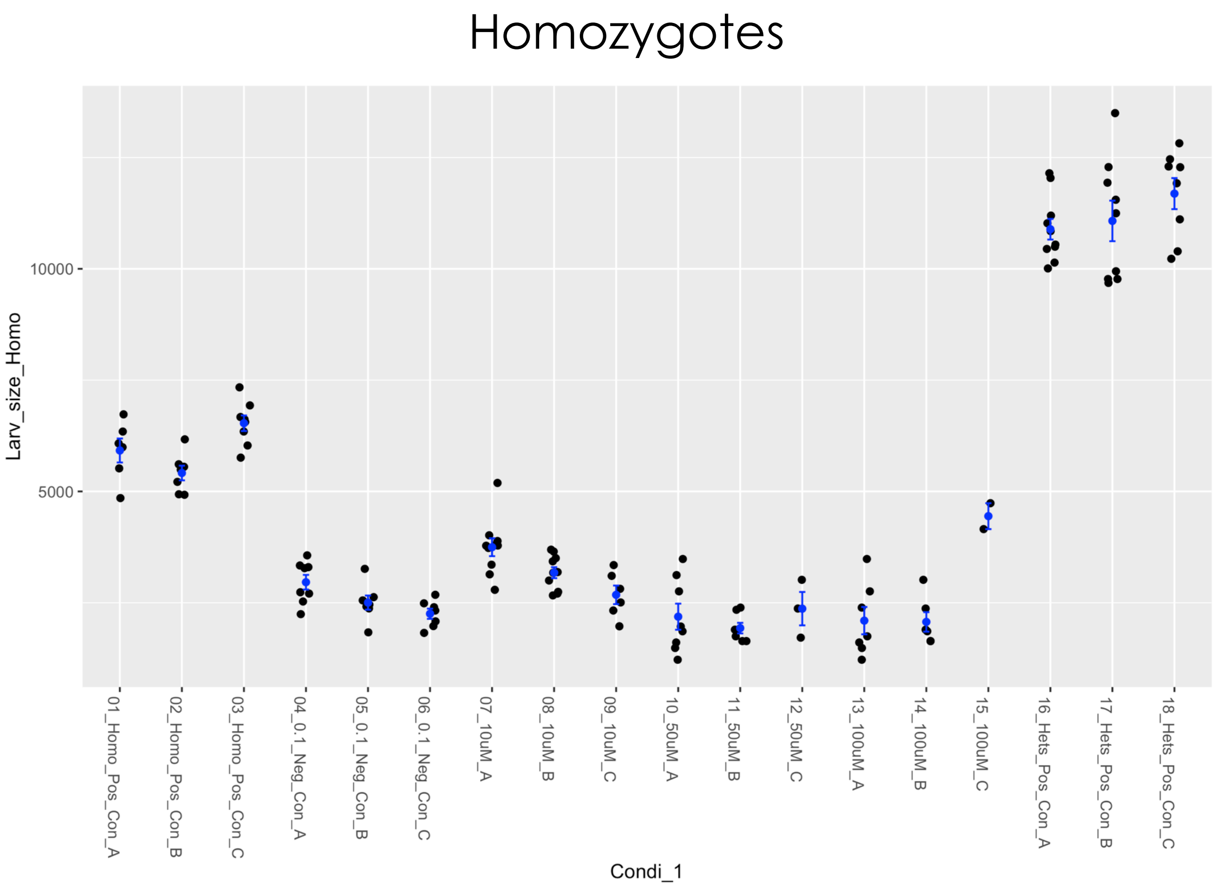


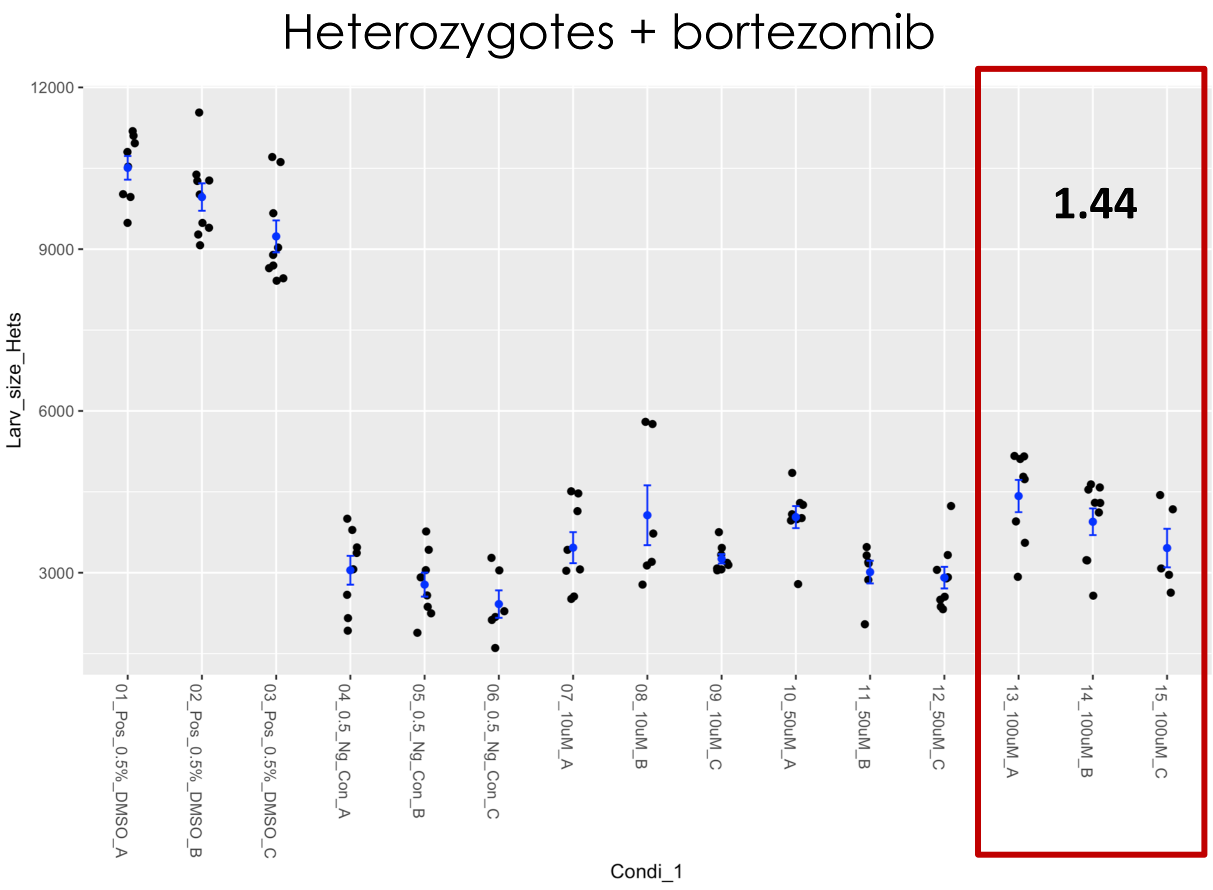

### Fly WormMS cross validation – KOPARIN

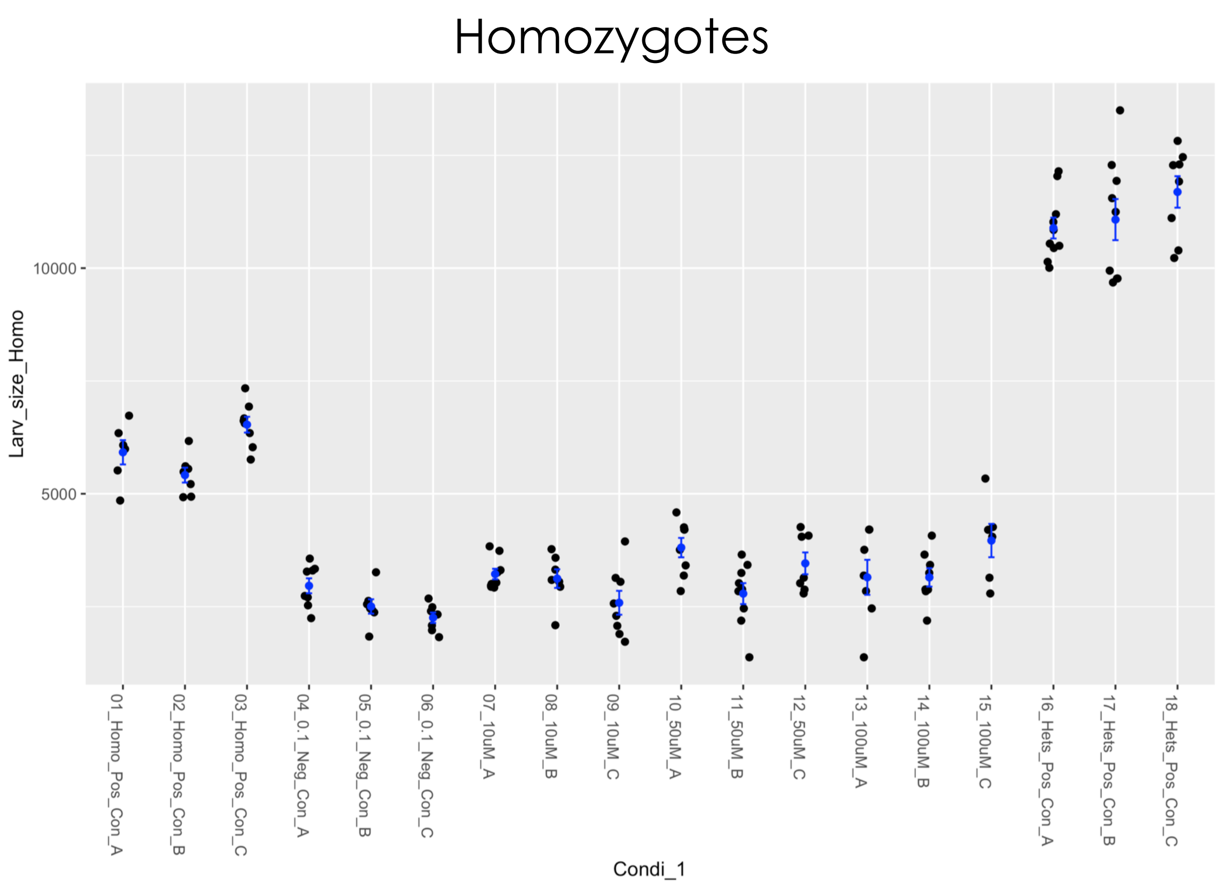


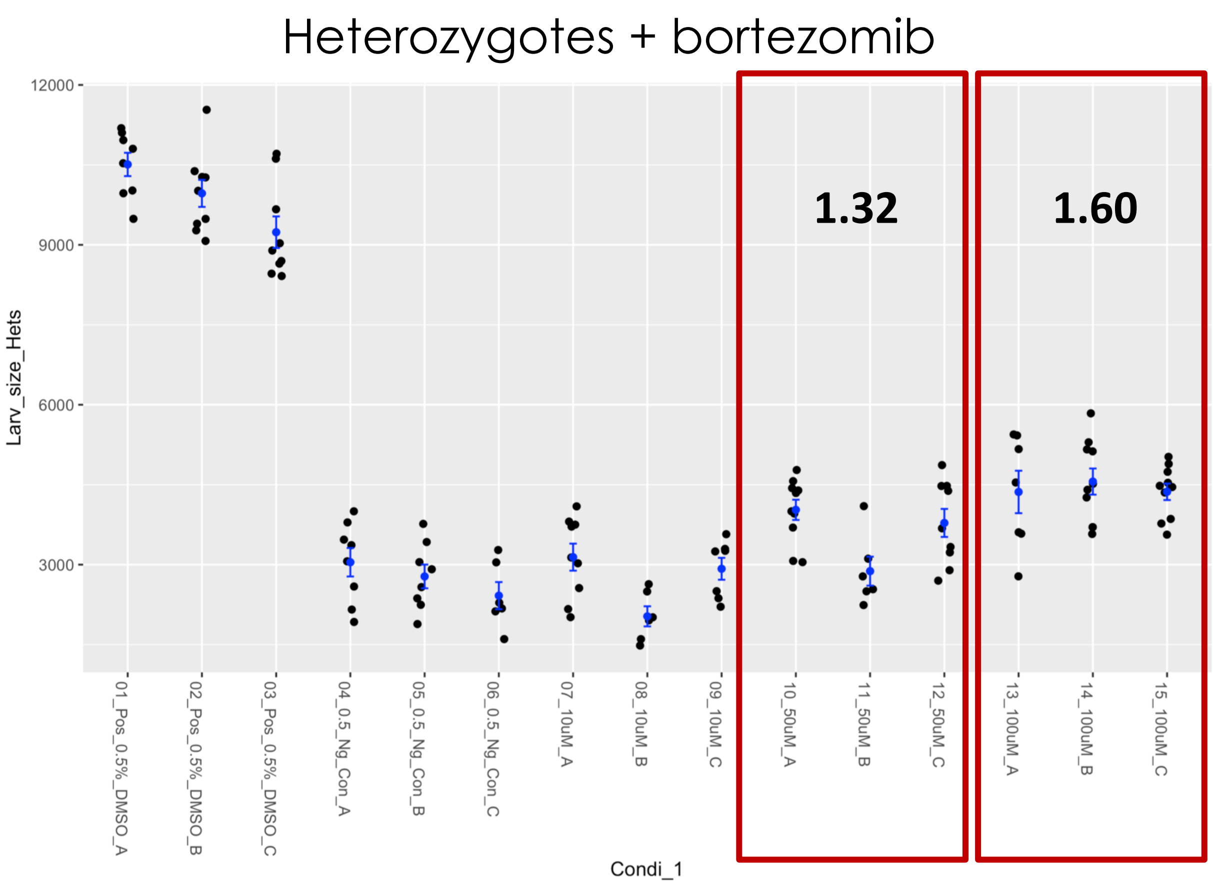

### Fly WormMS cross validation – PHENYLBUTAZONE

Homozygotes

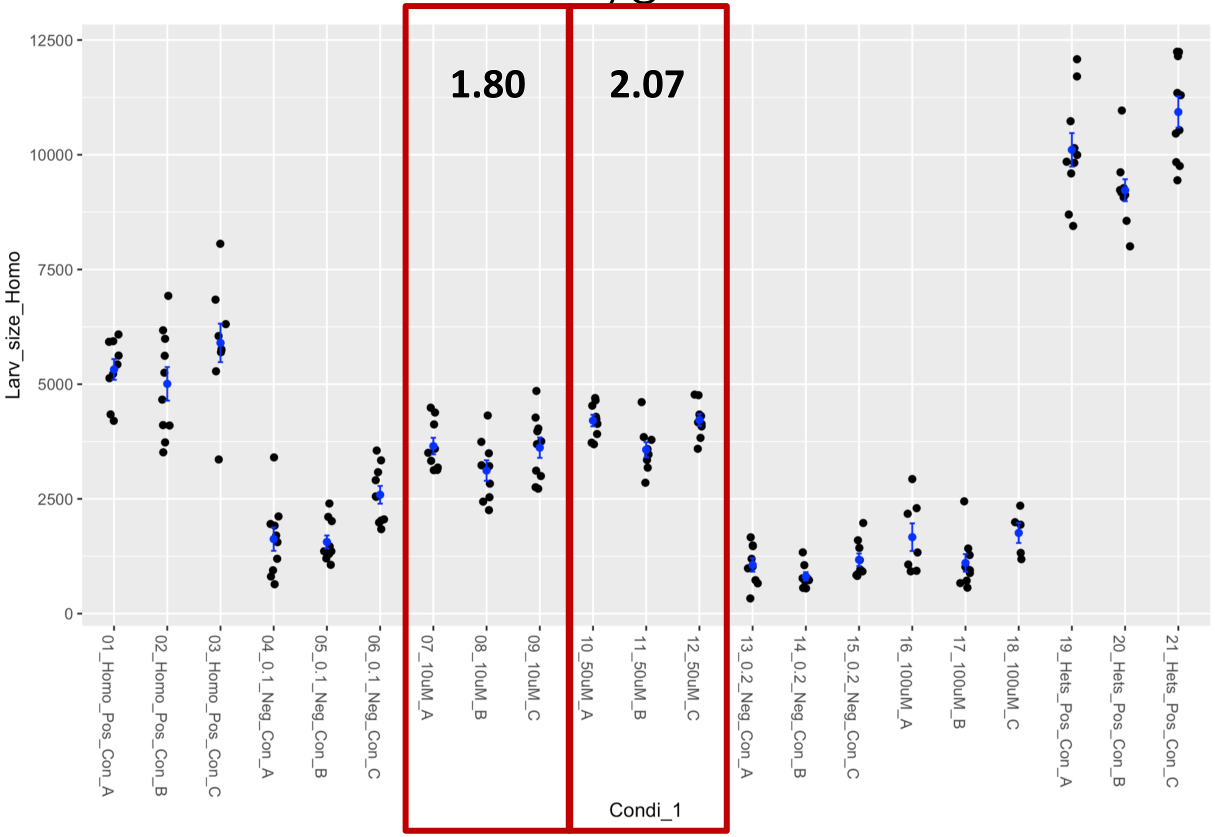


Heterozygotes + bortezomib

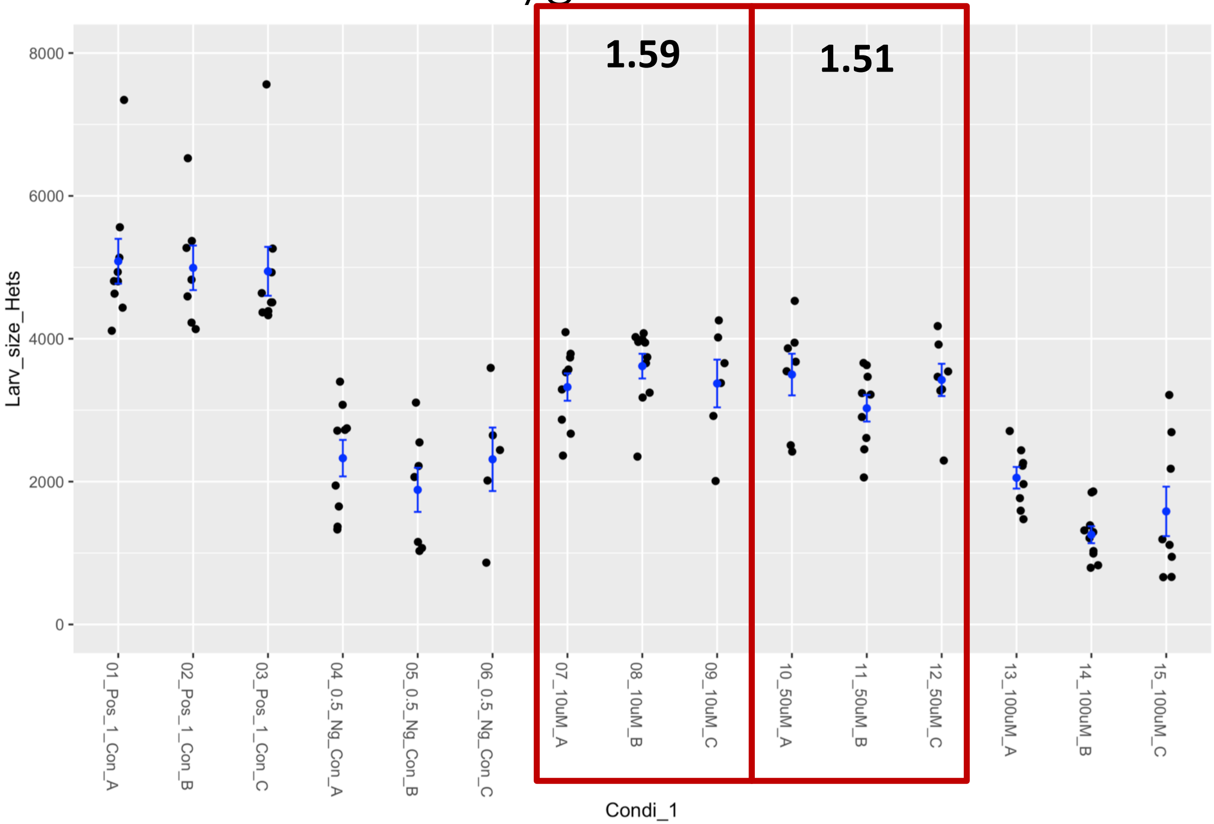

### Fly WormMS cross validation – POMIFERIN

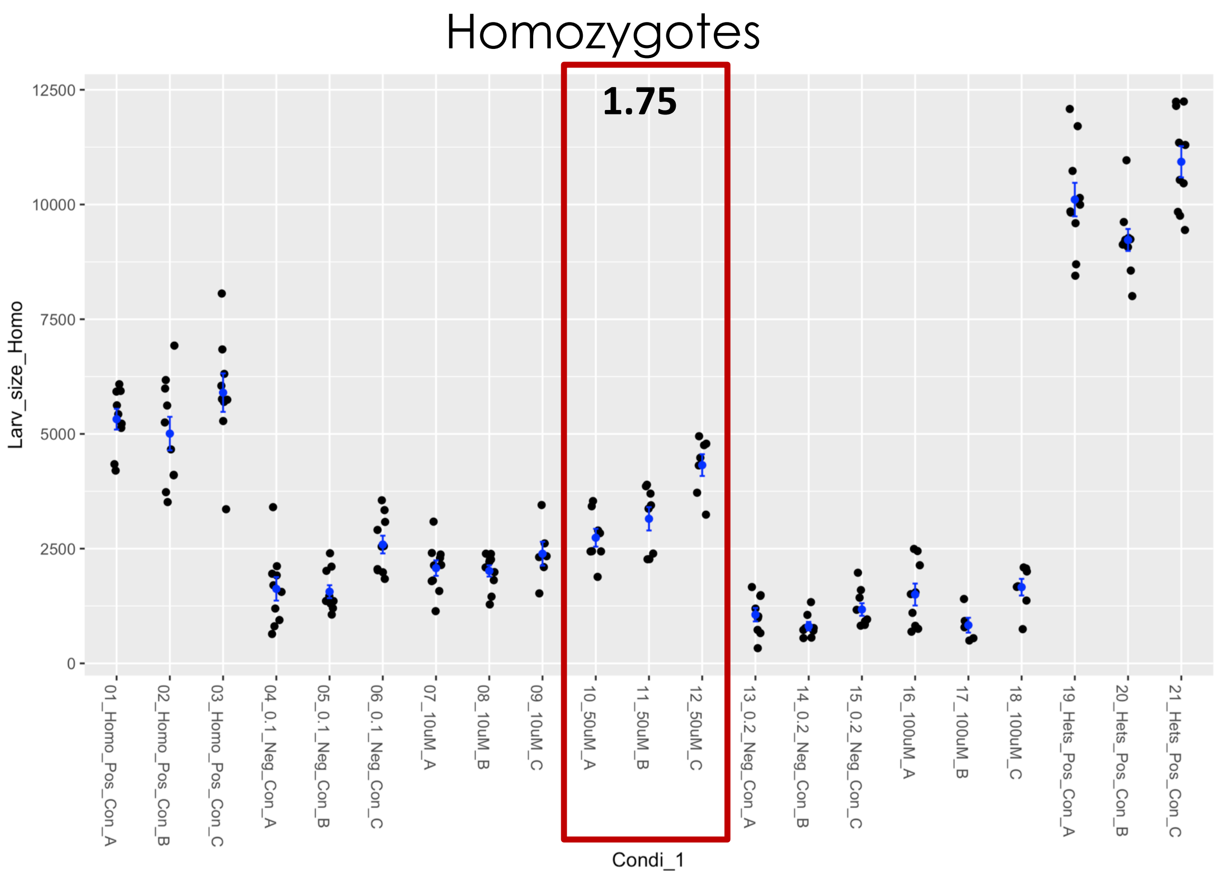


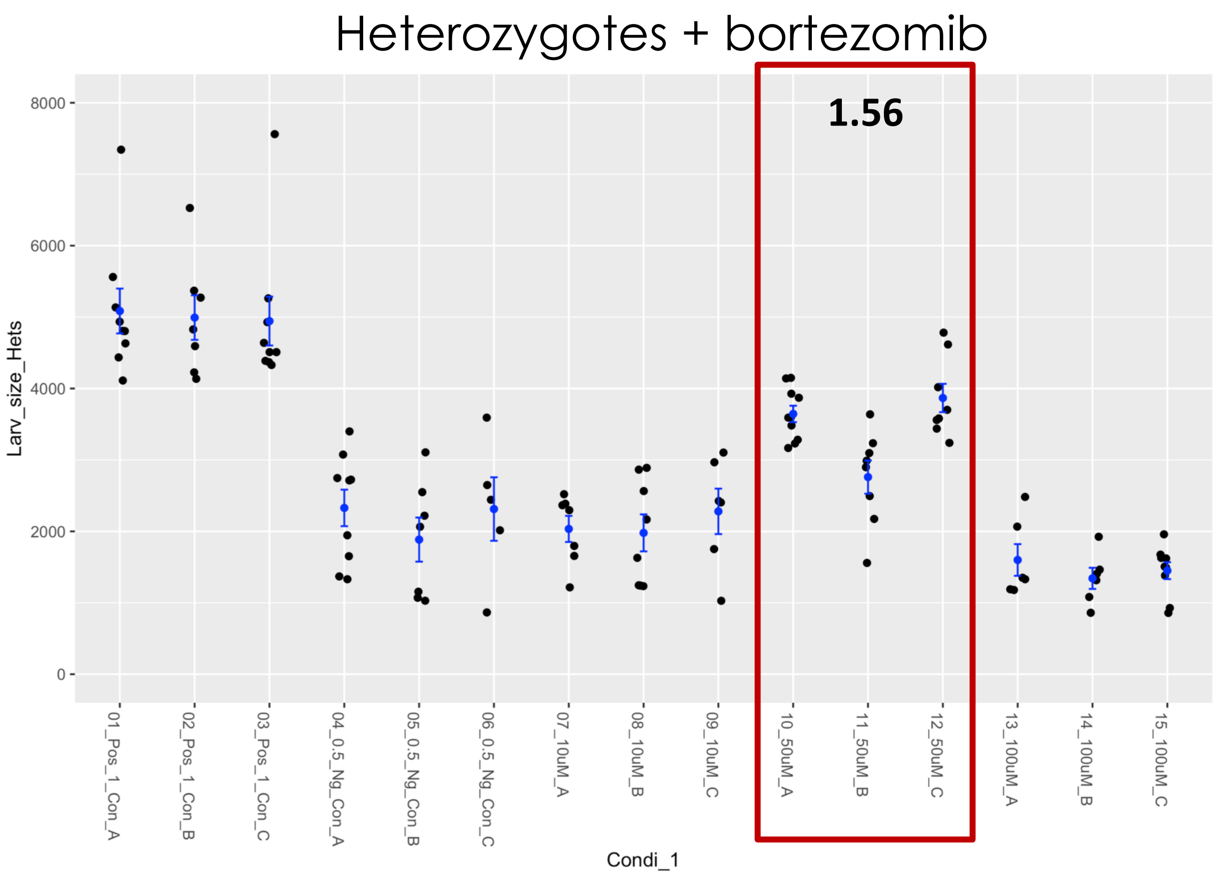

### Fly WormMS cross validation – PURPUROGALLIN-4-CARBOXYLIC ACID

Homozygotes

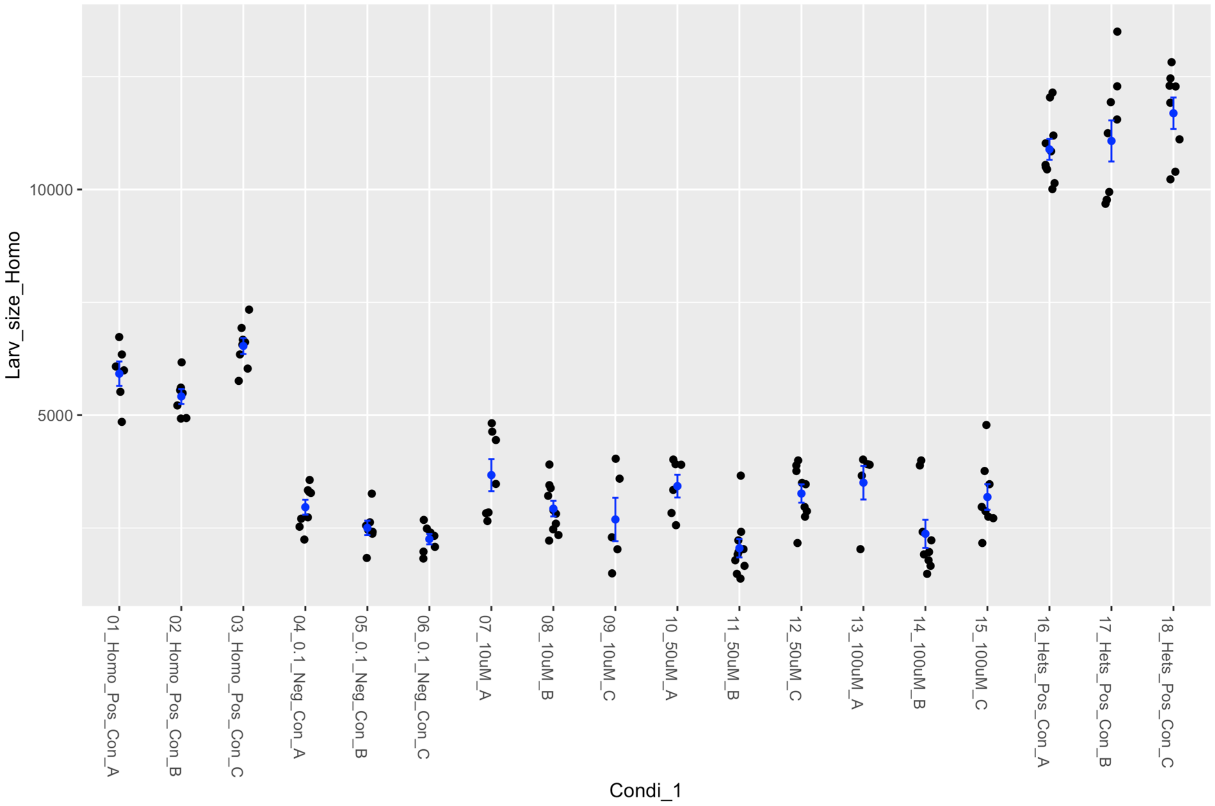


Heterozygotes + bortezomib

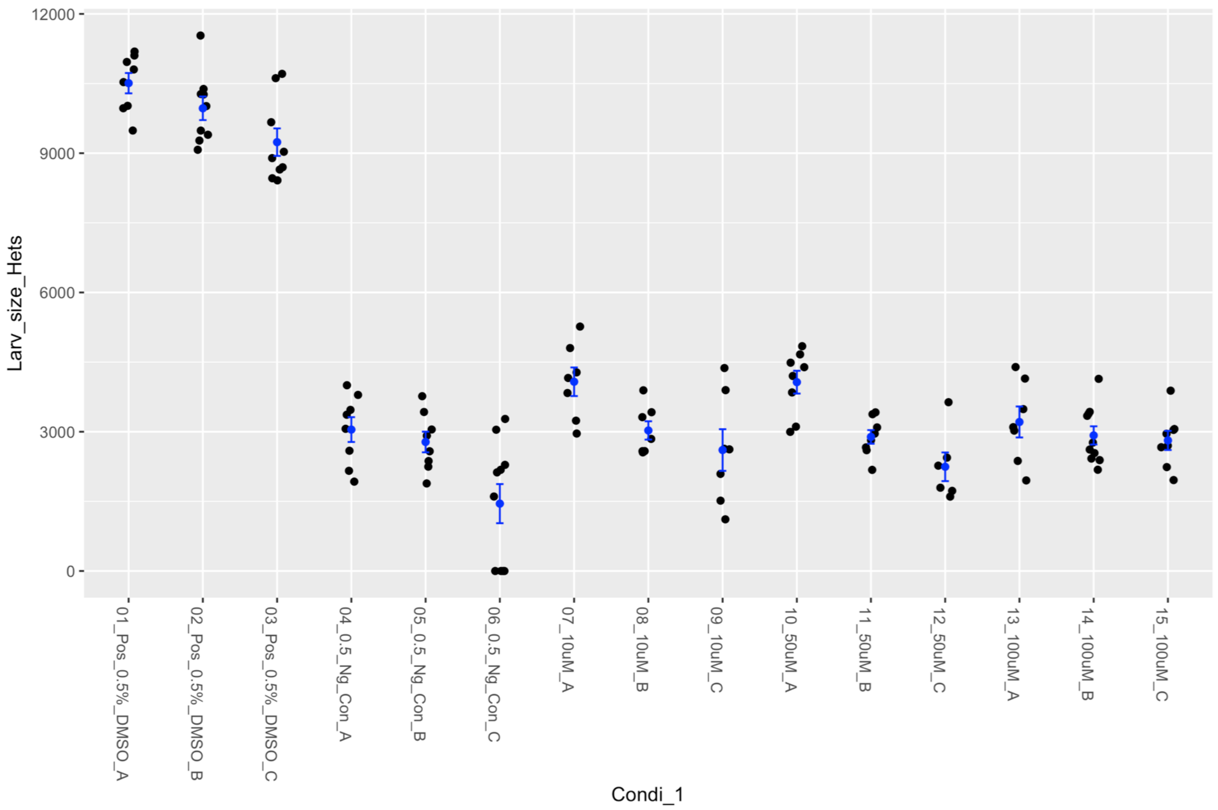

### Fly WormMS cross validation – QUERCETIN

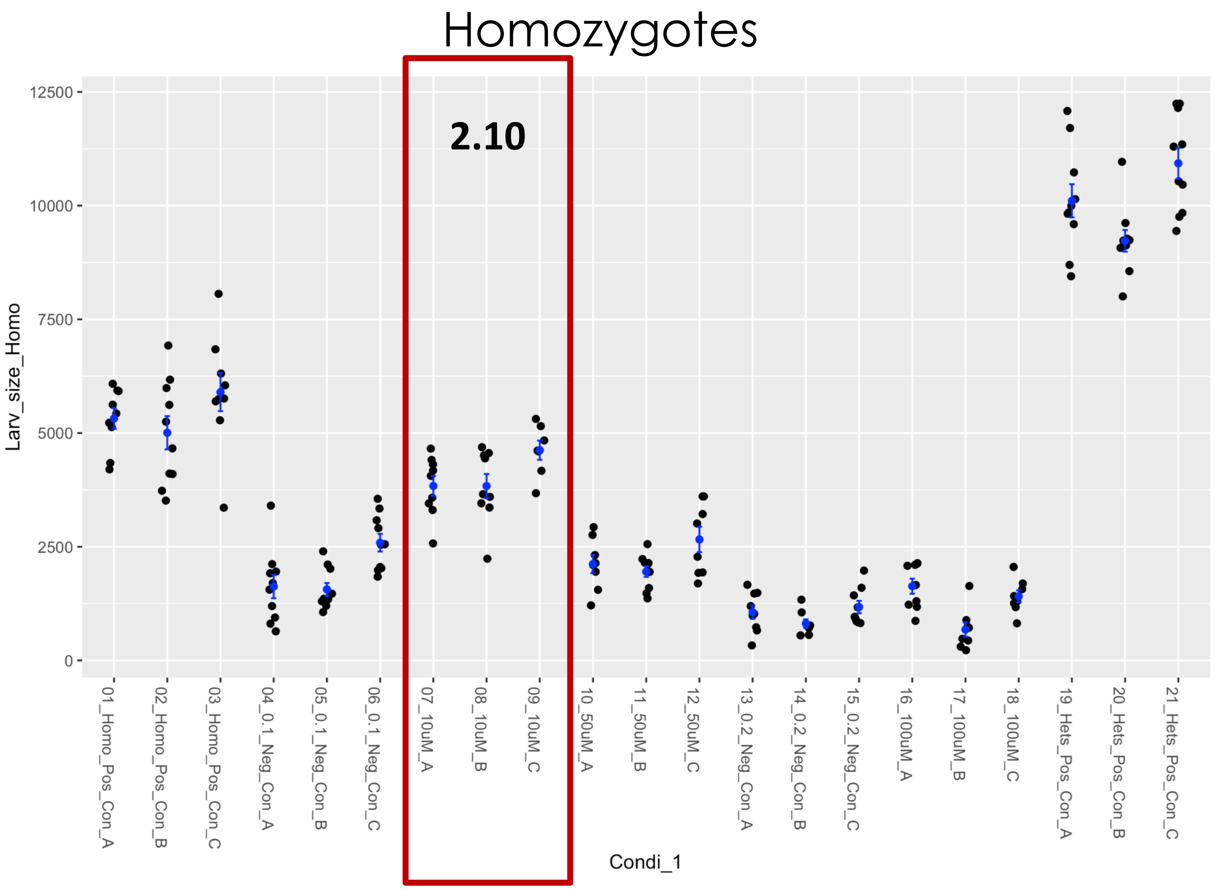


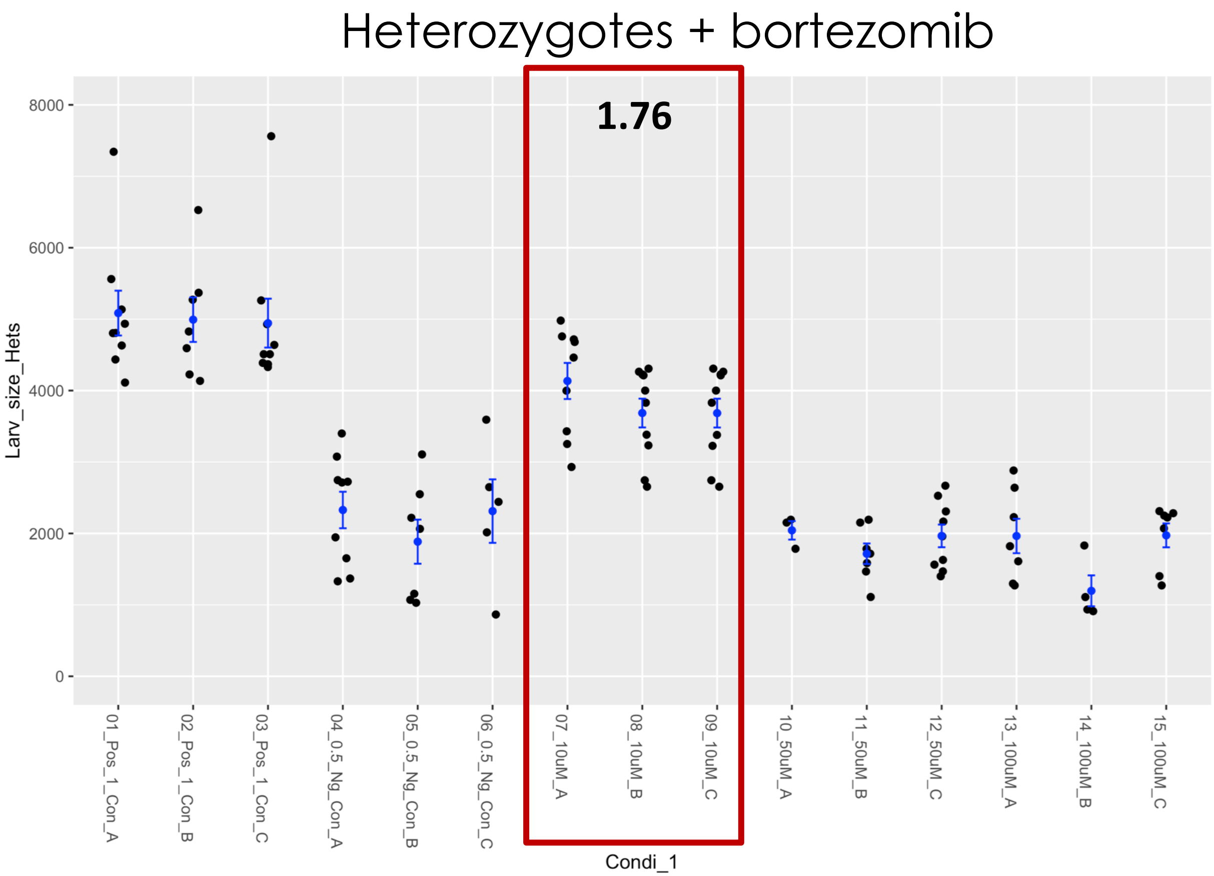

### Fly WormMS cross validation – THEAFLAVIN MONOGALLATE

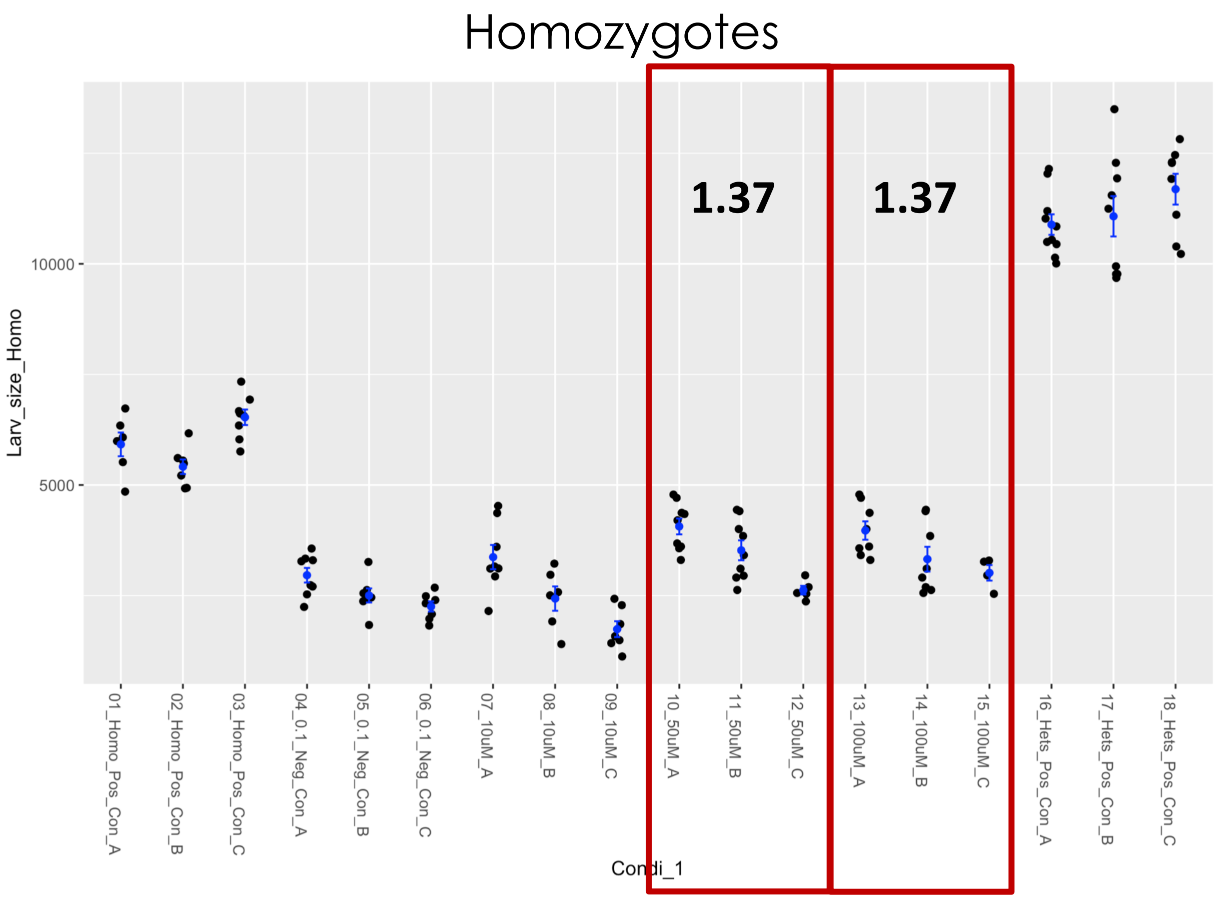


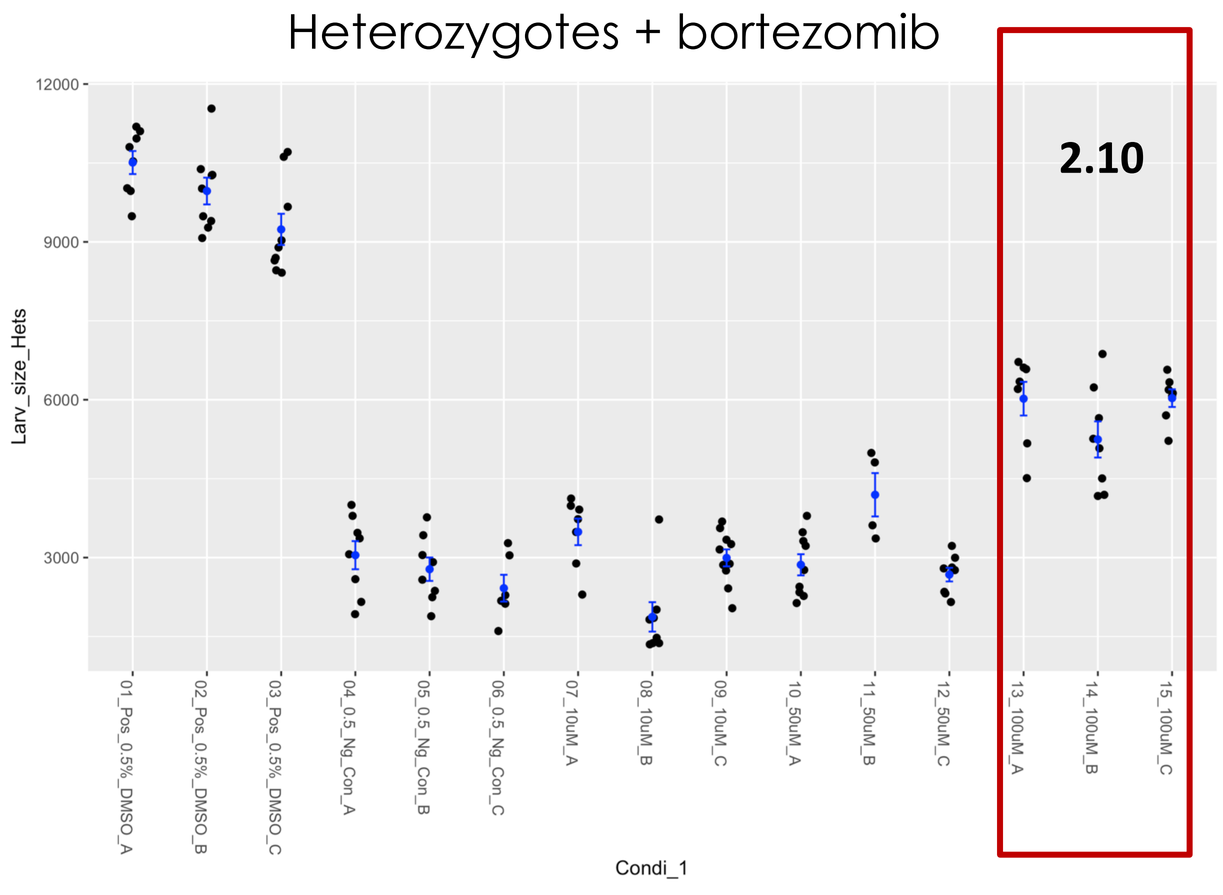

### Fly WormMS cross validation – TRIAMCINOLONE

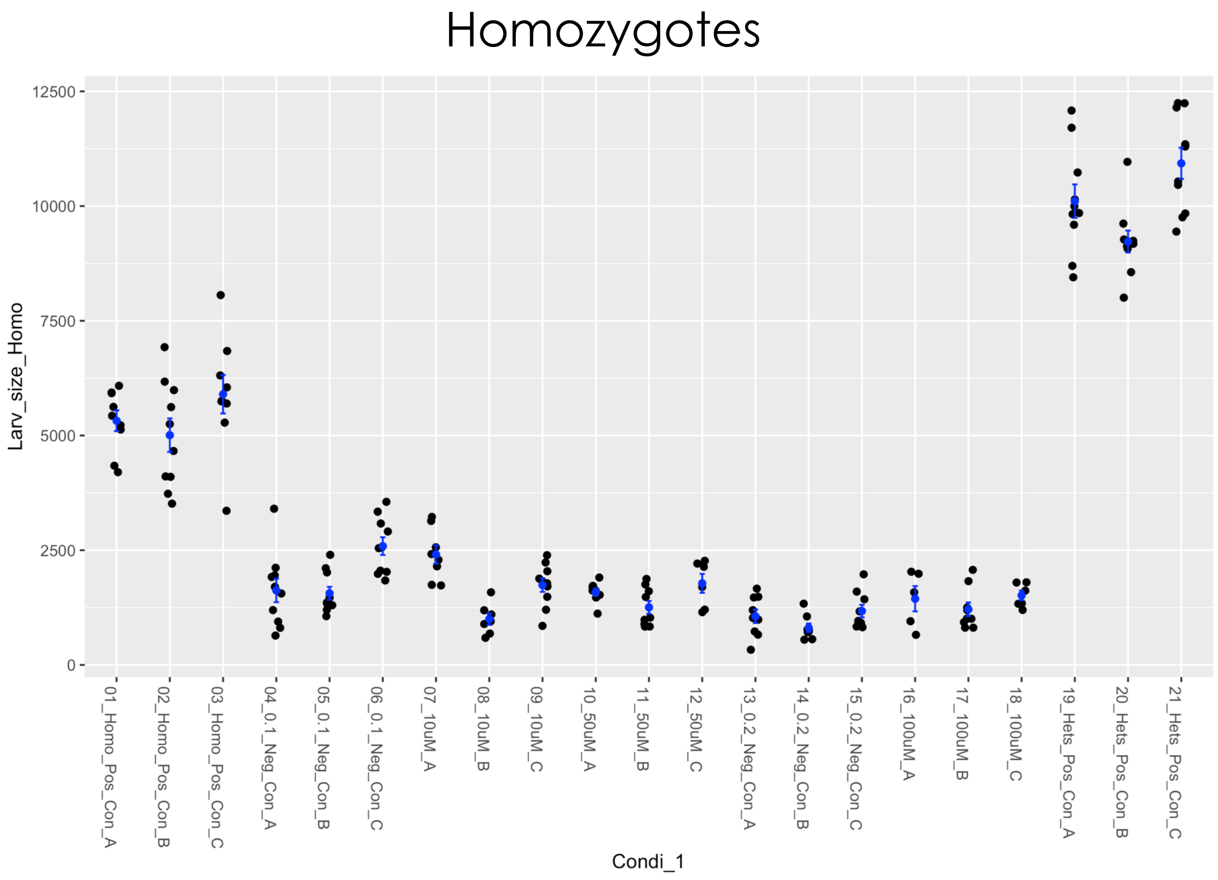


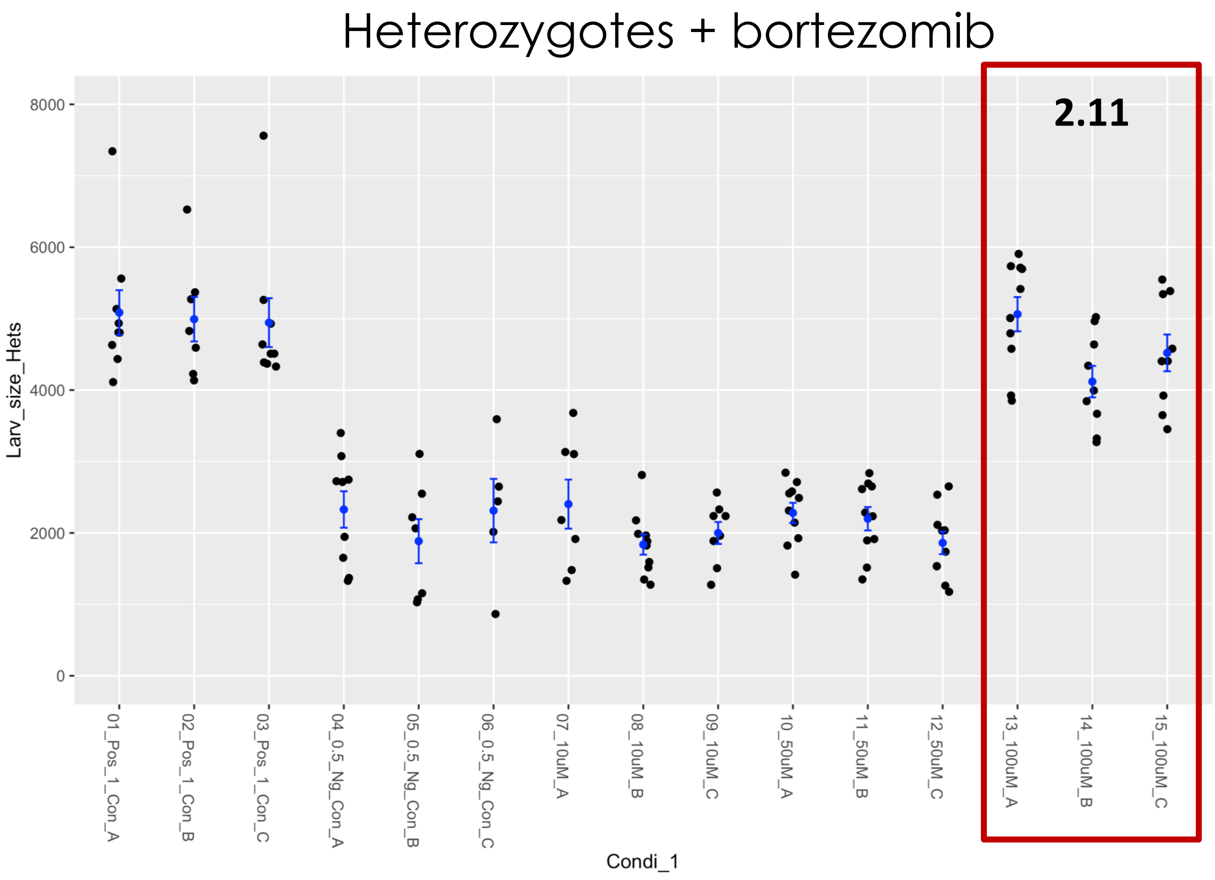

### Fly Microsource retest –2-METHYLENE-5-(2,5-DIOXOTETRAHYDROFURAN-3-YL)-6-OXO--10,10-DIMETHYL

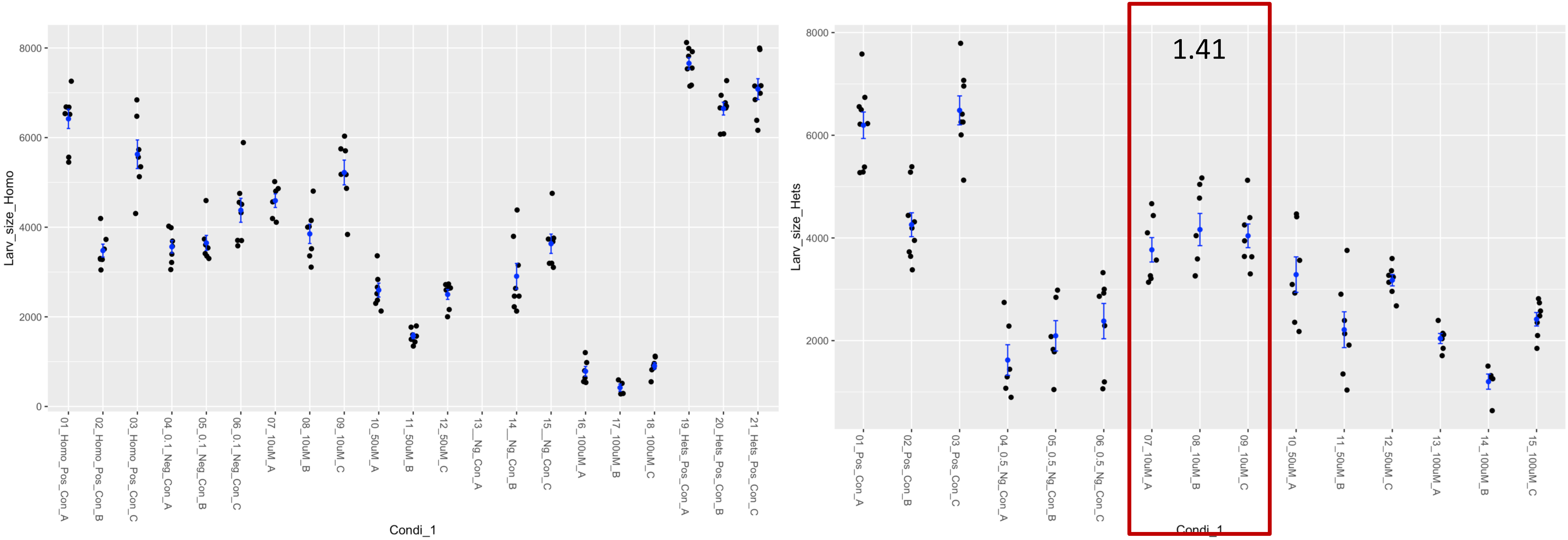

### Fly Microsource Retest – AMINACRINE

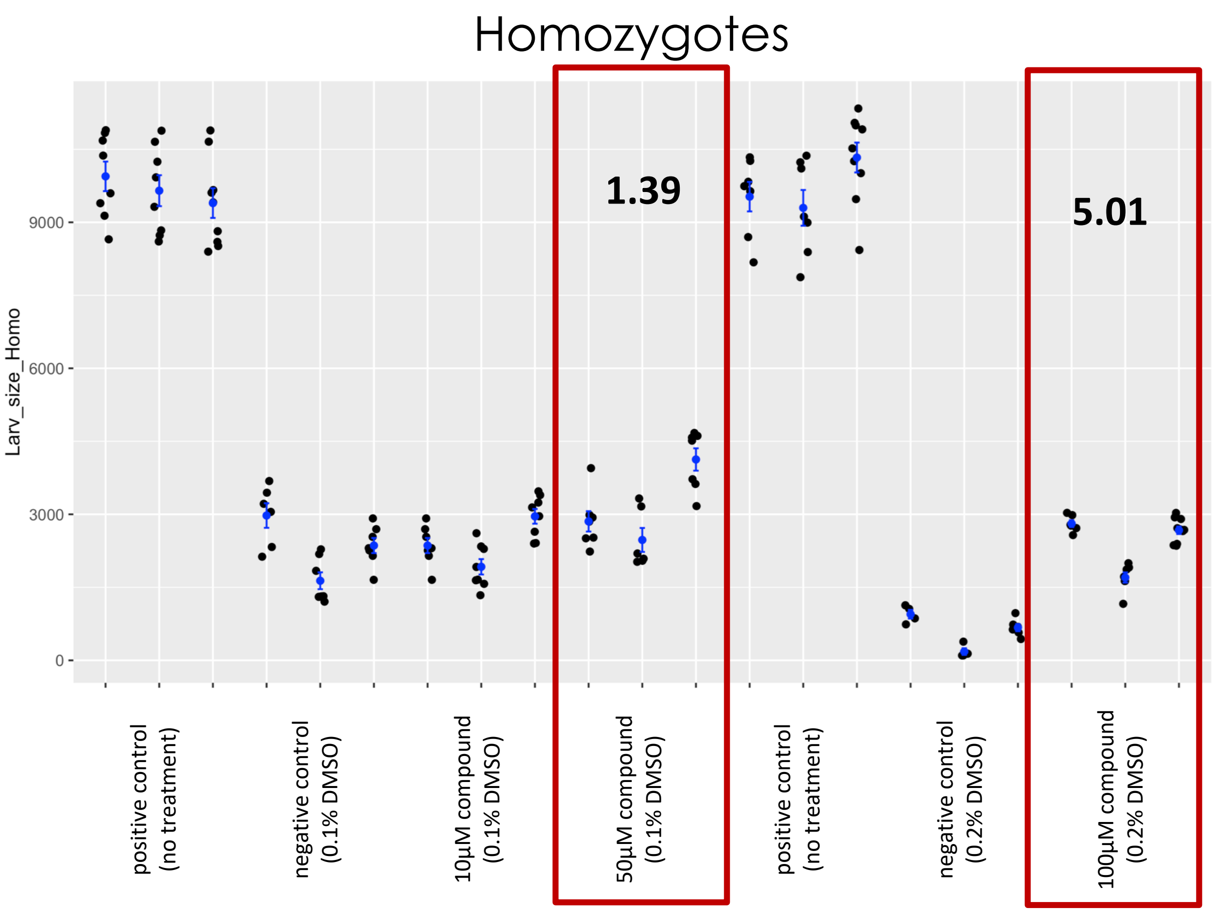


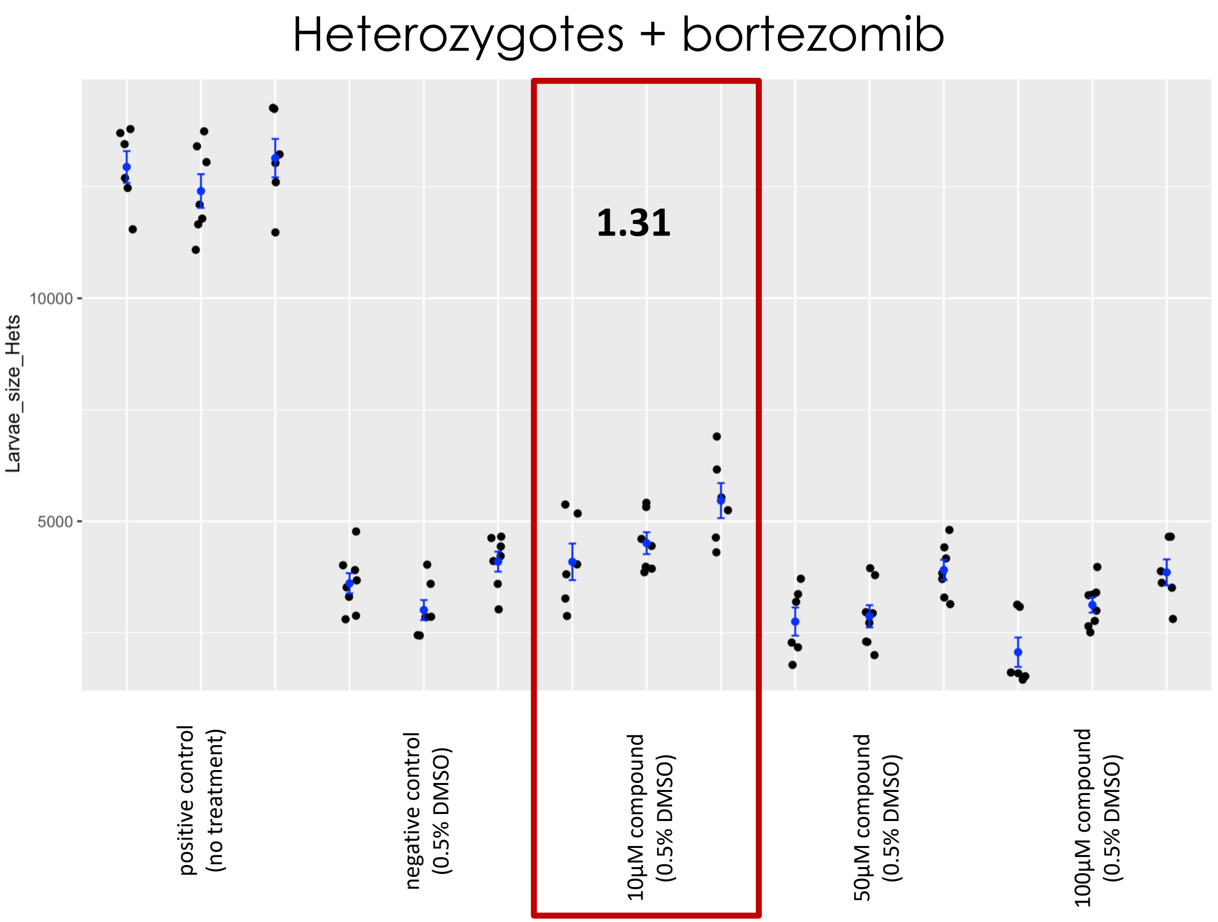

### Fly Microsource Retest – ARIPIPRAZOLE

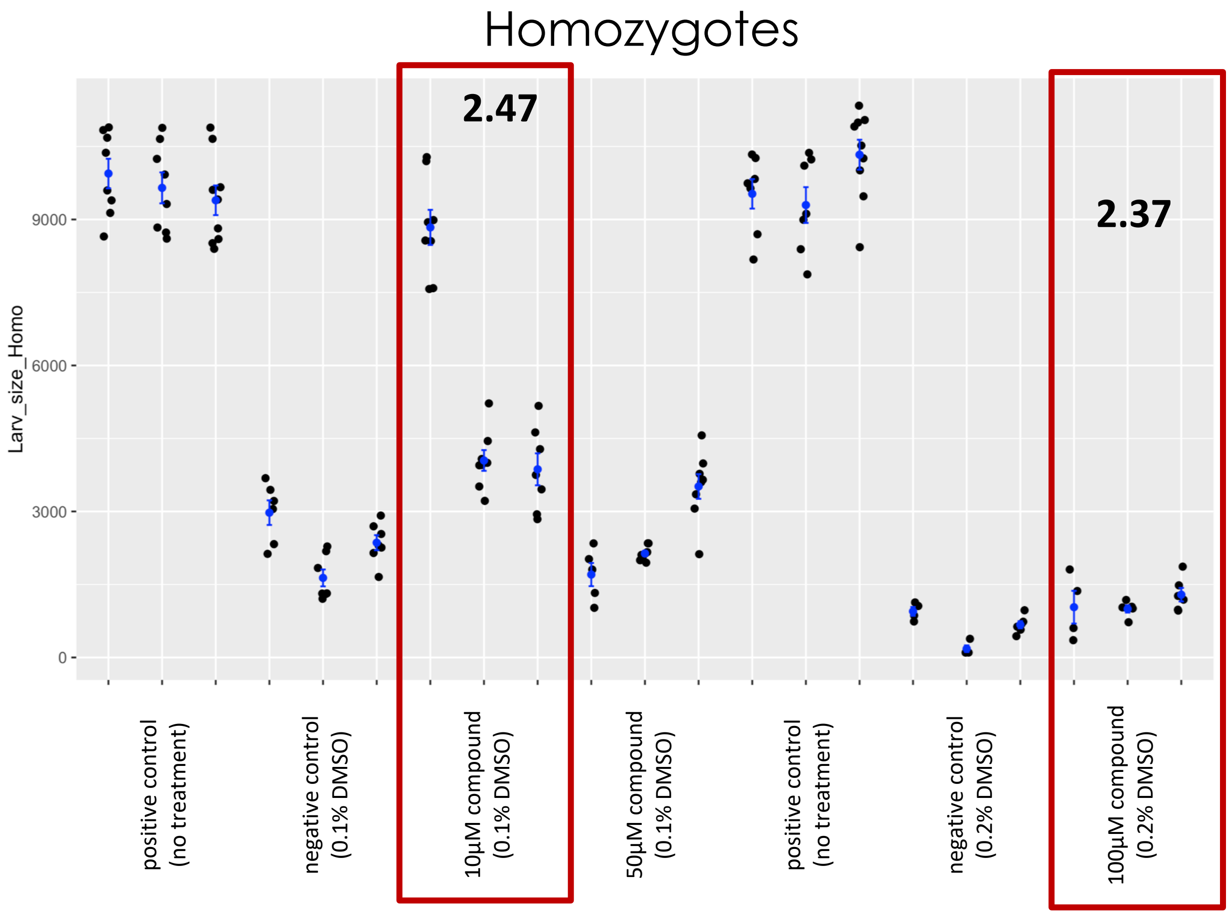


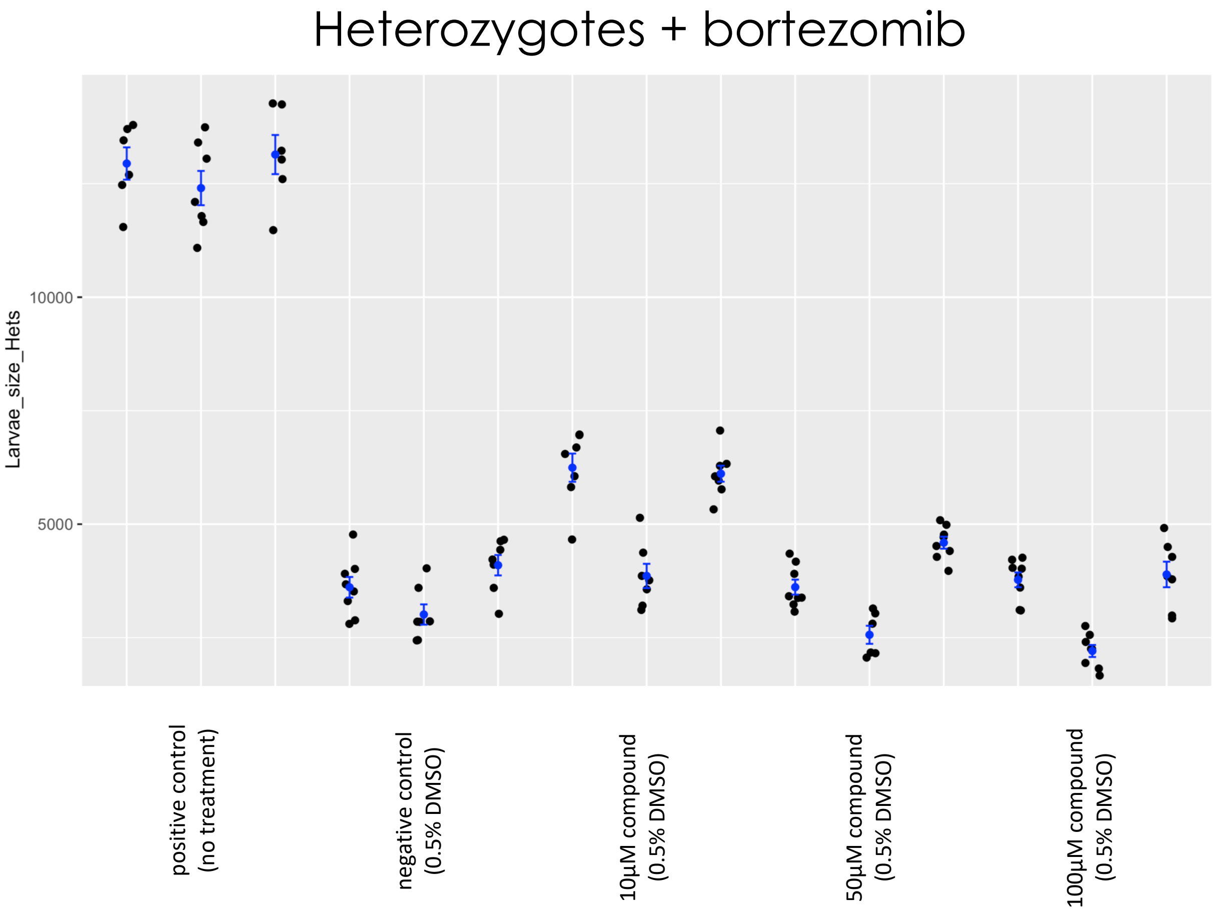

### Fly Microsource Retest – beta-AMYRIN

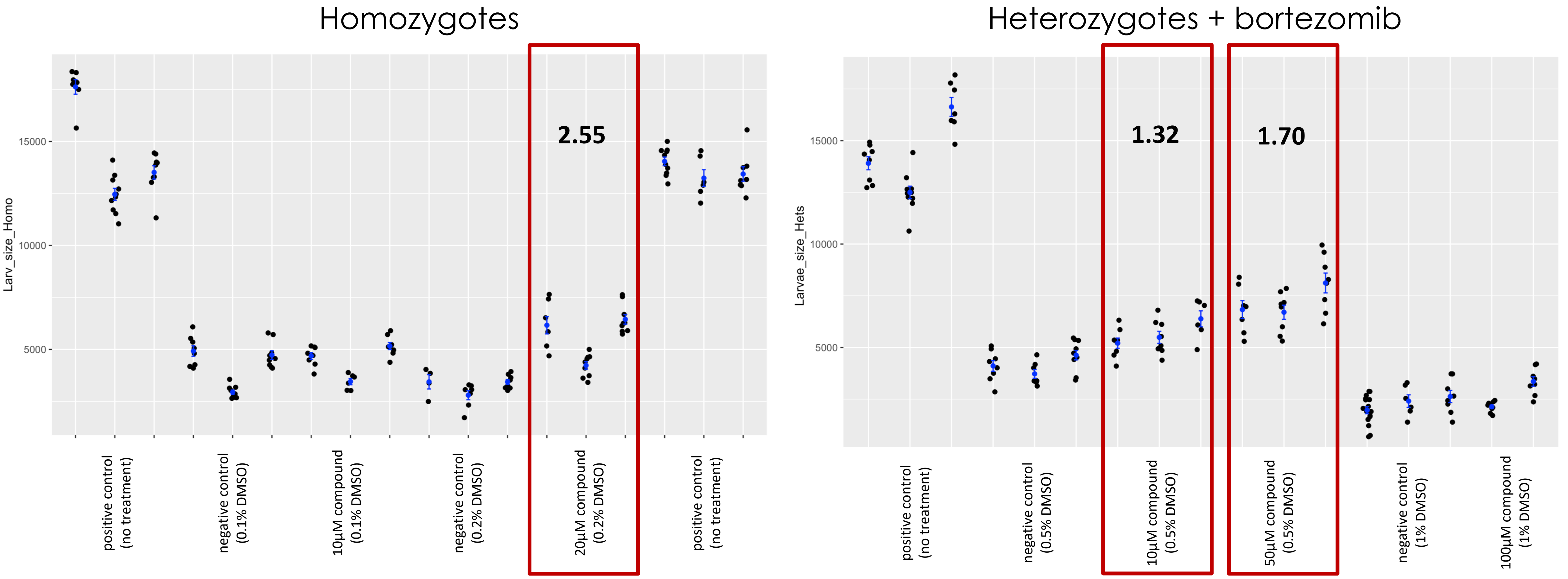

### Fly Microsource Retest – BUTOPYRONOXYL

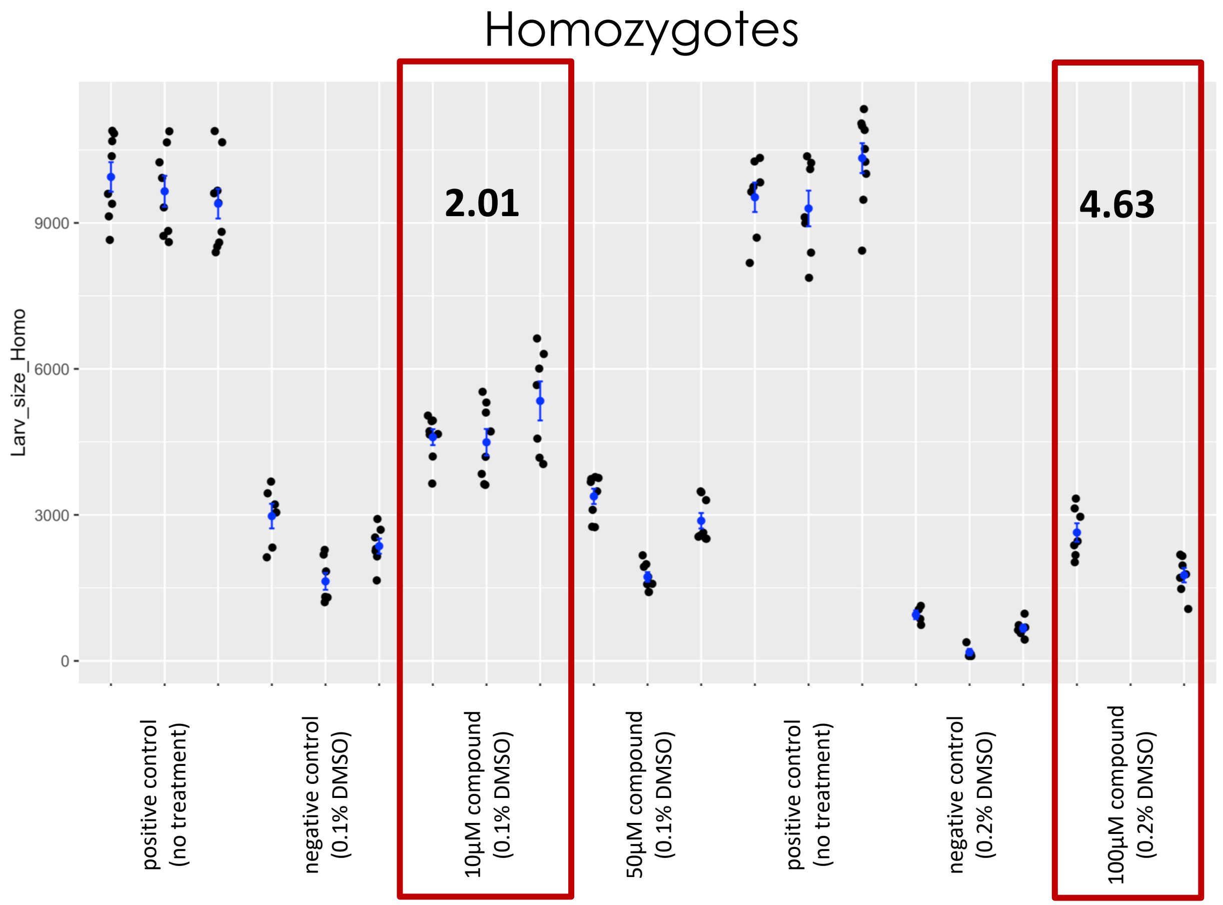


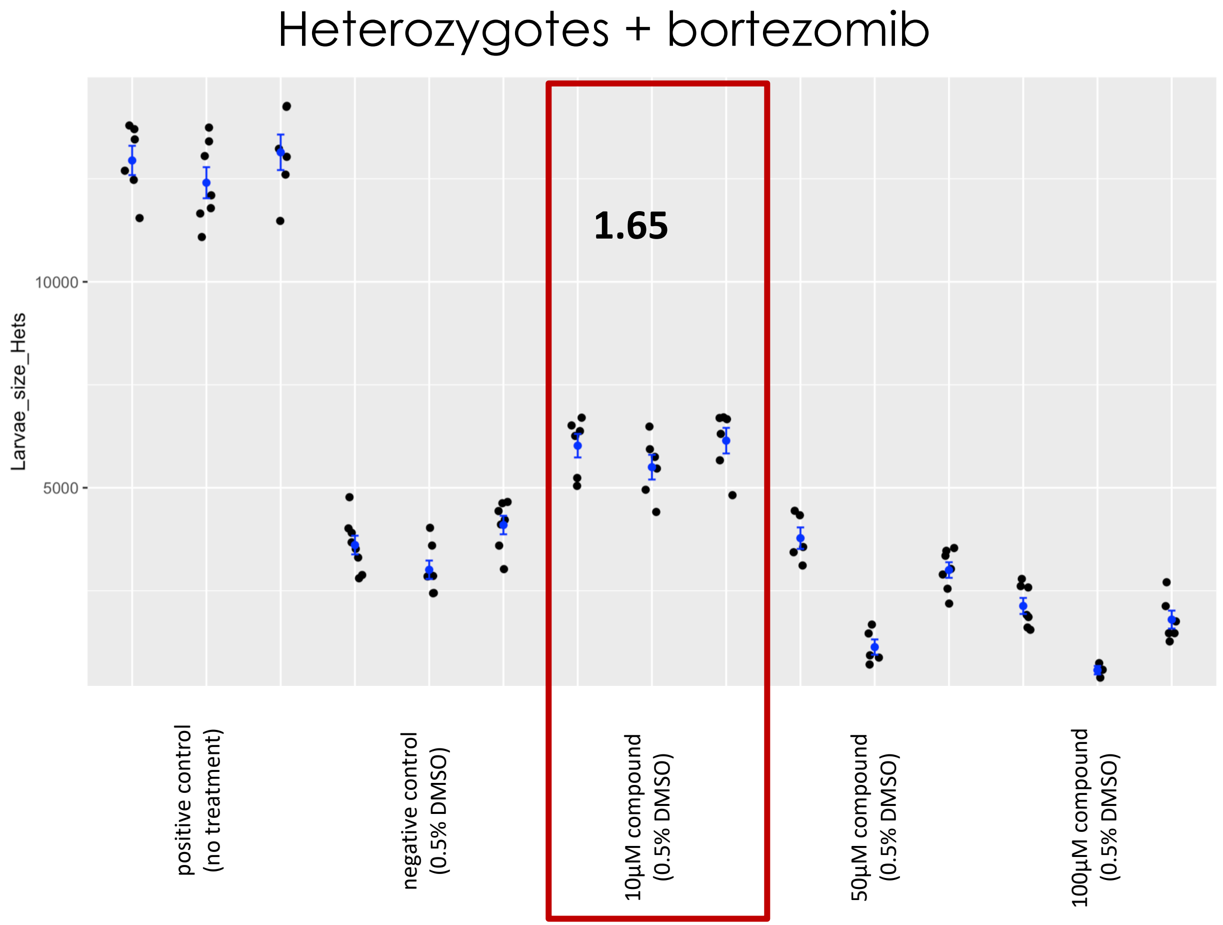

### Fly Microsource Retest – CIMICIFUGOSIDE

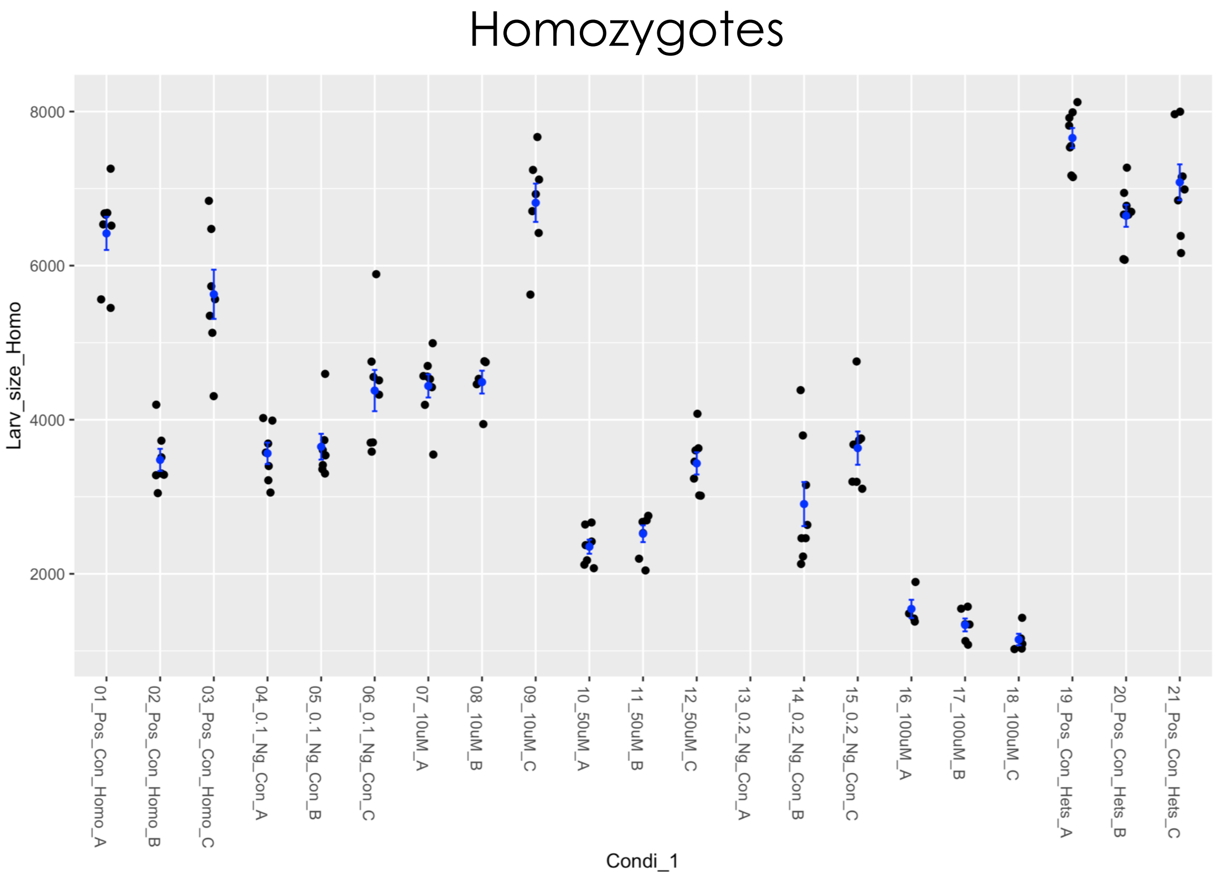


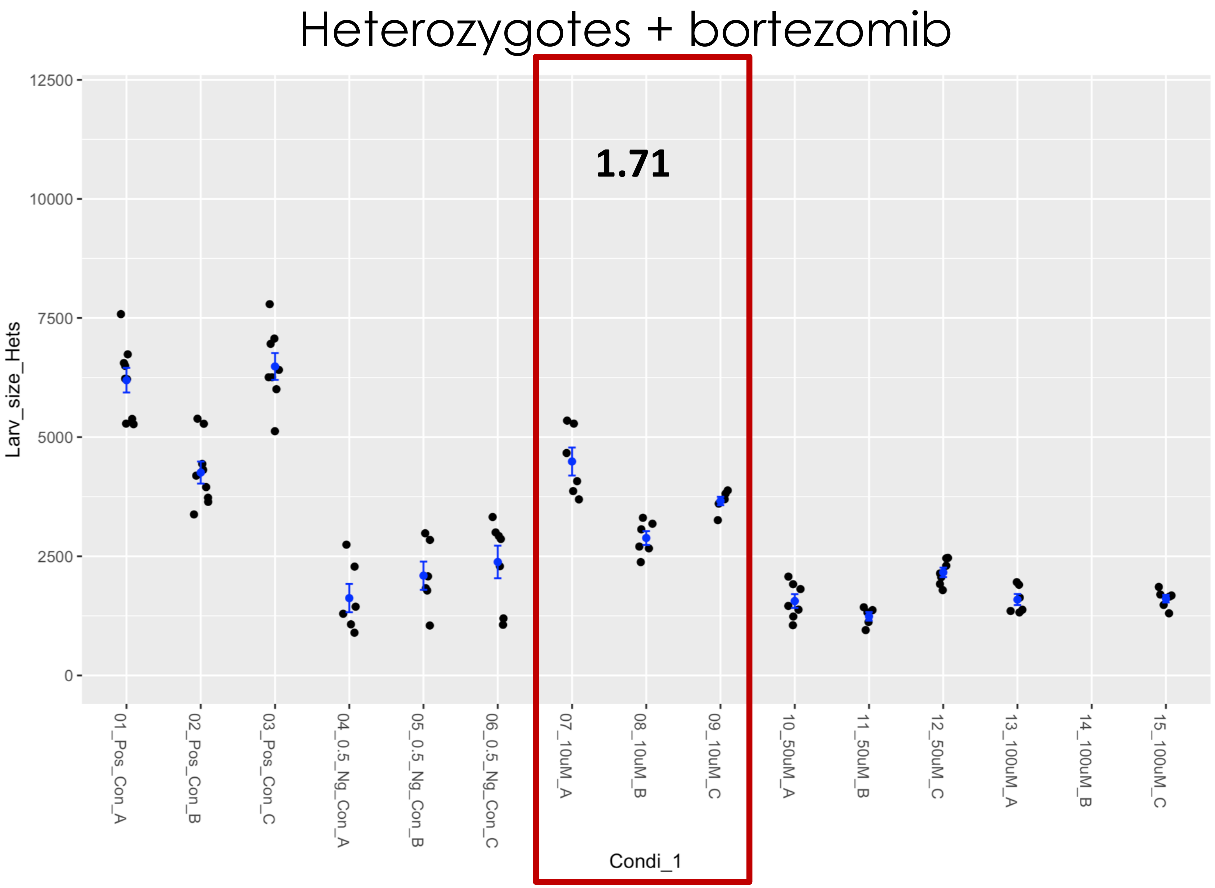

### Fly Microsource Retest – CINNARAZINE

Homozygotes

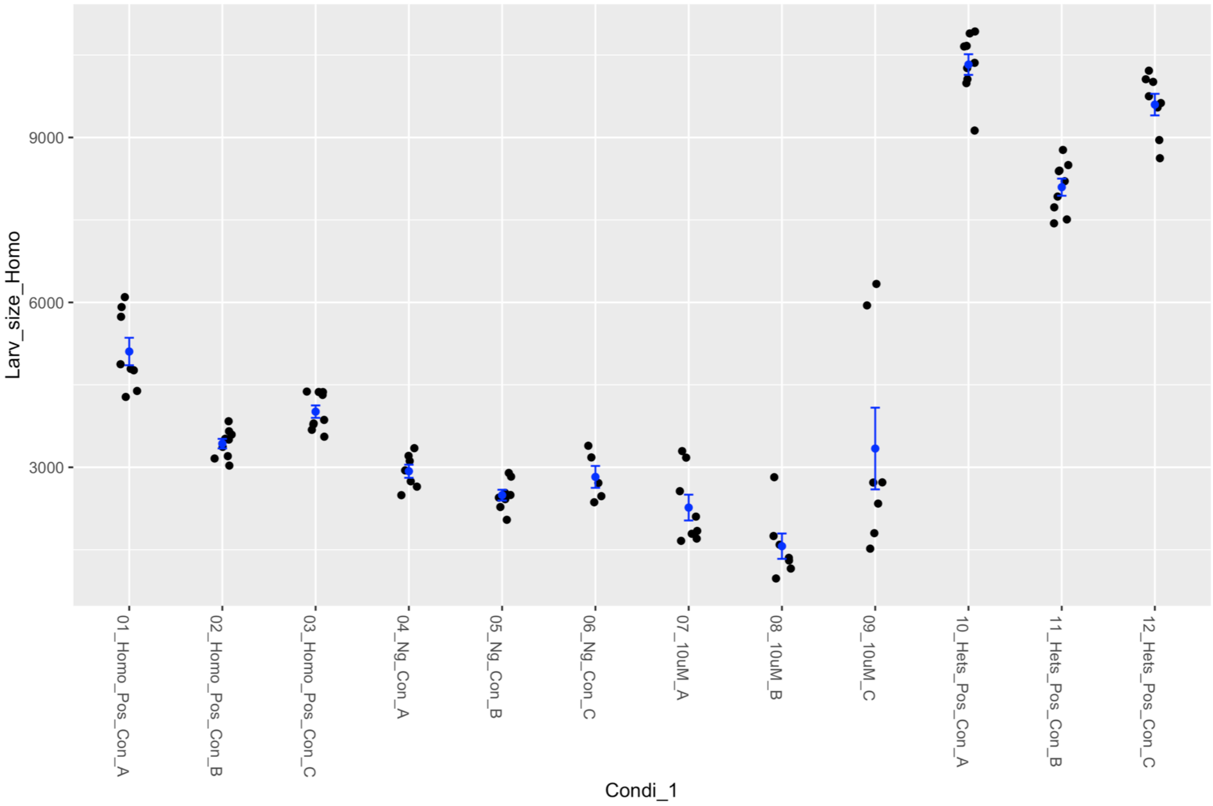


Heterozygotes + bortezomib

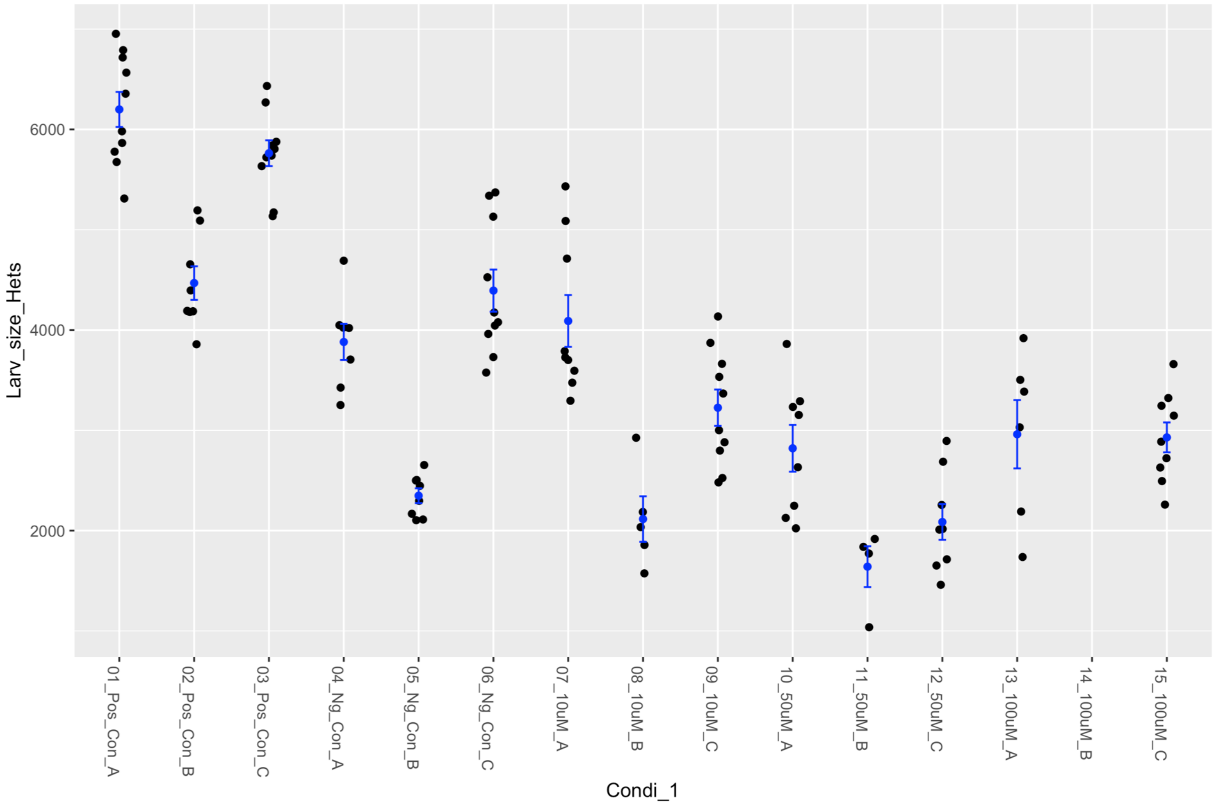

### Fly Microsource Retest – CLINDAMYCIN HYDROCHLORIDE

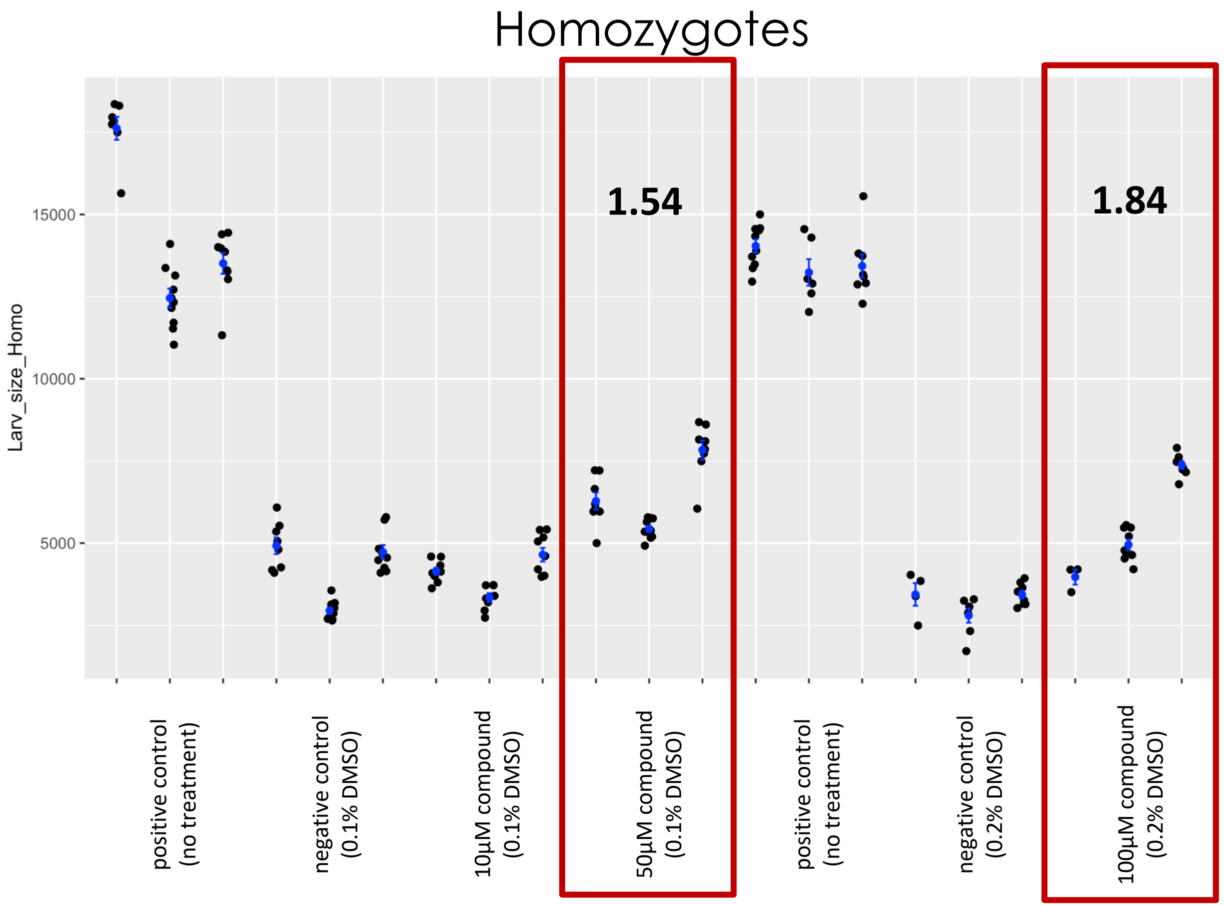


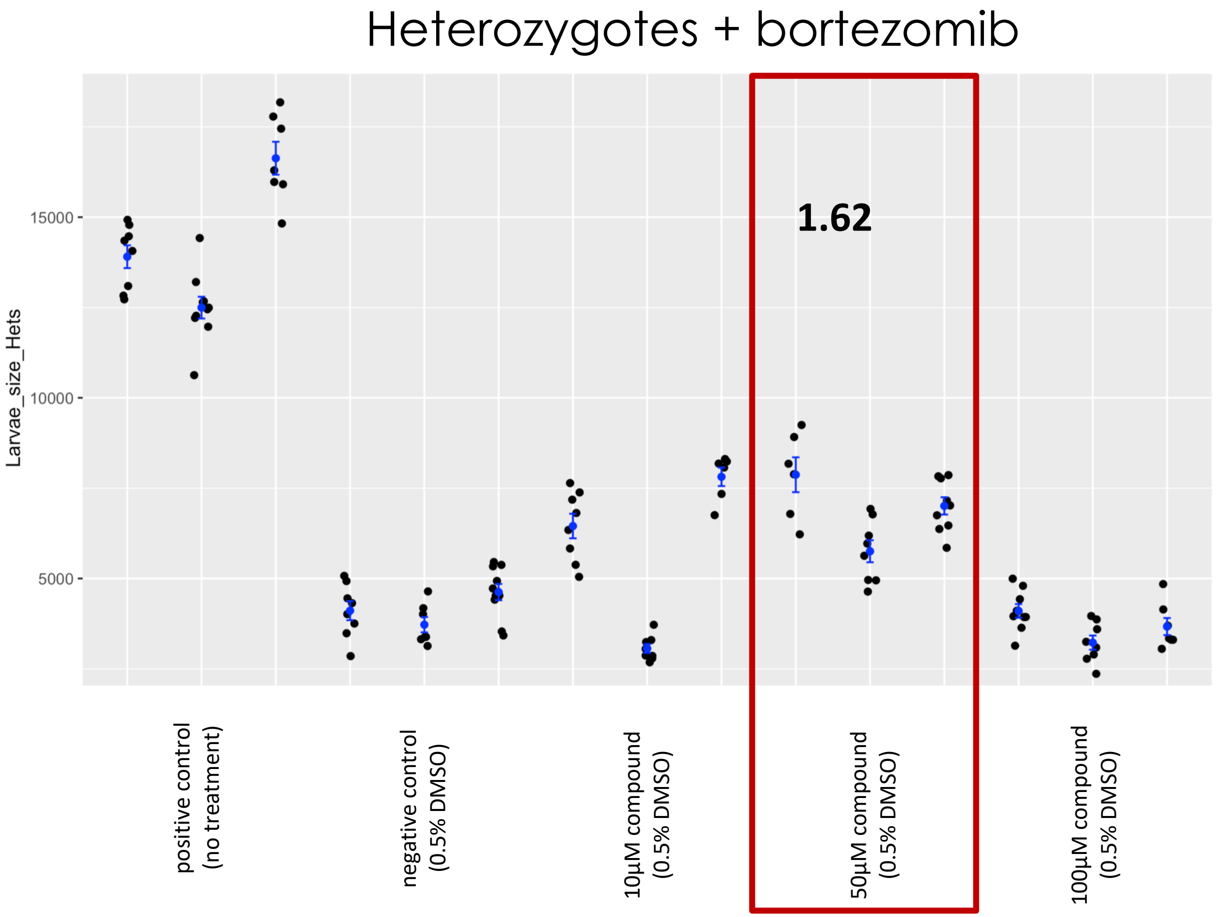

### Fly Microsource Retest – CLOPERASTINE HYDROCHLORIDE

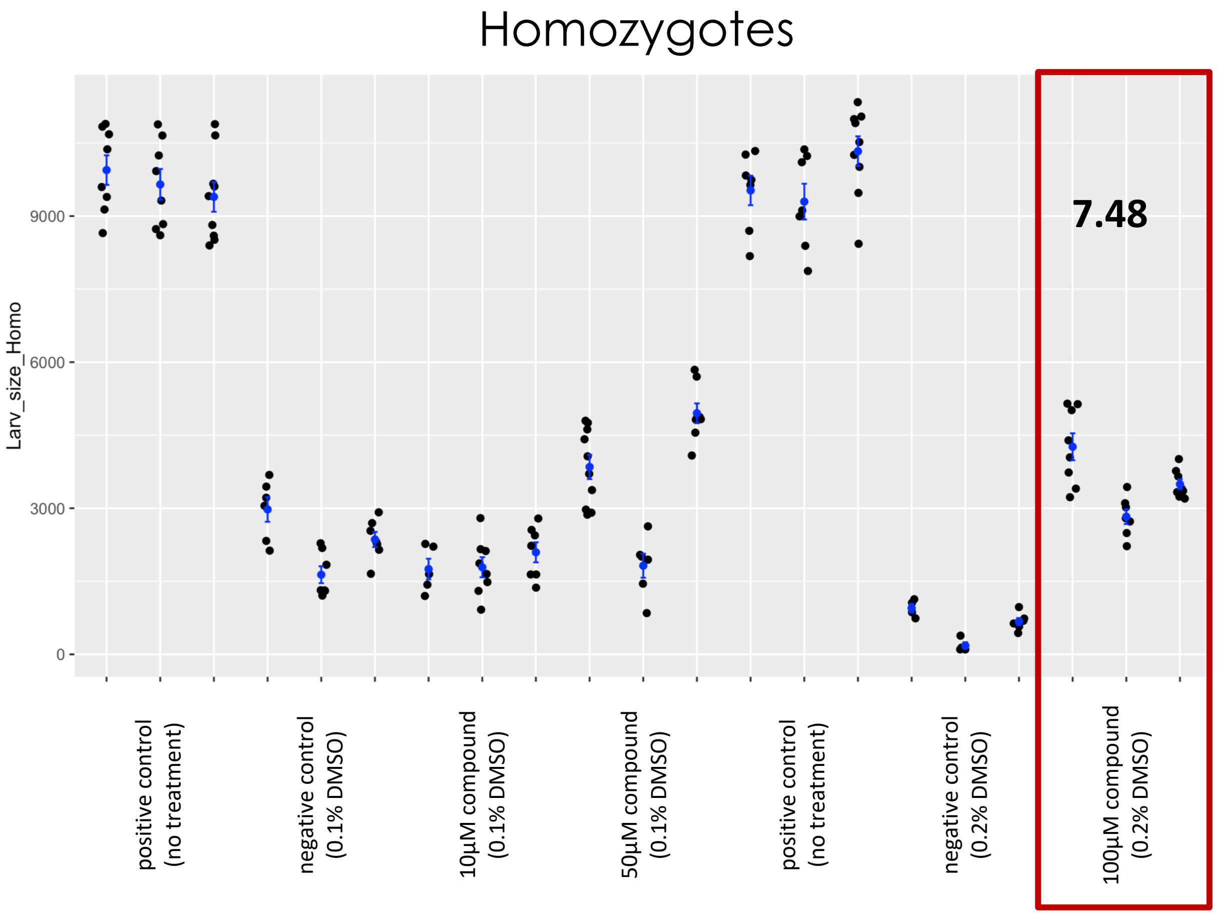


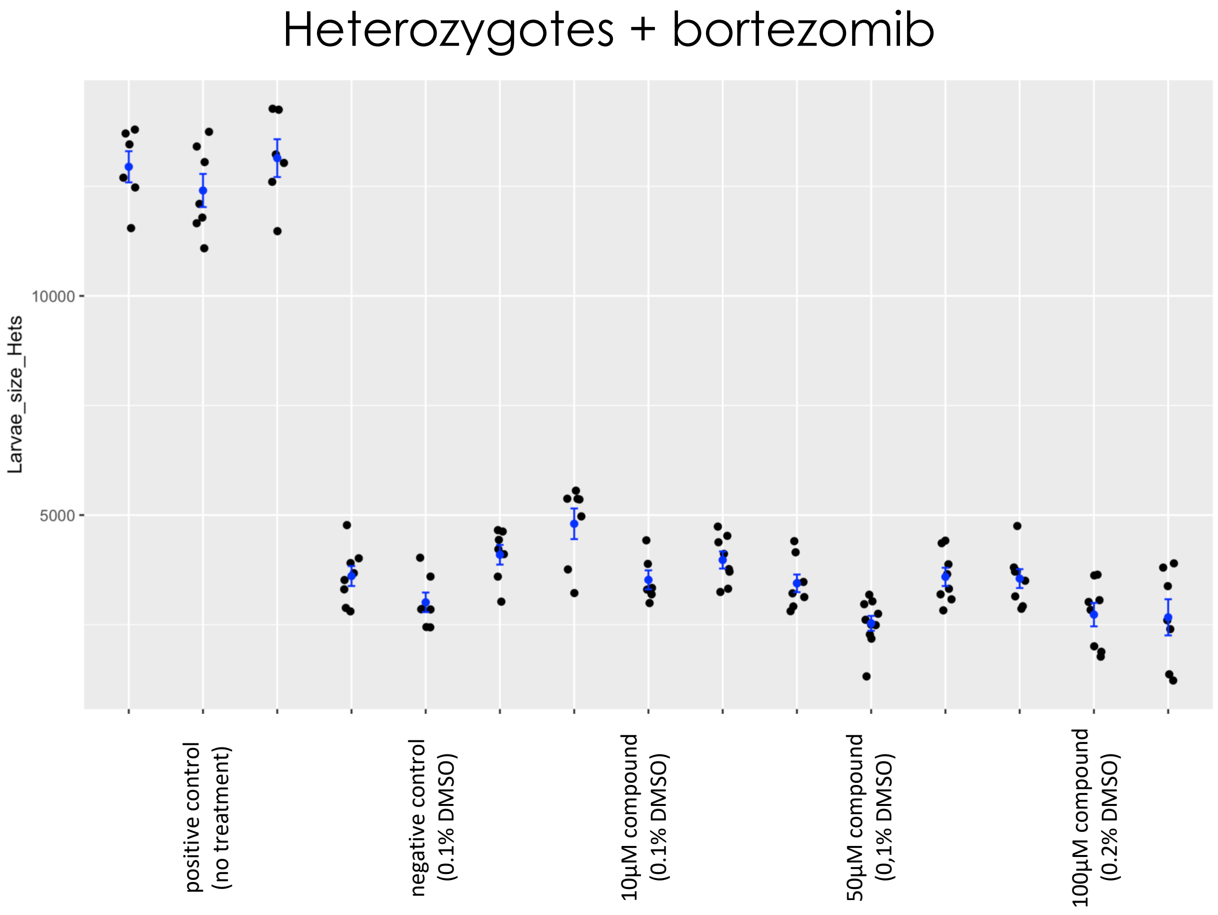

### Fly Microsource retest – CYCLANDELATE

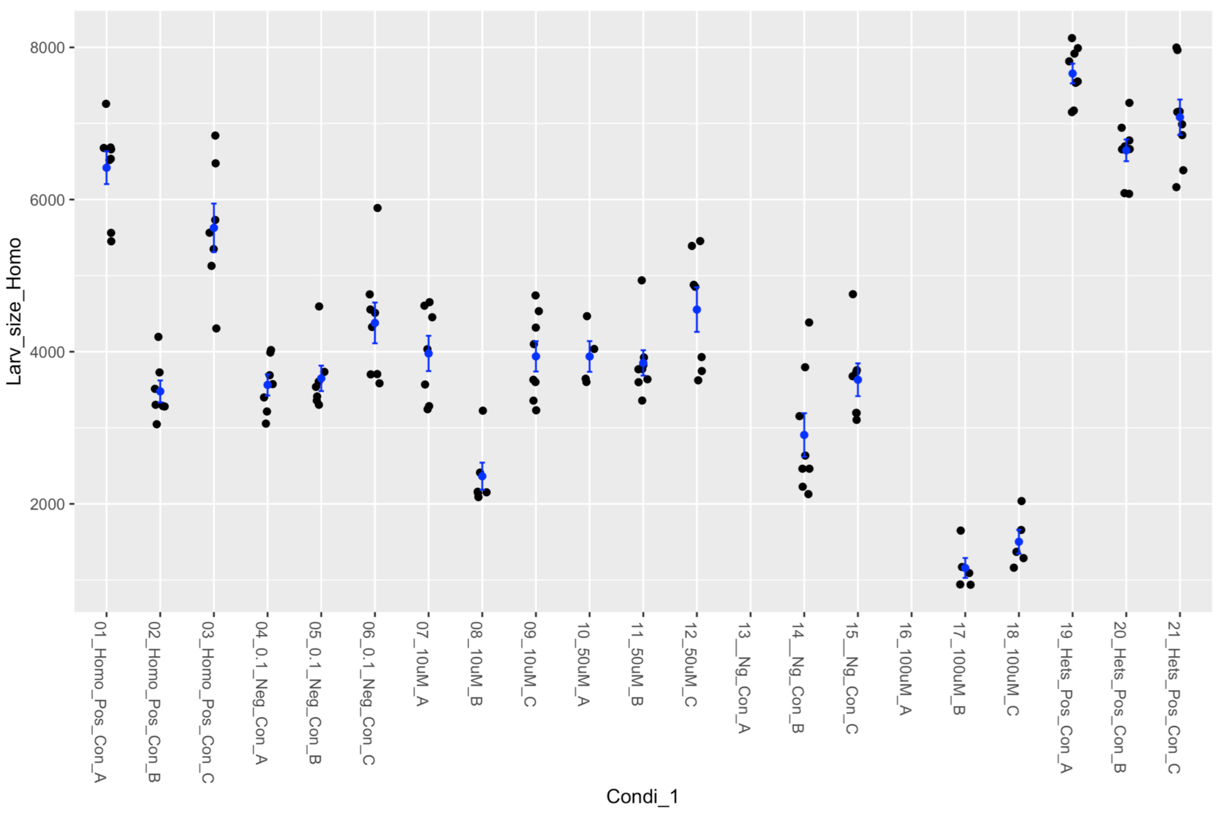


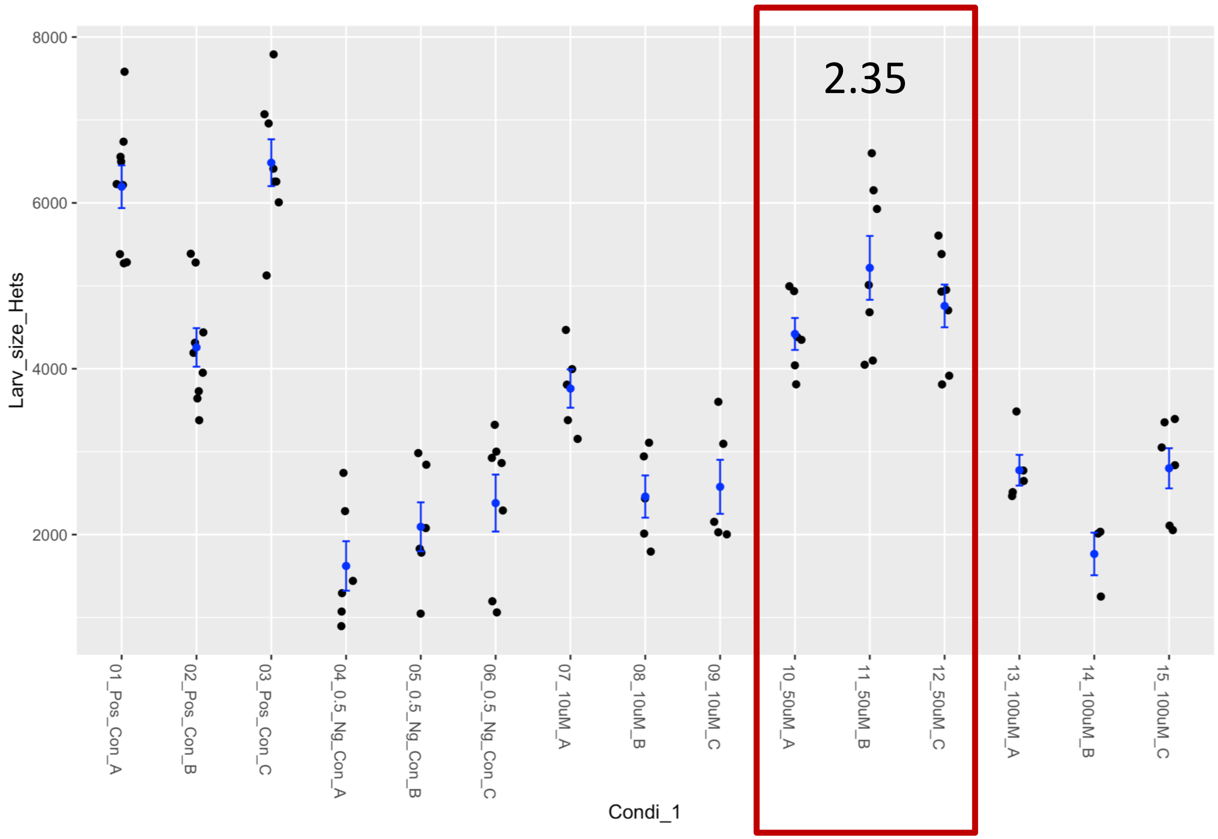

### Fly Microsource retest – CYSTINE

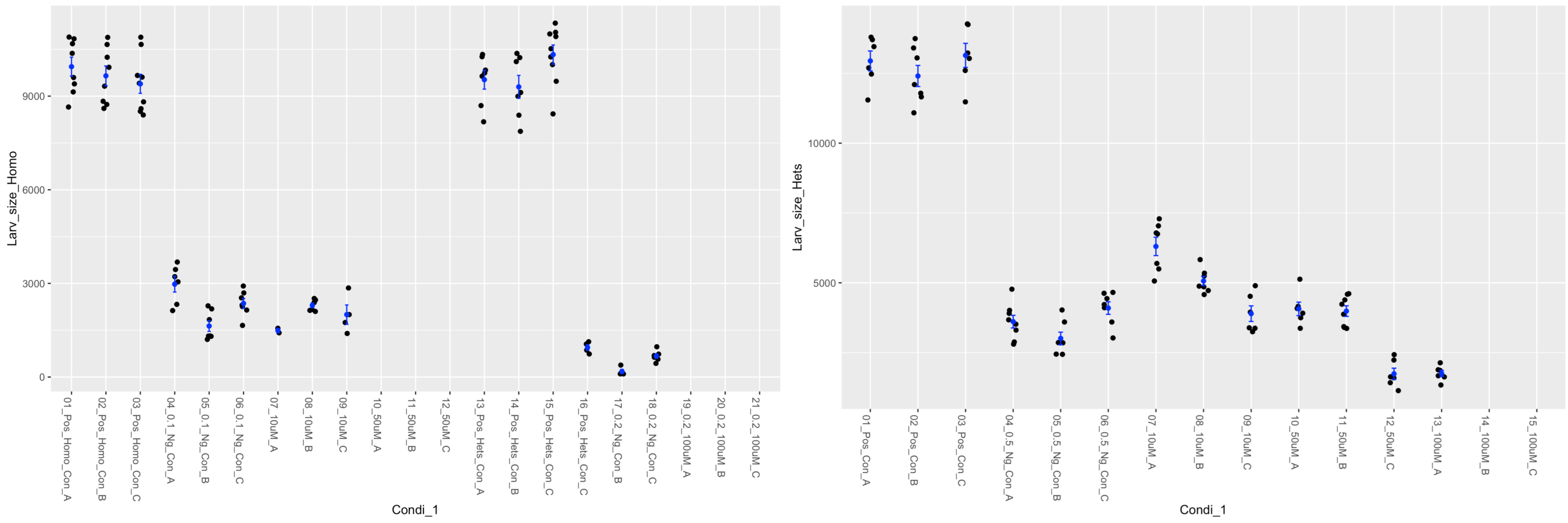

### Fly Microsource Retest – DYPHYLLINE

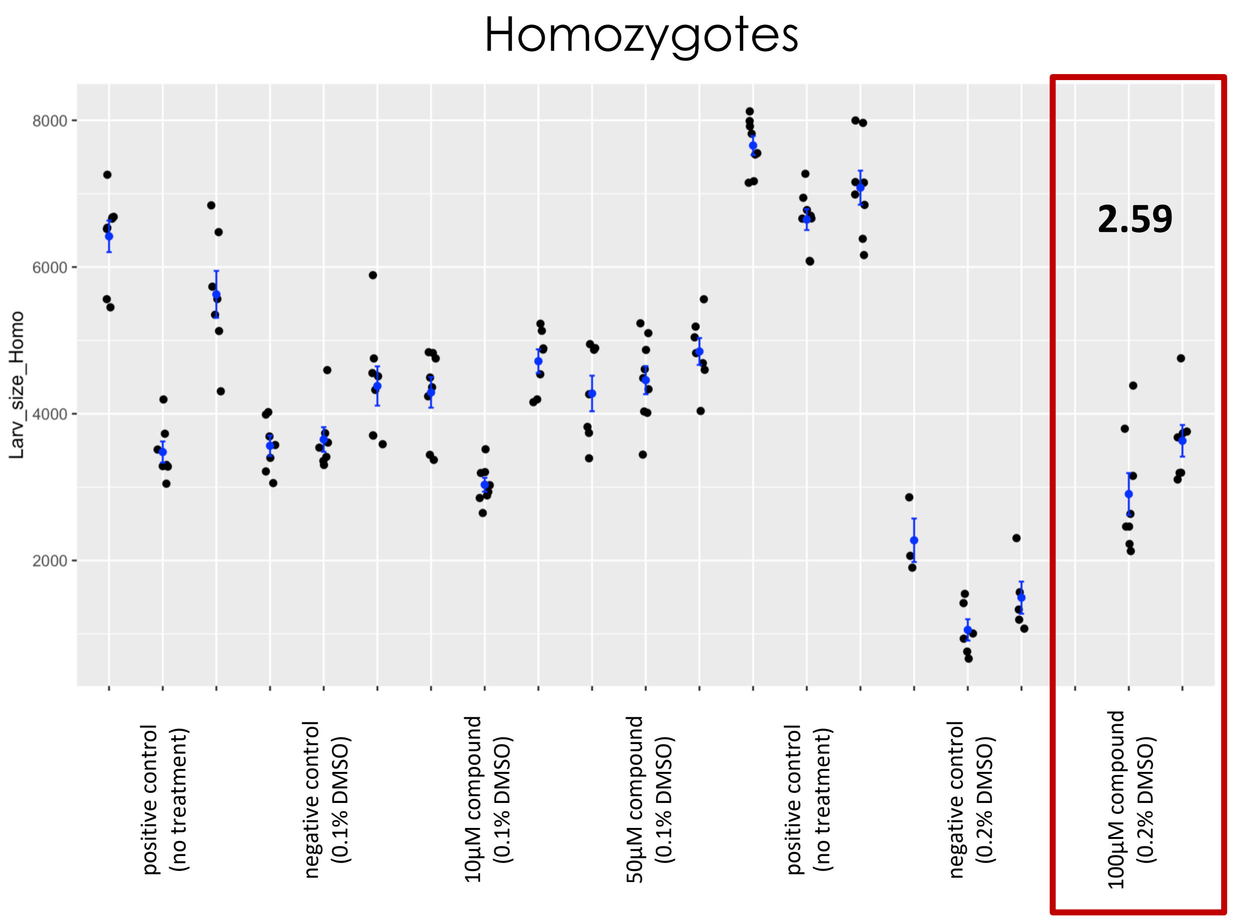


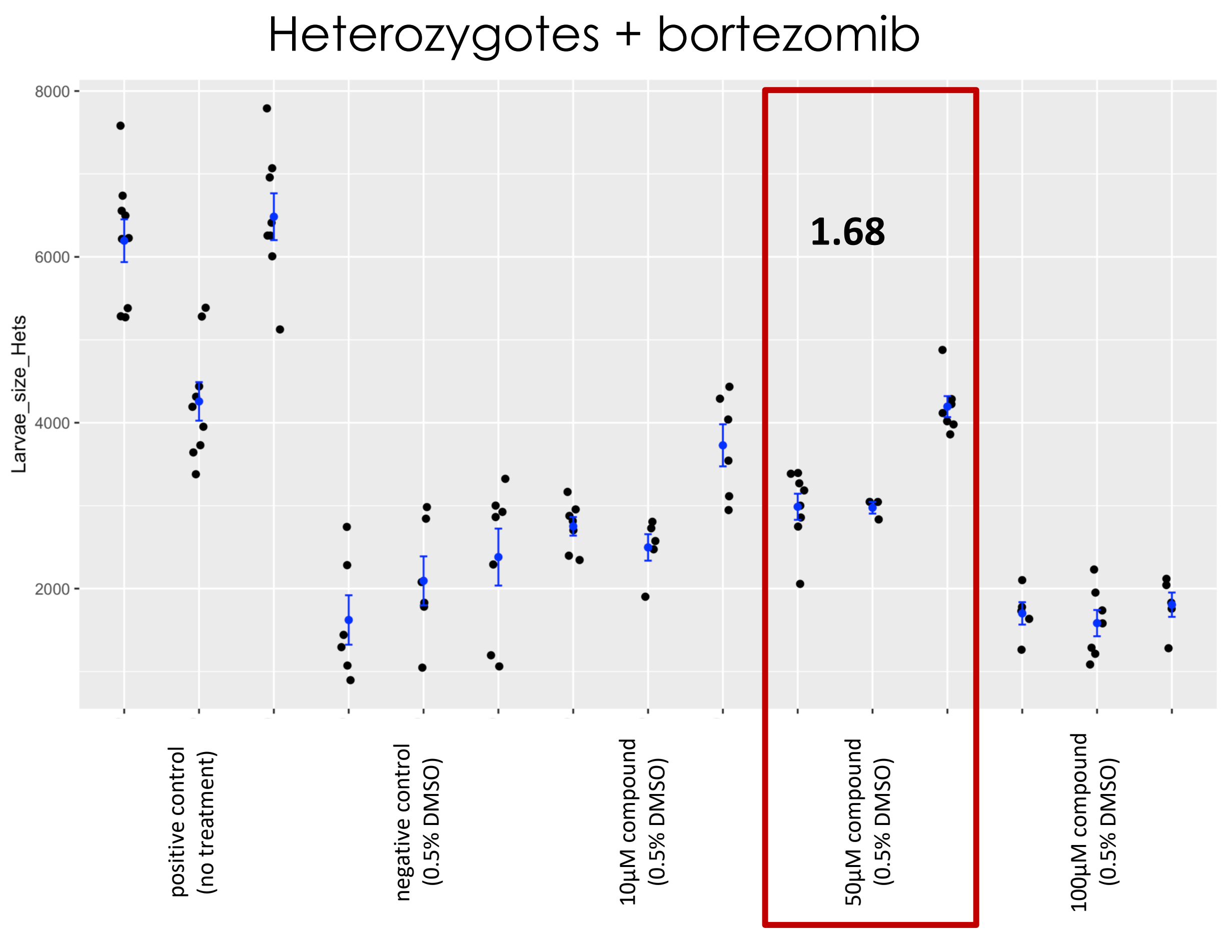

### Fly Microsource Retest – EPALRESTAT

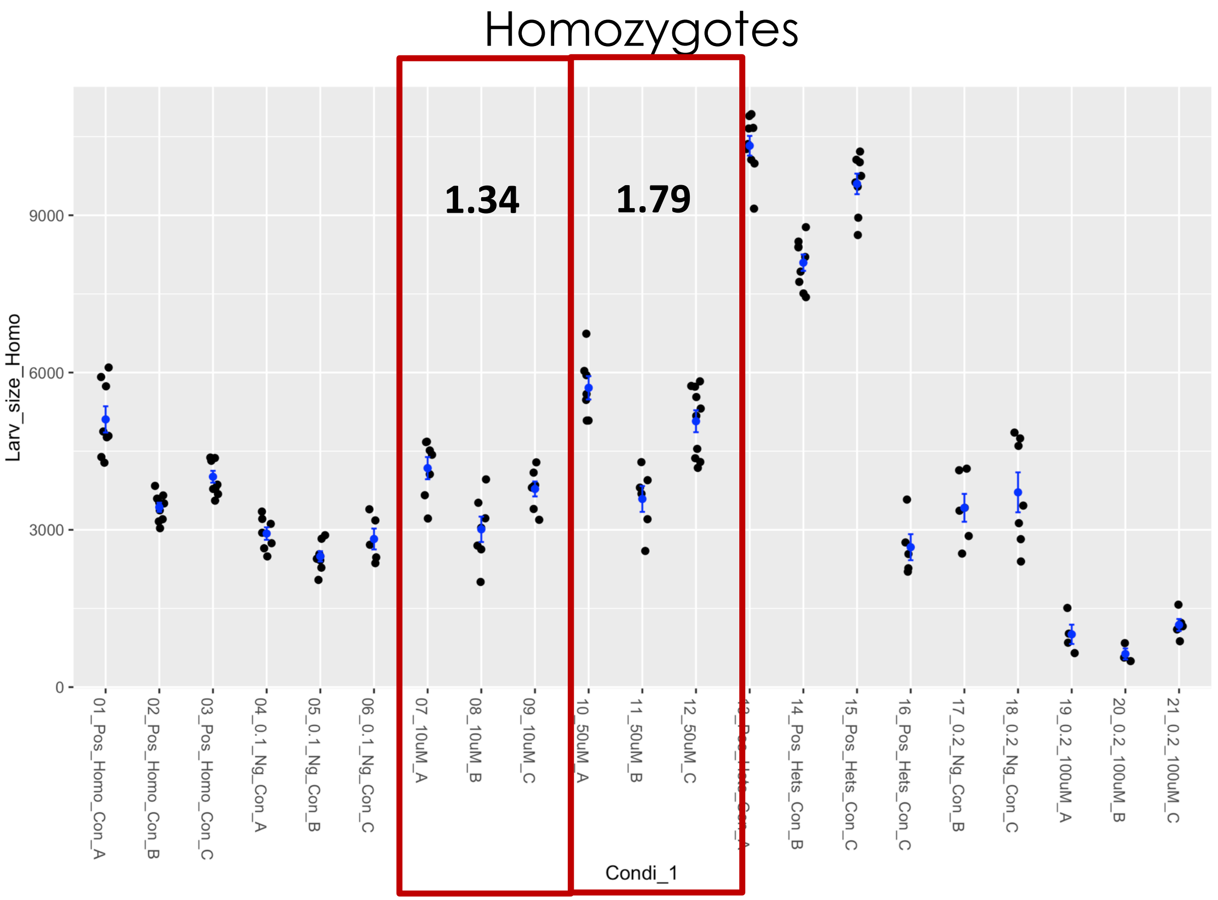


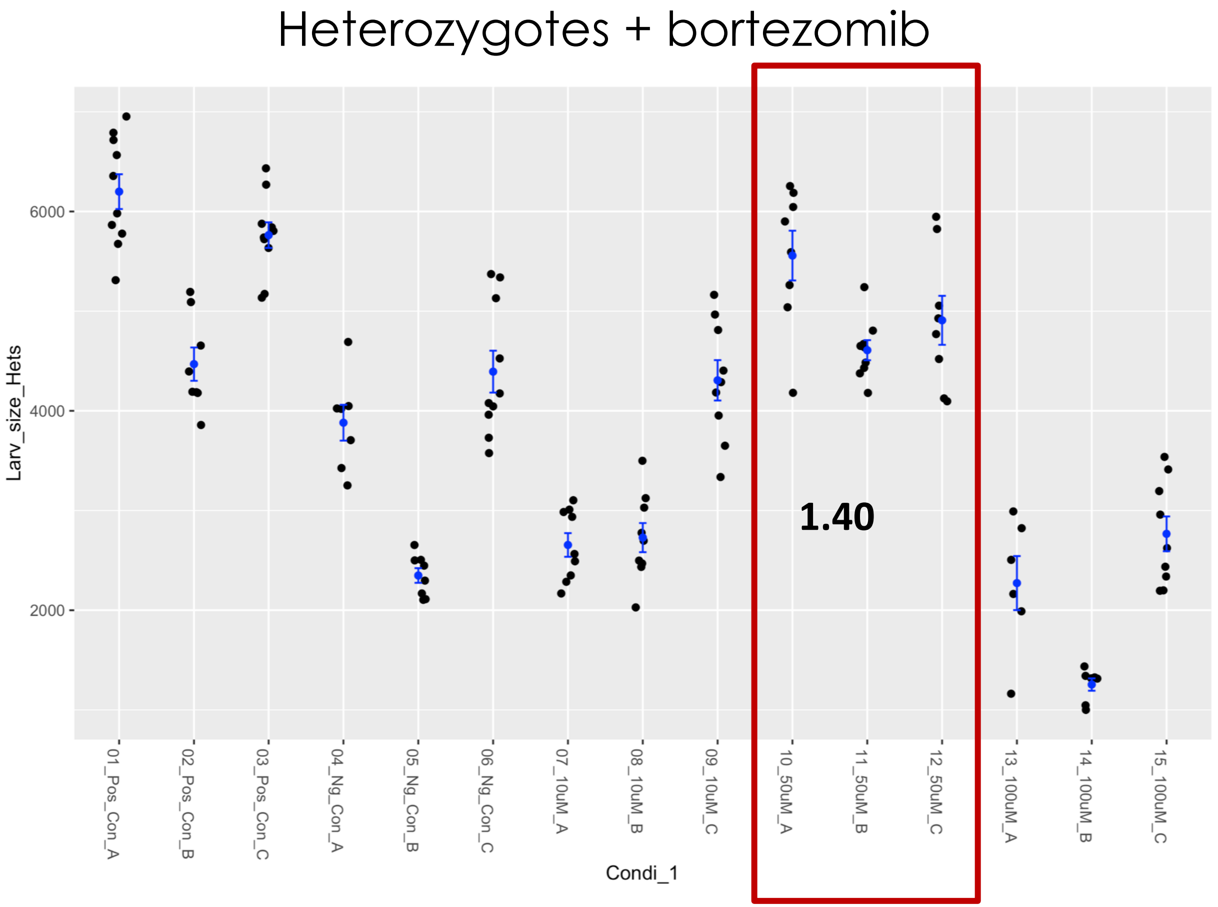

### Fly Microsource retest – ETHAVERINE HYDROCHLORIDE

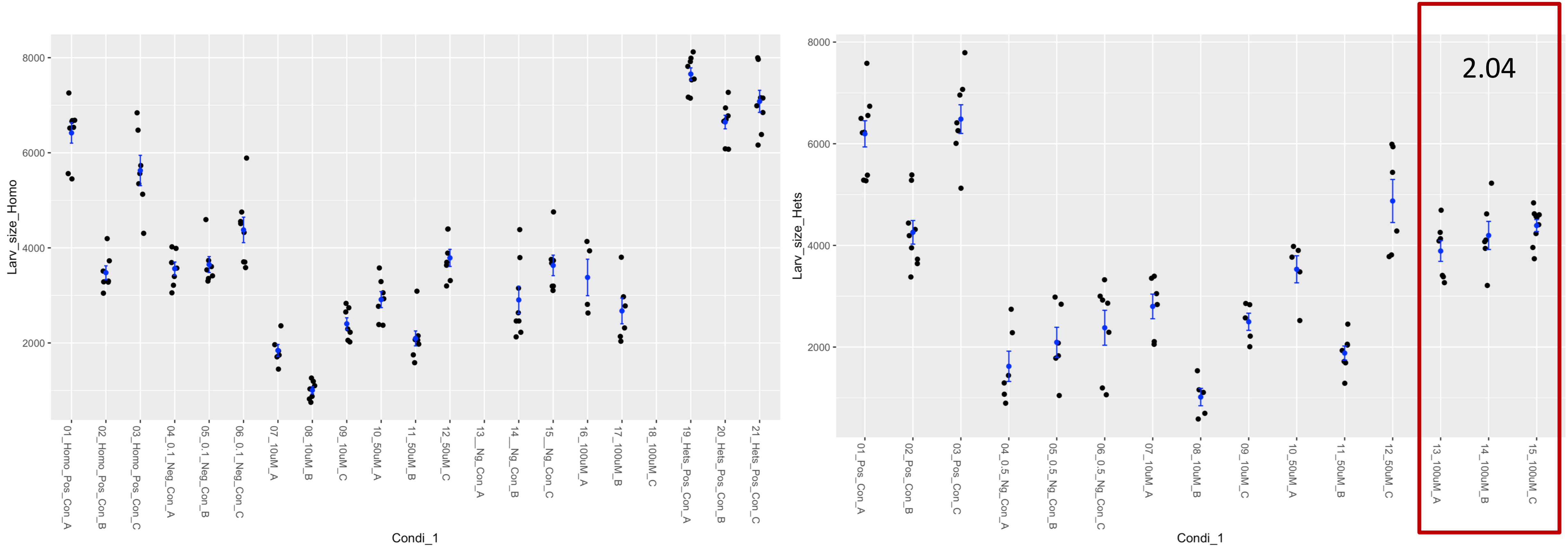

### Fly Microsource Retest – FOMEPIZOLE HYDROCHLORIDE

Homozygotes

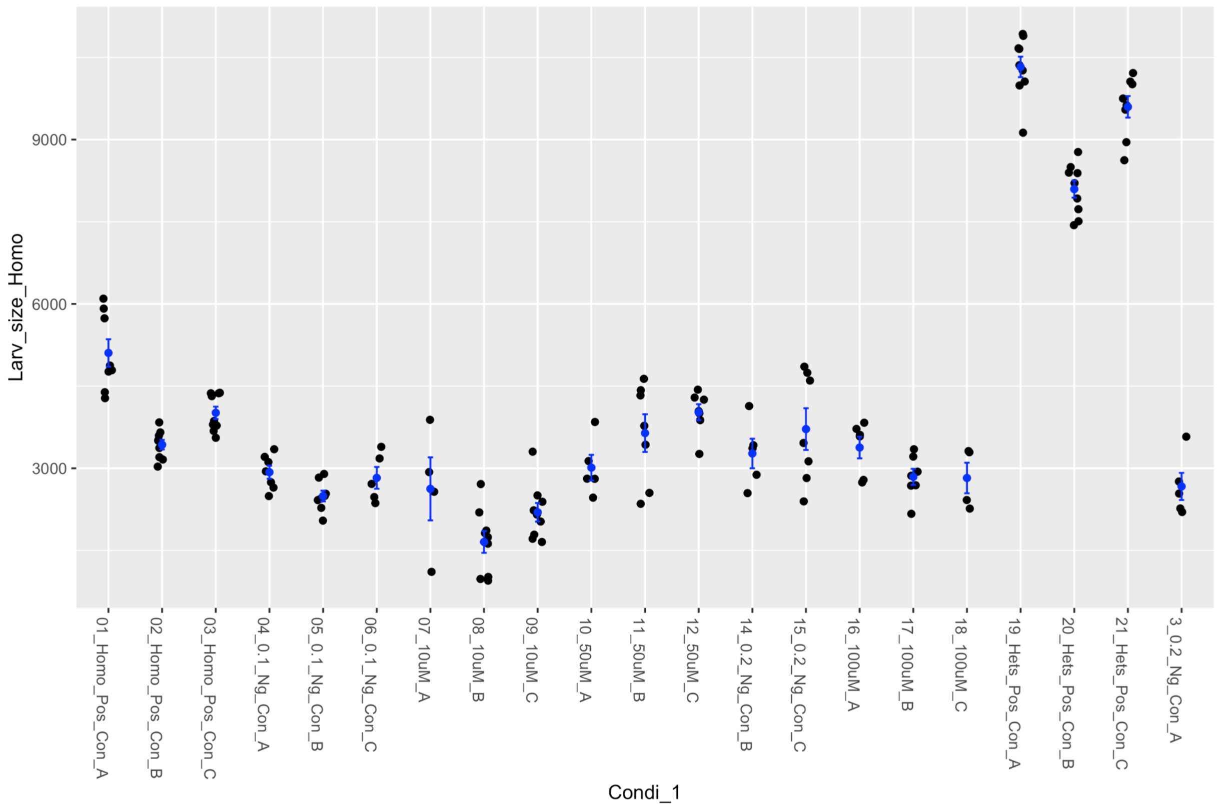


Heterozygotes + bortezomib

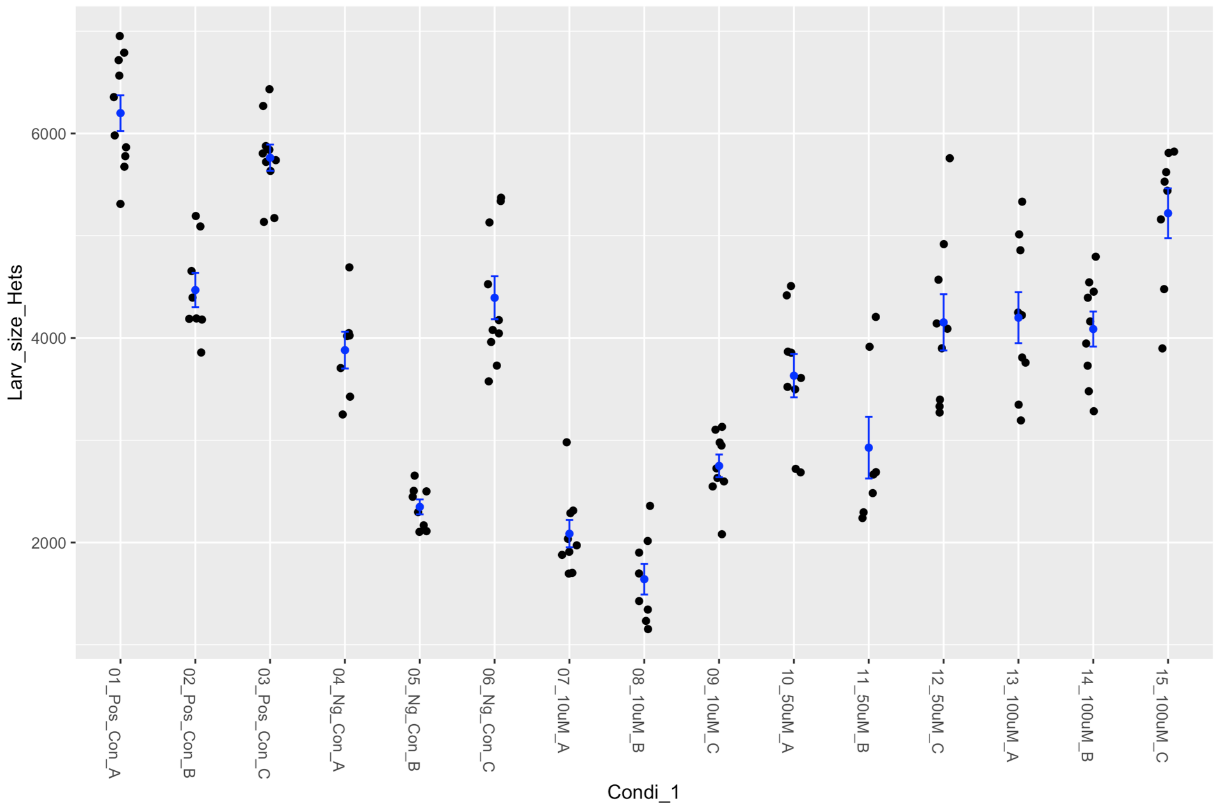

### Fly Microsource Retest – GLUTAMINE (L) HYDROCHLORIDE

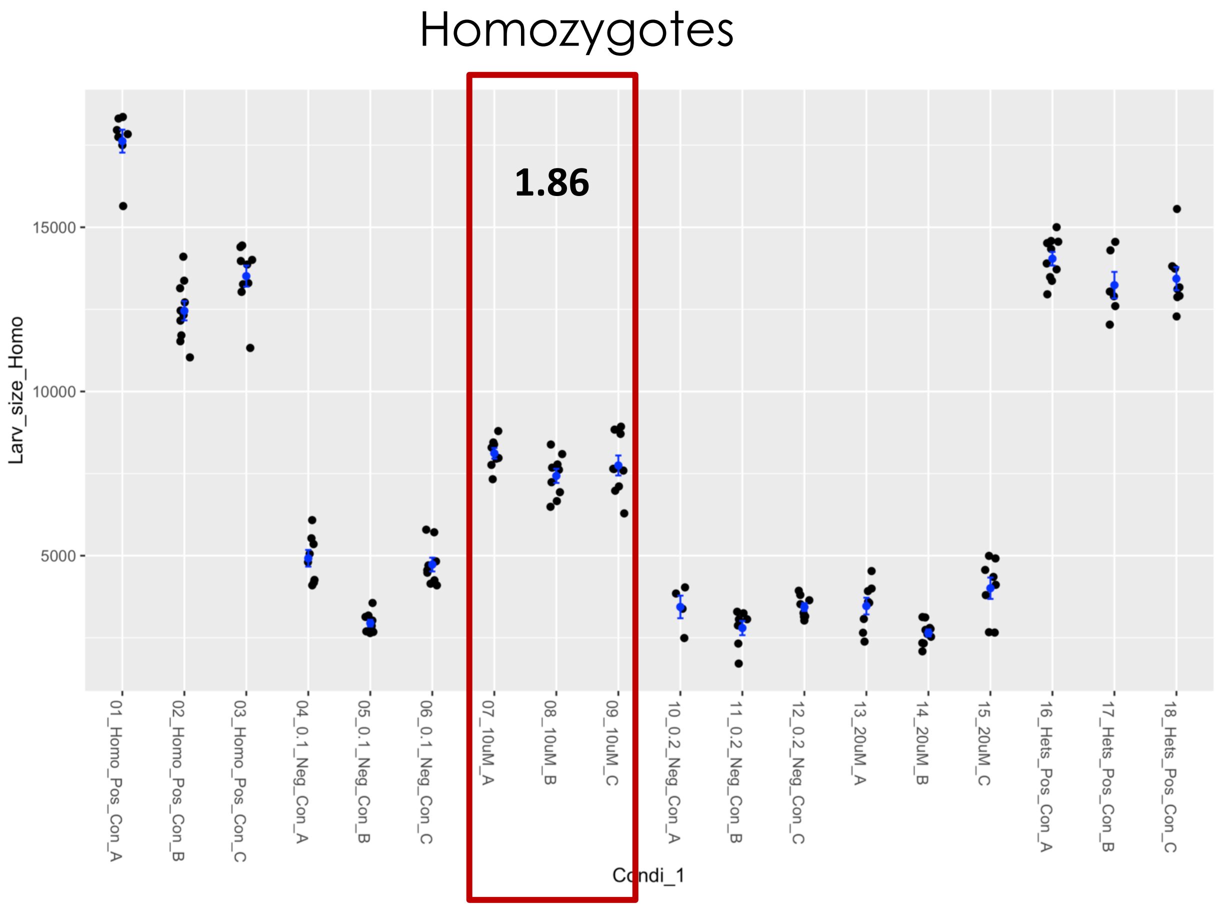


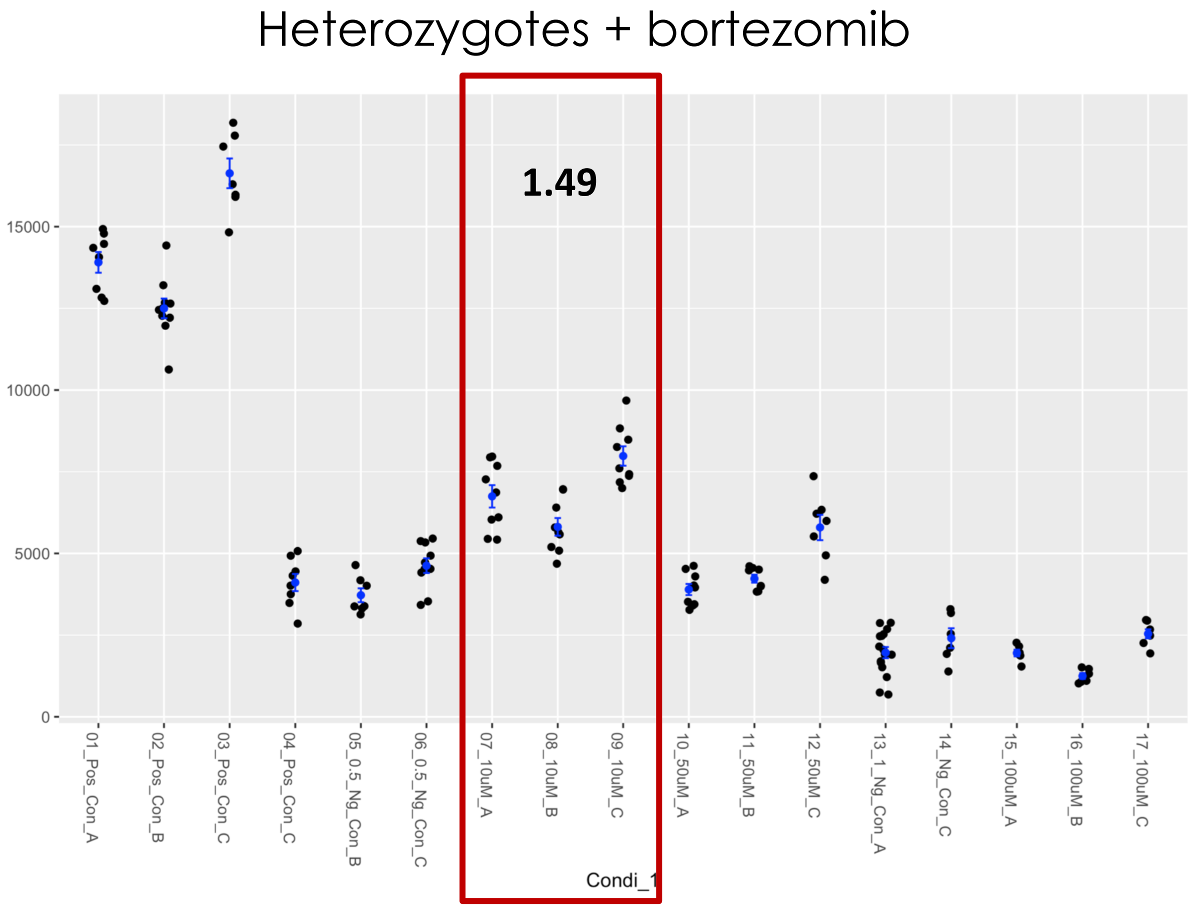

### Fly Microsource Retest – GOSSYPETIN

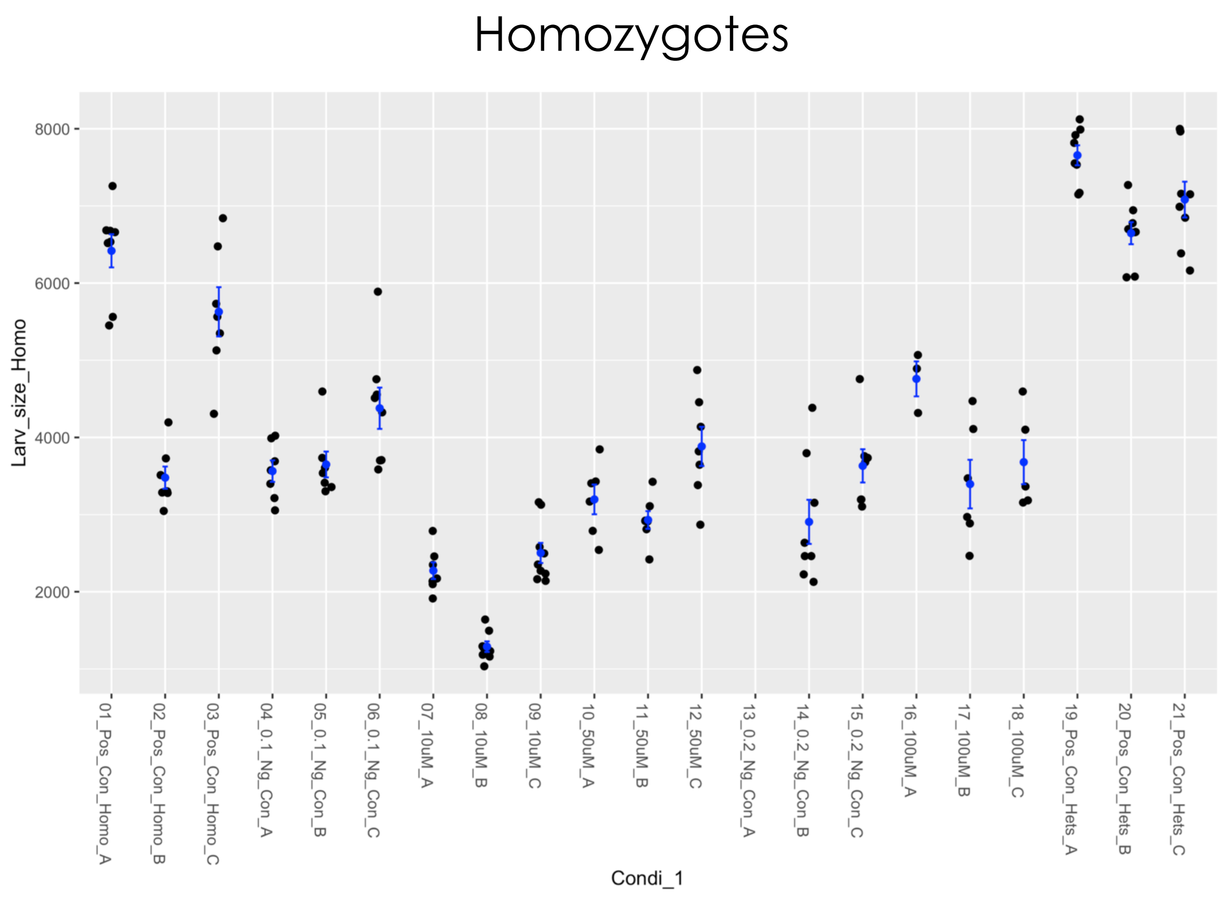


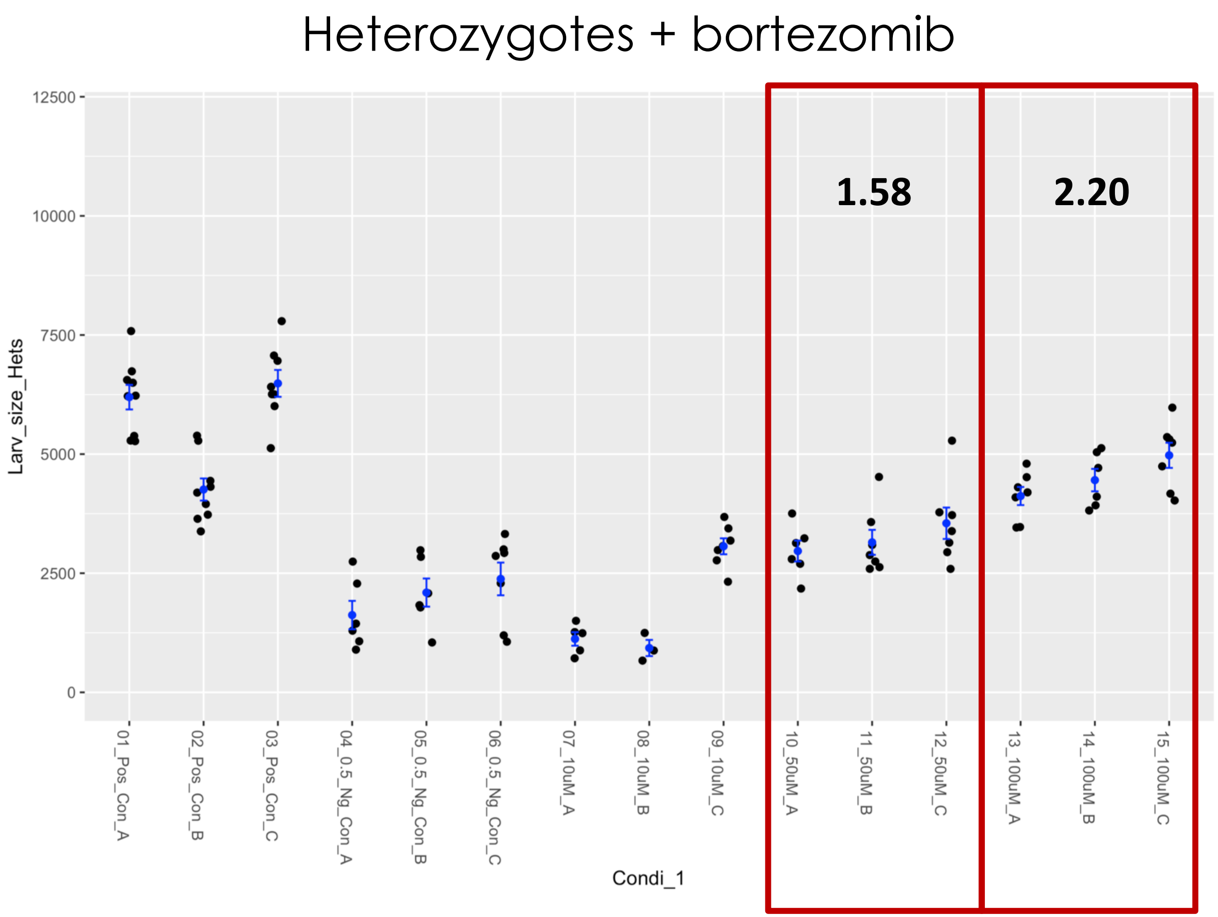

### Fly Microsource retest – HAEMATOXYLIN

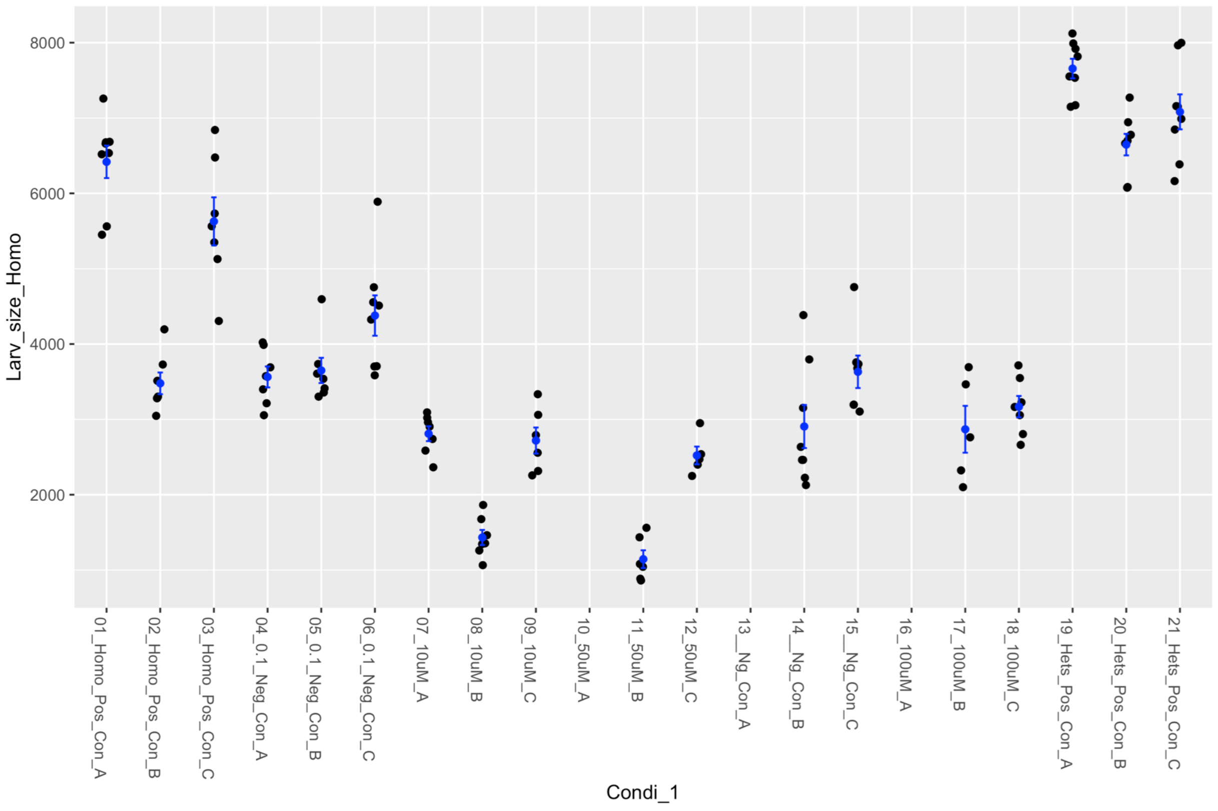


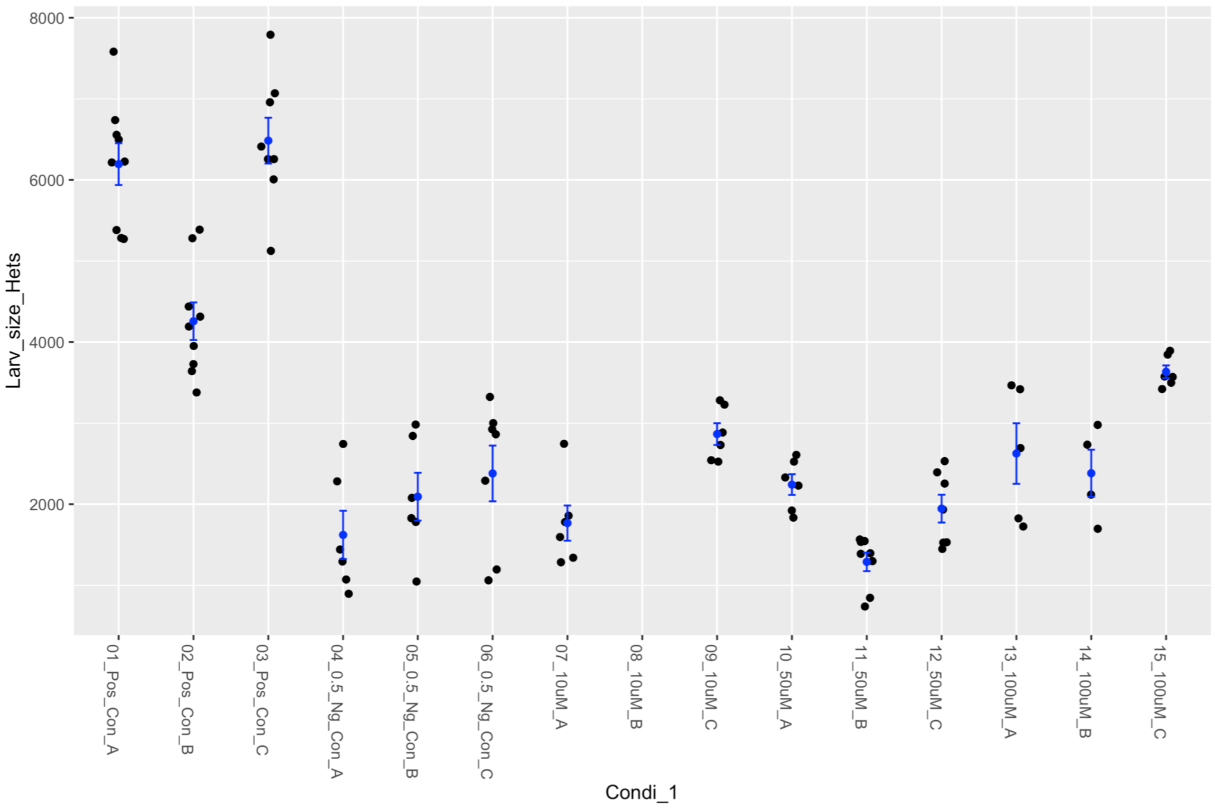

### Fly Microsource Retest – KETOROLAC TROMETHAMINE

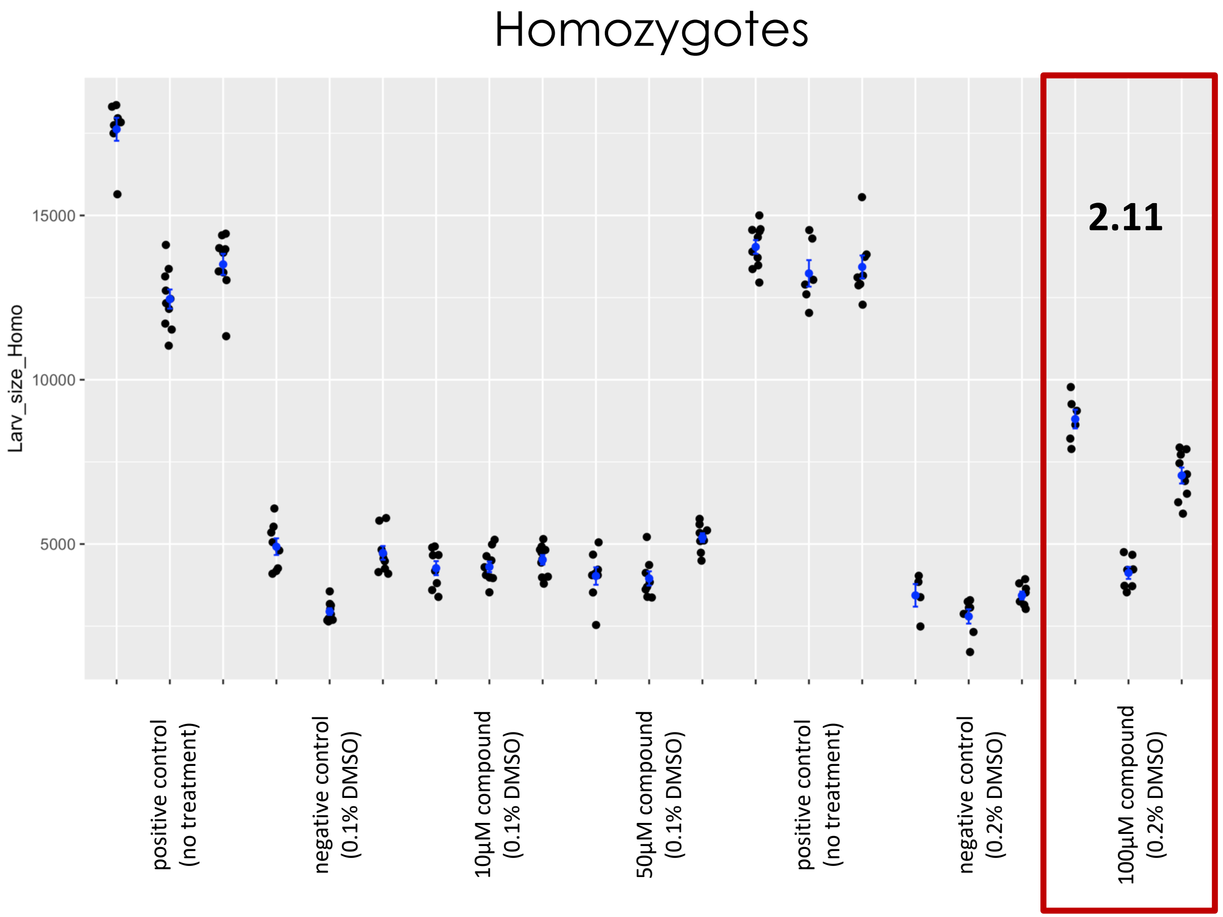


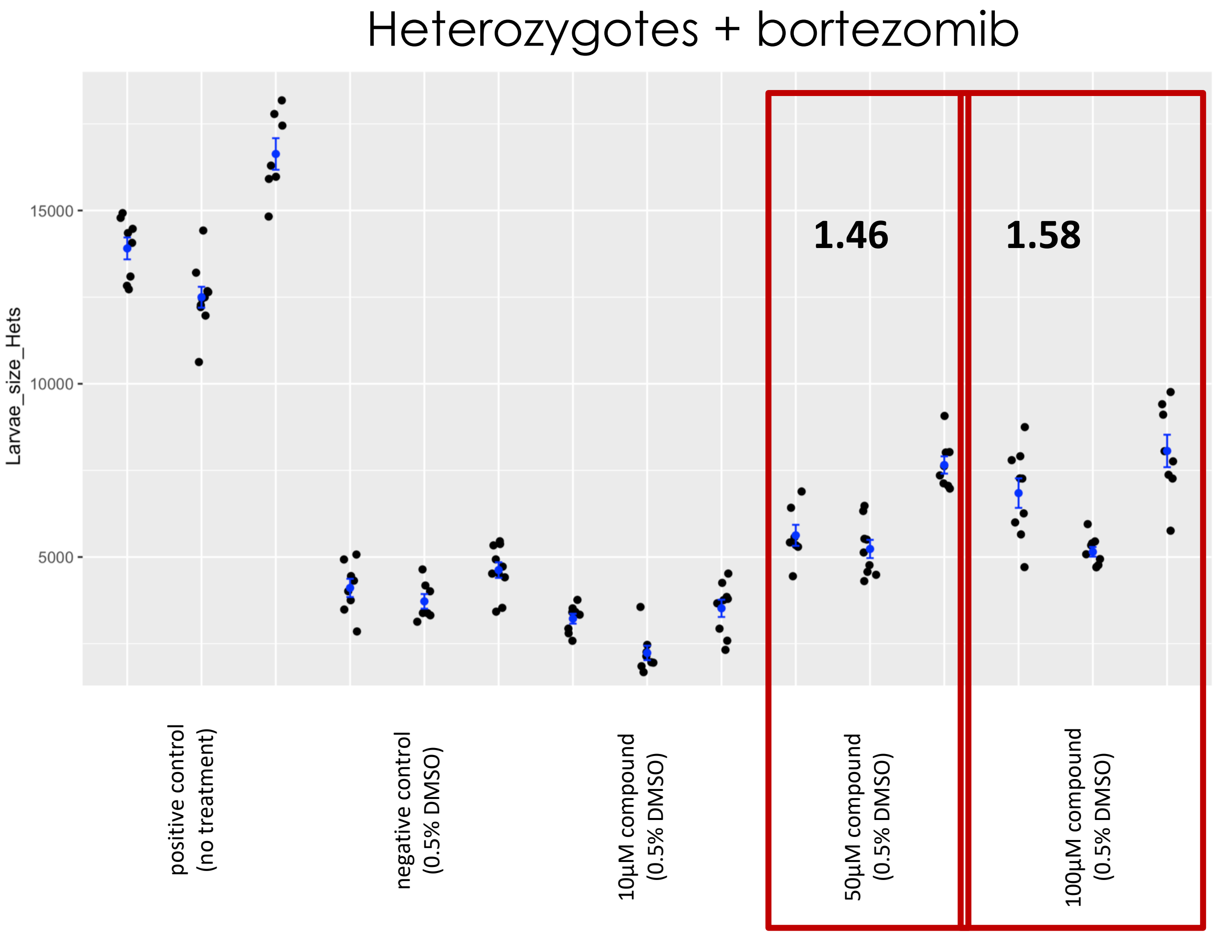

### Fly Microsource Retest – LUMEFANTRINE

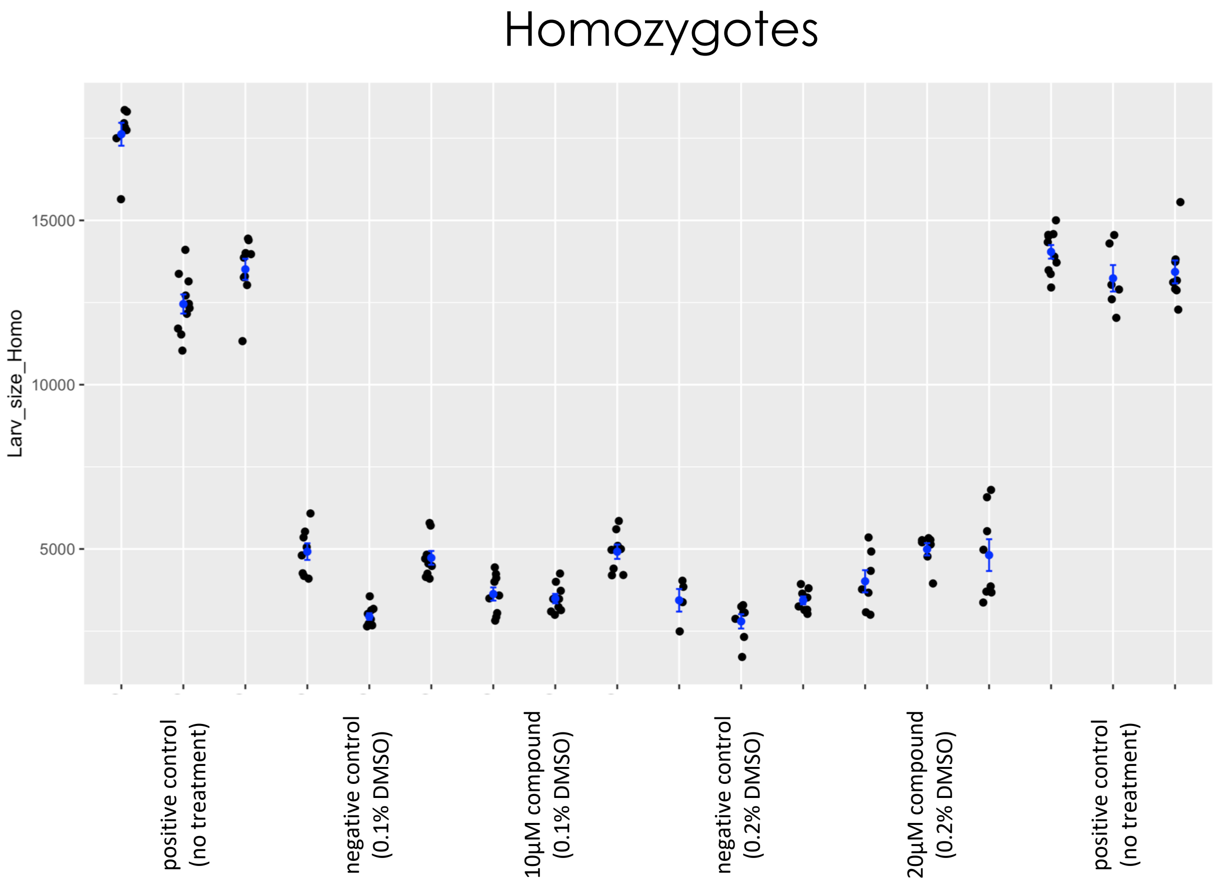


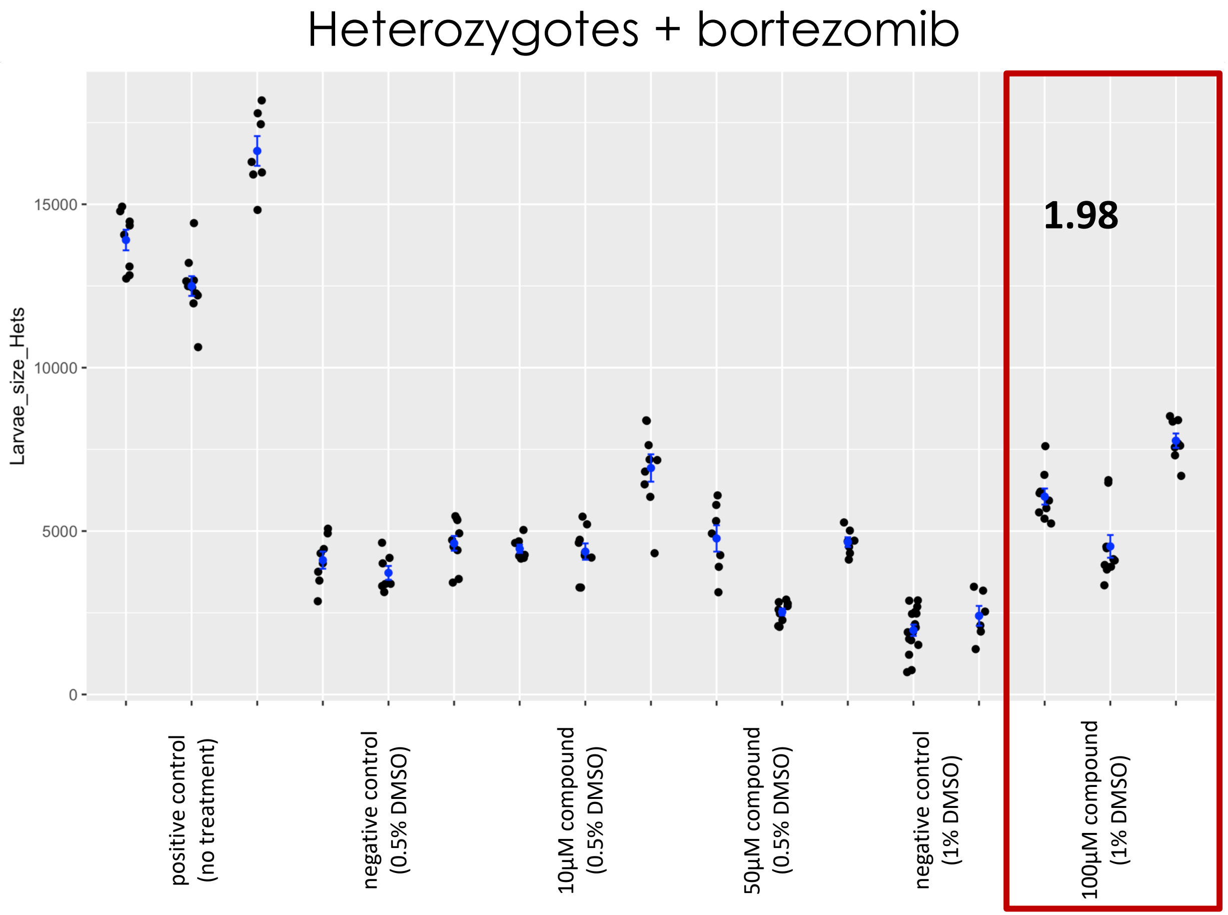

### Fly Microsource Retest – NICERGOLINE

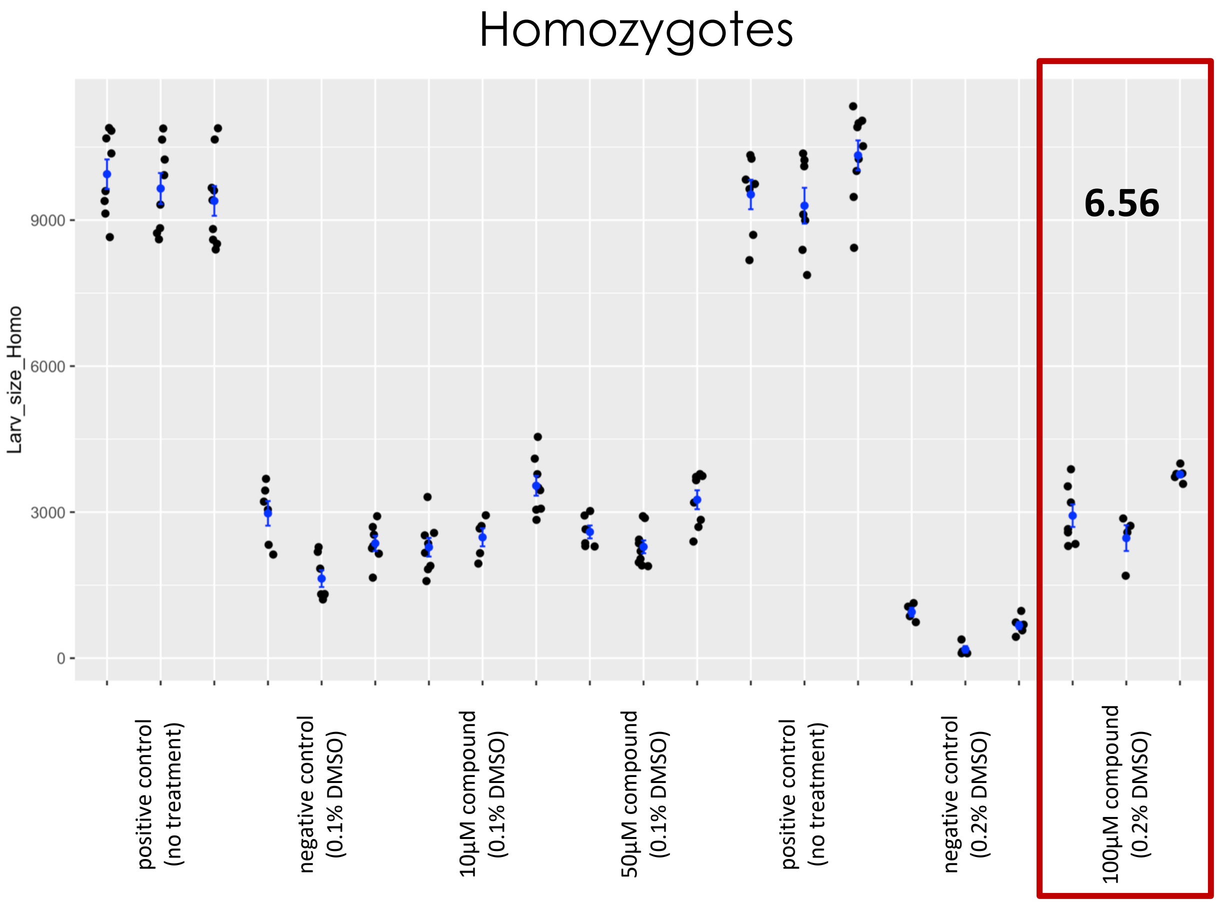


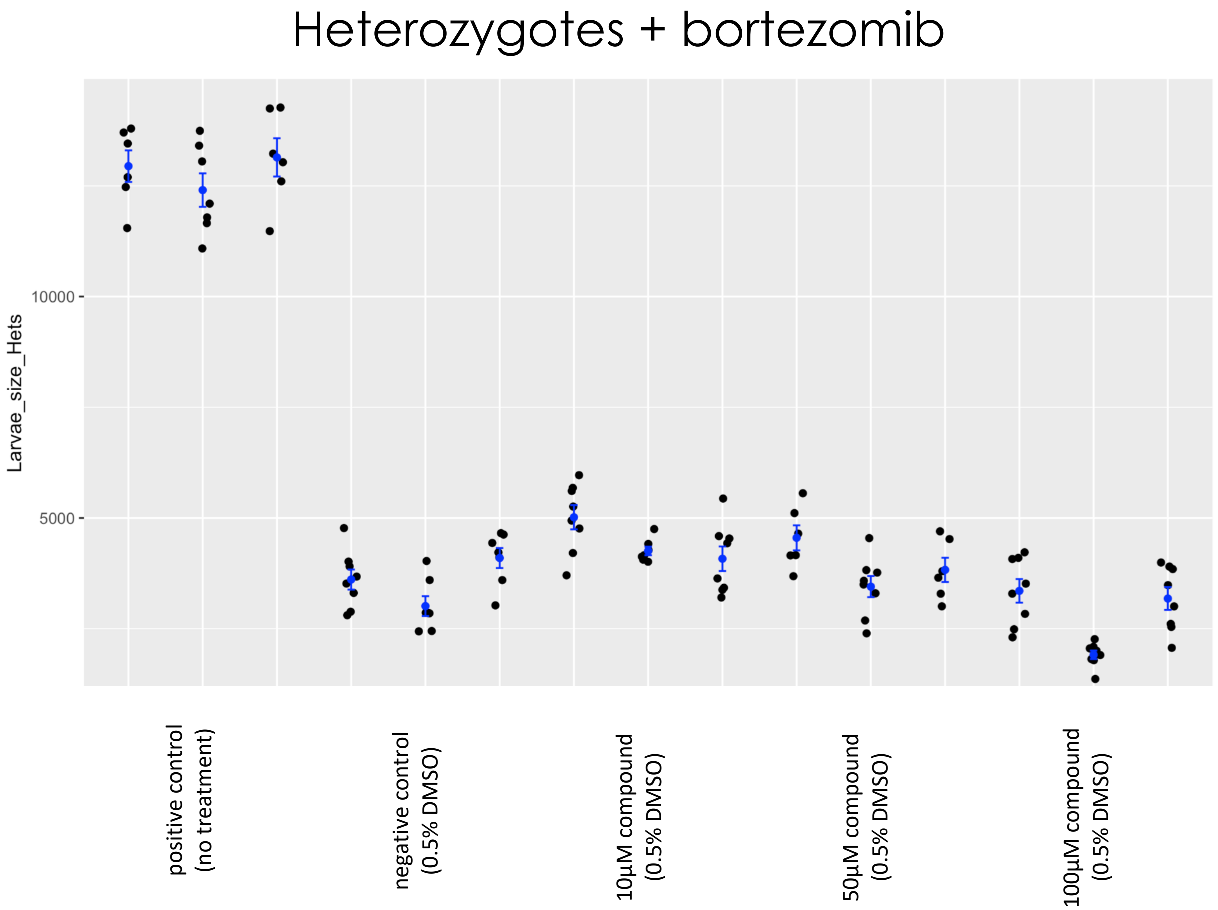

### Fly Microsource Retest – PIMETHIXENE MALEATE

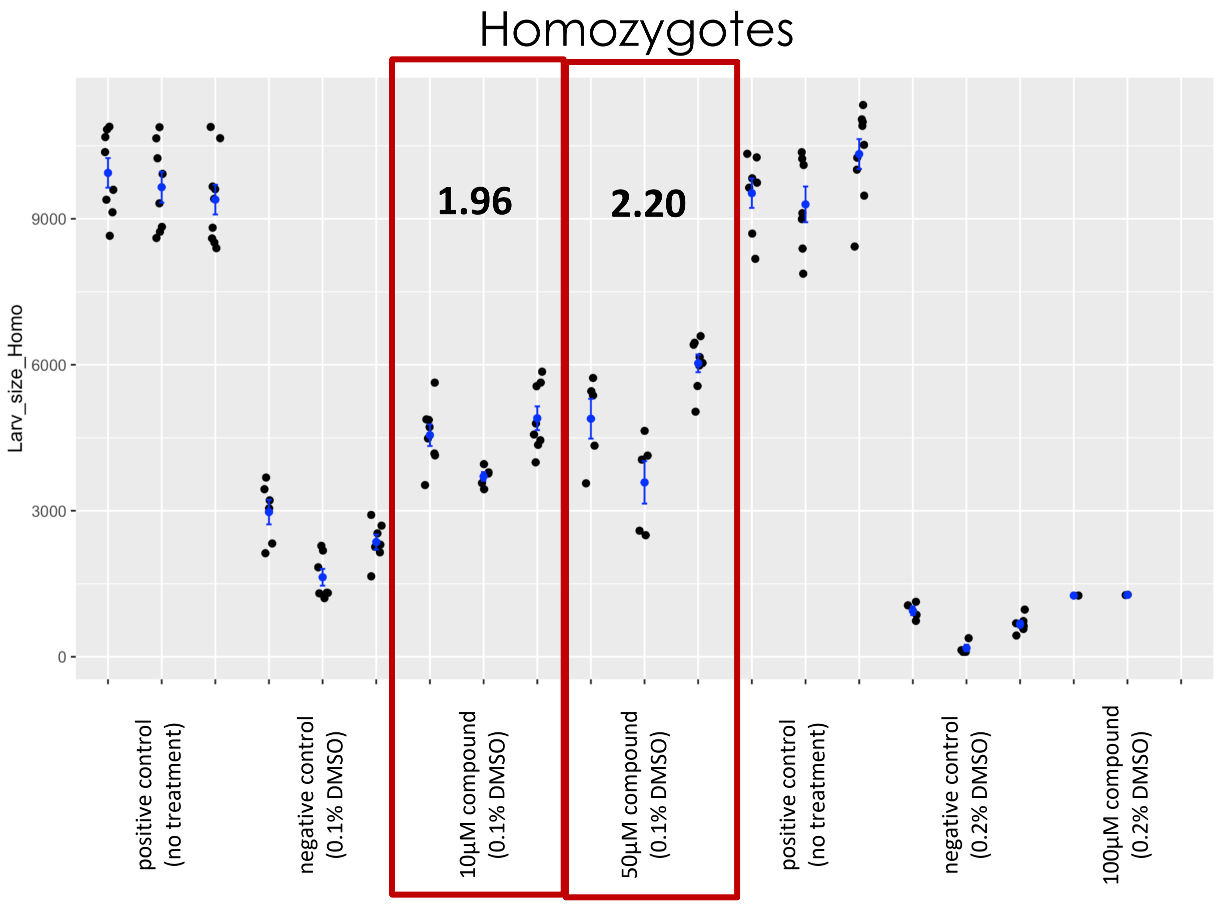


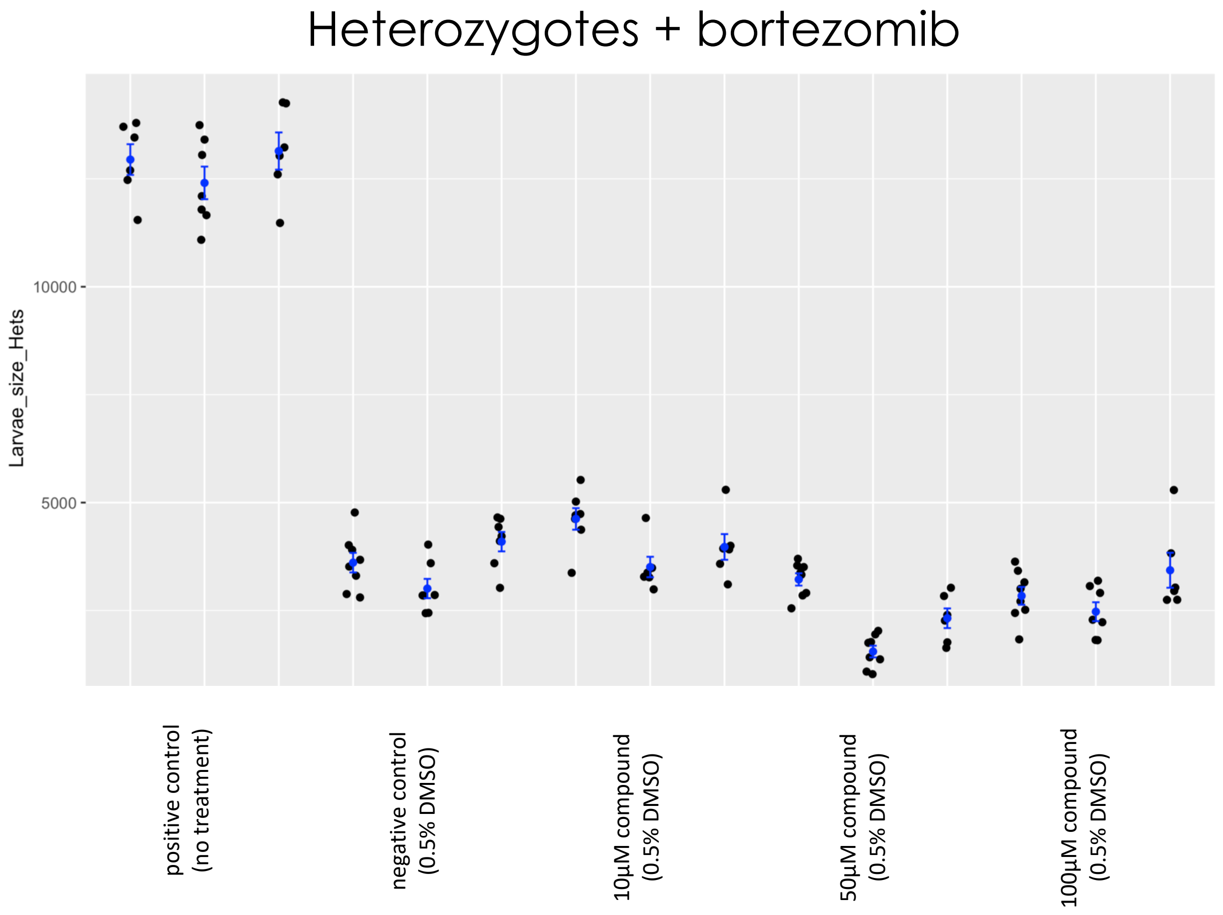

### Fly Microsource Retest – PIPERINE

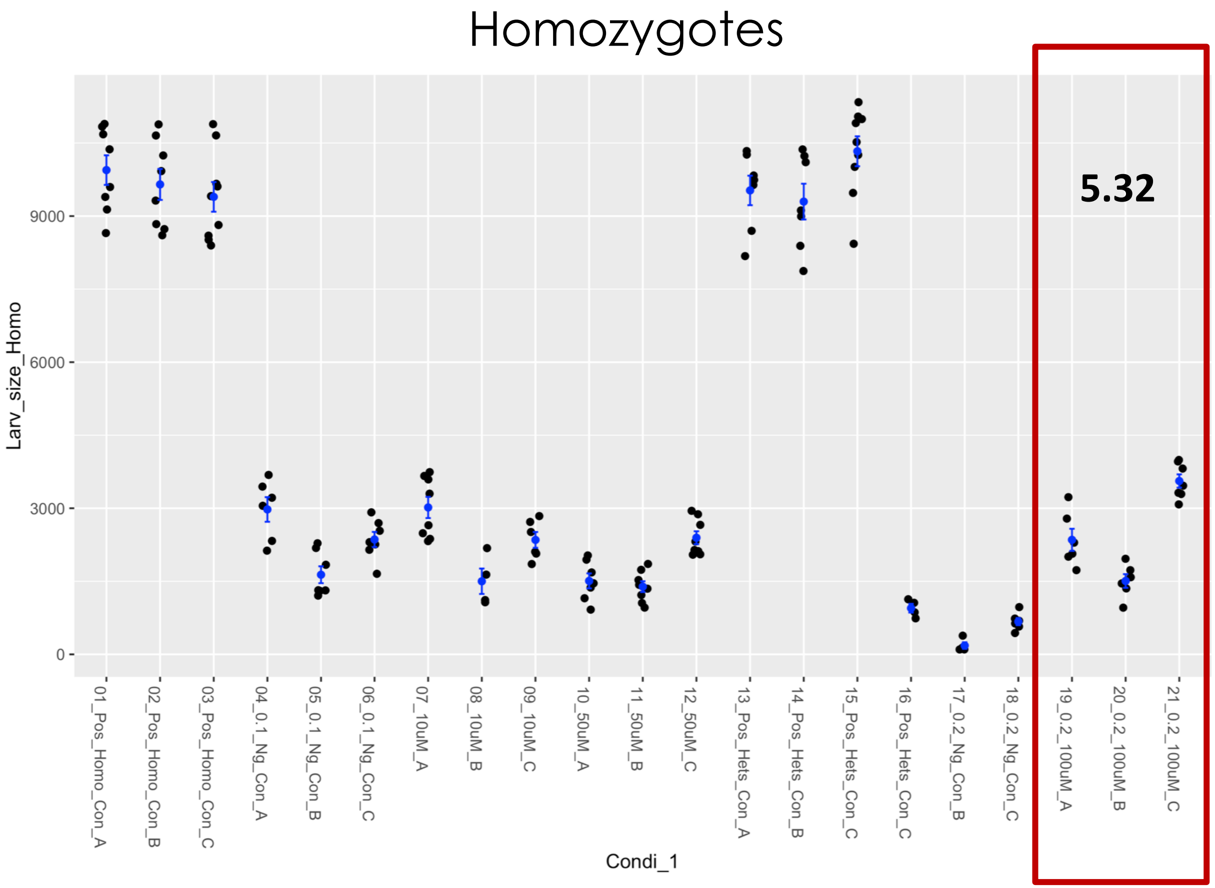


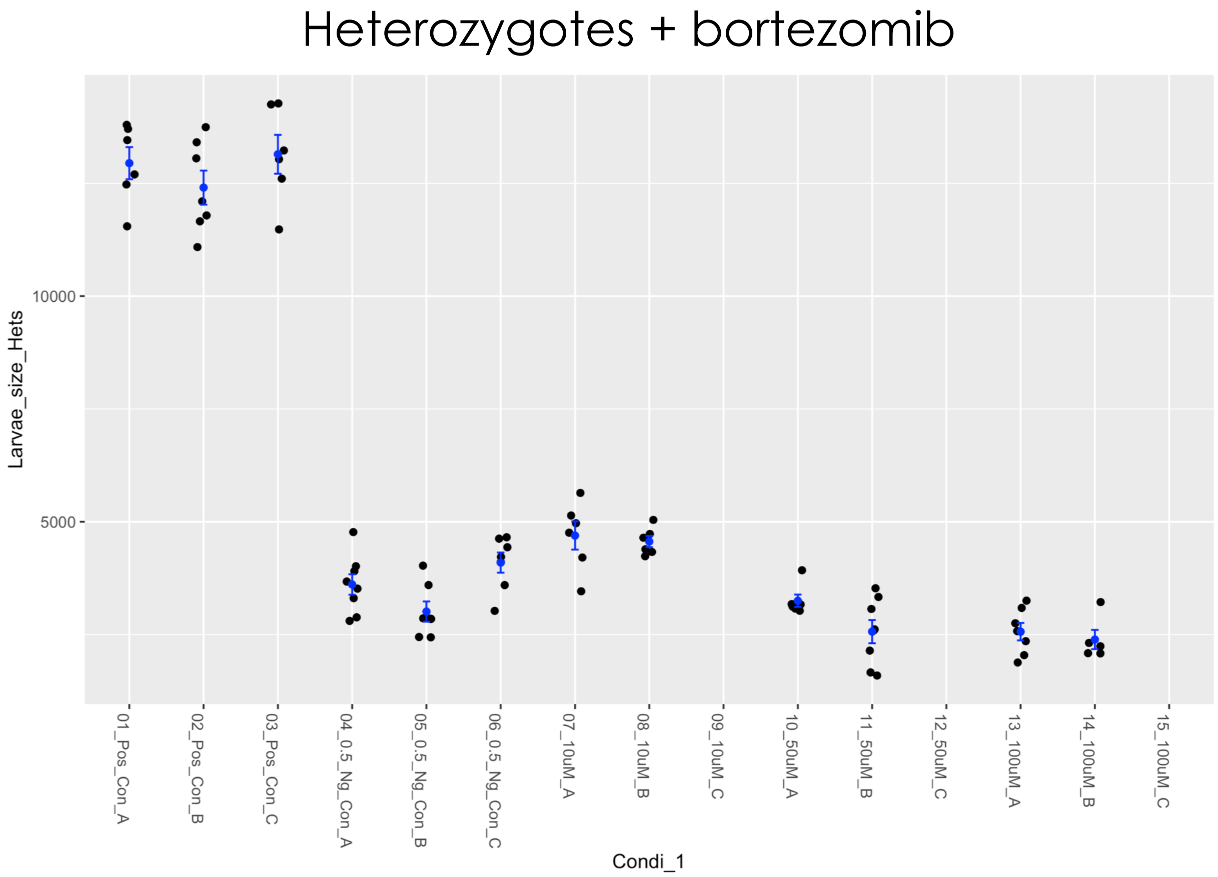

### Fly Microsource Retest – PRAXADINE HYDROCHLORIDE

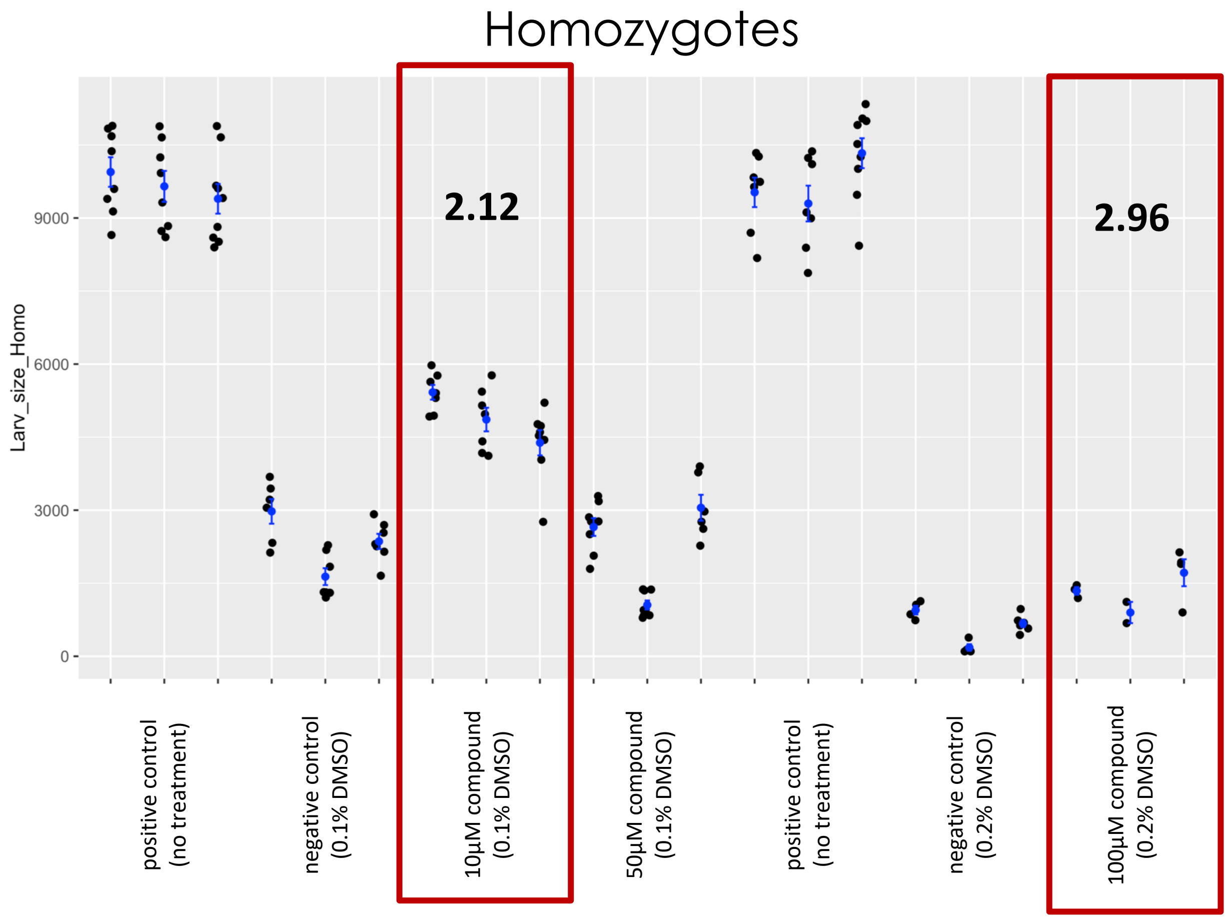


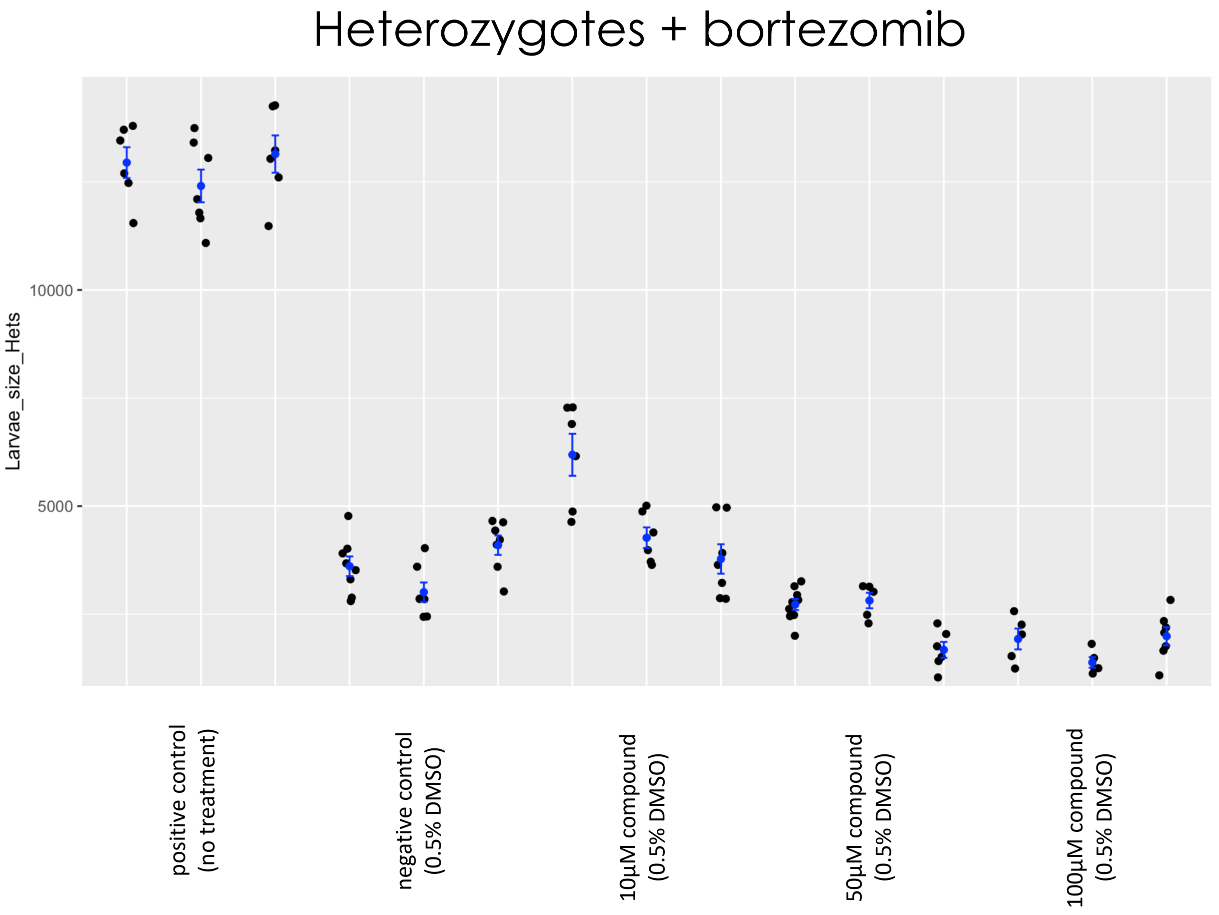

### Fly Microsource Retest – PROMAZINE HYDROCHLORIDE

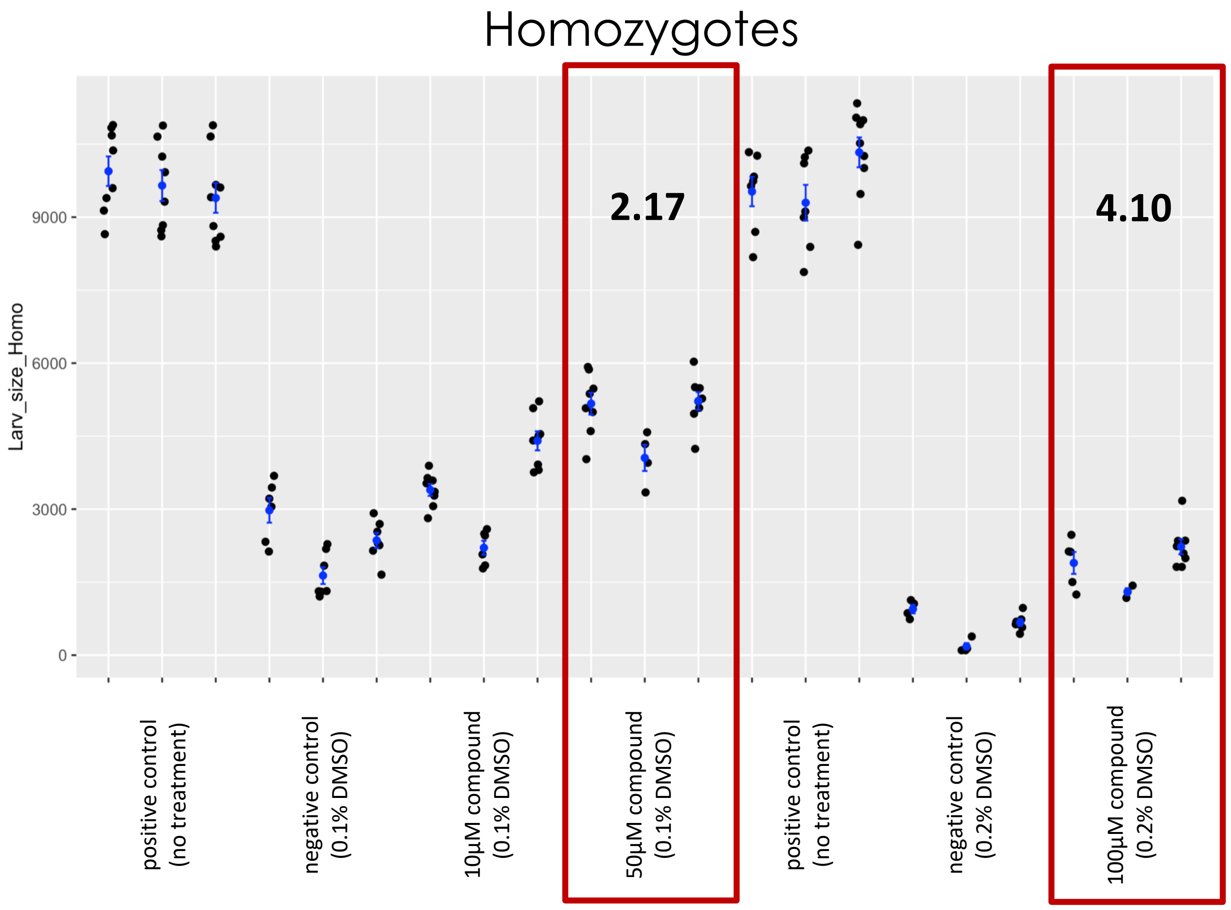


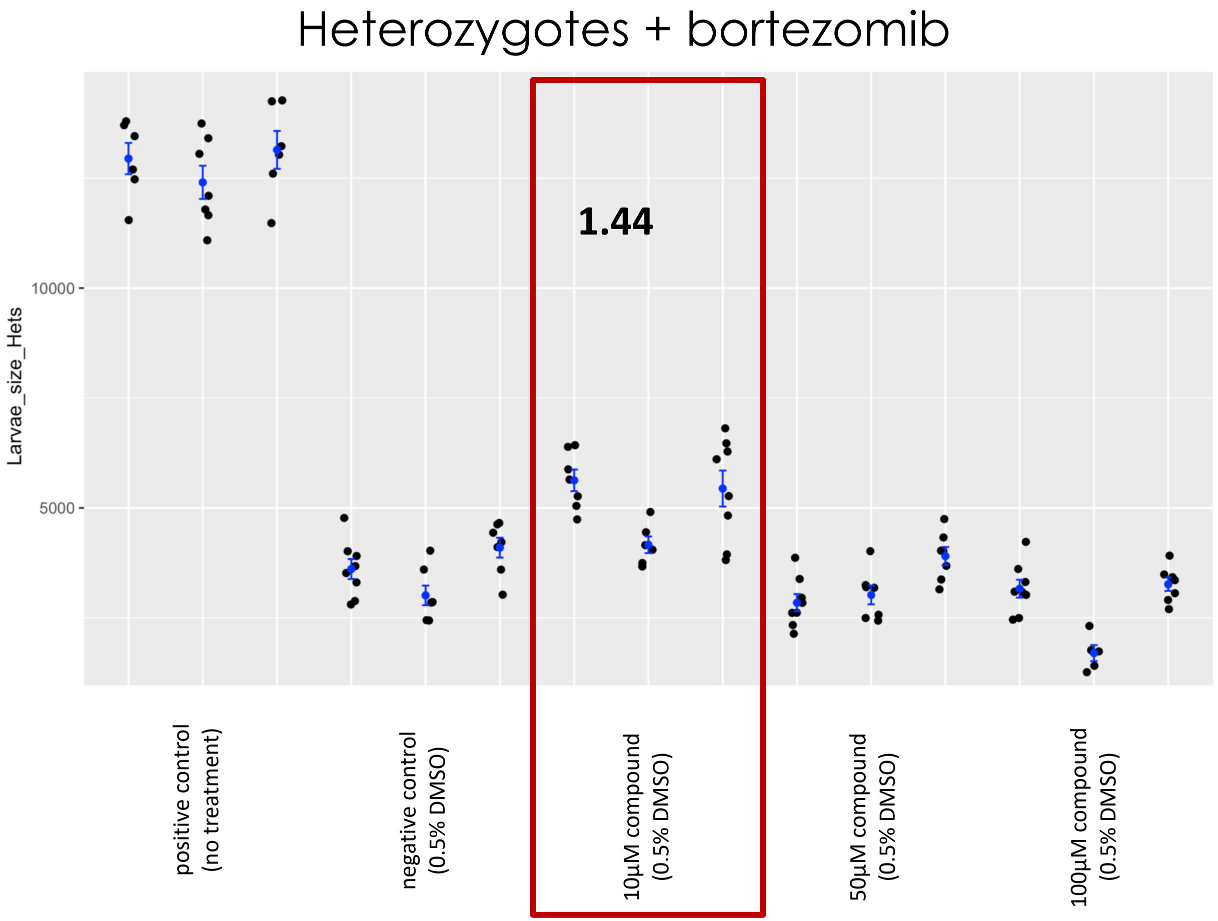

### Fly Microsource Retest – PYRIMETHAMINE

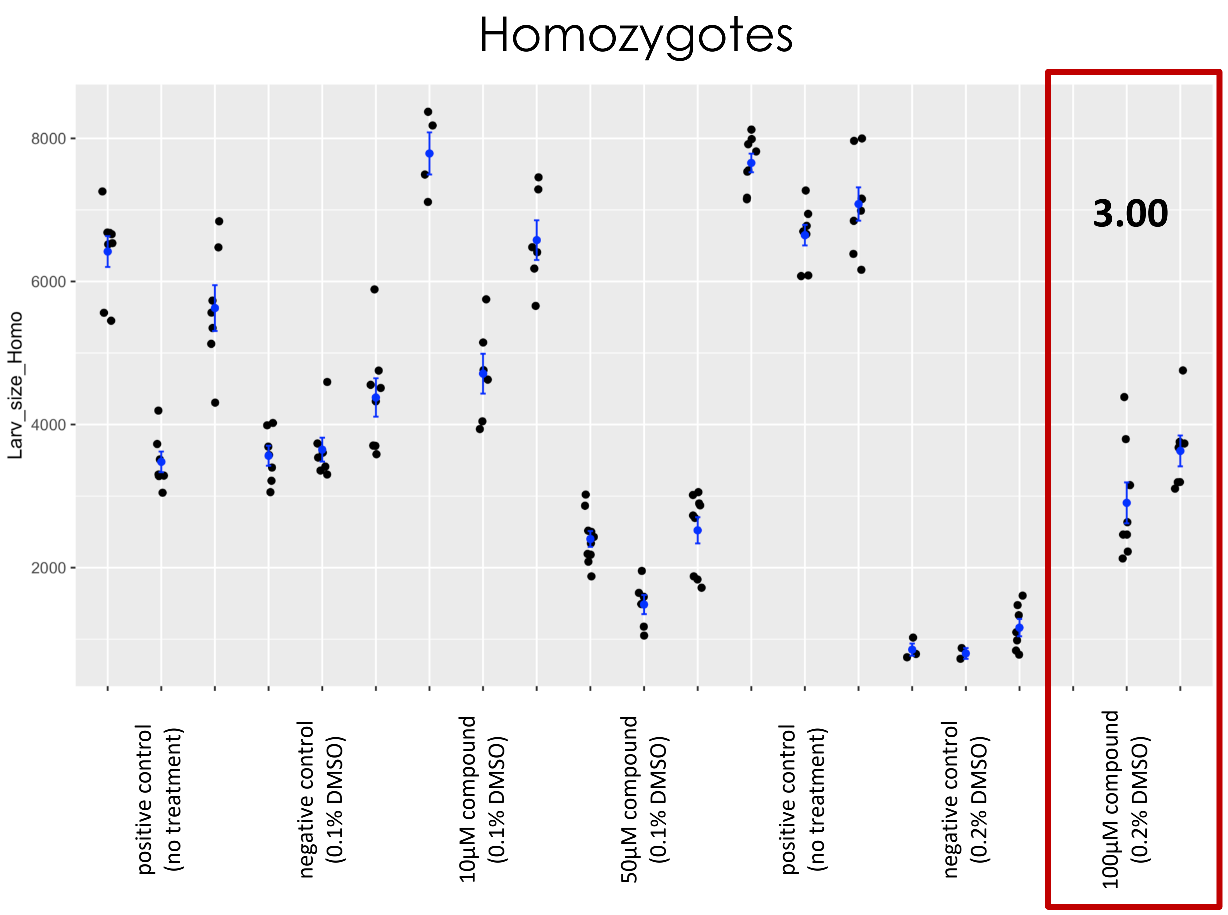


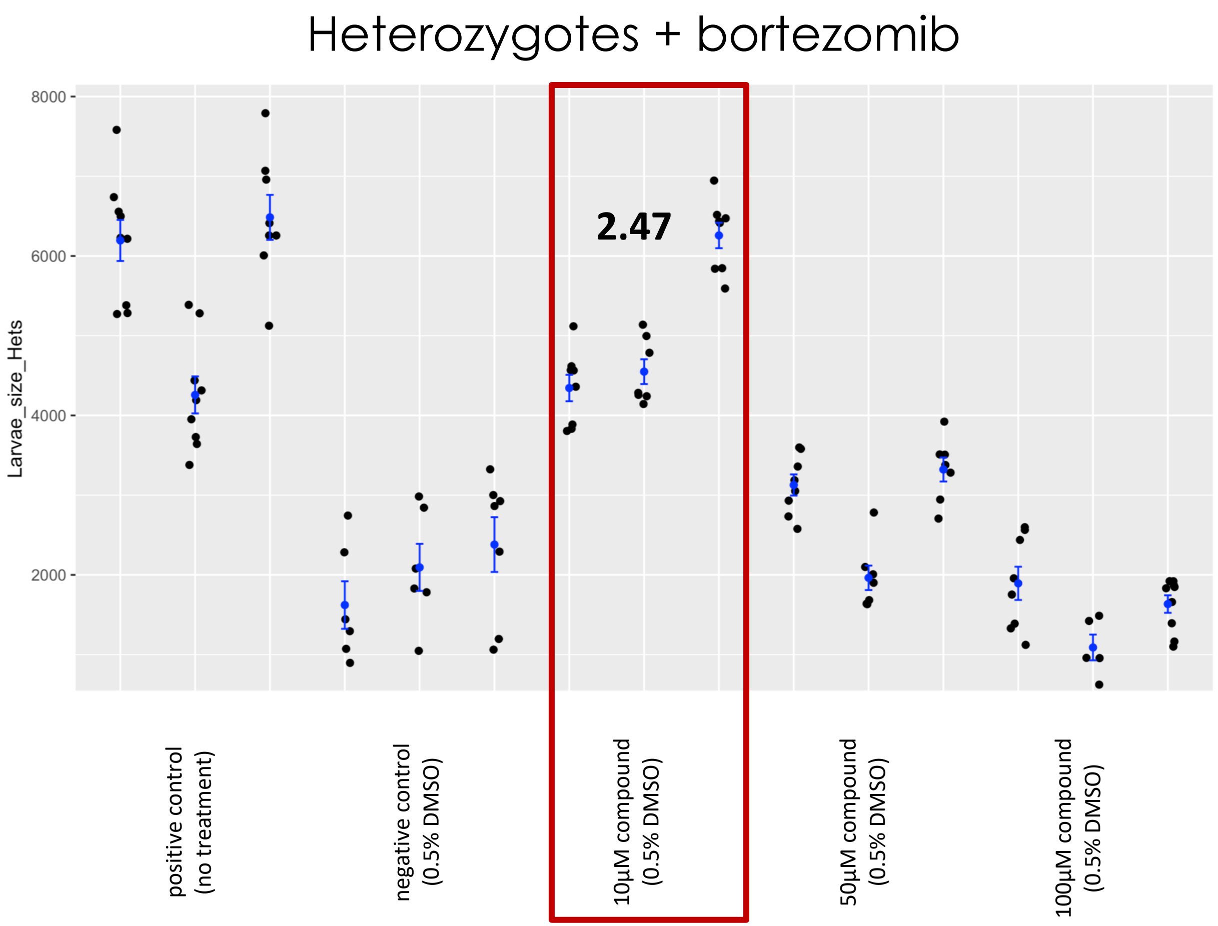

### Fly Microsource Retest – QUINIZARIN

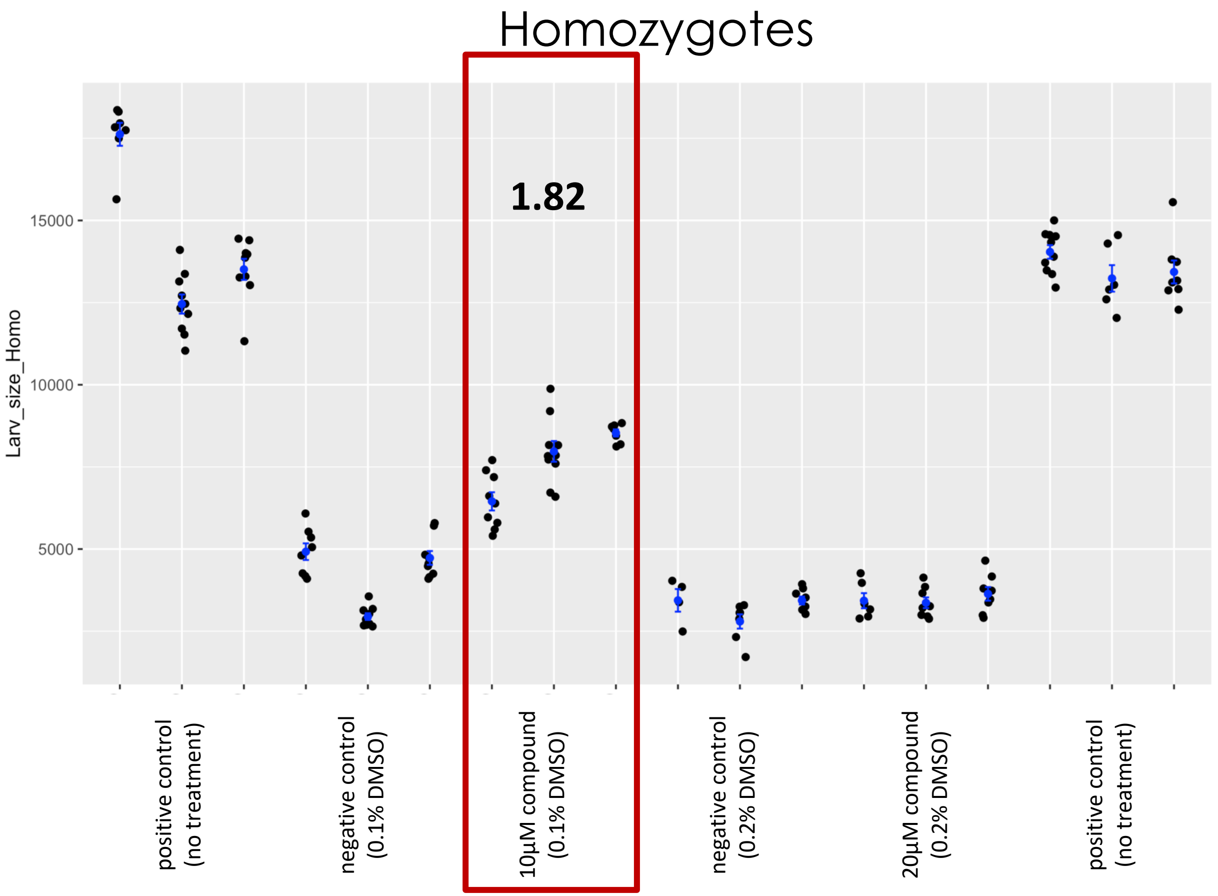


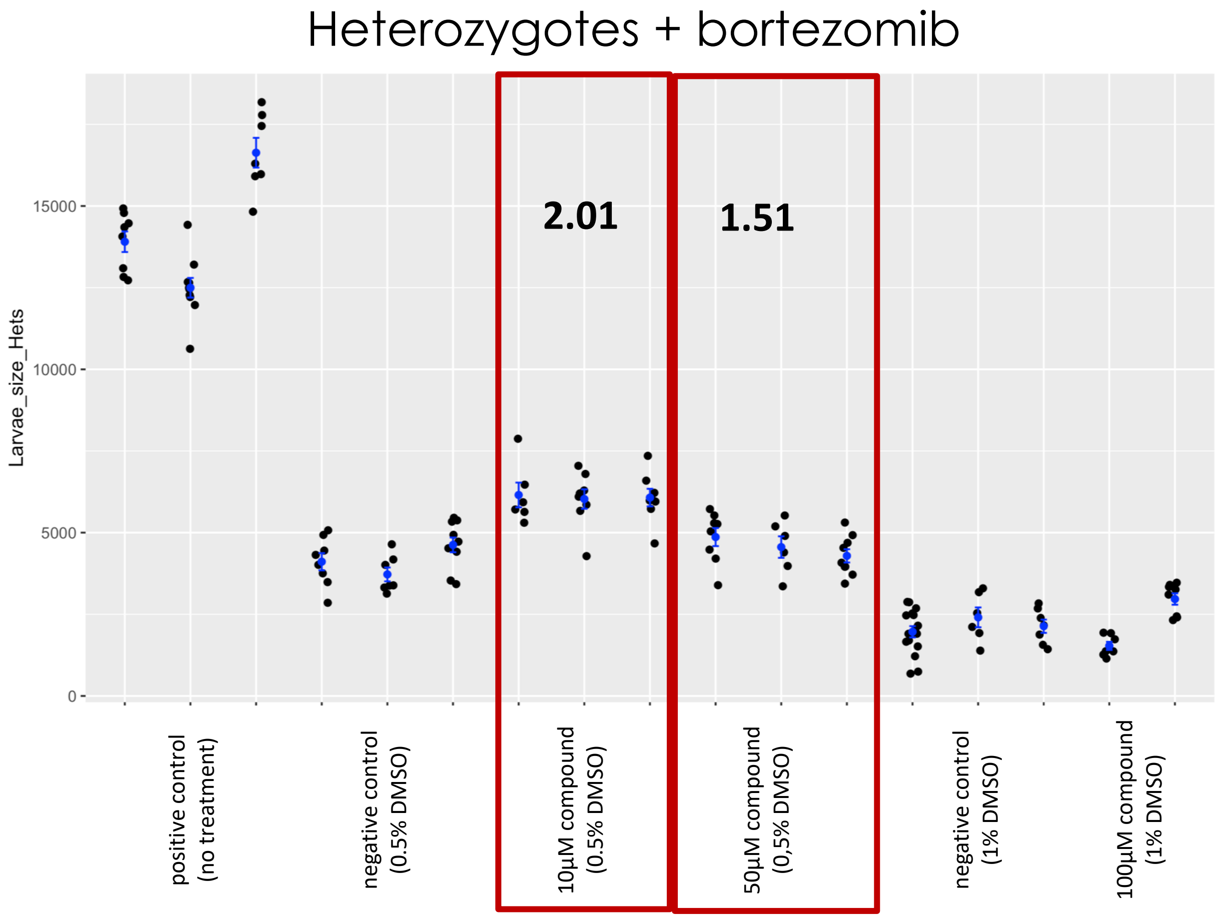

### Fly Microsource Retest – RHETSININE

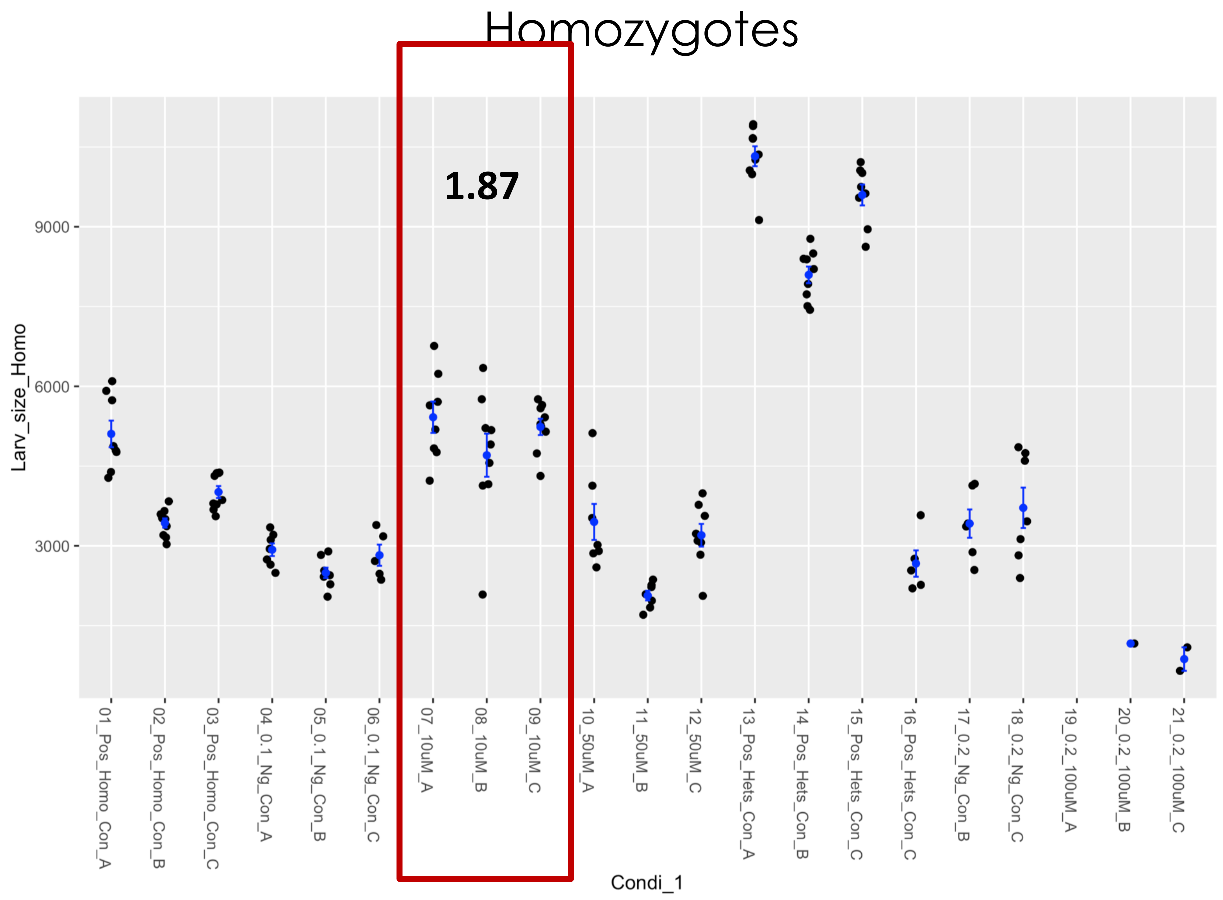


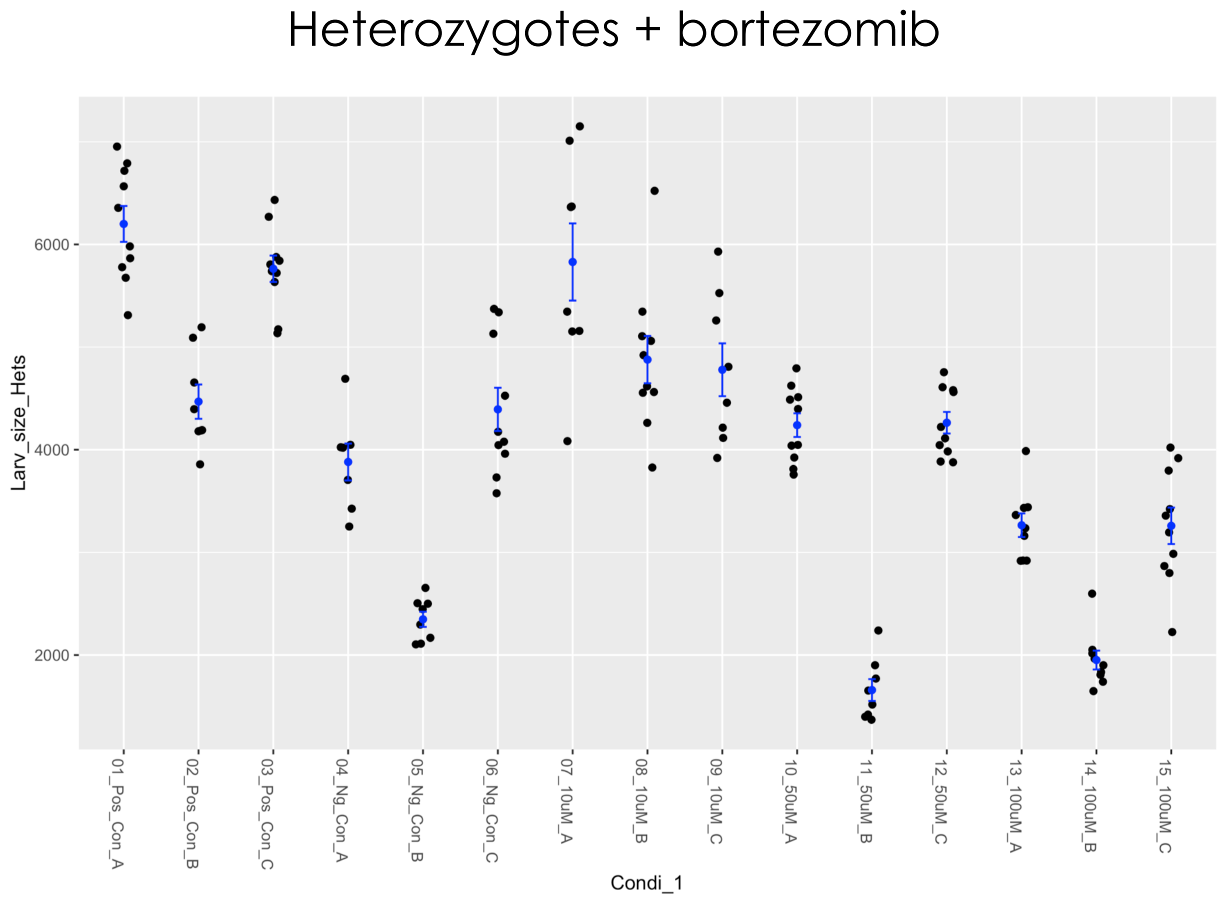

### Fly GSF tool compound test – SULFORAPHANE

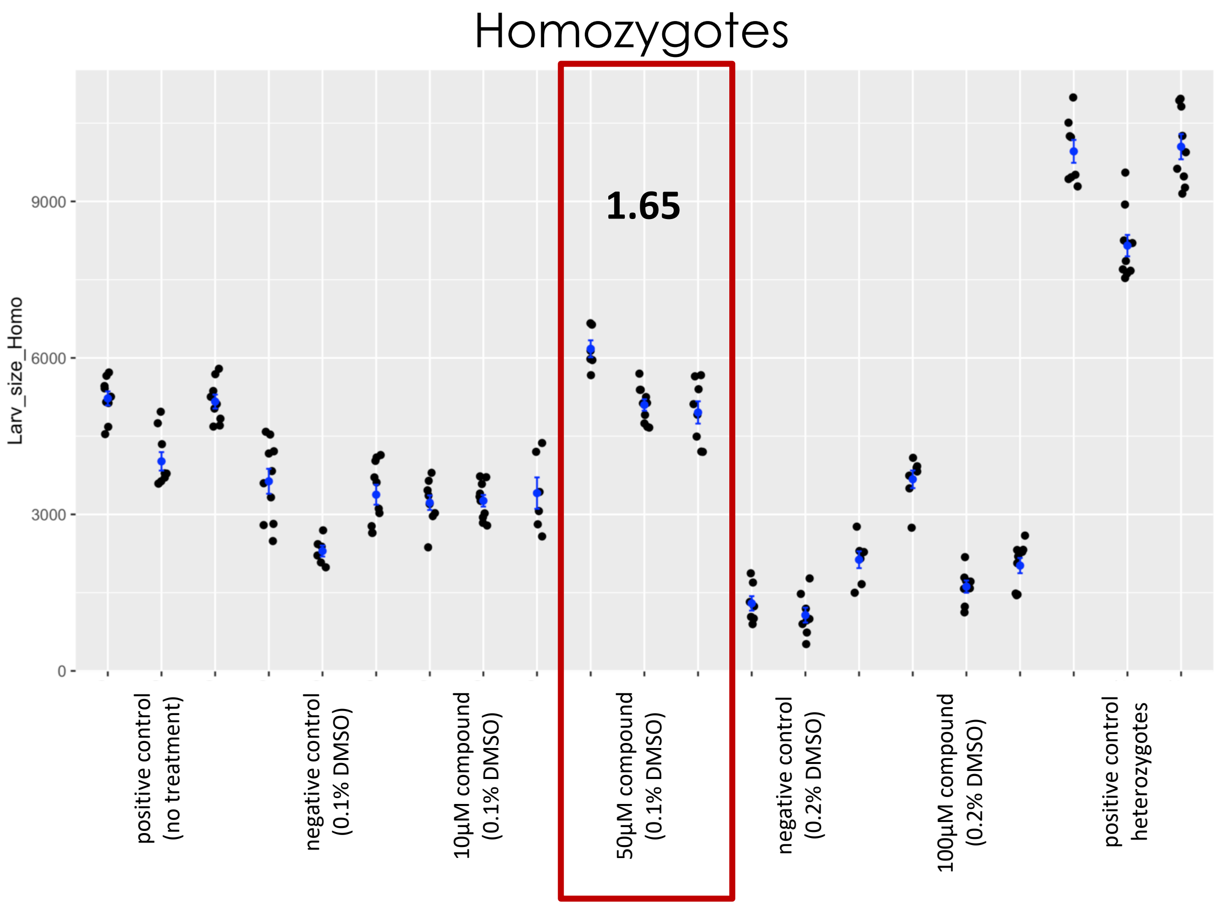


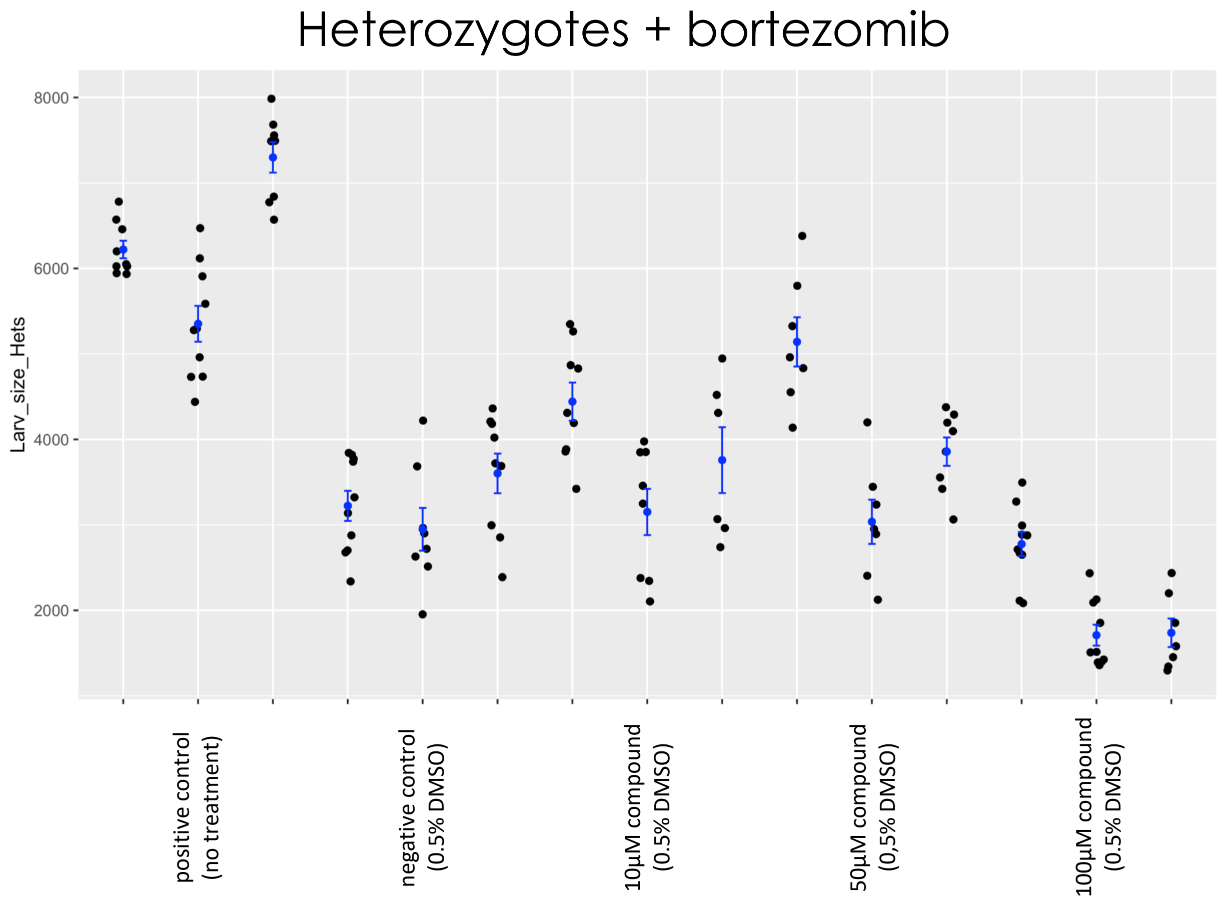

### Fly Microsource retest – TANNIC ACID

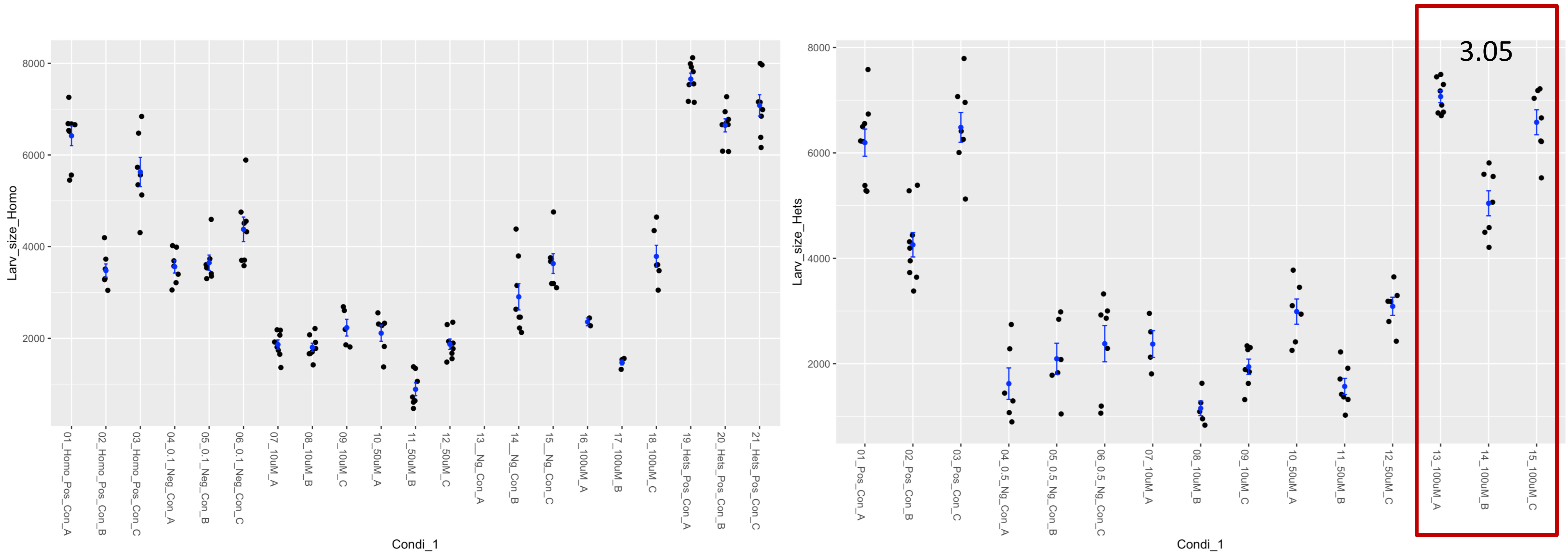

### Fly Microsource Retest – THIOGUANOSINE

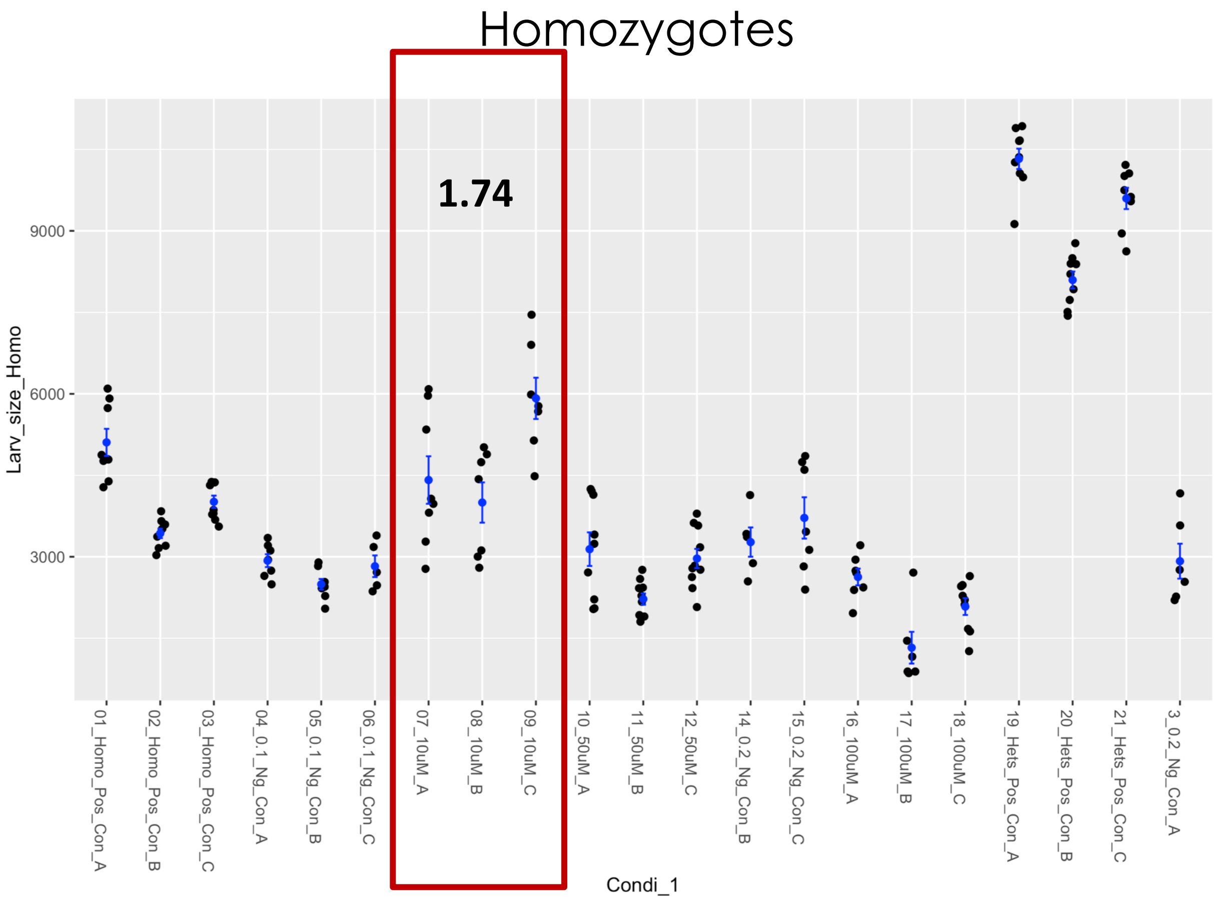


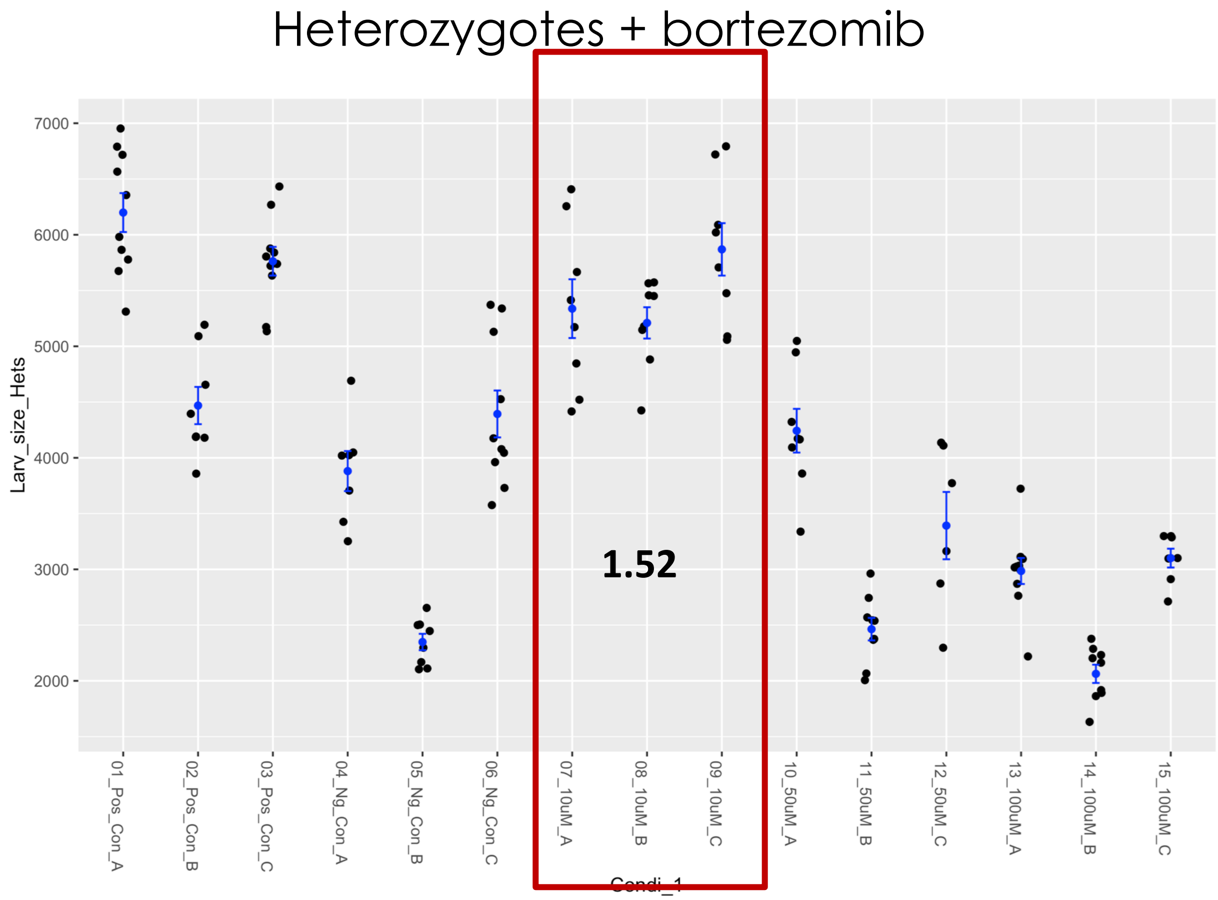

### Fly Microsource retest – TROLOX

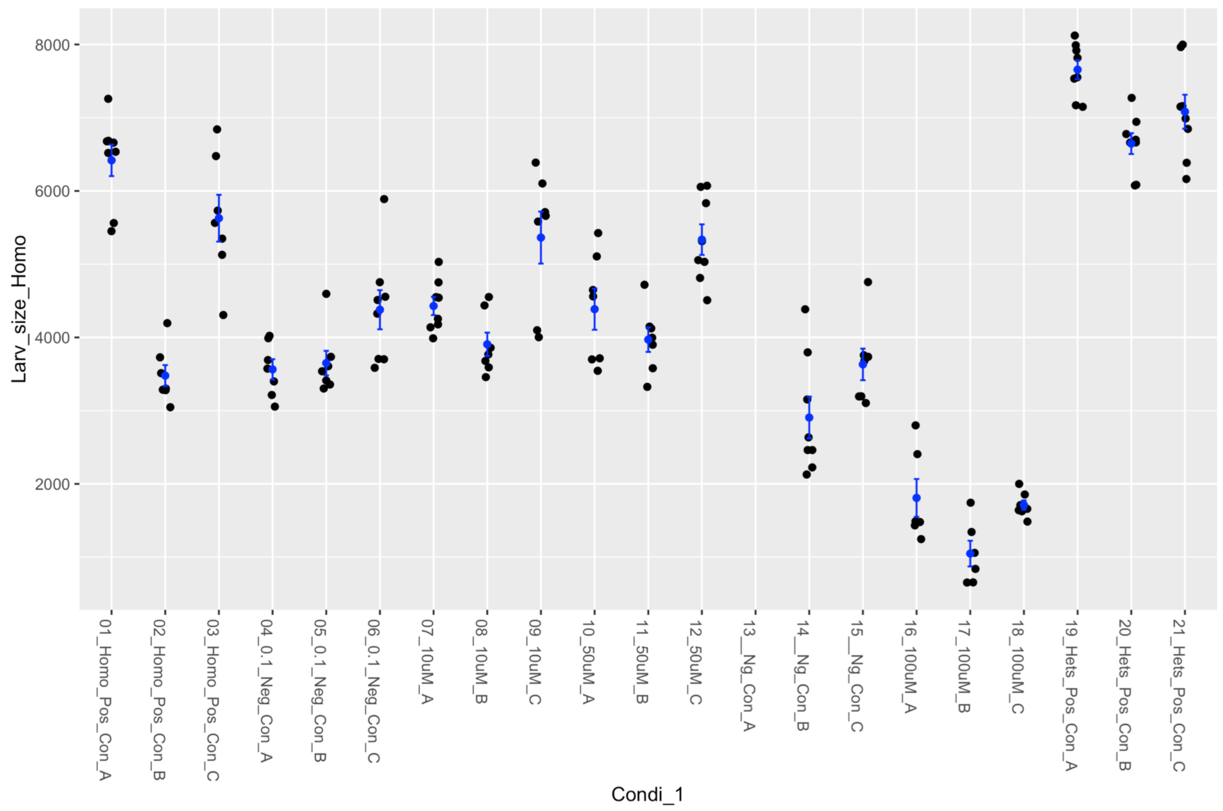


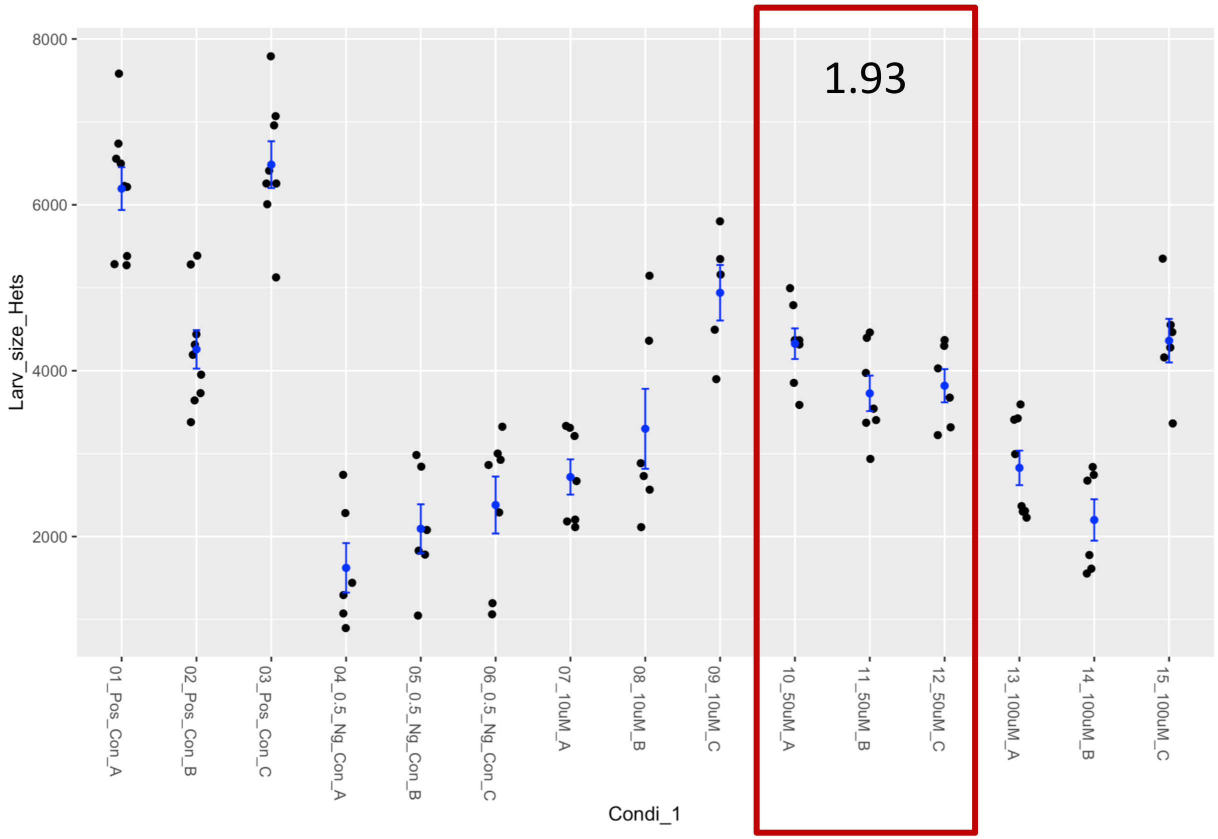

| Supplier ID | Structure |
| --- | --- |
| 4141722 | 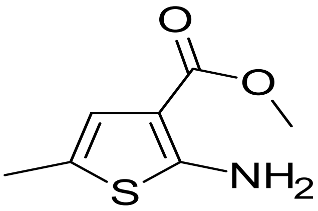 |
| 9223573 | 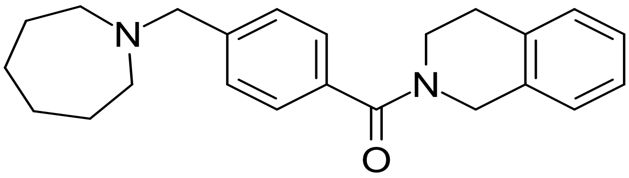 |
| 9268787 | 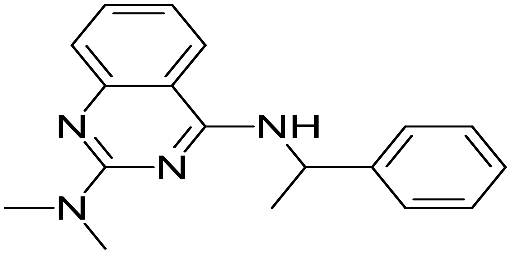 |
| 9283120 | 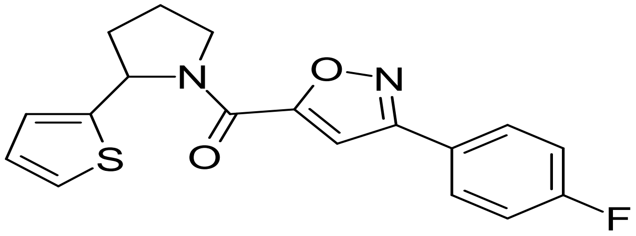 |
| 9313992 | 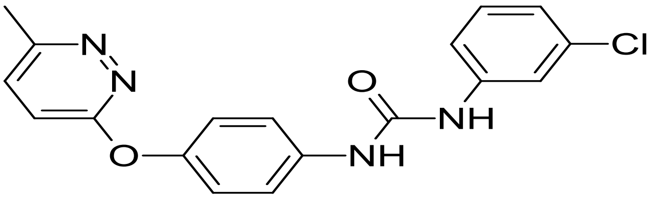 |

| Supplier ID | Structure |
|---|---|
| 4141722 | 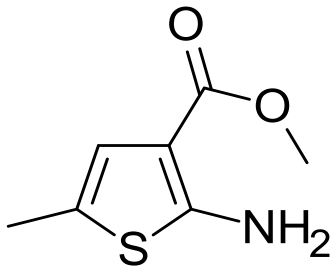 |
| 83371714 | 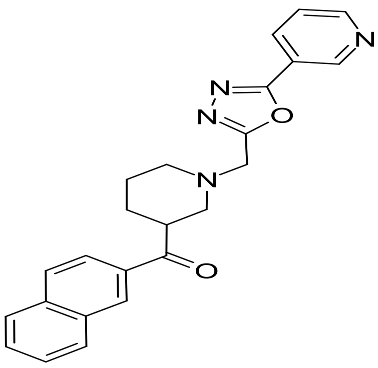 |
| 9193234 | 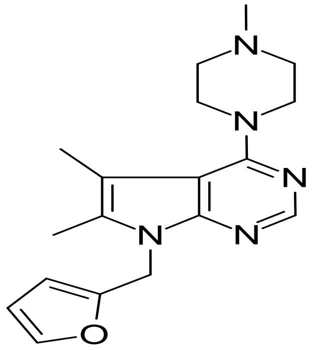 |
| 9223573 | 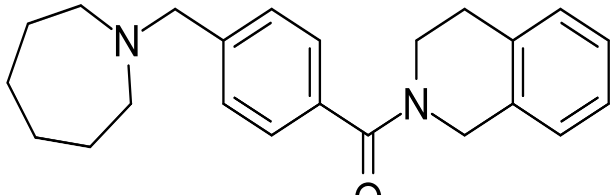 |

9243480

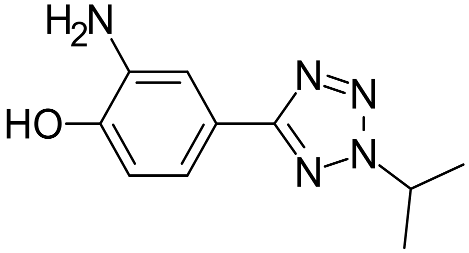


9253438

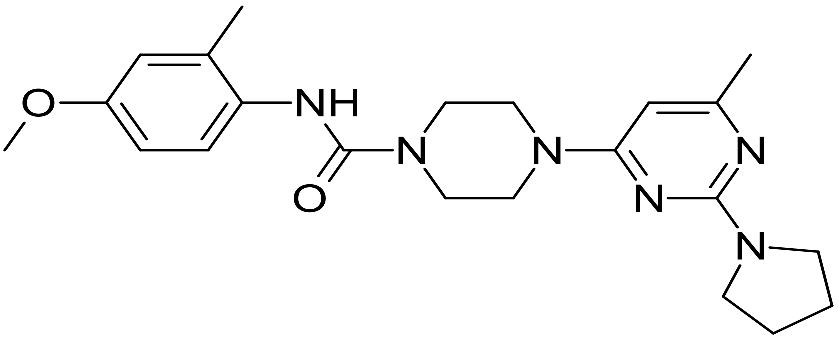


9258723

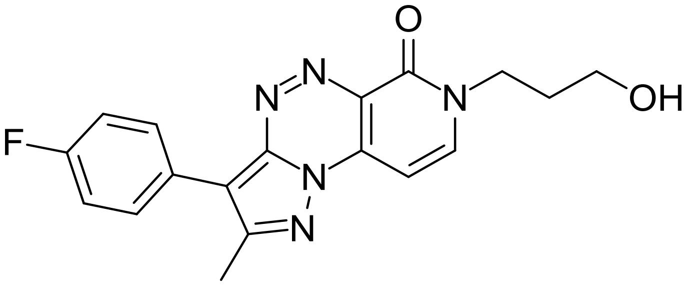


9266225

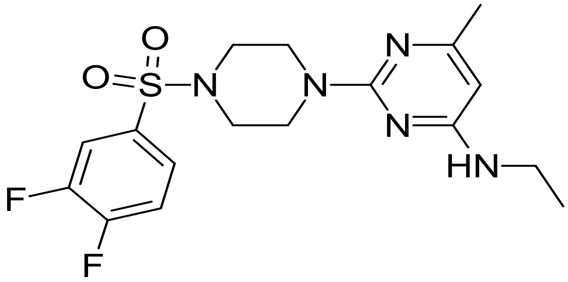

9266617

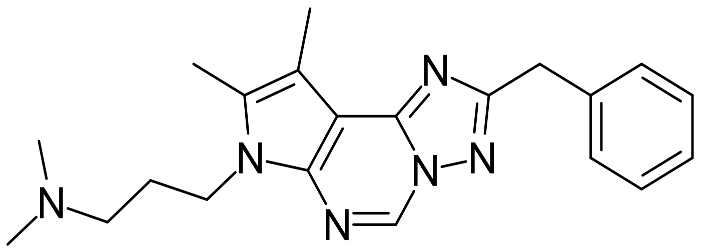


9268787

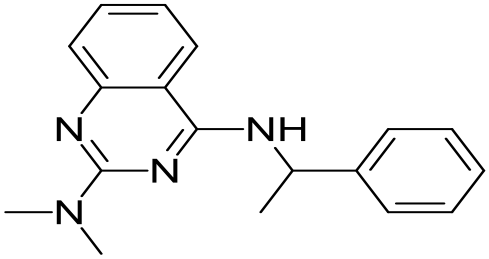


9283120

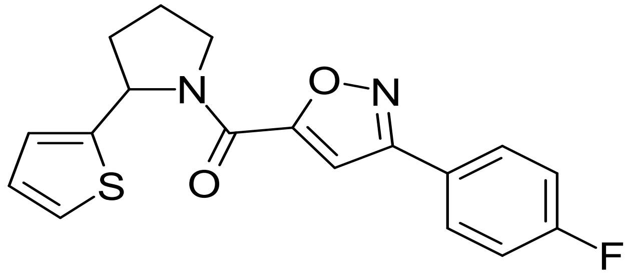


9313992

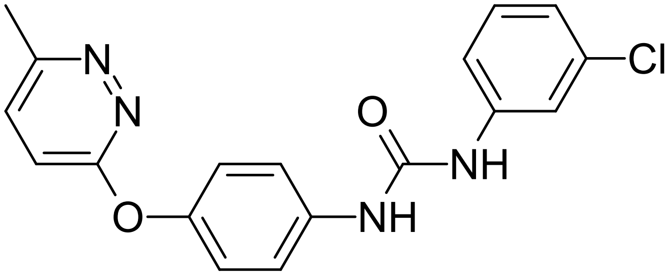

| Supplier ID | Structure |
| --- | --- |
| 4141722 | 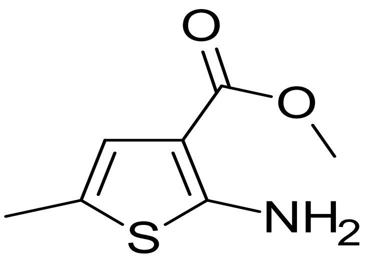 |
| 9223301 | 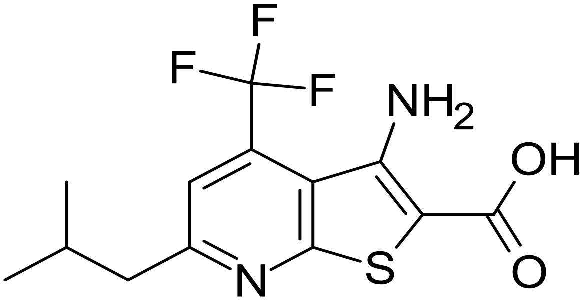 |
| 9223573 | 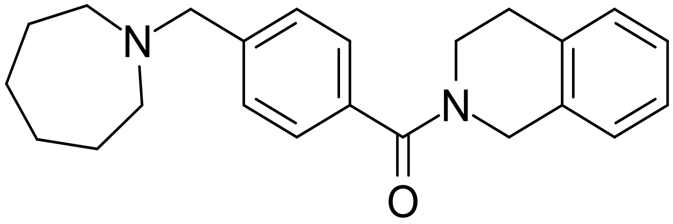 |
| 9236078 | 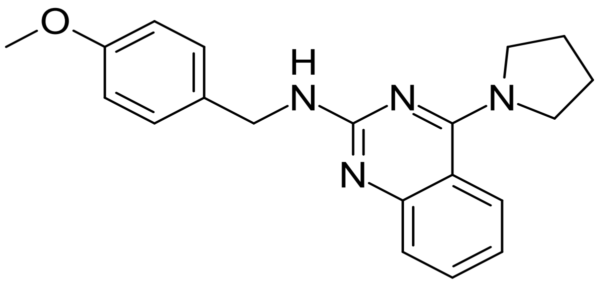 |

9256157

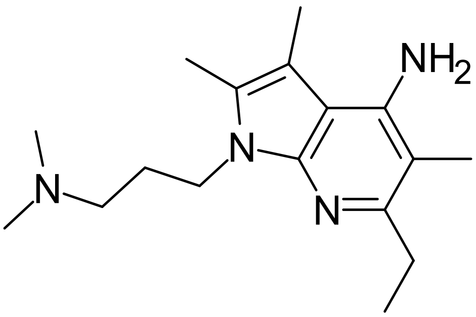

9257006

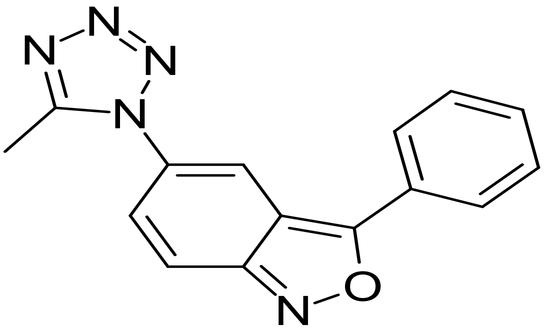

9260754

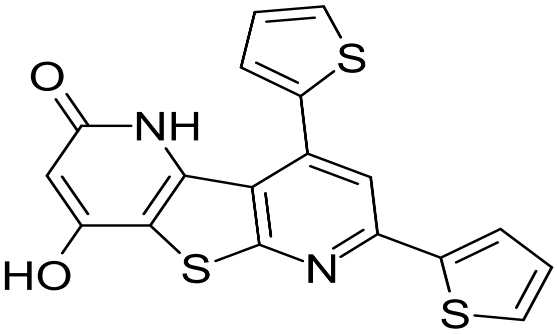

9268787

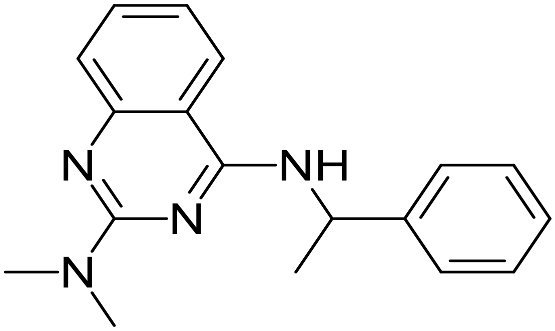

9283120

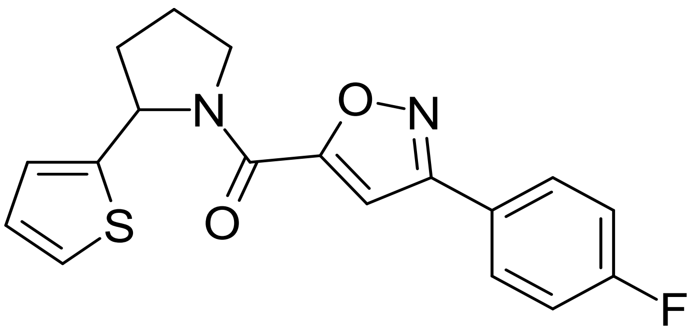


9313992

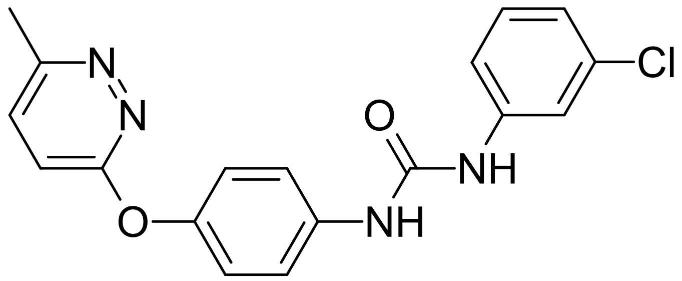
